# Comprehensive analysis of next-generation sequencing data in COVID-19 and its secondary complications

**DOI:** 10.1101/2022.02.03.478930

**Authors:** Muttanagouda Giriyappagoudar, Basavaraj Vastrad, Rajeshwari Horakeri, Chanabasayya Vastrad

**Affiliations:** Department of Radiation Oncology, Karnataka Institute of Medical Sciences (KIMS), Hubballi, Karnataka 580022, India; Department of Pharmaceutical Chemistry, K.L.E. Socitey’s College of Pharmacy, Gadag, Karnataka 582101, India; Department of Computer science, Govt First Grade College, Hubballi, Karnataka 580032, India; Biostatistics and Bioinformatics, Chanabasava Nilaya, Bharthinagar, Dharwad 580001, Karanataka, India

**Keywords:** COVID-19, differentially expressed genes, next-generation sequencing data analysis, hub genes, protein-protein interaction network

## Abstract

The ongoing pandemic of coronavirus disease 2019 (COVID-19) has made a serious public health threat globally. To discover key molecular changes in COVID-19 and its secondary complications, we analyzed next-generation sequencing (NGS) data of COVID-19. NGS data (GSE163151) was screened and downloaded from the Gene Expression Omnibus database (GEO). Differentially expressed genes (DEGs) were identified in the present study, using DESeq2 package in R programming software. Gene ontology (GO) and pathway enrichment analysis were performed, and the protein-protein interaction (PPI) network, module analysis, miRNA-hub gene regulatory network and TF-hub gene regulatory network were established. Subsequently, receiver operating characteristic curve (ROC) analysis was used to validate the diagonostics valuesof the hub genes. Firstly, 954 DEGs (477 up regulated and 477 down regulated) were identified from the four NGS dataset. GO enrichment analysis revealed enrichment of DEGs in genes related to the immune system process and multicellular organismal process, and REACTOME pathway enrichment analysis showed enrichment of DEGs in the immune system and formation of the cornified envelope. Hub genes were identified from the PPI network, module analysis, miRNA-hub gene regulatory network and TF-hub gene regulatory network. Furthermore, the ROC analysis indicate that COVID-19 and its secondary complications with following hub genes, namely, RPL10, FYN, FLNA, EEF1A1, UBA52, BMI1, ACTN2, CRMP1, TRIM42 and PTCH1, had good diagnostics values. This study identified several genes associated with COVID-19 and its secondary complications, which improves our knowledge of the disease mechanism.

## Introduction

Coronavirus Disease 2019 (COVID-19) also known as severe acute respiratory syndrome coronavirus-2 (SARS-CoV-2) and was first reported in Wuhan city, Hubei province, China in December 2019 [1]. Within a short period of time, COVID-19 has raced around the world and World Health Organization (WHO) officially declared the COVID-19 epidemic as a health emergency [2]. The number of infected cases and deaths due to COVID-19 is rising severely and on 17th December 2021, there are more than 273,327,260 confirmed cases with over 5,355,724 deaths across 200 countries (Worldometer 2020).

COVID-19 can kill the tissue of lungs [3], heart [4], central nervous system [5], liver [6] and kidneys [7]. COVID-19 infection is characterized by an increased risk of acute respiratory failure [8], pneumonia [9], acute respiratory distress syndrome [10], acute liver injury [11], acute cardiac injury [12], secondary infection [13], central nervous system (CNS) disorders [14], acute kidney injury [15], septic shock [16], diabetes mellitus [17], hypertension [18], disseminated intravascular coagulation [19], blood clots [20], multisystem inflammatory syndrome in children [21], chronic fatigue [22] and rhabdomyolysis [23]. The higher prevalence and limited treatments of COVID-19 infection lead to substantial public health and global burdens [24]. Therefore, it is necessary to improve our understanding of COVID-19 molecular pathogenesis and to develop better screening methods for COVID-19 infection and its secondary complications.

The COVID-19 infection and its secondary complications are an extremely complicated pathophysiological process and regulated by various genetic factors and pathways [25]. Five novel genetic factor for COVID-19 infection and its secondary complications including APOL1 [26], ACE2 [27], CXCL10 [28], IFNAR2 [29] and MBL2 [30] as potential biomarkers of COVID-19 infection and its secondary complications. Signaling pathway includes JAK-STAT signaling pathway [31], ACE-2\Mas\BDNF signaling pathway [32], IFN-AhR signaling pathway [33] and toll-like receptor 4-mediated inflammatory signaling pathway [34] were responsible for progression of COVID-19 infection and its secondary complications. Recently, next generation sequencing (NGS) technology is widely used in COVID-19 investigation, with the pronounced advantages relying on its ability to together determine the expression data of huge genes one time [35]. Based on the massive NGS datasets information stored in a large number of public databases such as Gene Expression Omnibus (GEO) database and ArrayExpress, the target data could be mined, integrated, and reanalyzed by bioinformatics method, which provide useful clues for the investigation of numerous diseases [36–37].

We downloaded NGS dataset GSE163151 [38], from GEO (http://www.ncbi.nlm.nih.gov/geo/) [39], which contain gene expression data from COVID-19 samples and non COVID-19 samples. We then performed bioinformatics analysis, including identifying DEGs, gene ontology (GO) enrichment analysis, pathway enrichment analysis and protein-protein interaction (PPI) network and module analysis. The functions of the hub genes were further assessed by miRNA-hub gene regulatory network and TF-hub gene regulatory network construction and analysis. The findings were further validated by receiver operating characteristic curve (ROC) analysis. The aim of this investigation was to identify DEGs and important signaling pathways, and to explore potential candidate biomarkers for the diagnosis and therapeutic targets in COVID-19 infection and its secondary complications.

## Materials and methods

### Data resources

We used “COVID-19” as a keyword on the Gene Expression Omnibus (GEO) database, and NGS dataset (GSE163151) [38] was collected. GSE163151 was in GPL24676 platform Illumina NovaSeq 6000 (Homo sapiens), which included 148 COVID-19 samples and 31 non COVID-19 samples.

### Identification of DEGs

The DESeq2 package of R software [40] was used to analyze the DEGs between COVID-19 and non COVID-19 in the NGS data of GSE163151. The adjusted P-value and [logFC] were calculated. The Benjamini & Hochberg false discovery rate method was used as a correction factor for the adjusted P-value in DESeq2 [41]. The statistically significant DEGs were identified according to adjusted P < .05, and [logFC] > 2.576 for up regulated genes and [logFC] < −2.813 for down regulated genes. ggplot2 and gplot in R software was subsequently performed to plot the volcano plot and heat map.

### GO and pathway enrichment analyses of DEGs

The g:Profiler (http://biit.cs.ut.ee/gprofiler/) [42] is an online functional annotation tool was used to perform Gene Ontology (GO) and REACTOME pathway enrichment analyses. GO enrichment analysis (http://www.geneontology.org) [43] has three independent branches: biological process (BP), cellular component (CC). and molecular function (MF). The REACTOME (https://reactome.org/) [44] pathway database facilitates the systematic analysis of the intracellular metabolic pathways and functions of gene products. P value <0.05 as the cutoff criterion was considered statistically significant.

### Construction of the PPI network and module analysis

IID (Integrated Interactions Database) interactome (http://iid.ophid.utoronto.ca/search_by_proteins/) [45] is a database that provides a function for predicted protein interactions and providing systematic data regarding cellular processes. Subsequently, PPI networks was constructed using the Cytoscape software version 3.8.2 (http://www.cytoscape.org/) [46]. In addition, the node degree [47], betweenness centrality [48], stress centrality [49] and closeness centrality [50] of each protein node in the PPI network was calculated using Network Analyzer. In order to screen the clusters of original PPI network, module analysis by PEWCC1 plug-in [51] was carried out in Cytoscape software. The criteria for the network module screening were set beyond the number of nodes and PEWCC1 scores. Moreover, the GO and pathway enrichment analysis for DEGs were performed by g:Profiler software in the modules with a threshold of p<0.05.

### MiRNA-hub gene regulatory network construction

The miRNet database (https://www.mirnet.ca/) [52] is an open-source platform mainly focusing on miRNA-hub gene interactions. miRNet utilizes fourteen established miRNA-hub gene prediction databases, including TarBase, miRTarBase, miRecords, miRanda (S mansoni only), miR2Disease, HMDD, PhenomiR, SM2miR, PharmacomiR, EpimiR, starBase, TransmiR, ADmiRE, and TAM 2.0. Subsequently, the network of the hub genes and their targeted miRNAs was visualized by Cytoscape software version 3.8.2 [46].

### TF-hub gene regulatory network construction

The NetworkAnalyst database (https://www.networkanalyst.ca/) [53] is an open-source platform mainly focusing on TF-hub gene interactions. NetworkAnalyst database utilizes fourteen established TF-hub gene prediction database Jasper.. Subsequently, the network of the hub genes and their targeted TFs was visualized by Cytoscape software version 3.8.2 [46].

### Receiver operating characteristic curve (ROC) analysis

ROC curve analyses to resolve the specificity, sensitivity, likelihood ratios, positive predictive values, and negative predictive values for all potential thresholds of the ROC curve were accomplished using the pROC package in R statistical software [54]. ROC analysis was performed to discriminate COVID-19 from non COVID-19. The area under the curve (AUC) was calculated and used to compare the diagnostic value of hub genes.

## Results

### Identification of DEGs

To identify DEGs in the omental white adipocytes between ISO and IRO subjects, we retrieved relevant Expression profiling by high throughput sequencing of GSE163151 from GEO database. A total of 954 DEGs were identified from the NGS dataset, including 477 up regulated genes and 477 down regulated genes COVID-19 compared to non COVID-19. The expression levels of the up regulated and down regulated genes are displayed in Table 1. The expression levels of these DEGs are visualized in the form of a heatmap (Fig. 1). Volcano plots were generated to visualize fold changes of the DEGs had [logFC] > 2.576 for up regulated genes and [logFC] < −2.813 for down regulated genes, and adjusted P<0.05 (Fig. 2).

**Table 1.**
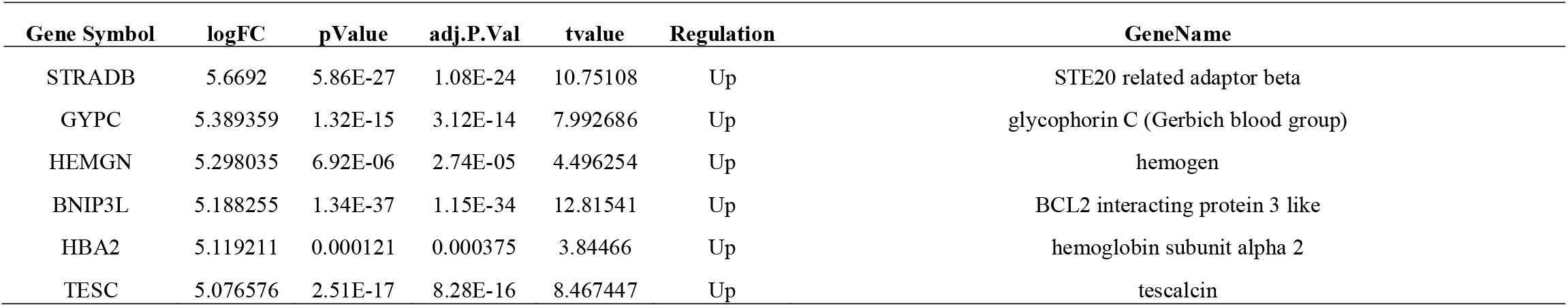
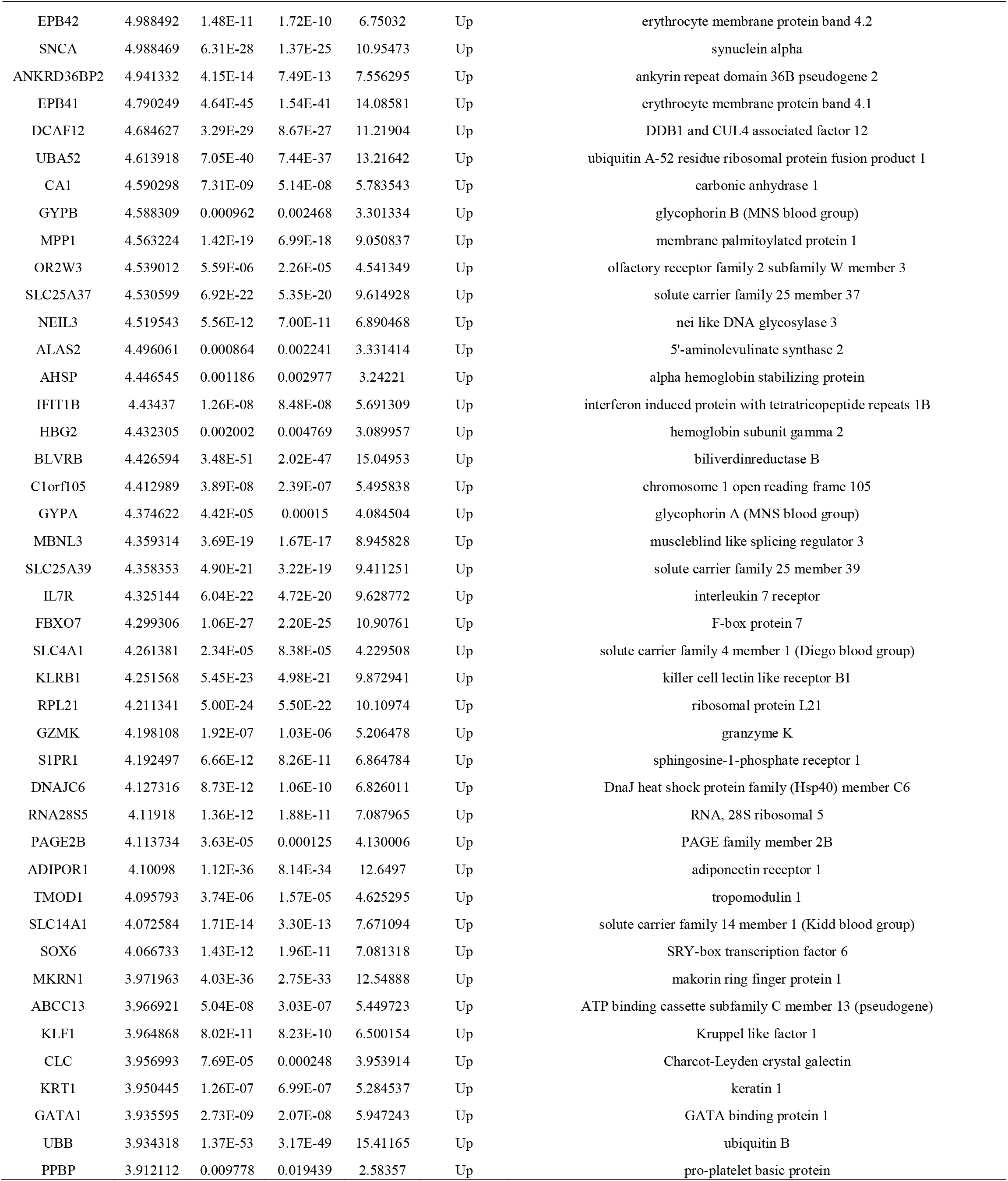
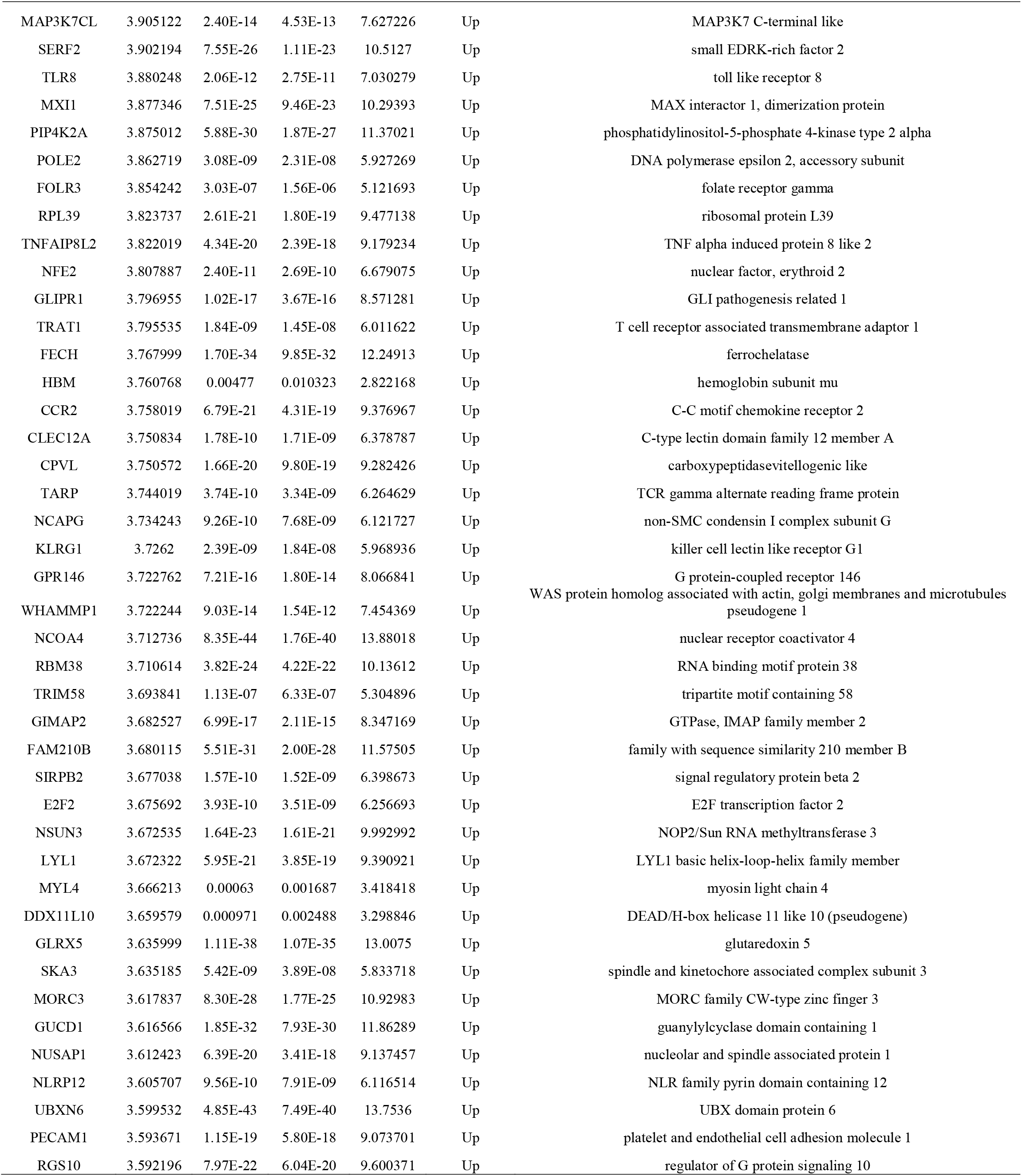
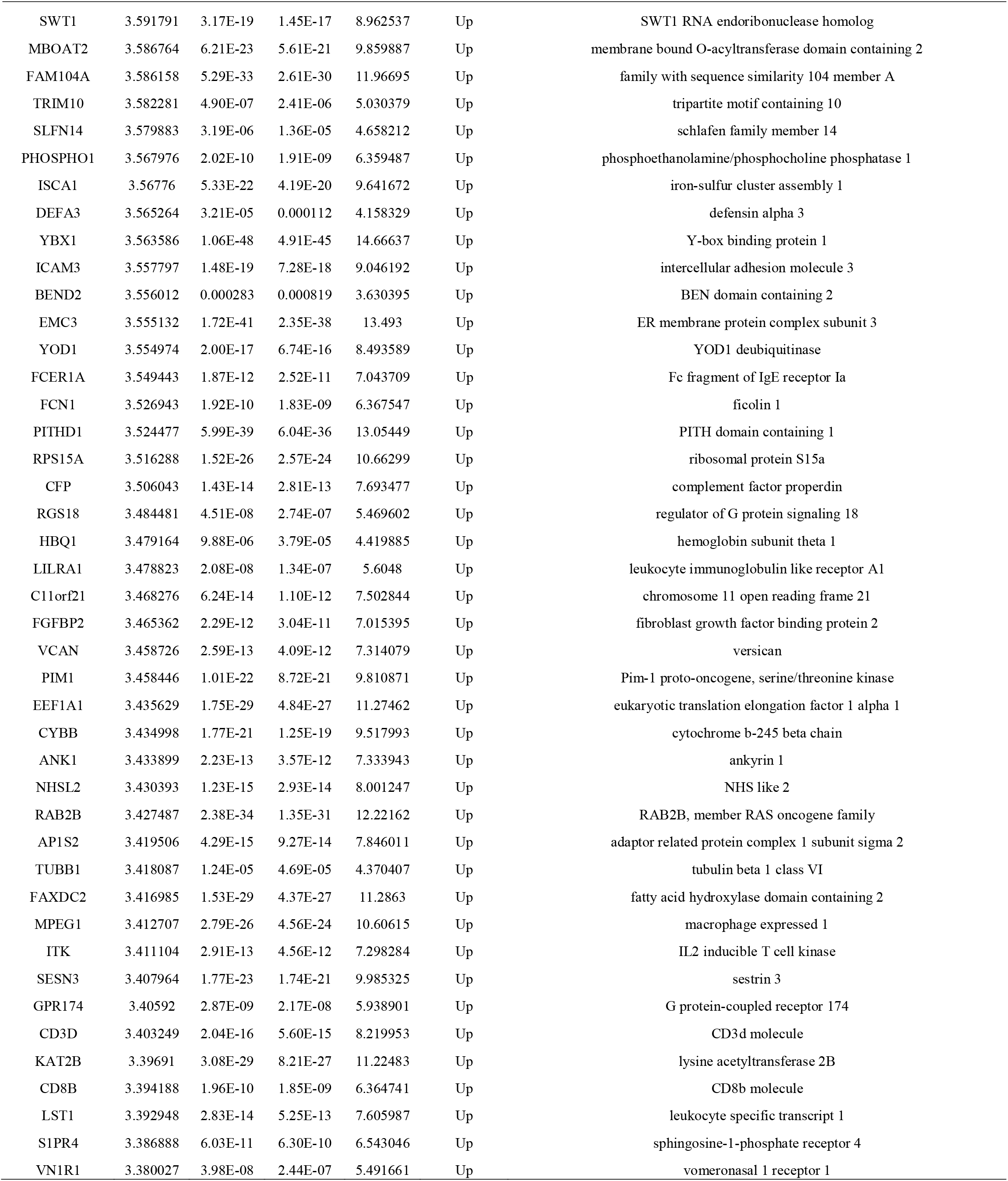
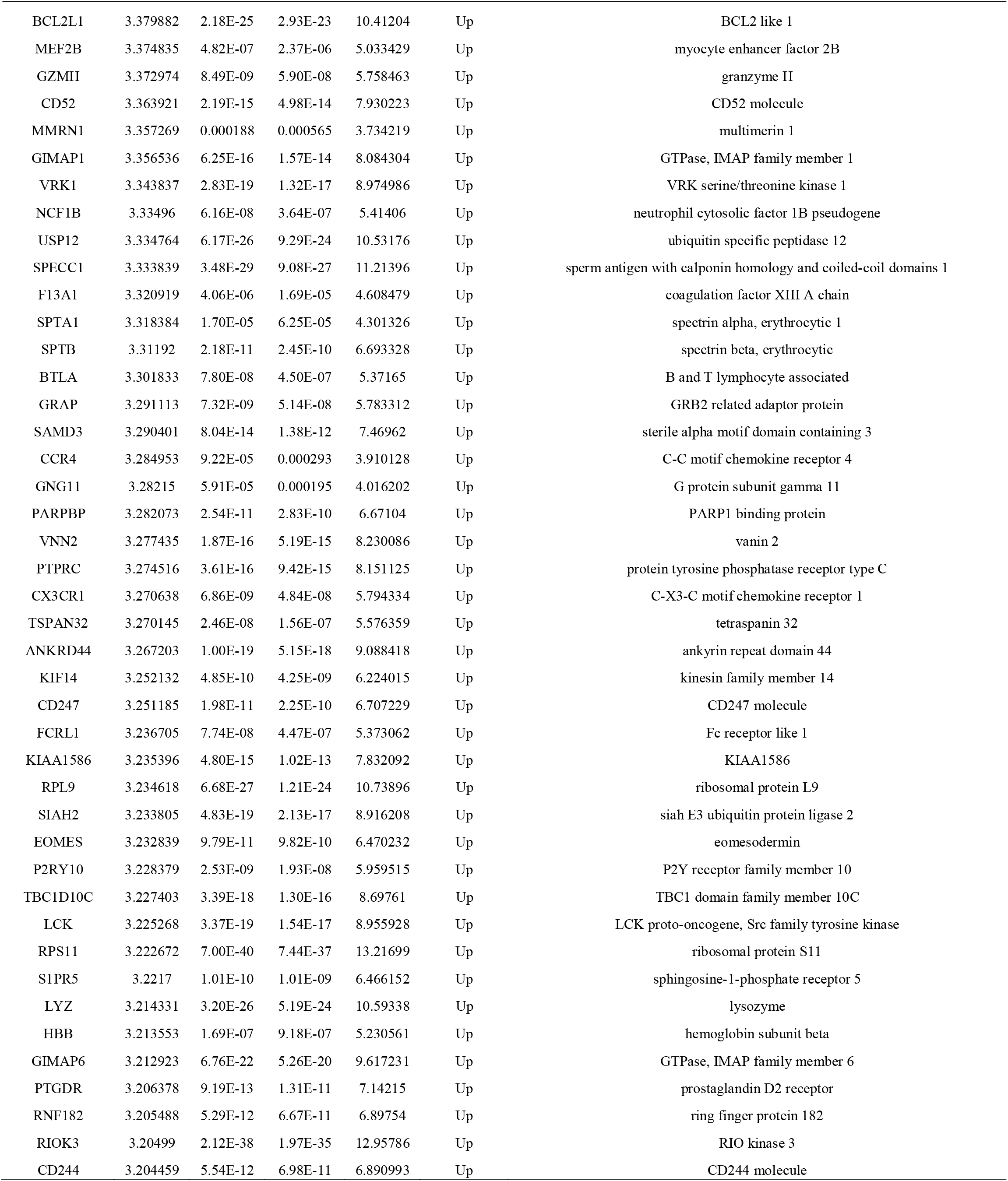
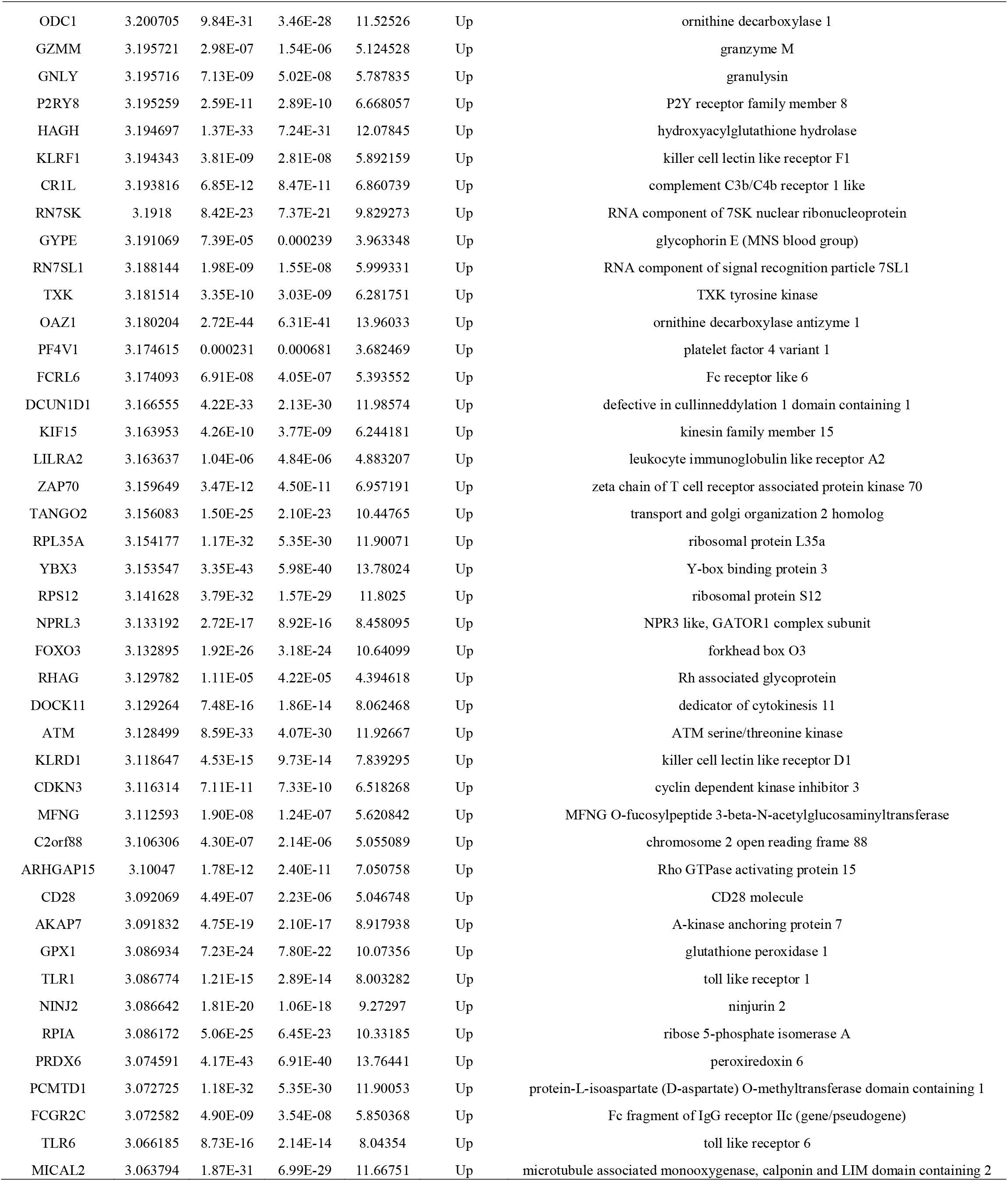
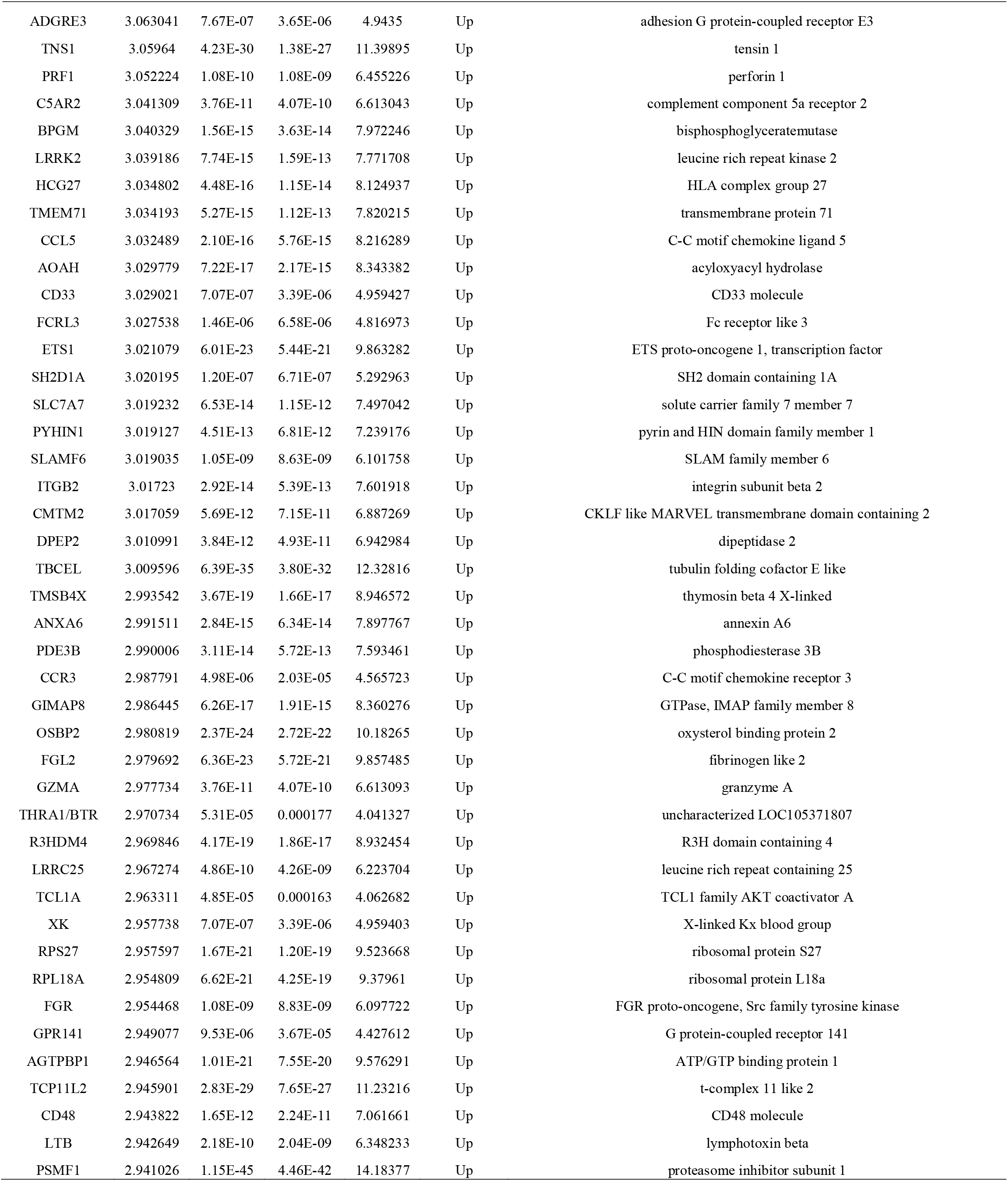
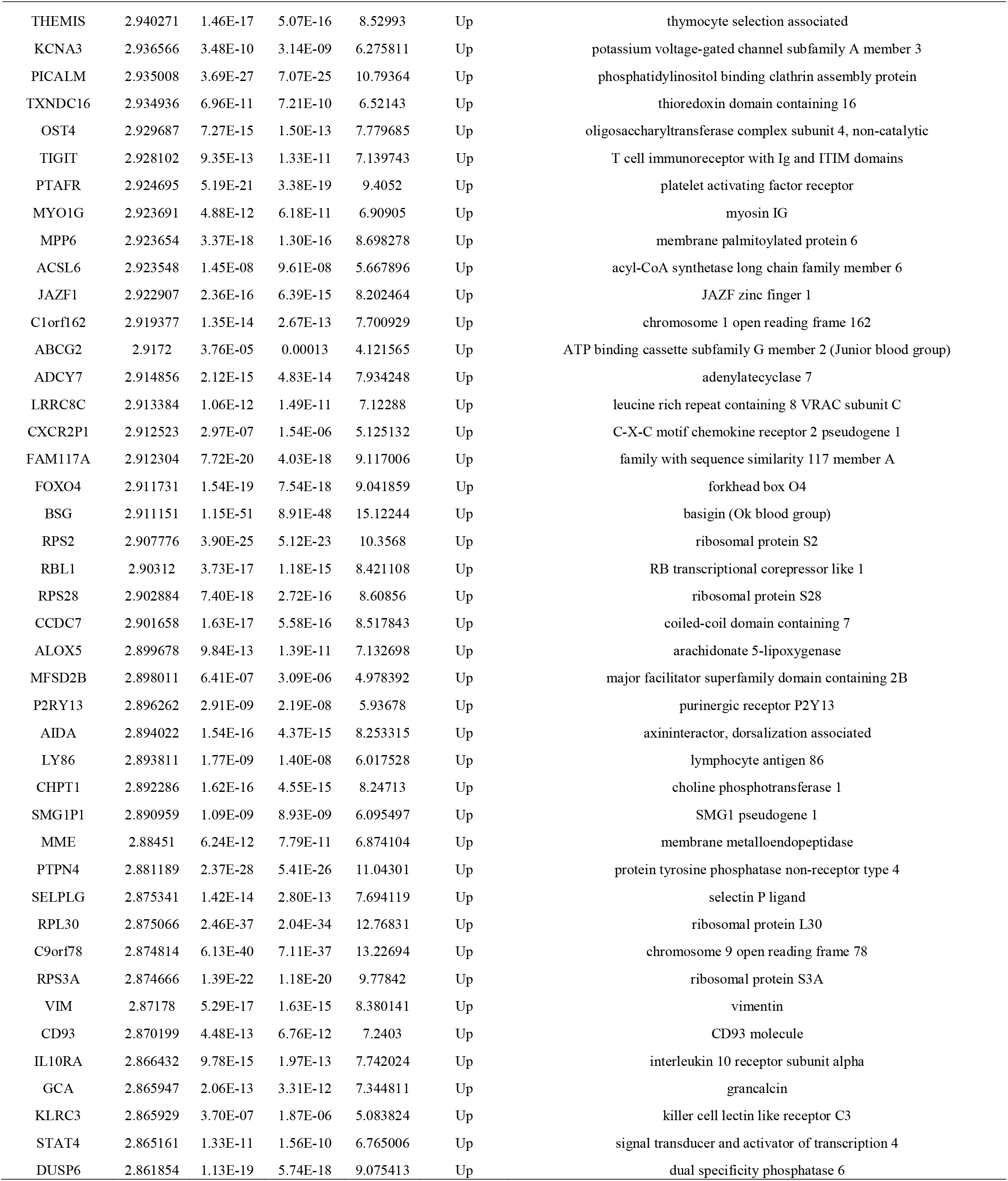
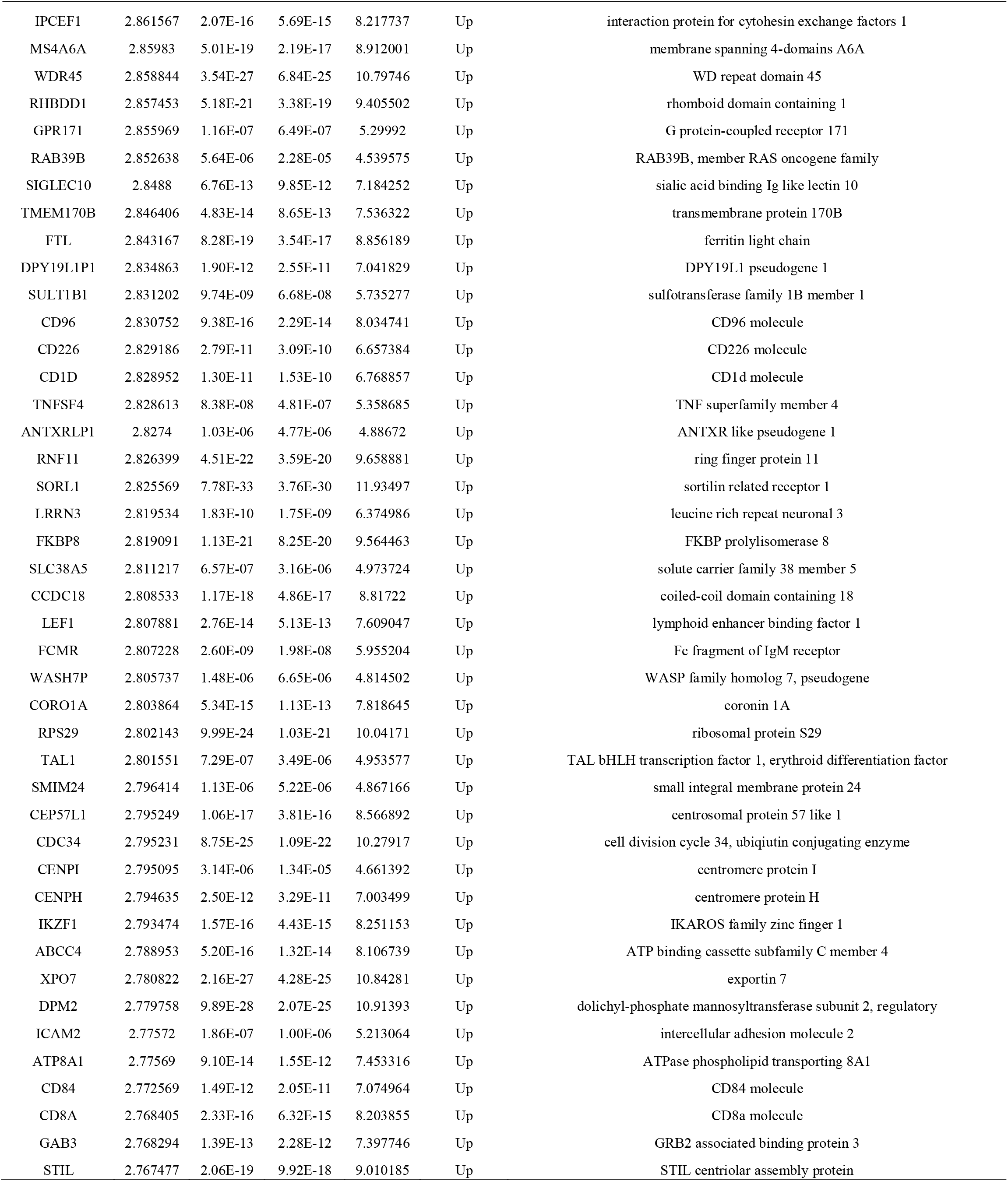
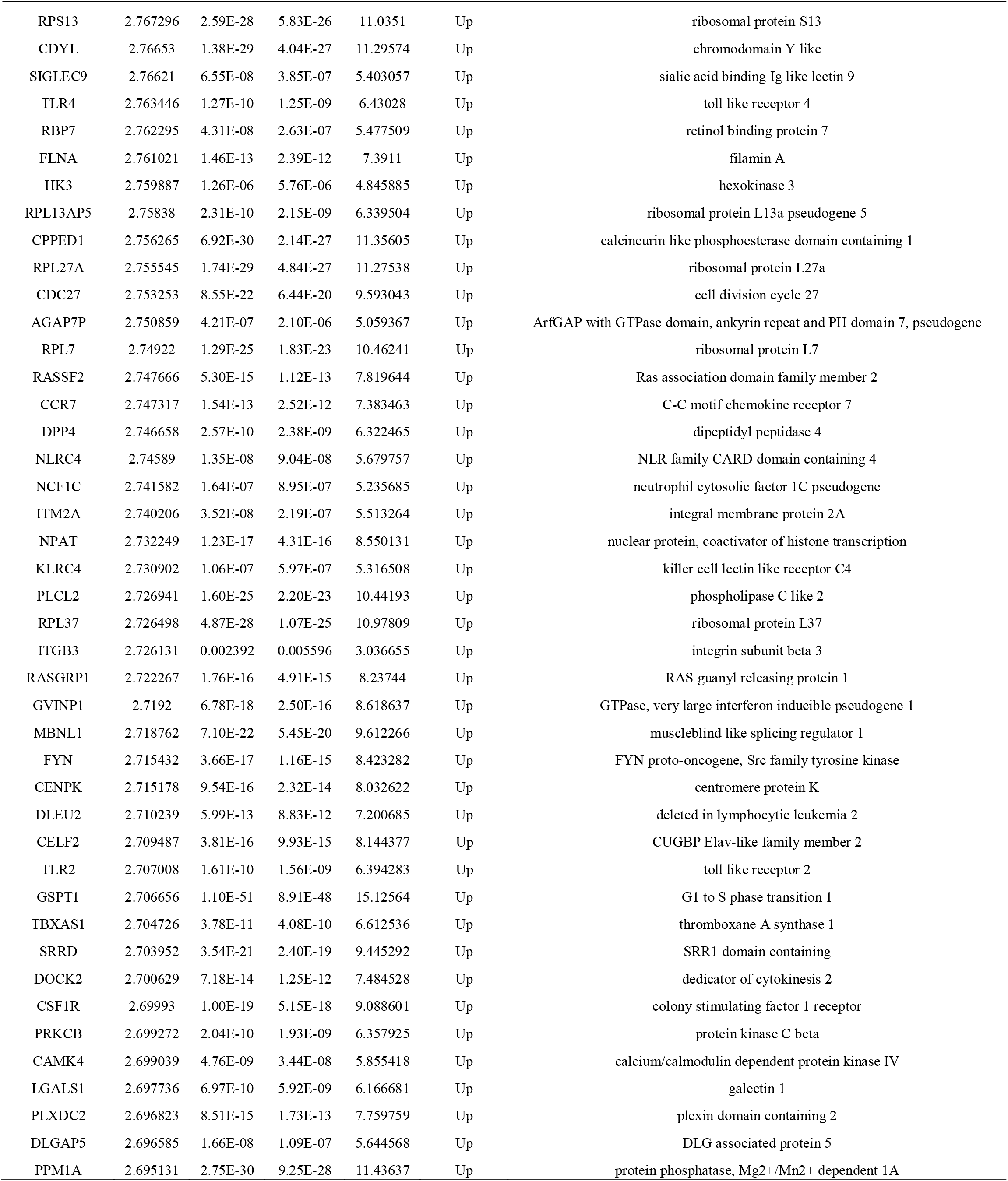
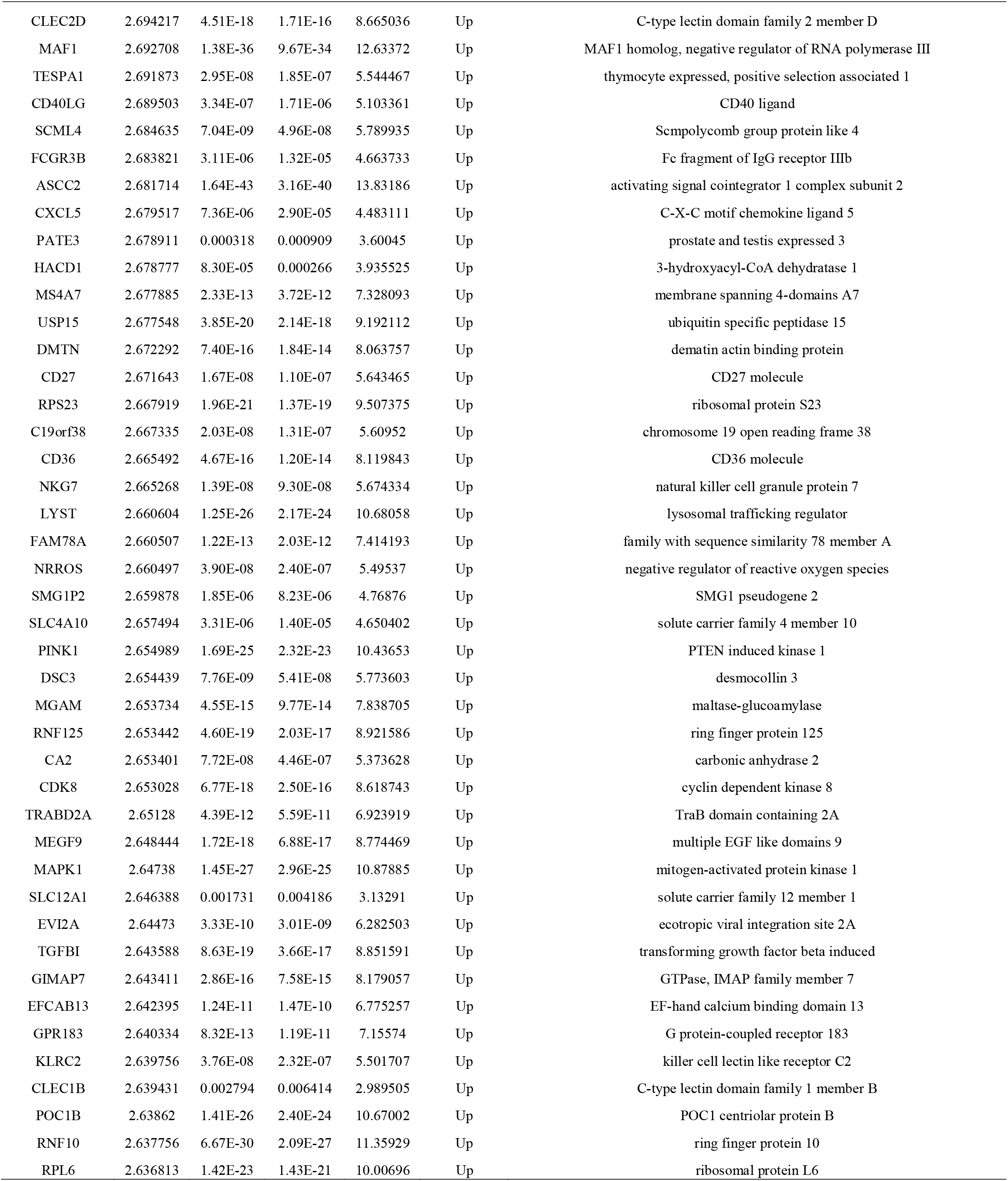
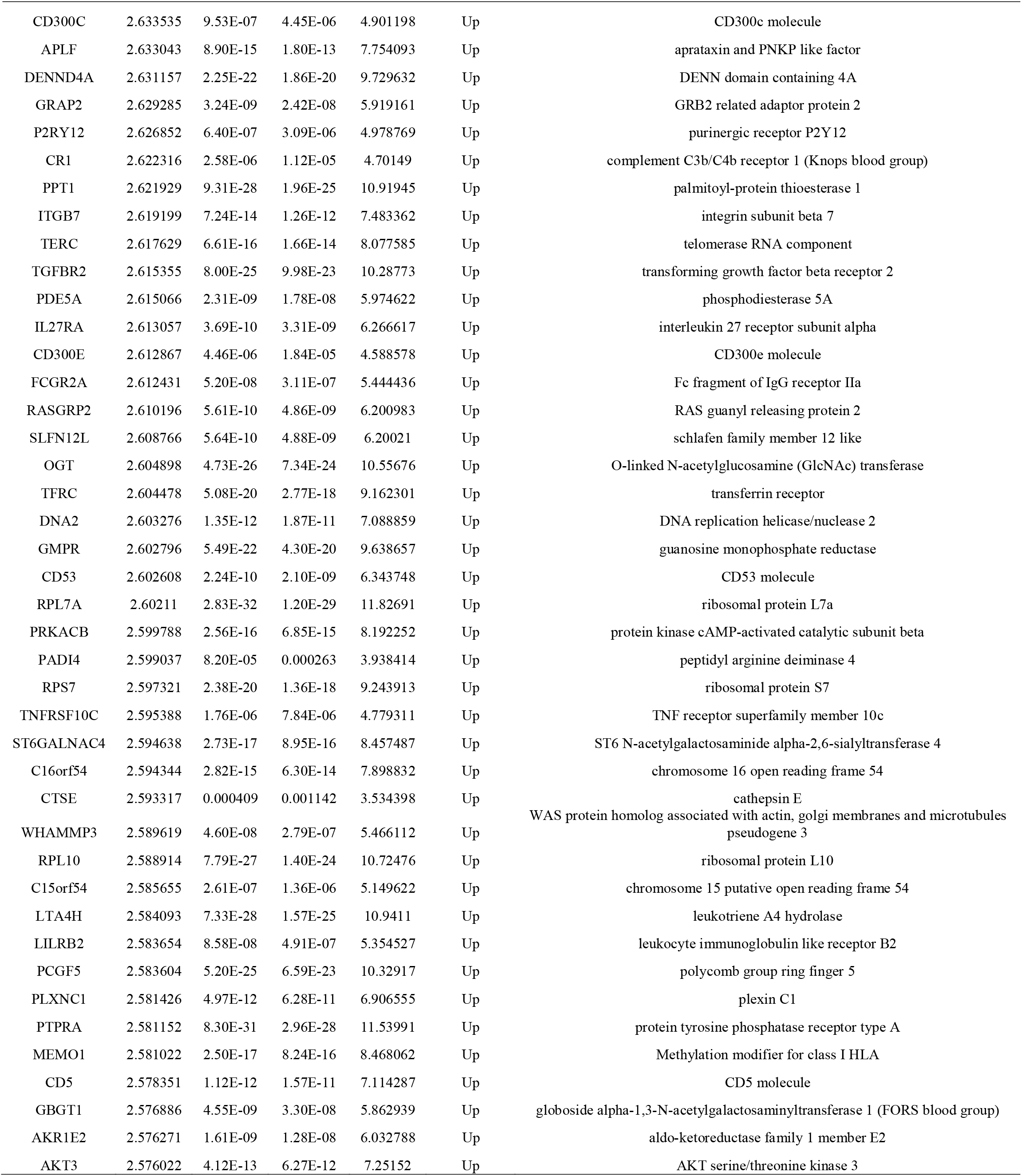
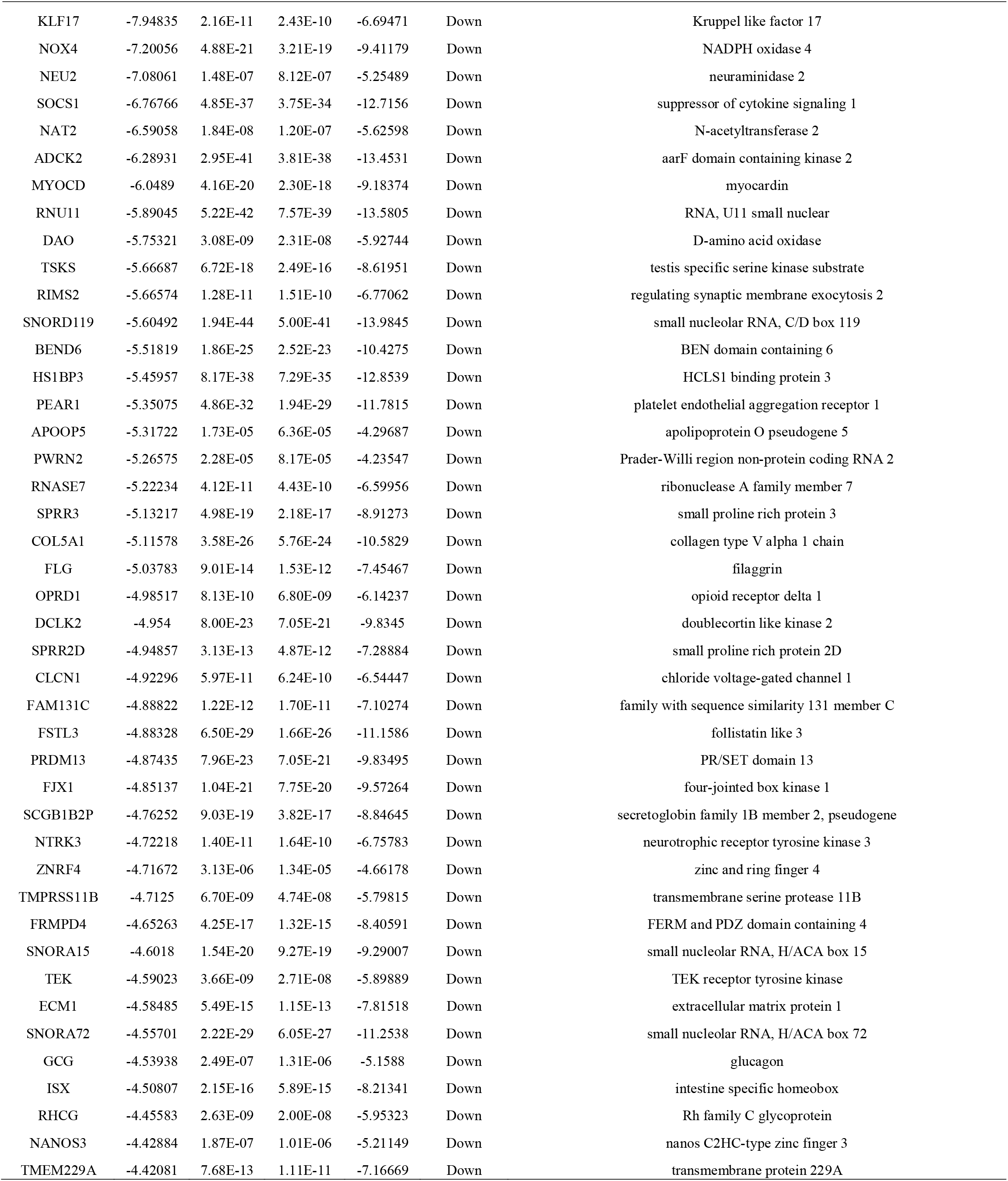
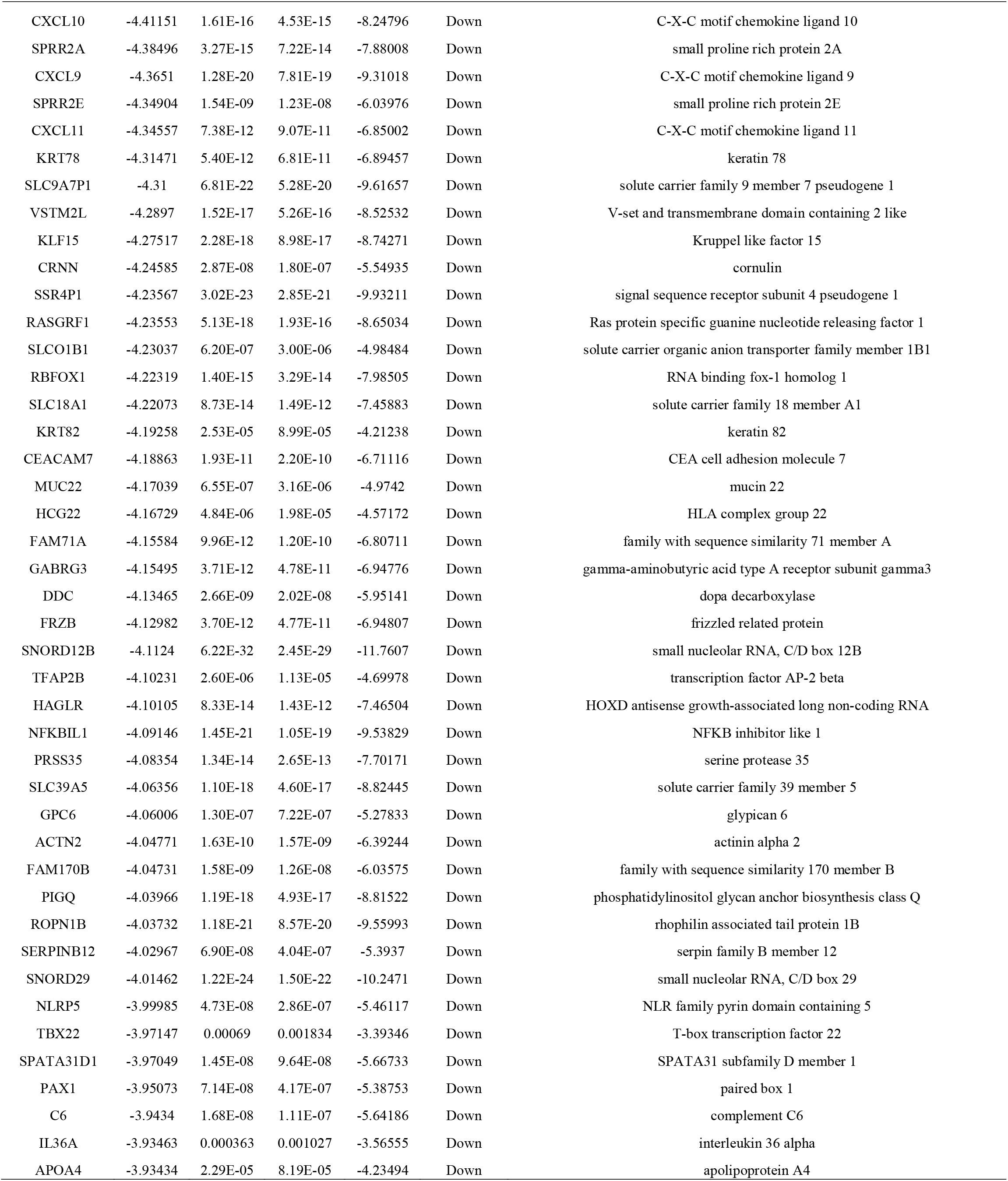
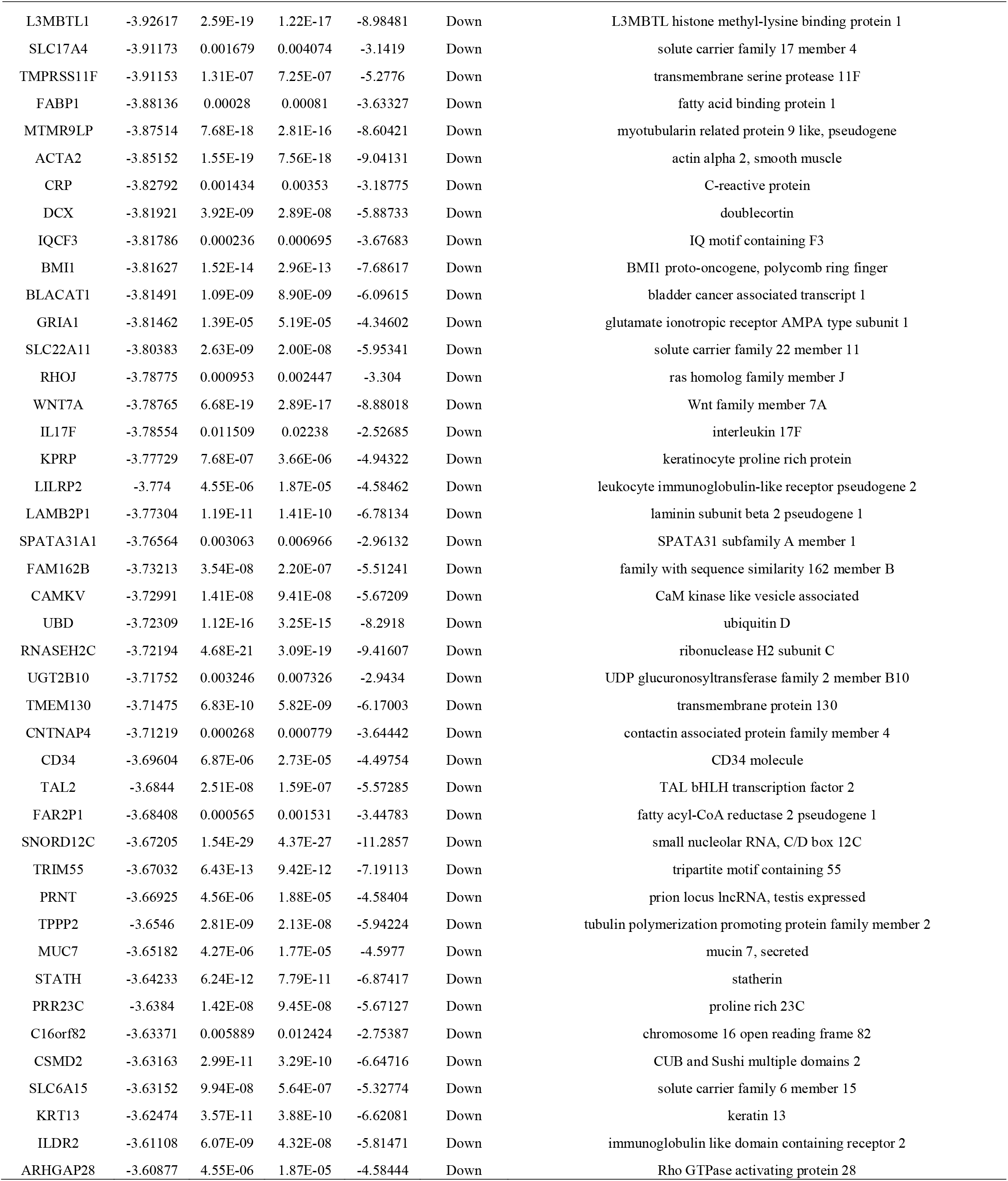
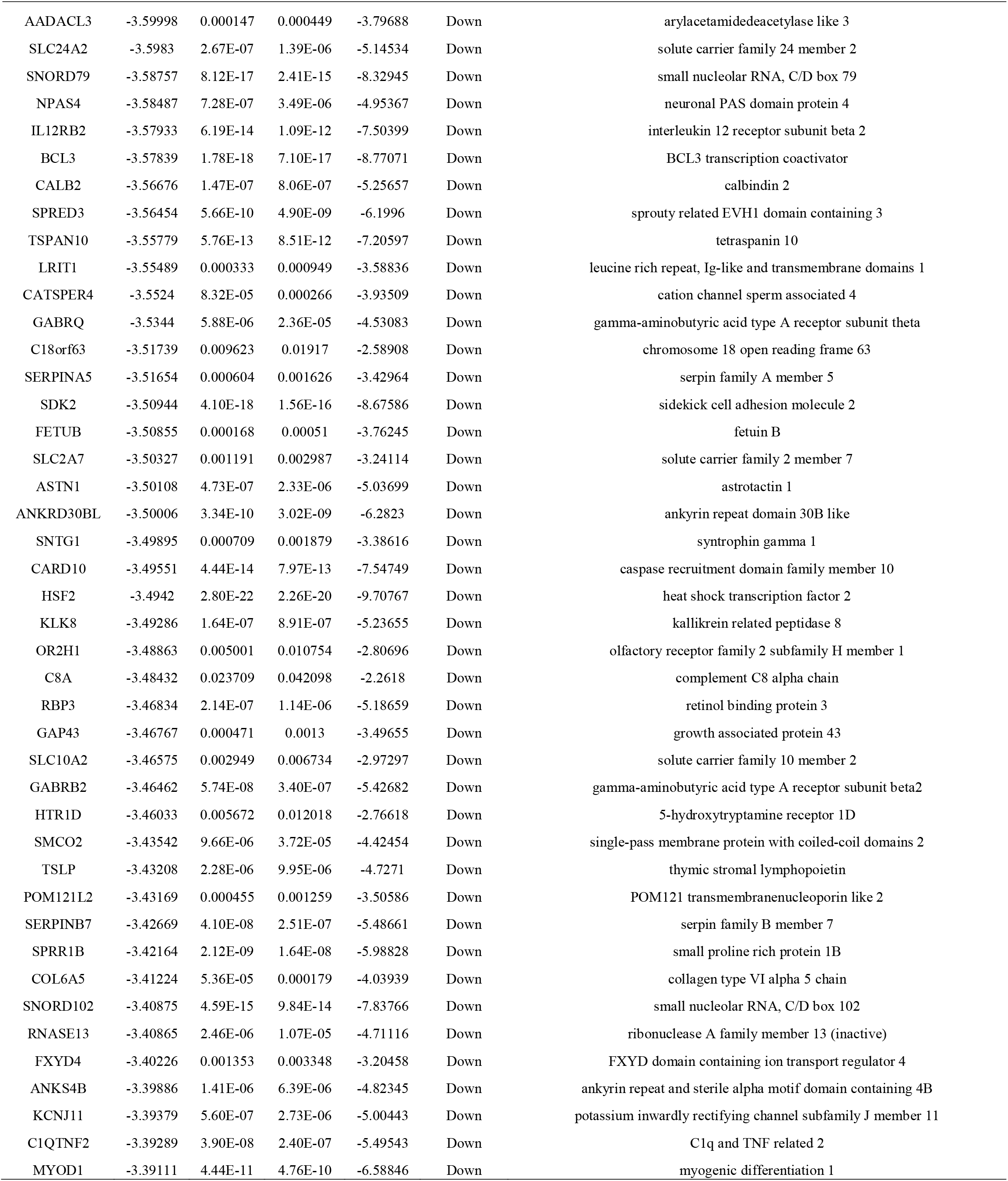
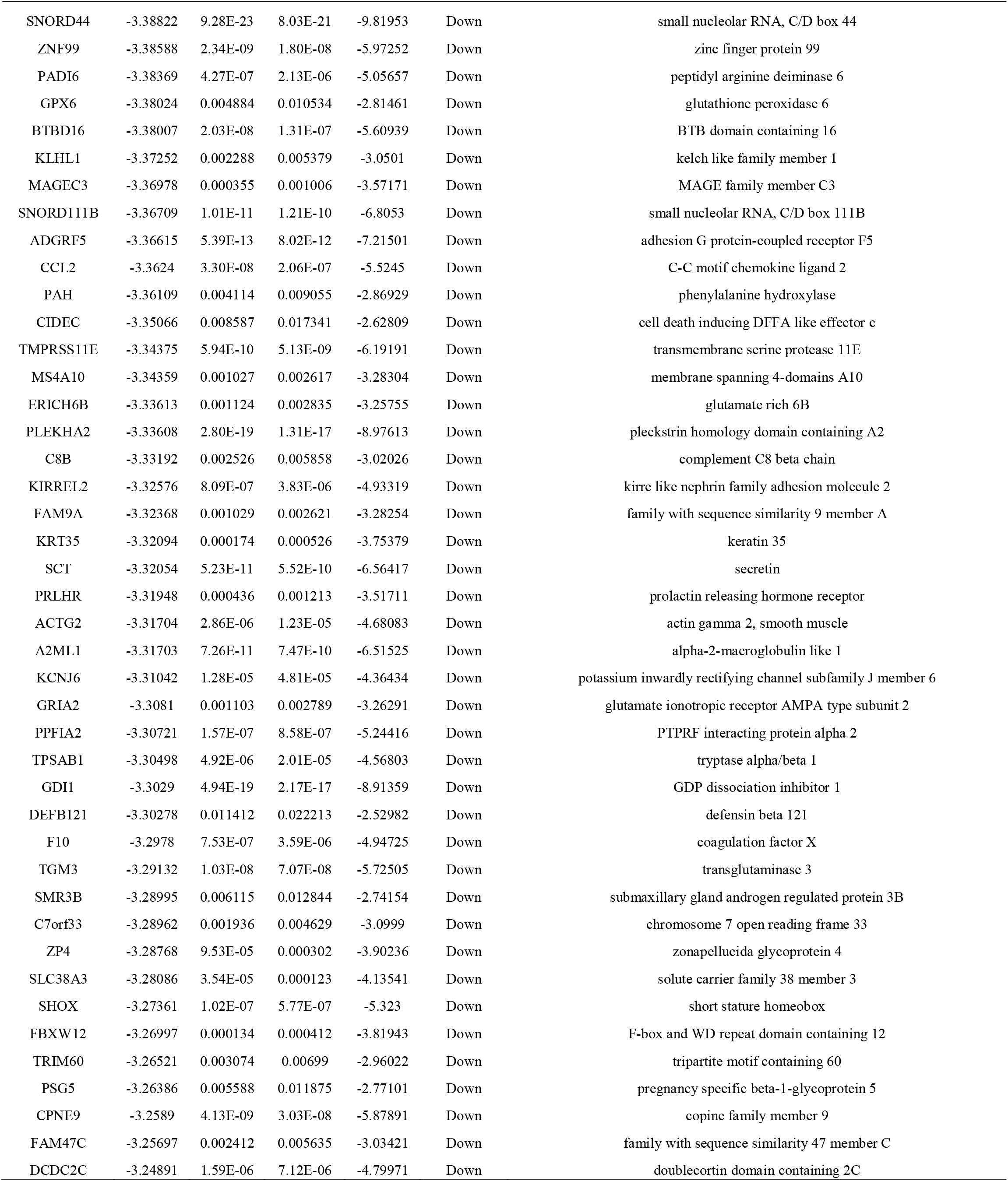
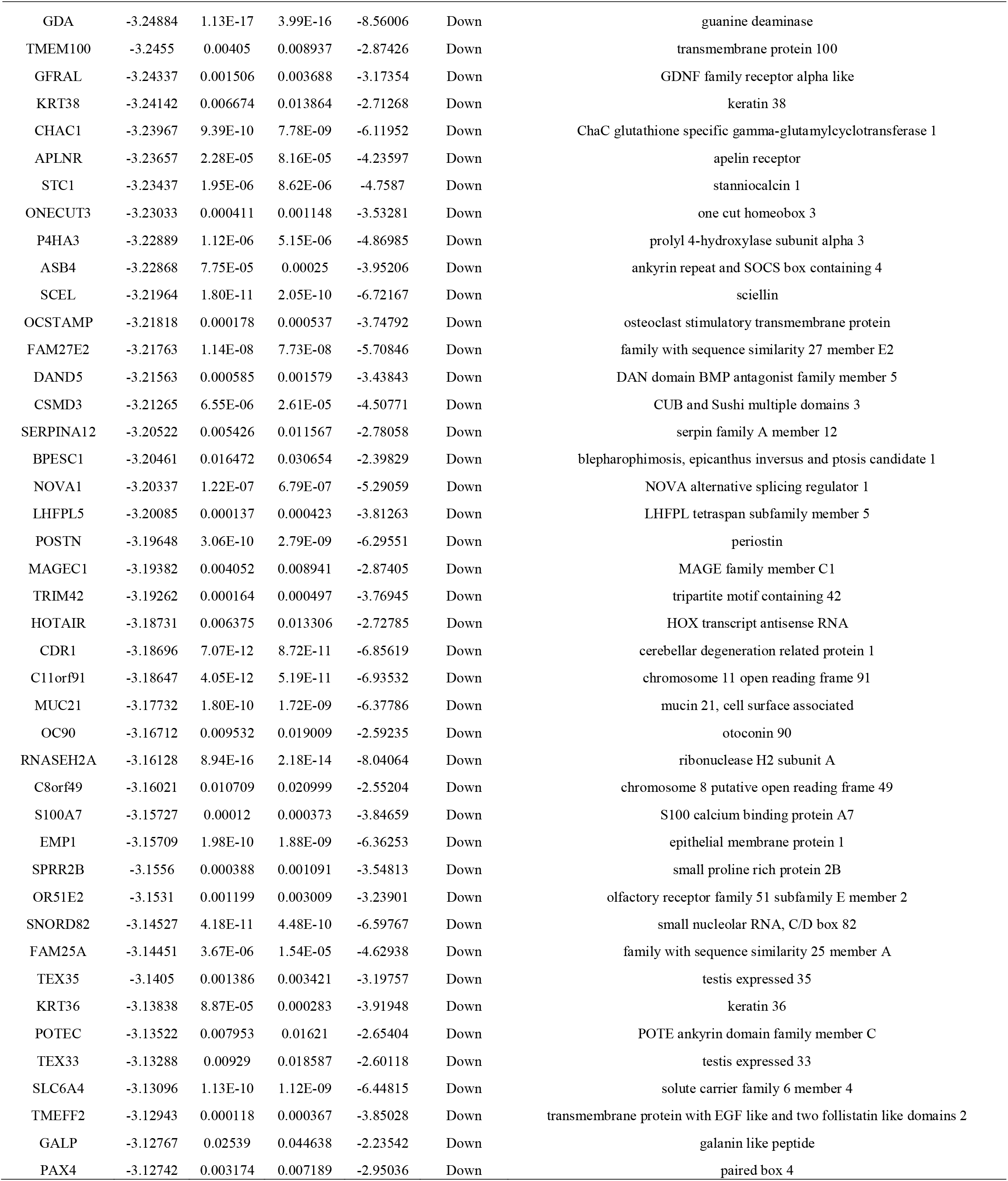
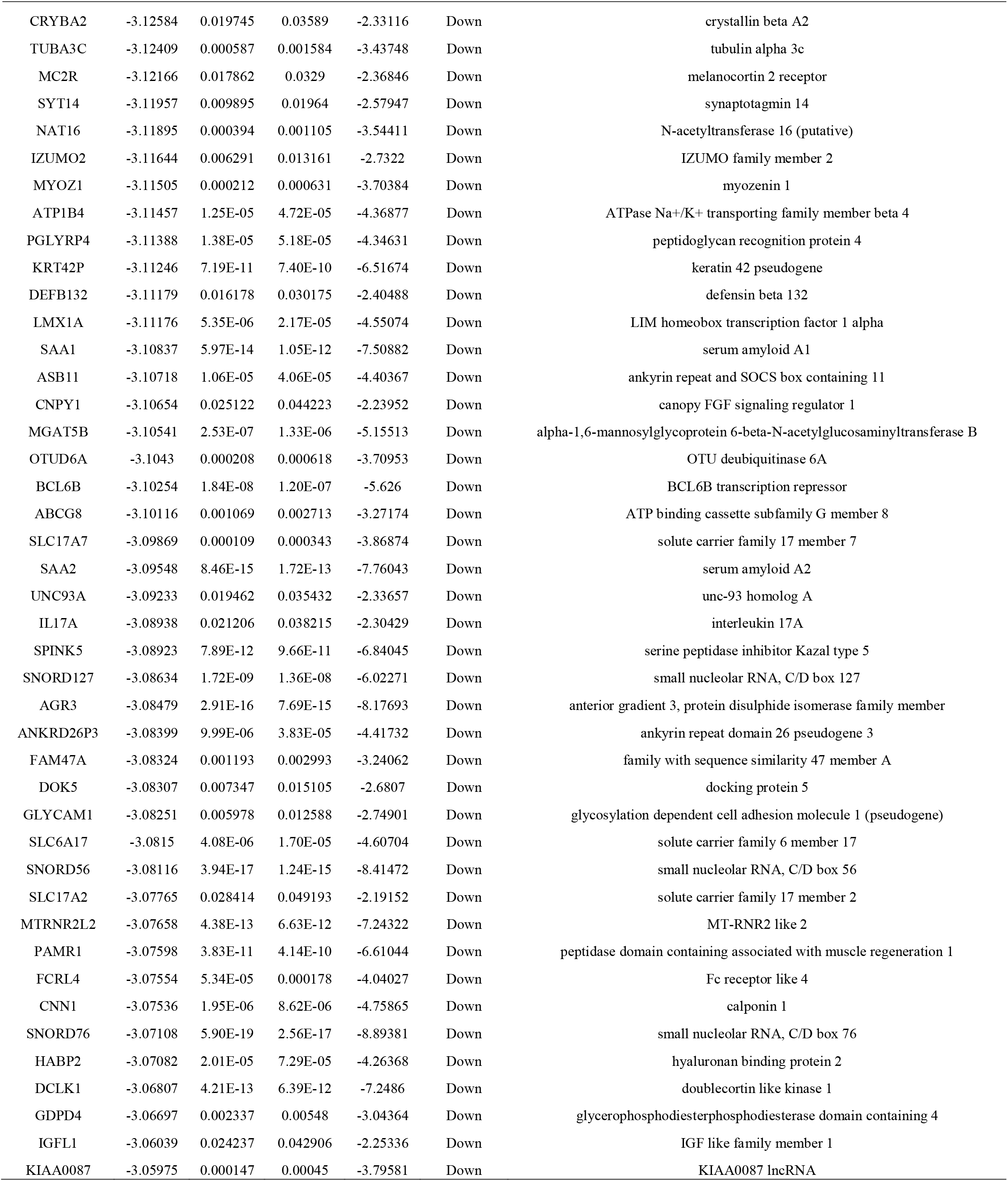
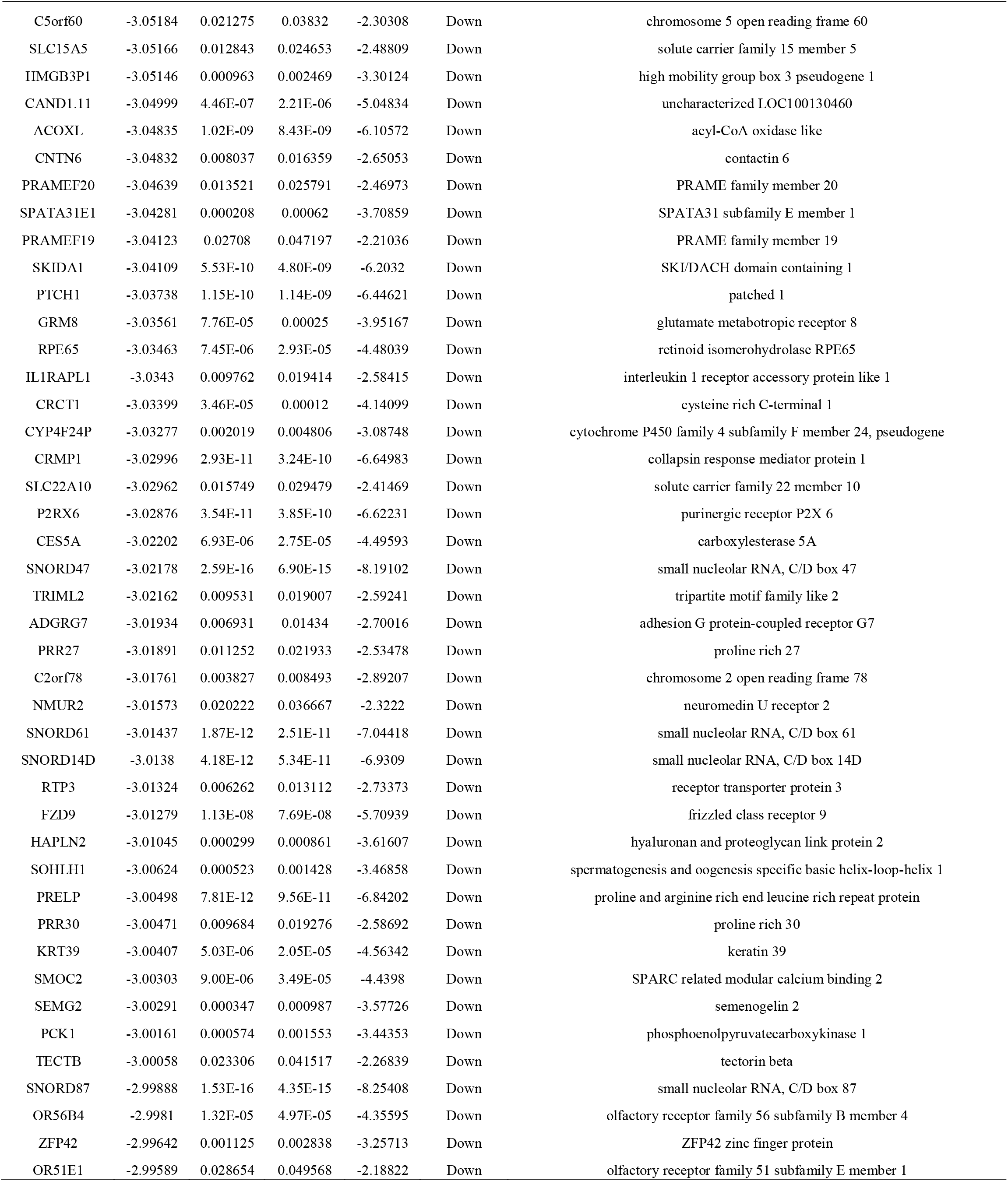
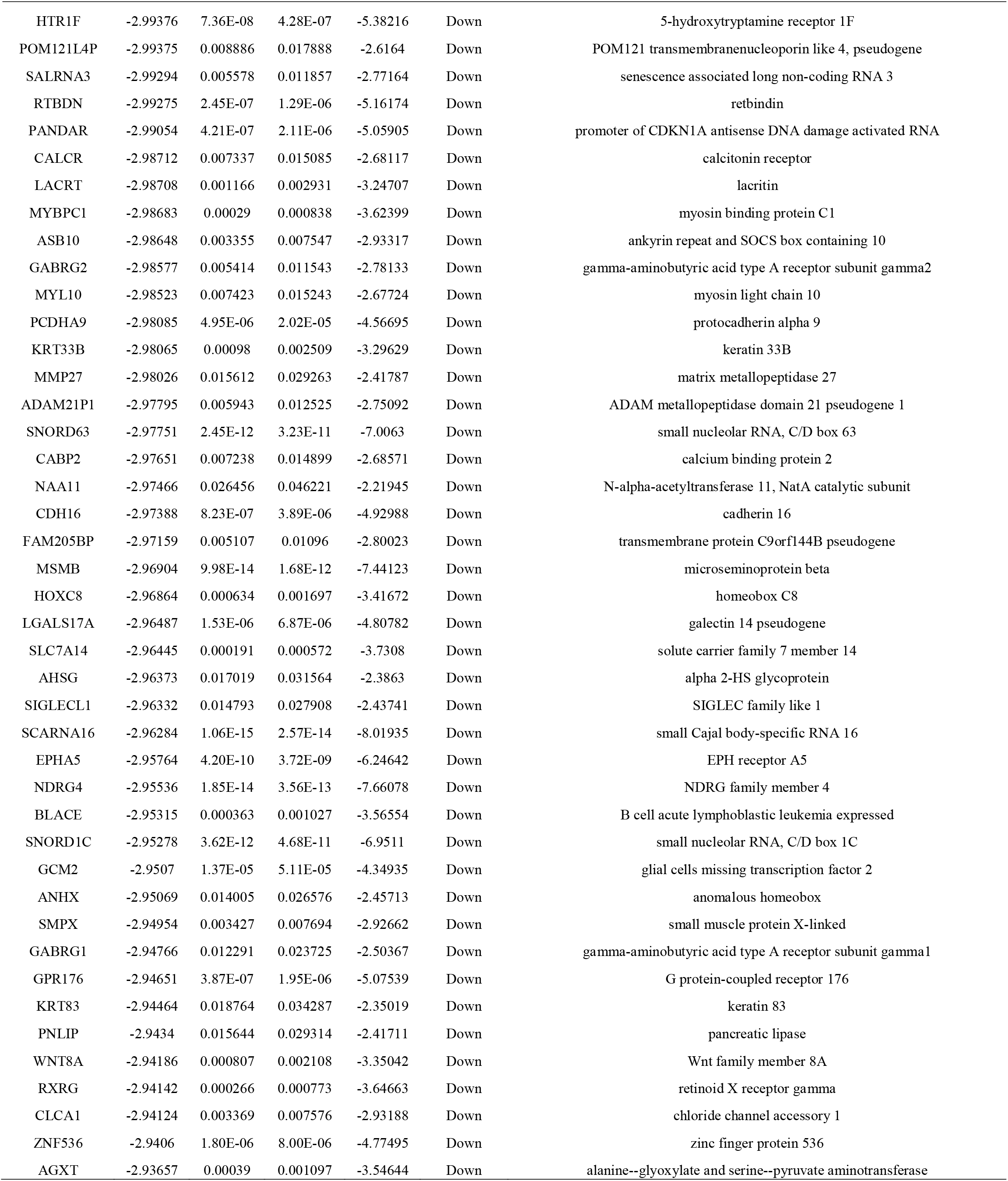
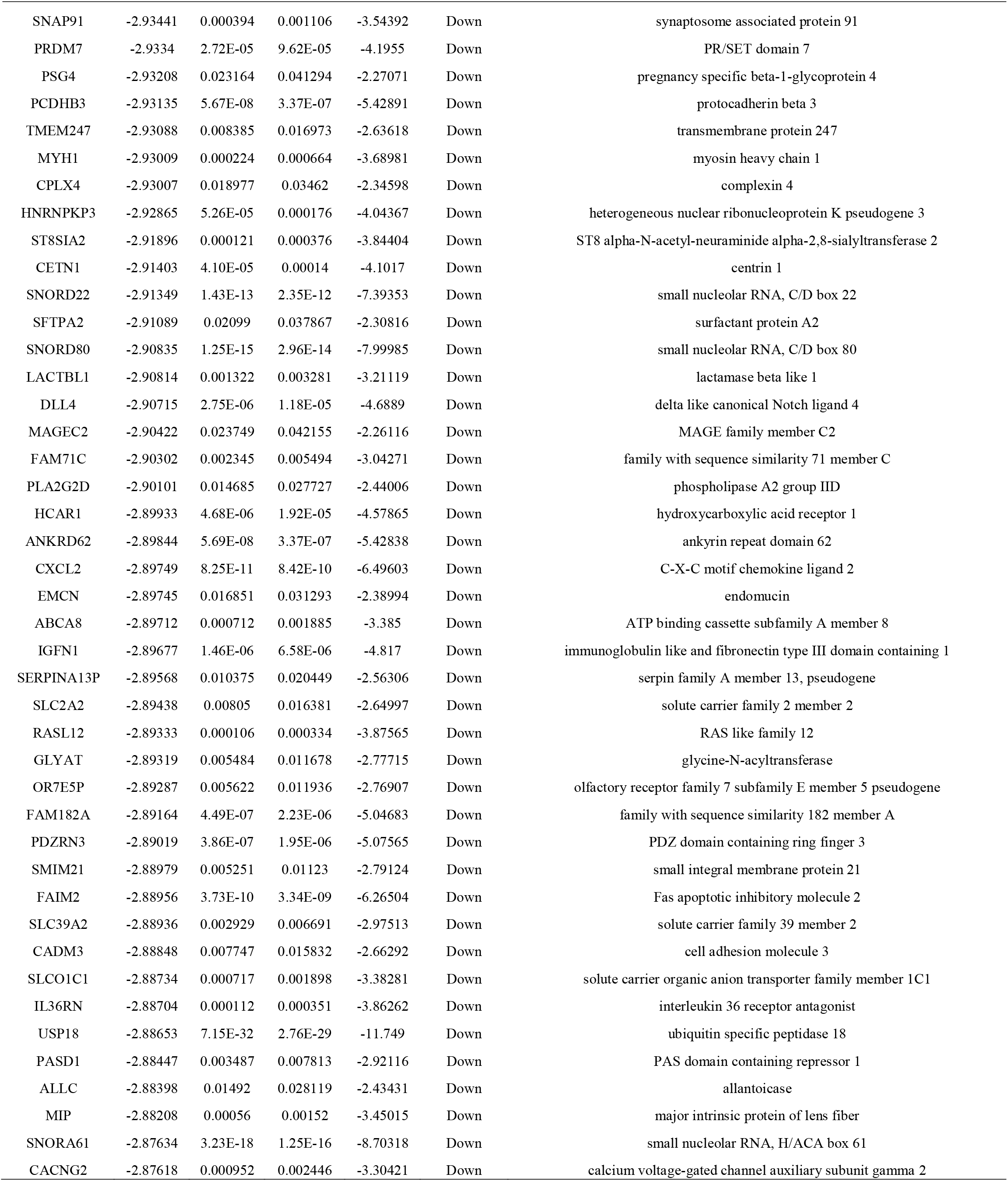
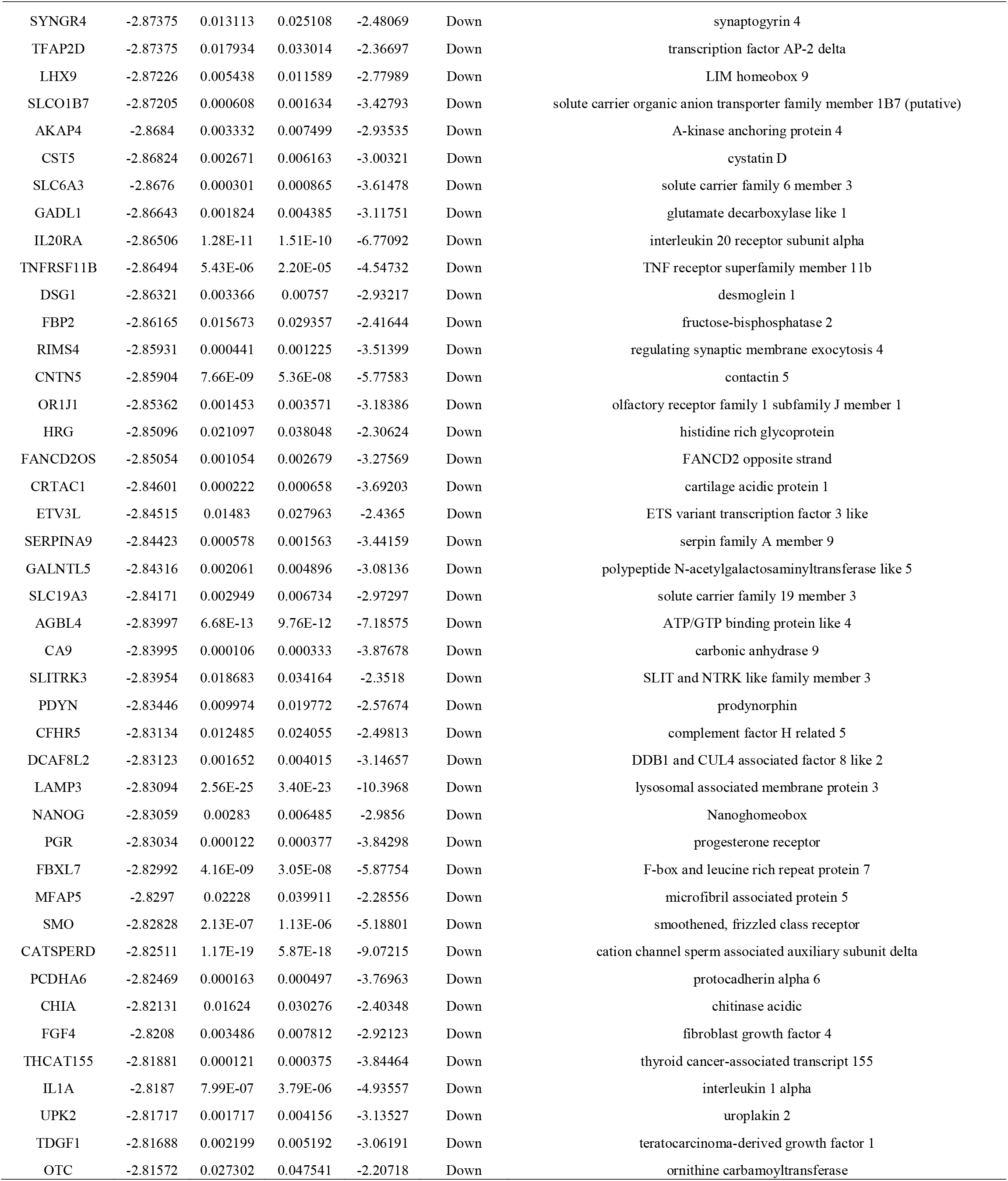
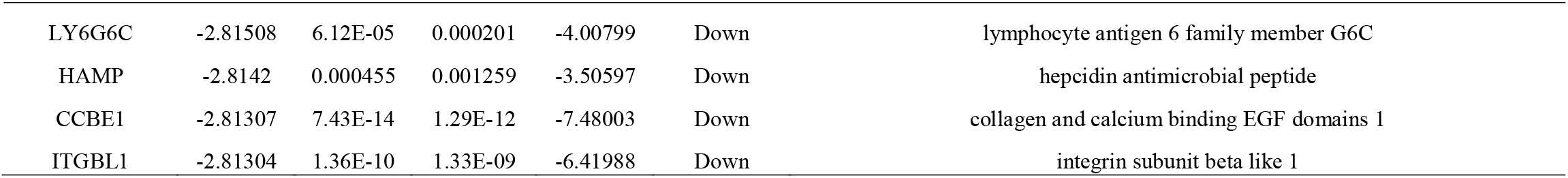
The statistical metrics for key differentially expressed genes (DEGs)

**Fig. 1.**
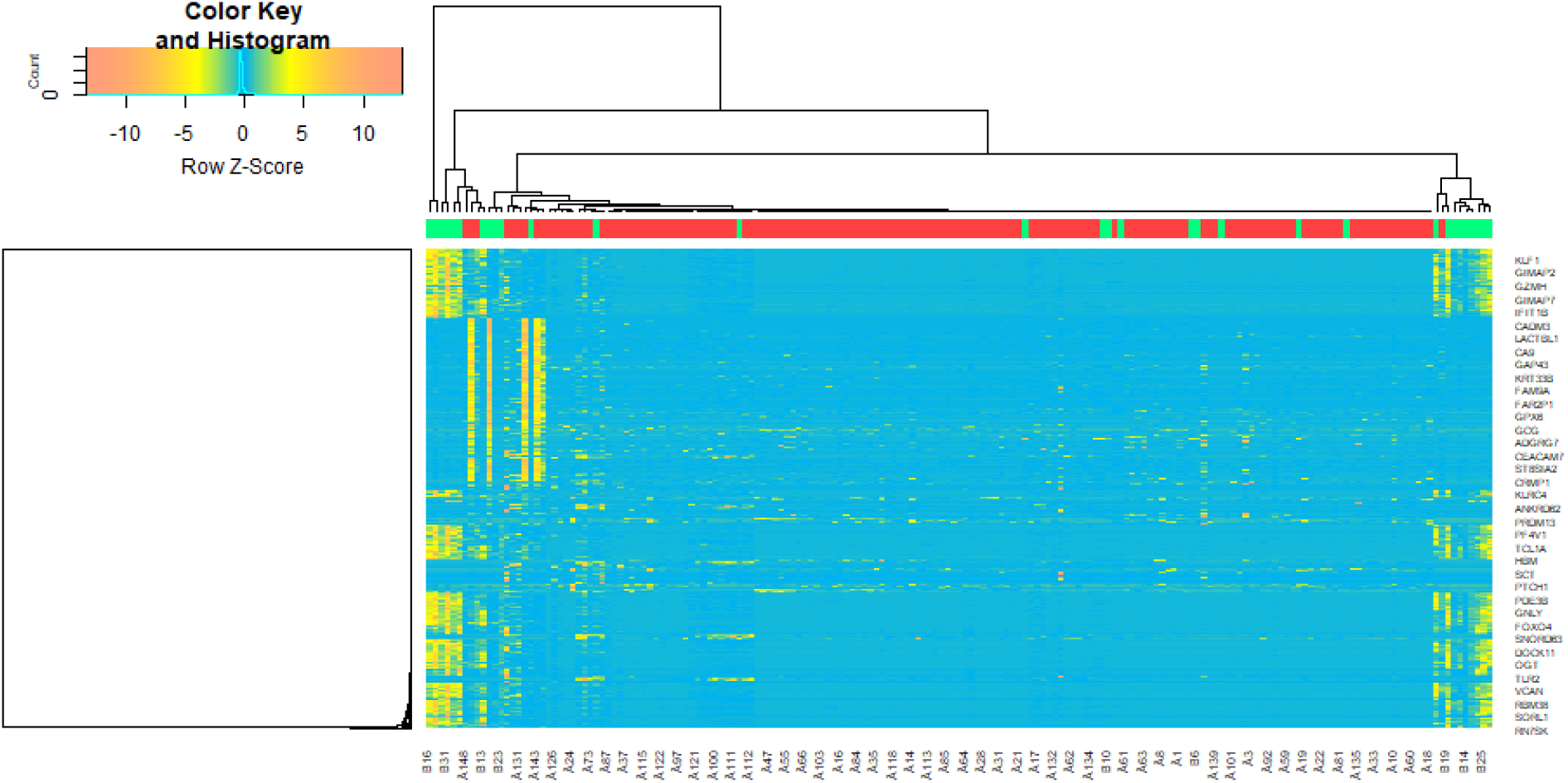
Volcano plot of differentially expressed genes. Genes with a significant change of more than two-fold were selected. Green dot represented up regulated significant genes and red dot represented down regulated significant genes.

**Fig. 2.**
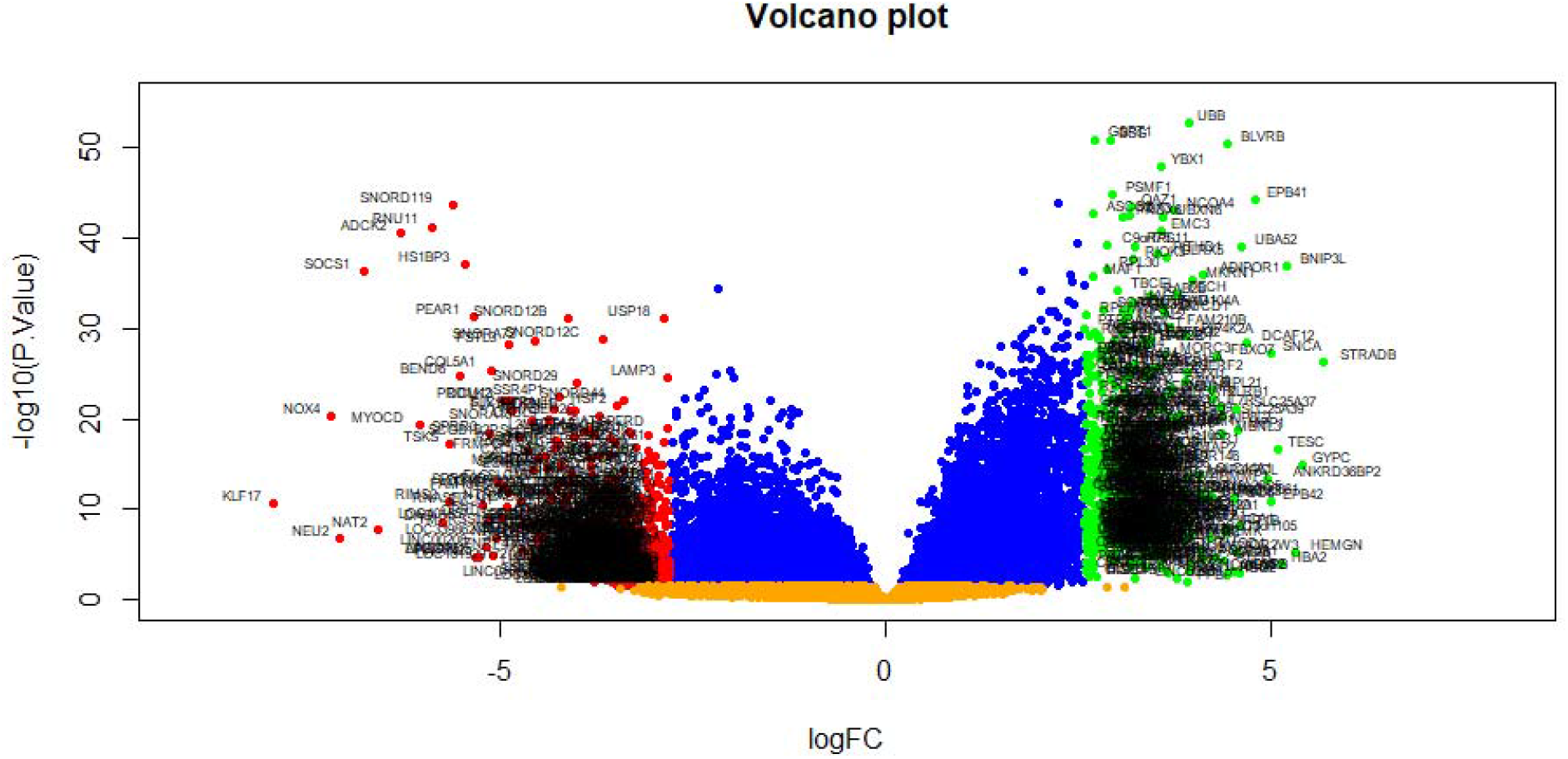
Heat map of differentially expressed genes. Legend on the top left indicate log fold change of genes. (A1 – A37 = normal control samples; B1 – B47 = HF samples)

### GO and pathway enrichment analyses of DEGs

To identify key genes related to sepsis, gene functions were annotated using the g:Profiler online software database. DEGs were classified into the three functional groups of GO analysis: BP, CC and MF (Table 2). GO enrichment analysis showed that DEGs were primarily enriched in immune system process, regulation of immune system process, multicellular organismal process and anatomical structure development were significantly enriched in BP; membrane, cell periphery, cell periphery and extracellular region were significantly enriched in CC; and in the MF, DEGs were mainly enriched in molecular transducer activity, protein binding, transporter activity and receptor ligand activity. DEGs were found to be significantly enriched in nine classical pathways, including the immune system, peptide chain elongation, formation of the cornified envelope and GPCR ligand binding and are listed in Table 3.

**Table 2.**
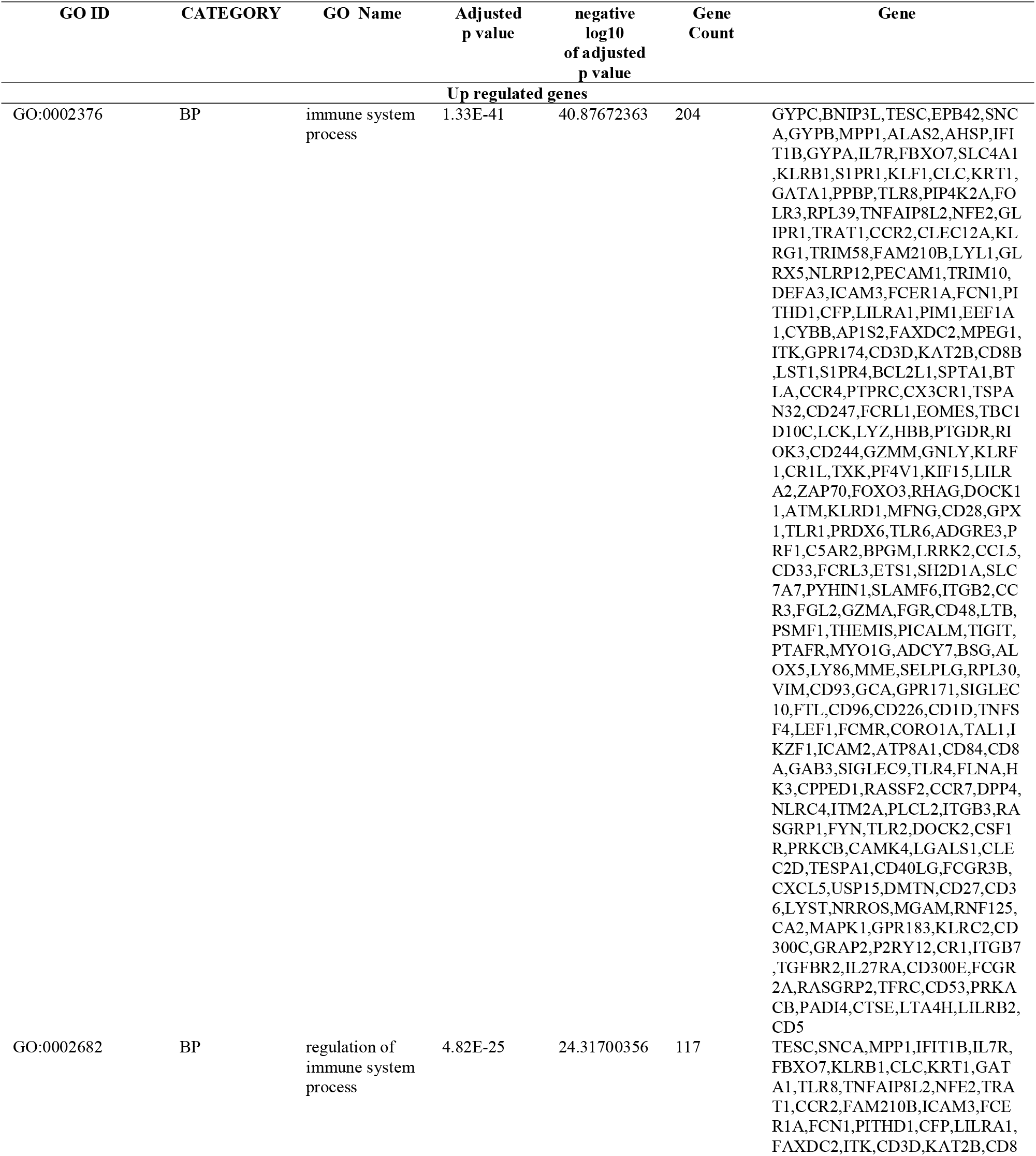
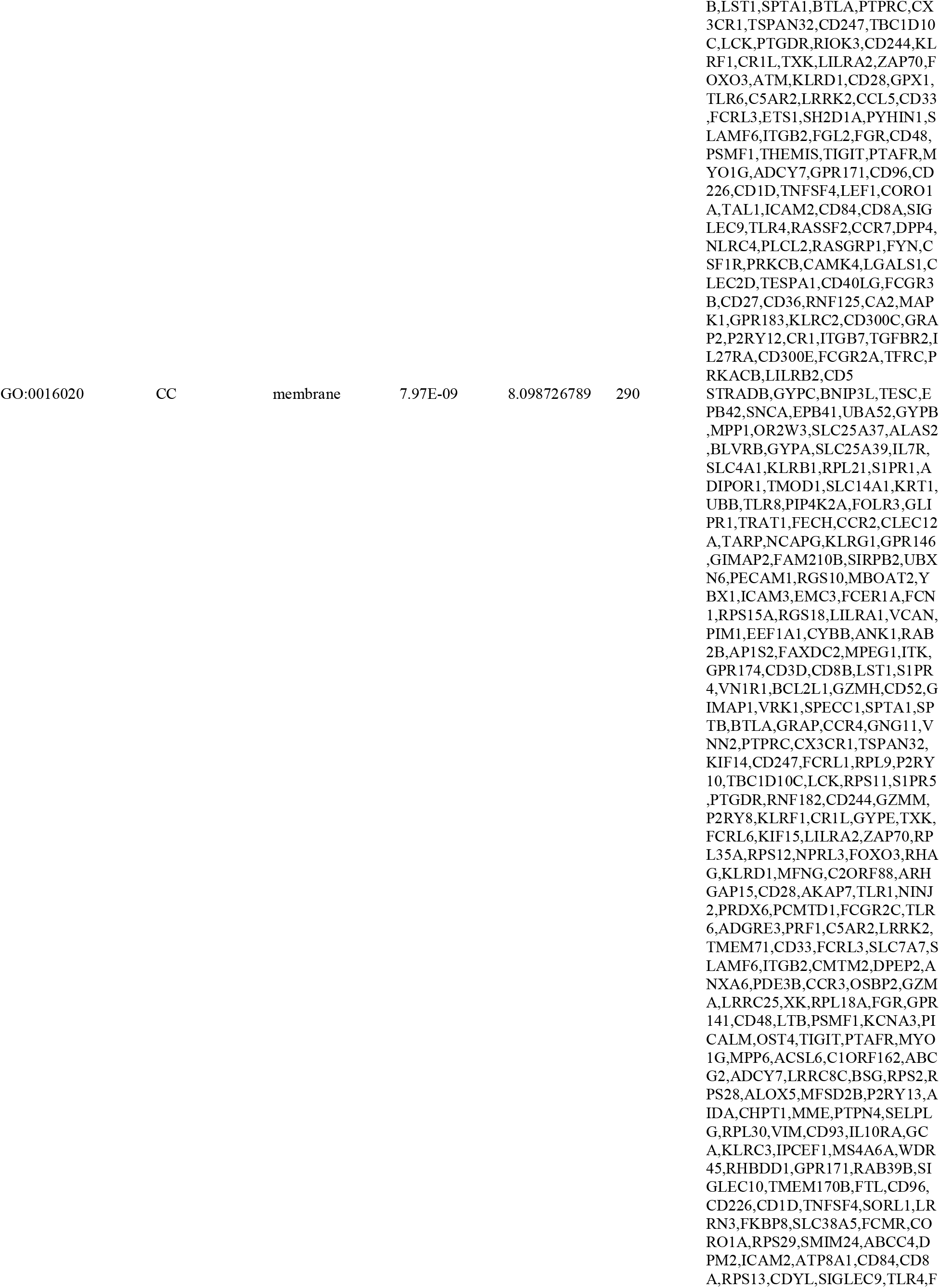
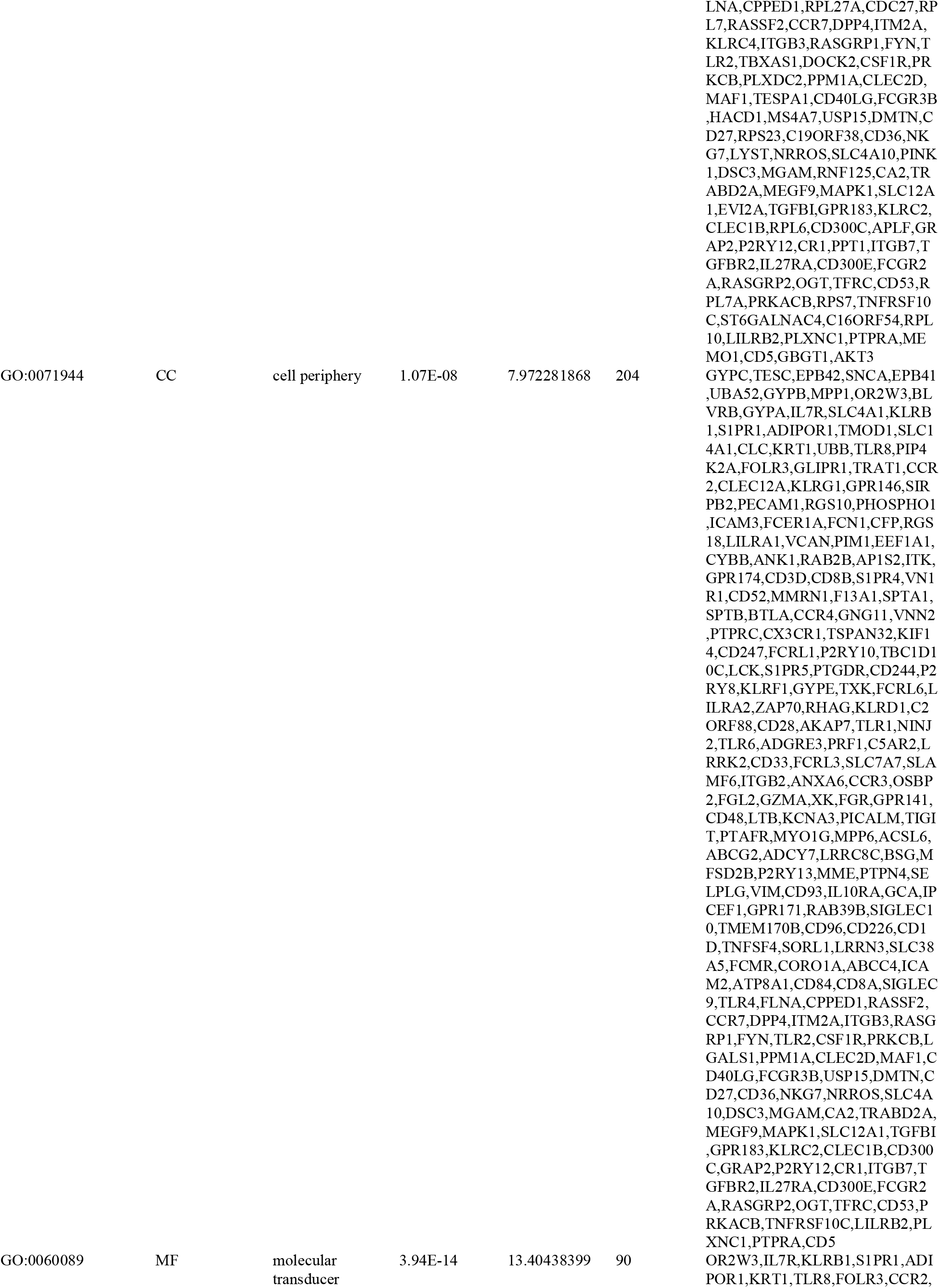
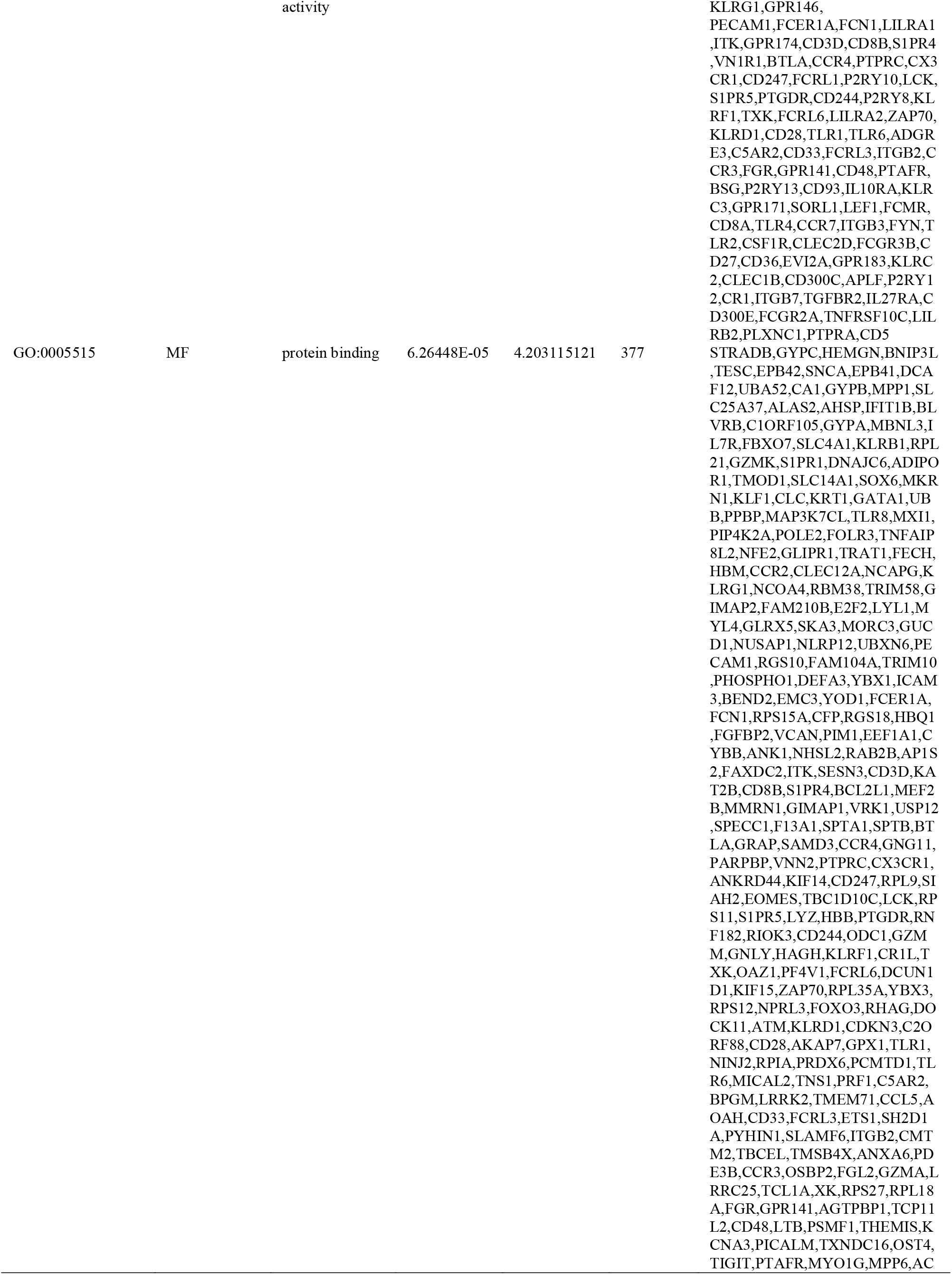
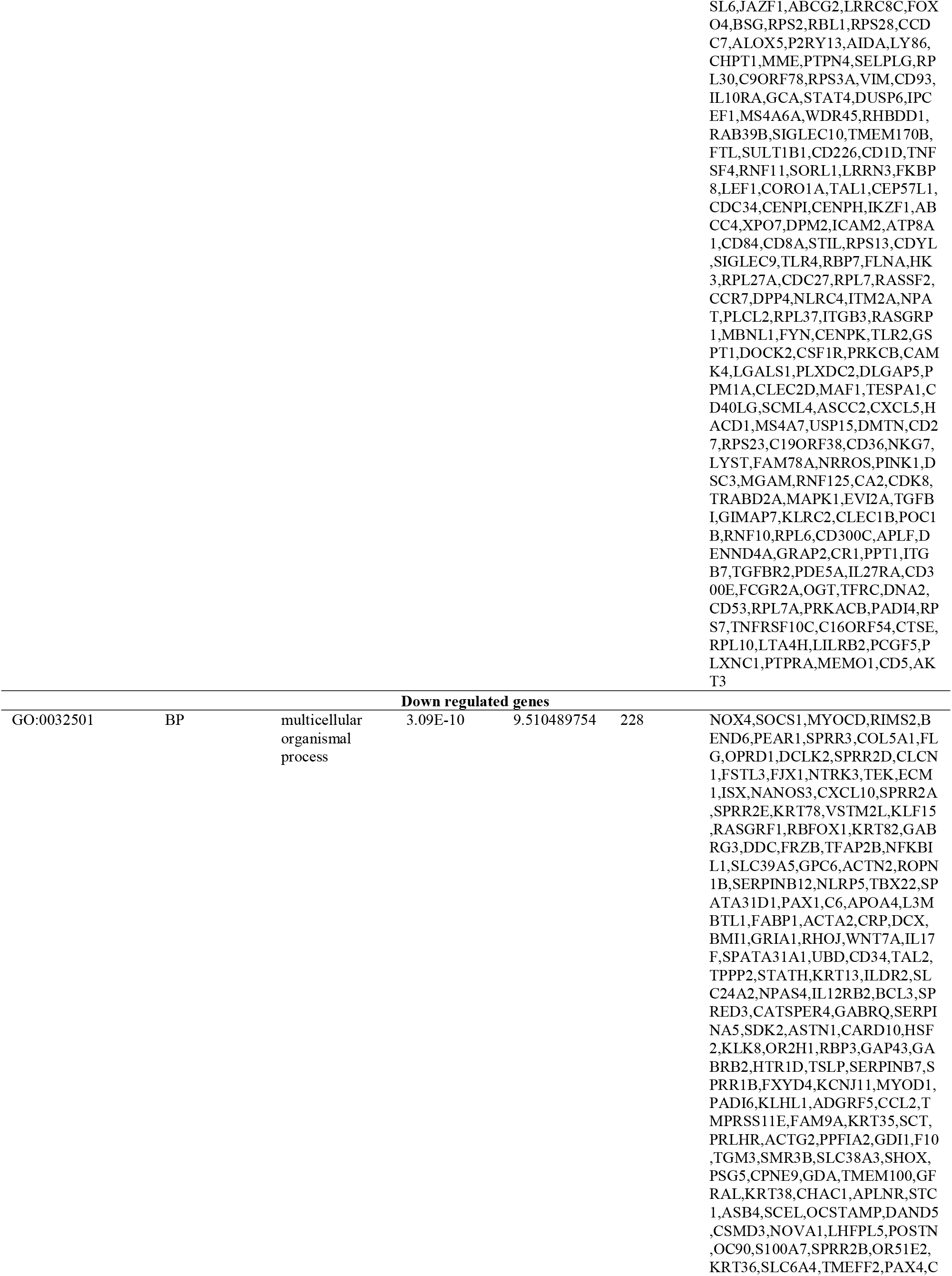
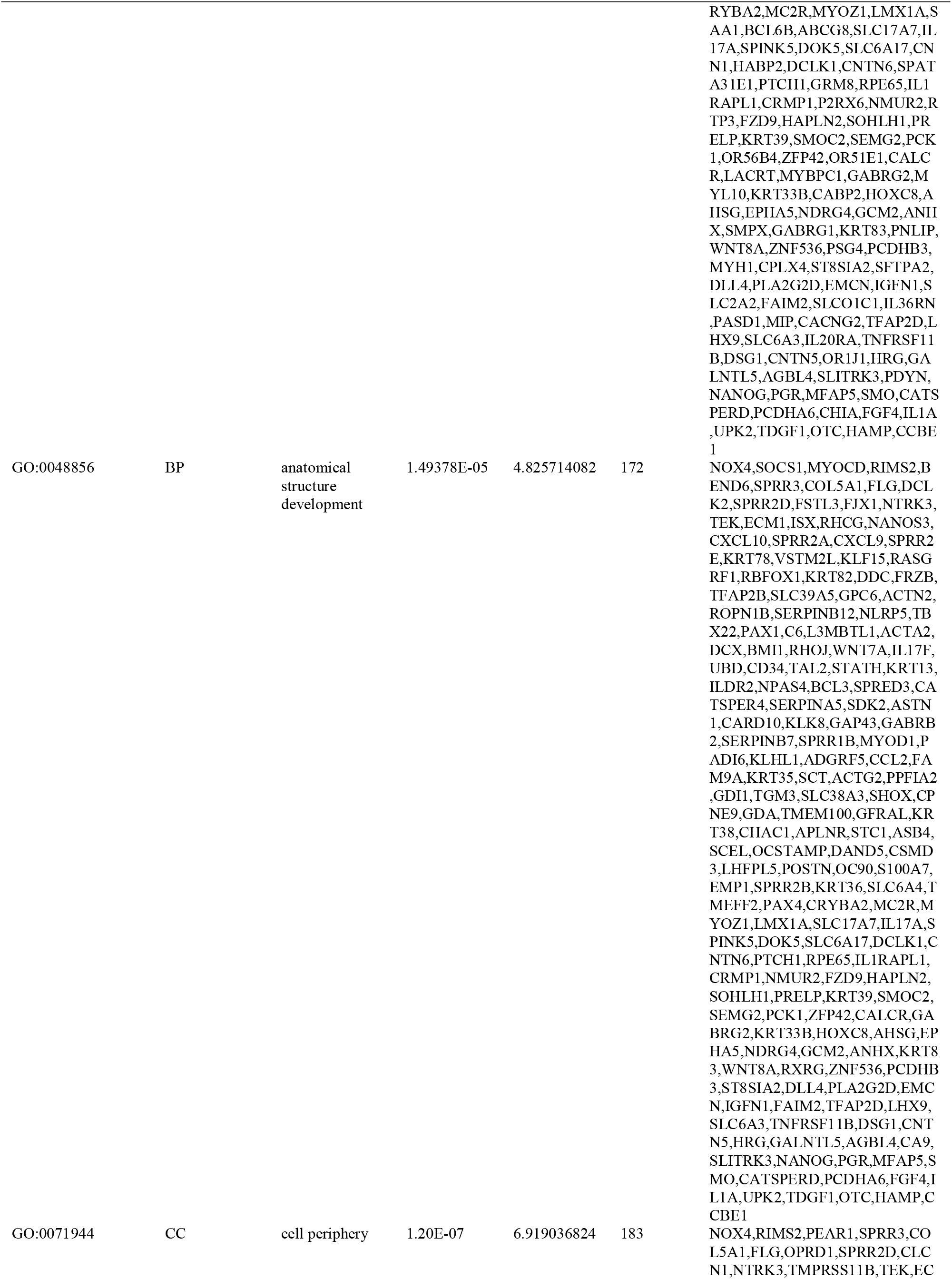
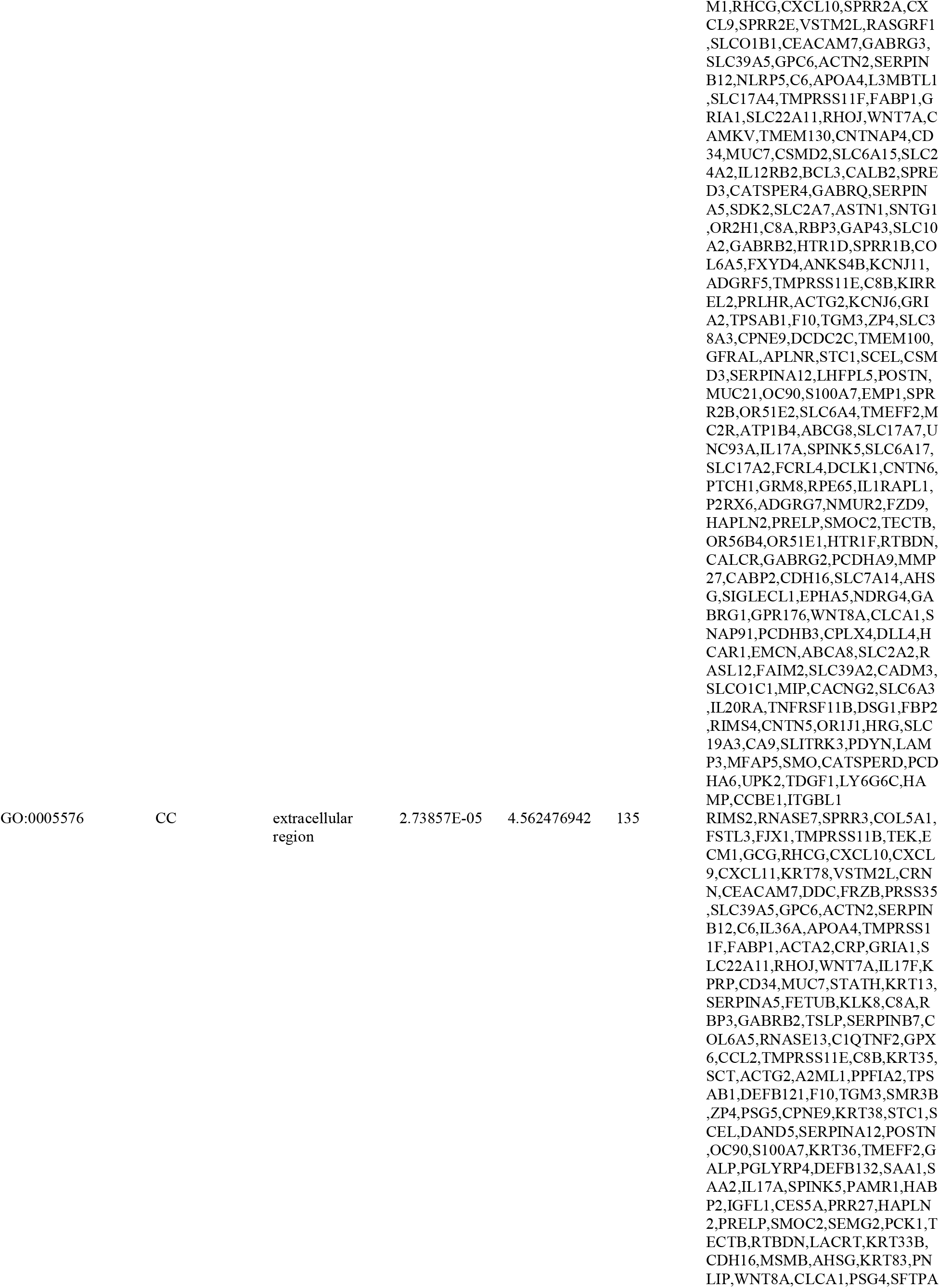
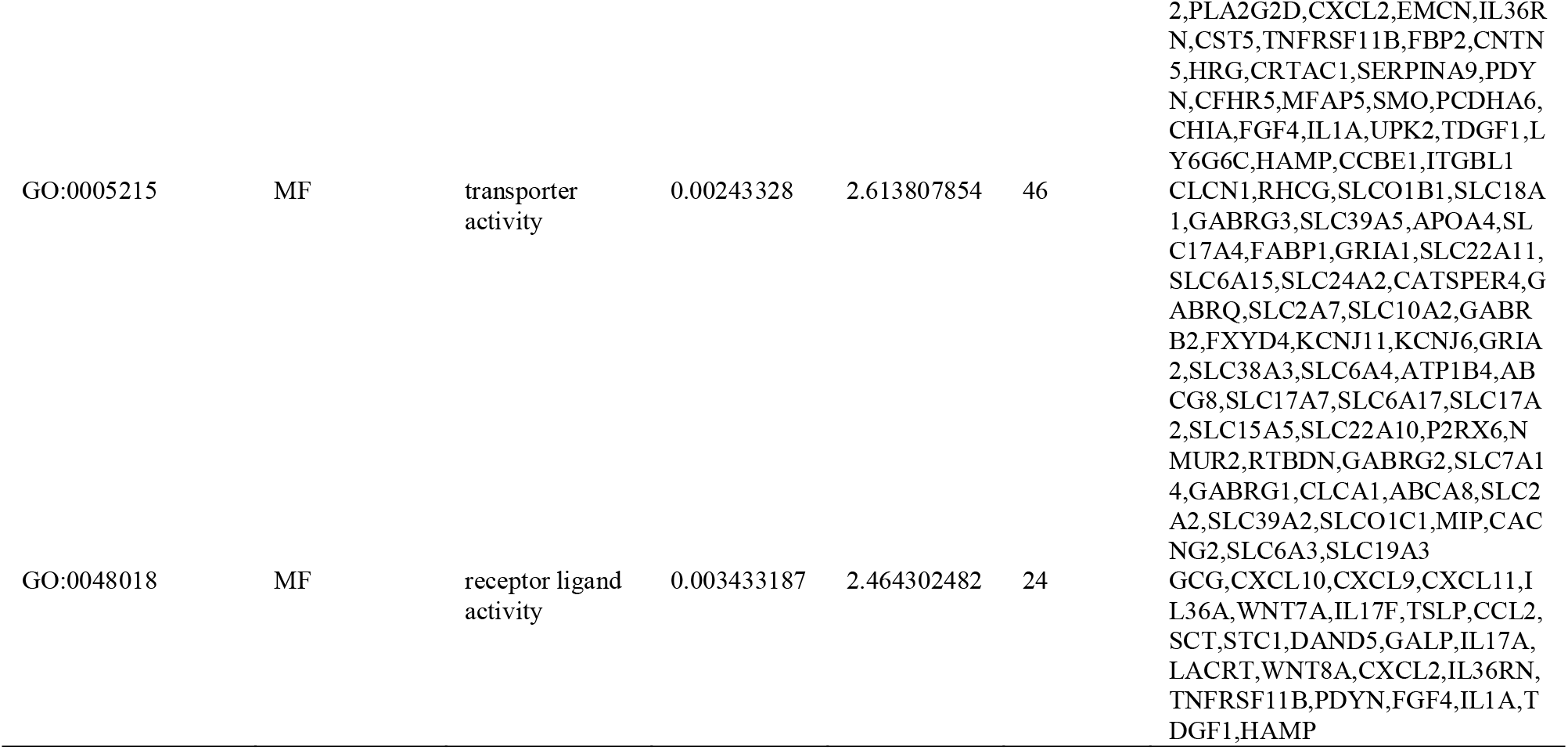
The enriched GO terms of the up and down regulated differentially expressed genes.

**Table 3.**
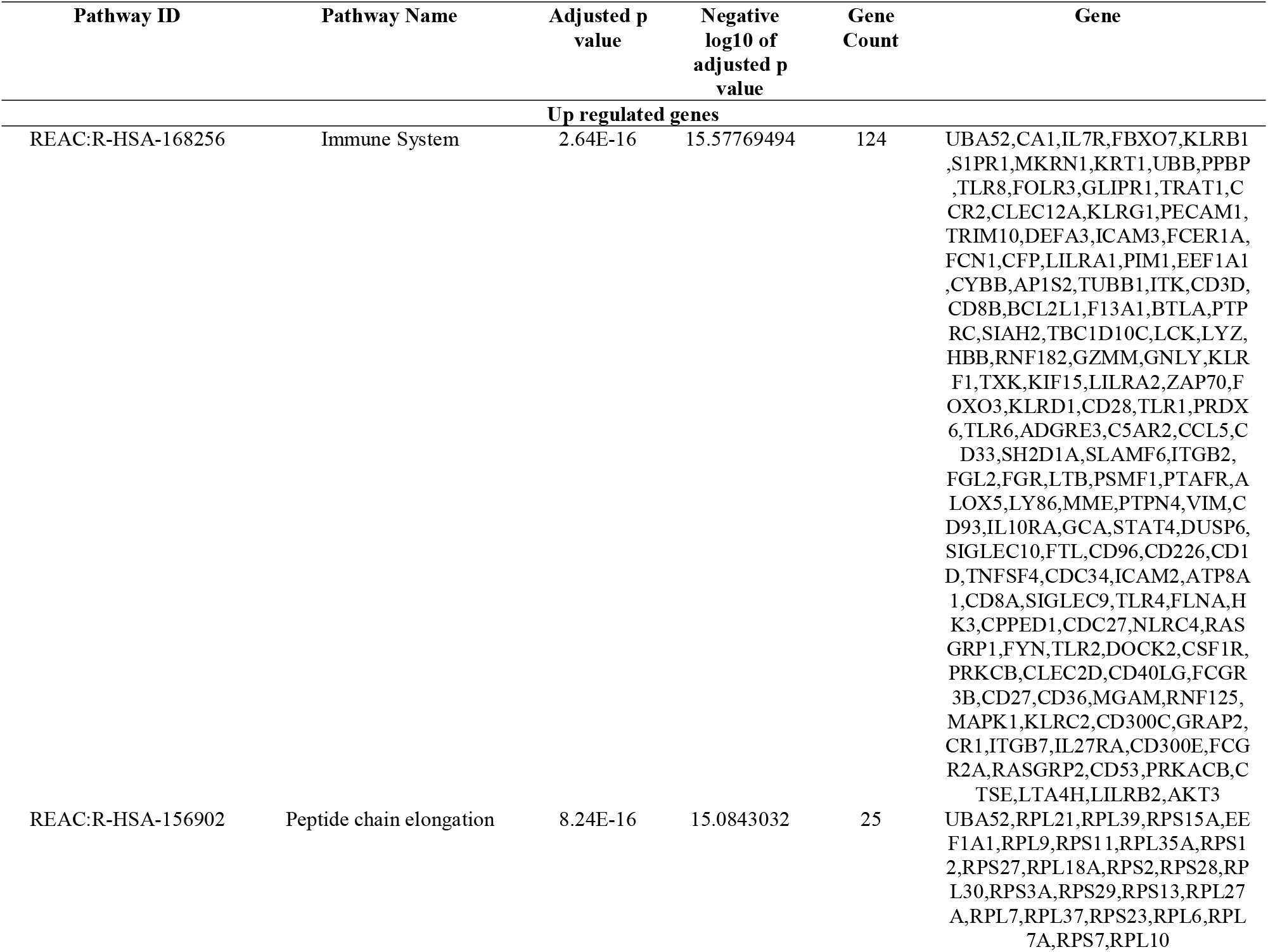
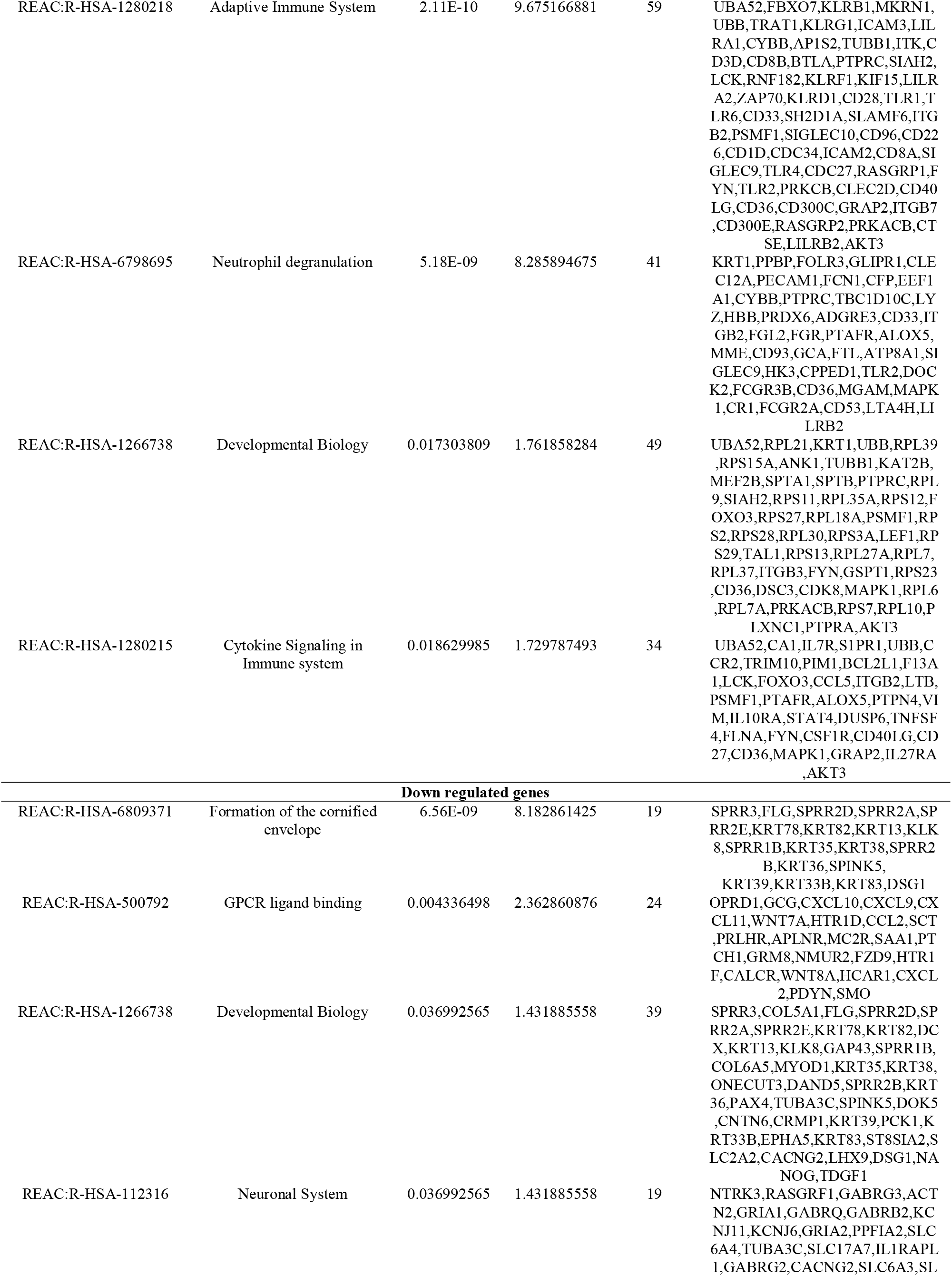
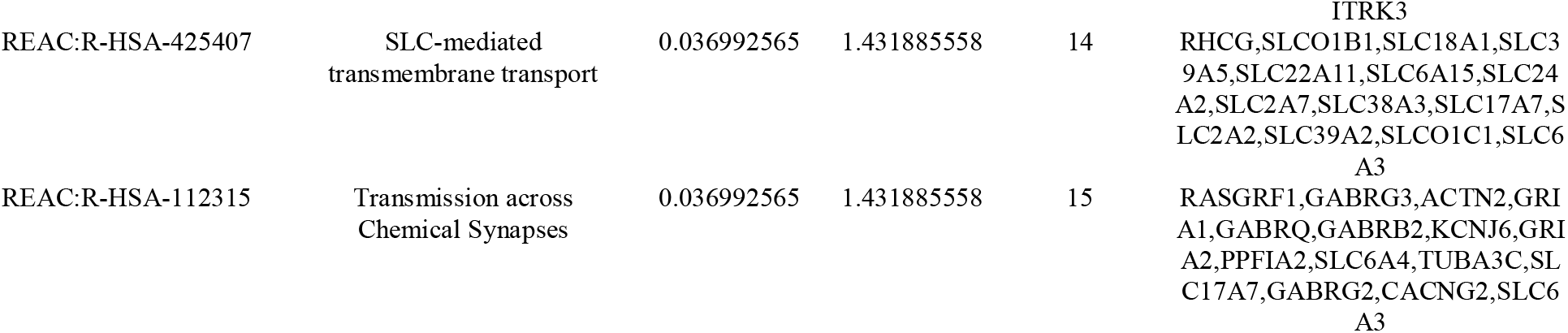
The enriched pathway terms of the up and down regulated differentially expressed genes.

### Construction of the PPI network and module analysis

The PPI network of the DEGs in COVID-19 was constructed based on the information obtained from the IID interactome database. The PPI network included 4020 nodes and 7417 edges, representative of proteins and protein-protein interactions, respectively (Fig. 3). Moreover, RPL10, FYN, FLNA, EEF1A1, UBA52, BMI1, ACTN2, CRMP1, TRIM42 and PTCH1 were identified as hub genes. Table 4 shows the hub genes with the highest node degree, betweenness centrality, stress centrality and closeness centrality, whereas the hub selected using the Network Analyzer plugin. PEWCC1 was used to identify the significant modules in the PPI network and the two significant modules were selected (Fig. 4A and Fig. 4B). Following GO annotation and pathway screening, the module 1 (19 nodes and 41 edges) was revealed to be associated with immune system, adaptive immune system, immune system process, cytokine signaling in immune system and regulation of immune system process, and module 2 (7 nodes and 7 edges) was revealed to be associated with developmental biology, neuronal system, cell periphery and multicellular organismal process.

**Fig. 3.**
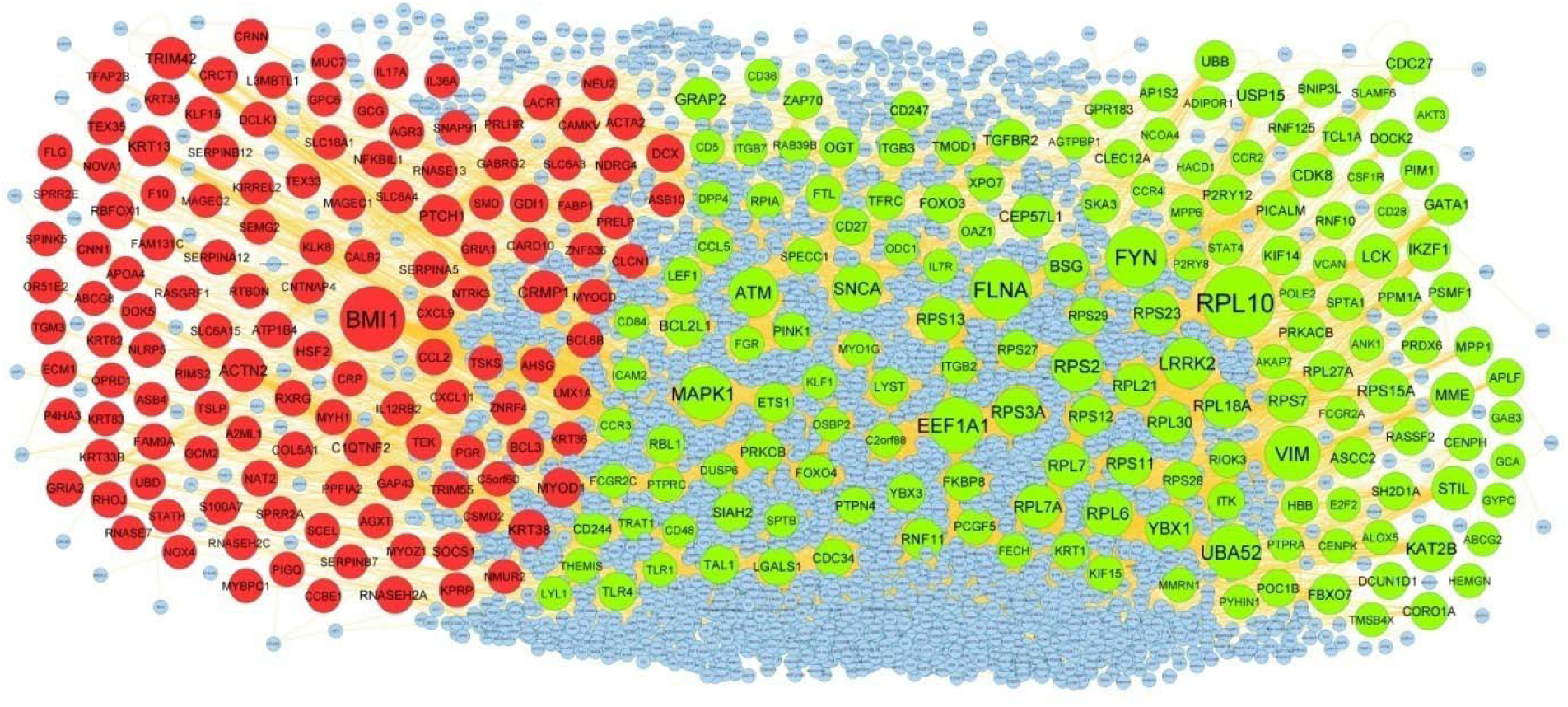
PPI network of DEGs. Up regulated genes are marked in green; down regulated genes are marked in red

**Table 4.**
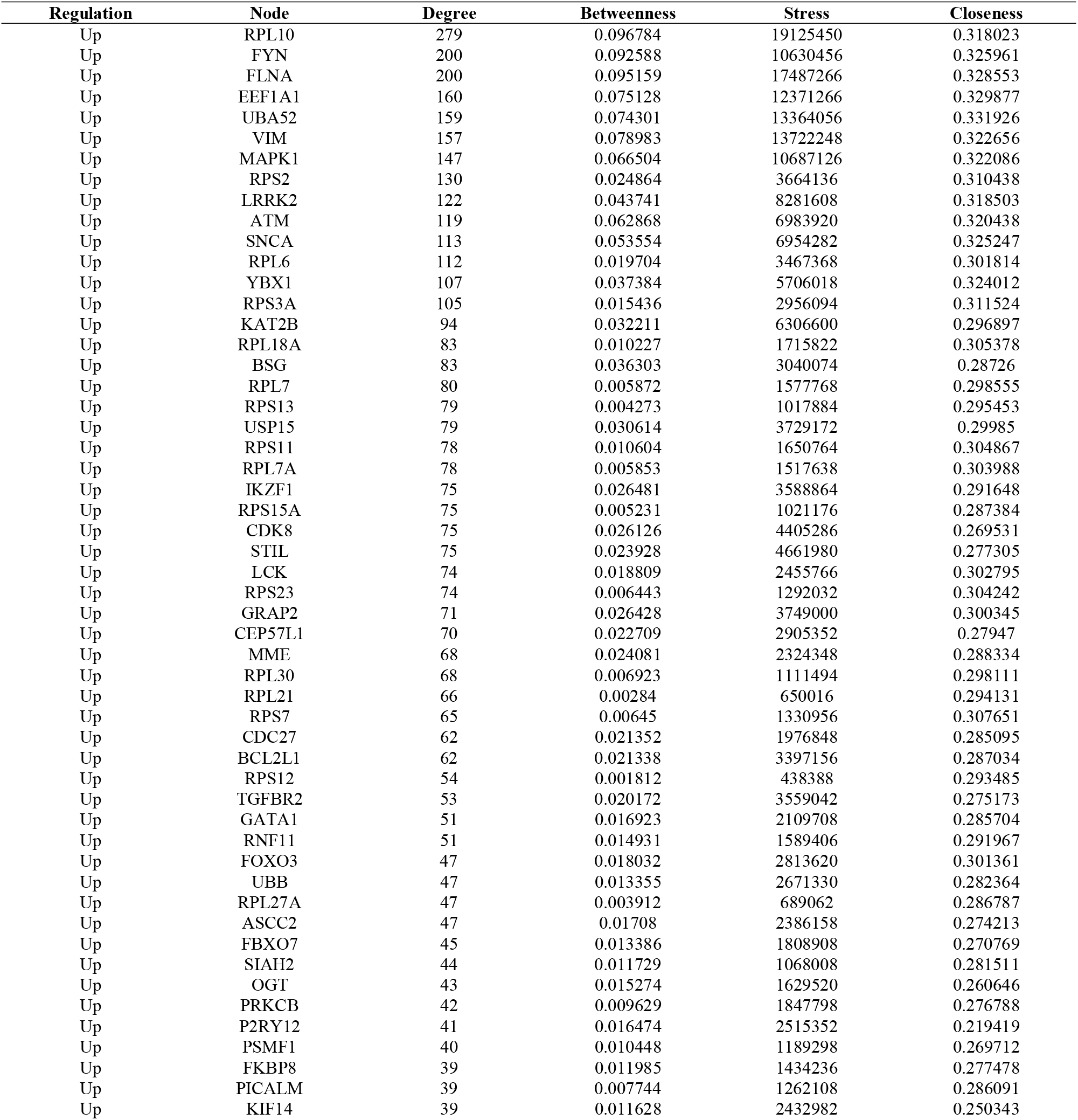
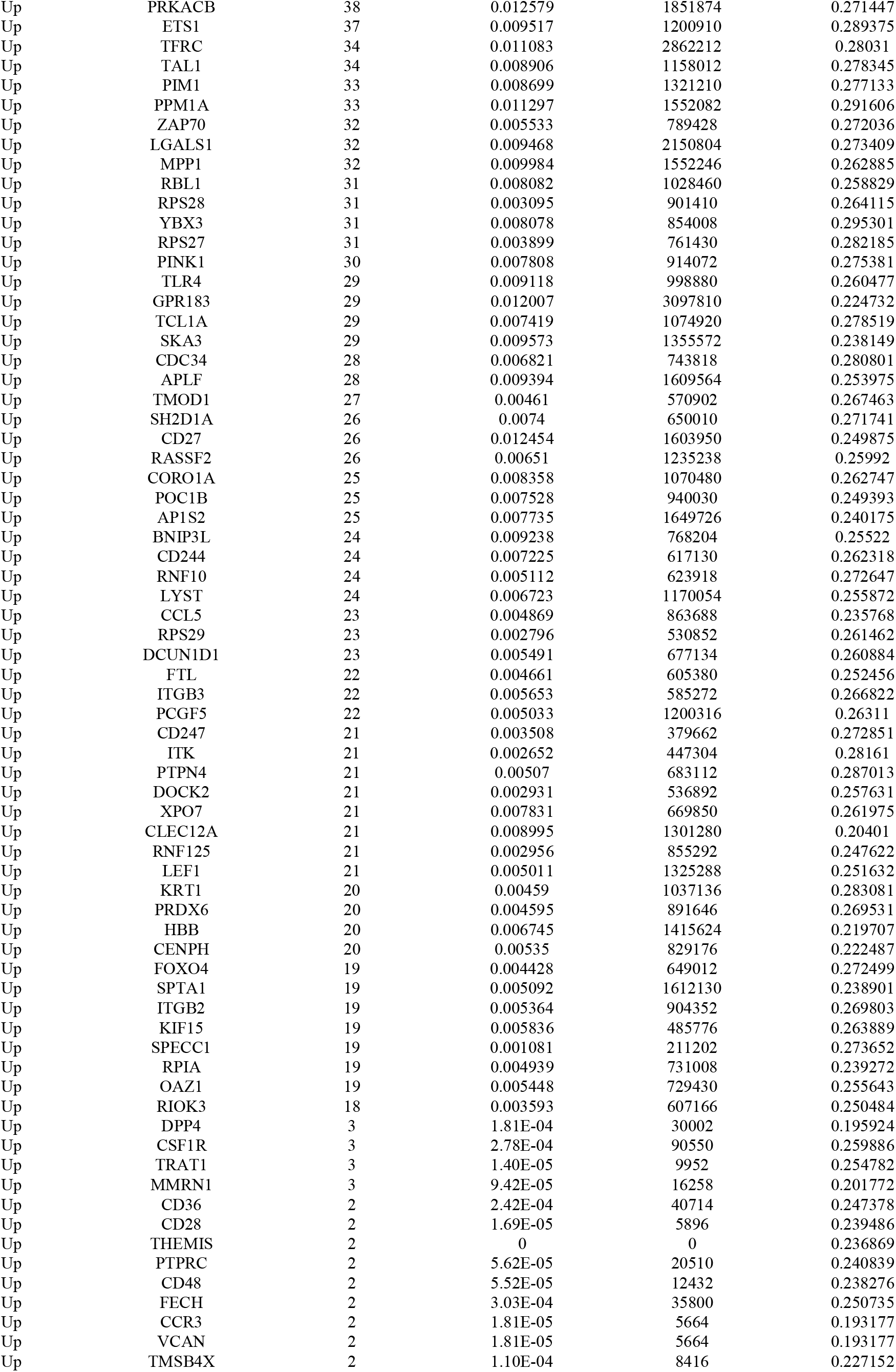
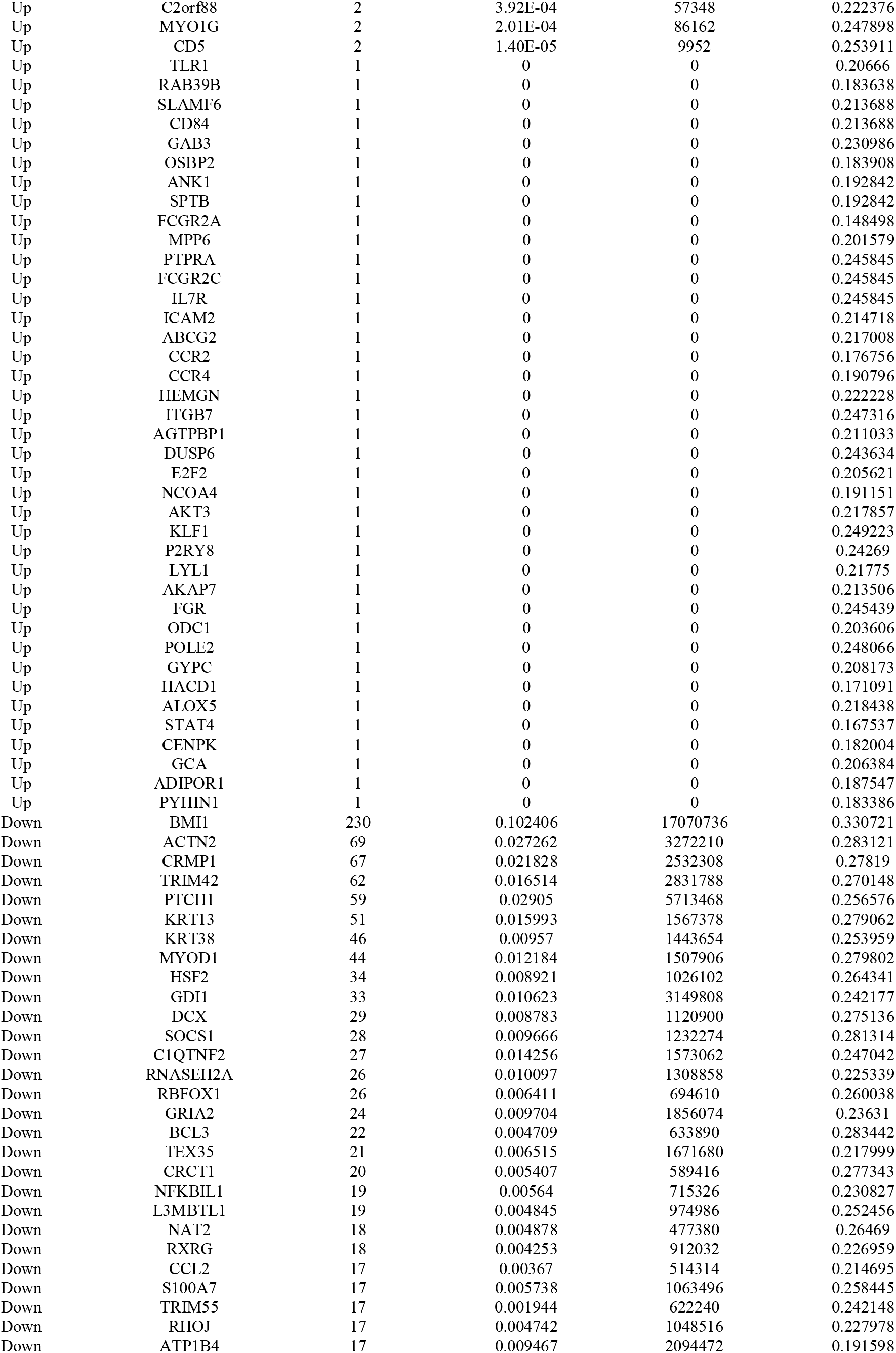
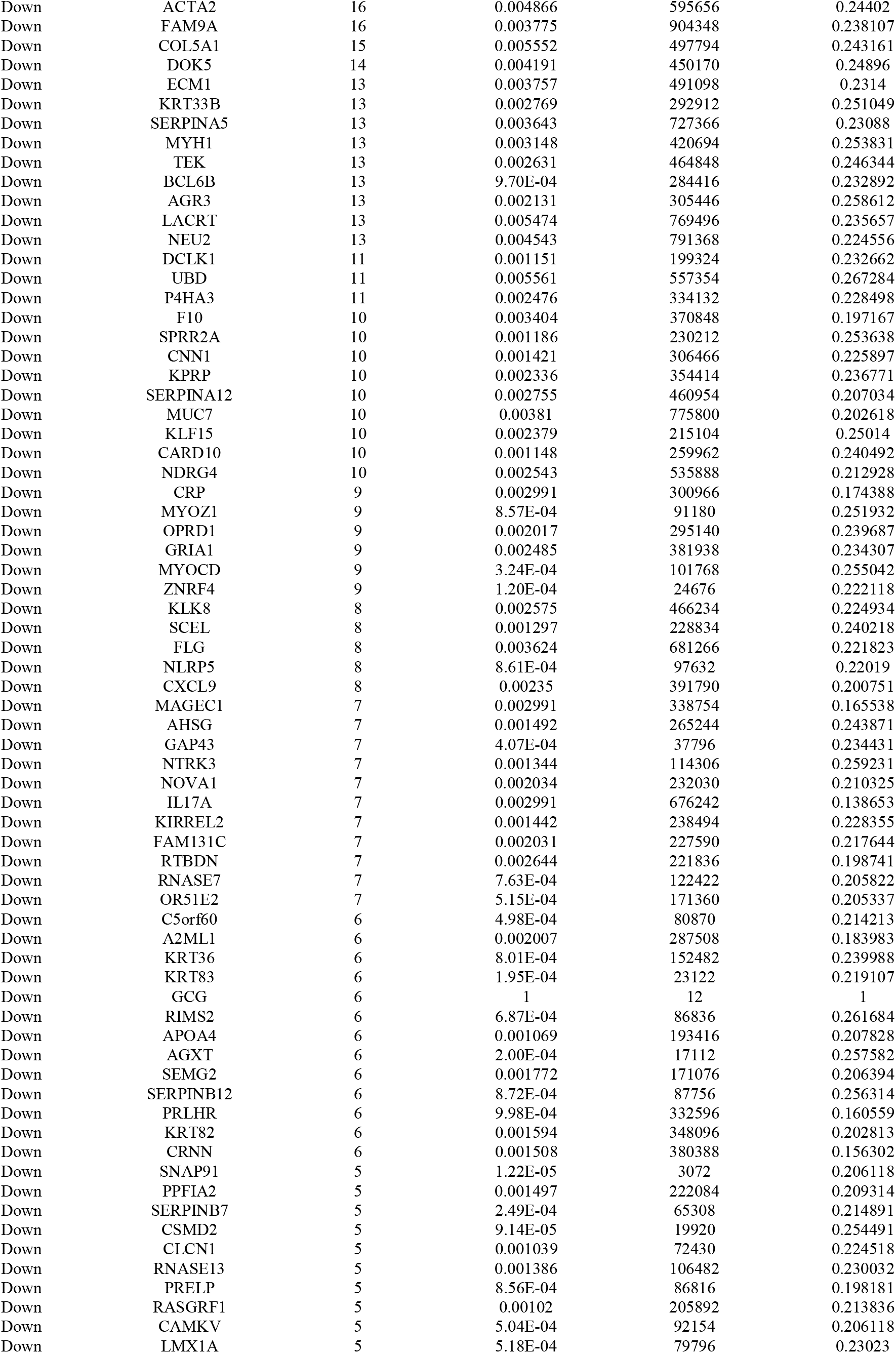
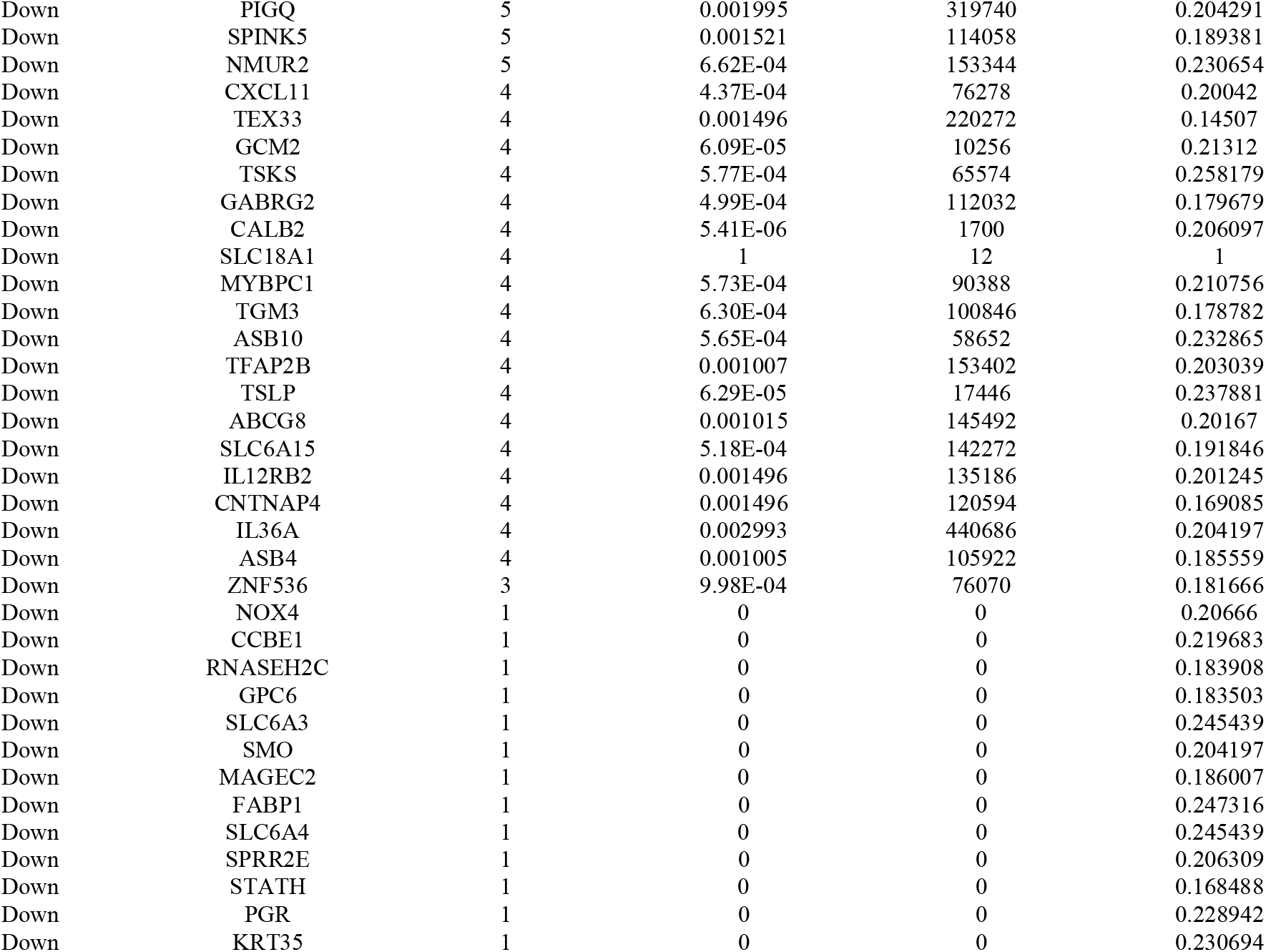
Topology table for up and down regulated genes.

**Fig. 4.**
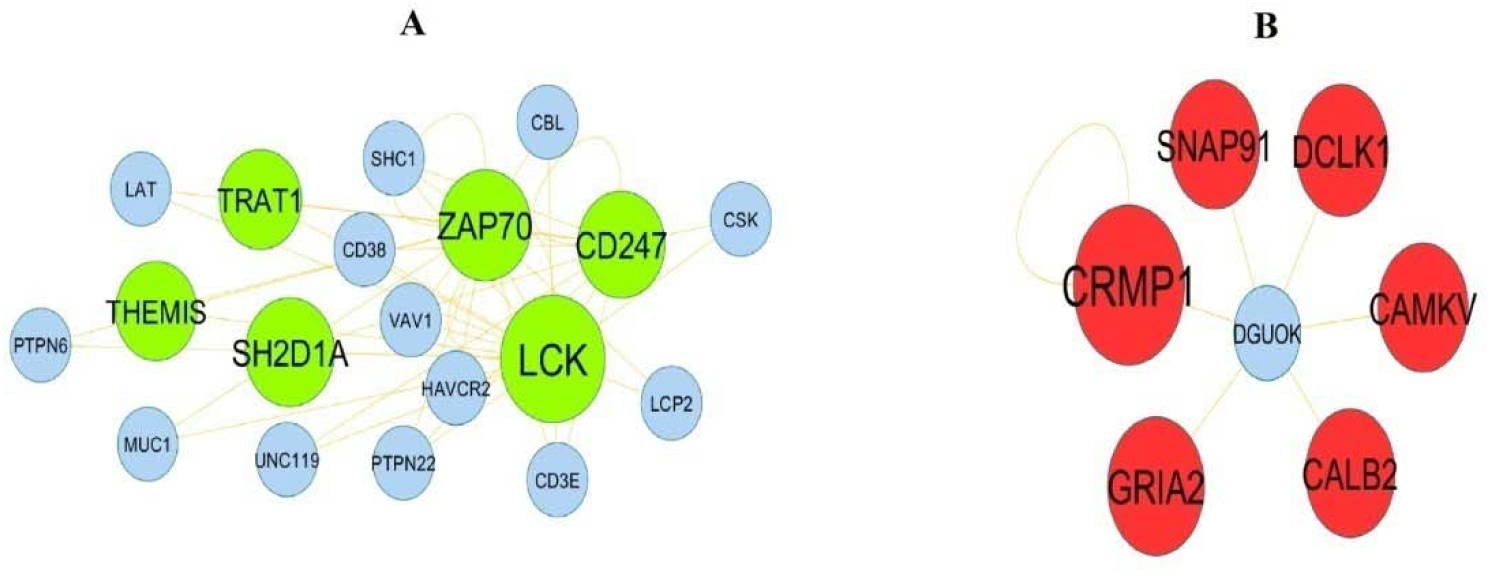
Modules of isolated form PPI of DEGs. (A) The most significant module was obtained from PPI network with 19 nodes and 41 edges for up regulated genes (B) The most significant module was obtained from PPI network with 7 nodes and 7 edges for down regulated genes. Up regulated genes are marked in green; down regulated genes are marked in red

### MiRNA-hub gene regulatory network construction

miRNet was applied to screen the targeted miRNAs of the hub genes. As illustrated in Fig. 5, the interaction network consists of nodes 2287 (293 hub genes and 1994 miRNAs) and 10353 edges. The top targeted hub genes for miRNAs were MAPK1 that was modulated by 286 miRNAs (ex; hsa-mir-320a), ATM that was modulated by 134 miRNAs (ex; hsa-mir-1910-3p), VIM that was modulated by 117 miRNAs (ex; hsa-mir-766-3p), FLNA that was modulated by 106 miRNAs (ex; hsa-mir-124-3p), EEF1A1 that was modulated by 96 miRNAs (ex; hsa-mir-4736), BMI1 that was modulated by 115 miRNAs (ex; hsa-mir-559), GDI1 that was modulated by 104 miRNAs (ex; hsa-mir-186-3p), SOCS1 that was modulated by 62 miRNAs (ex; hsa-mir-4495), PTCH1 that was modulated by 52 miRNAs (ex; hsa-mir-3689a-5p) and DCX that was modulated by 39 miRNAs (ex; hsa-mir-6842-5p), and are listed in Table 5.

**Fig. 5.**
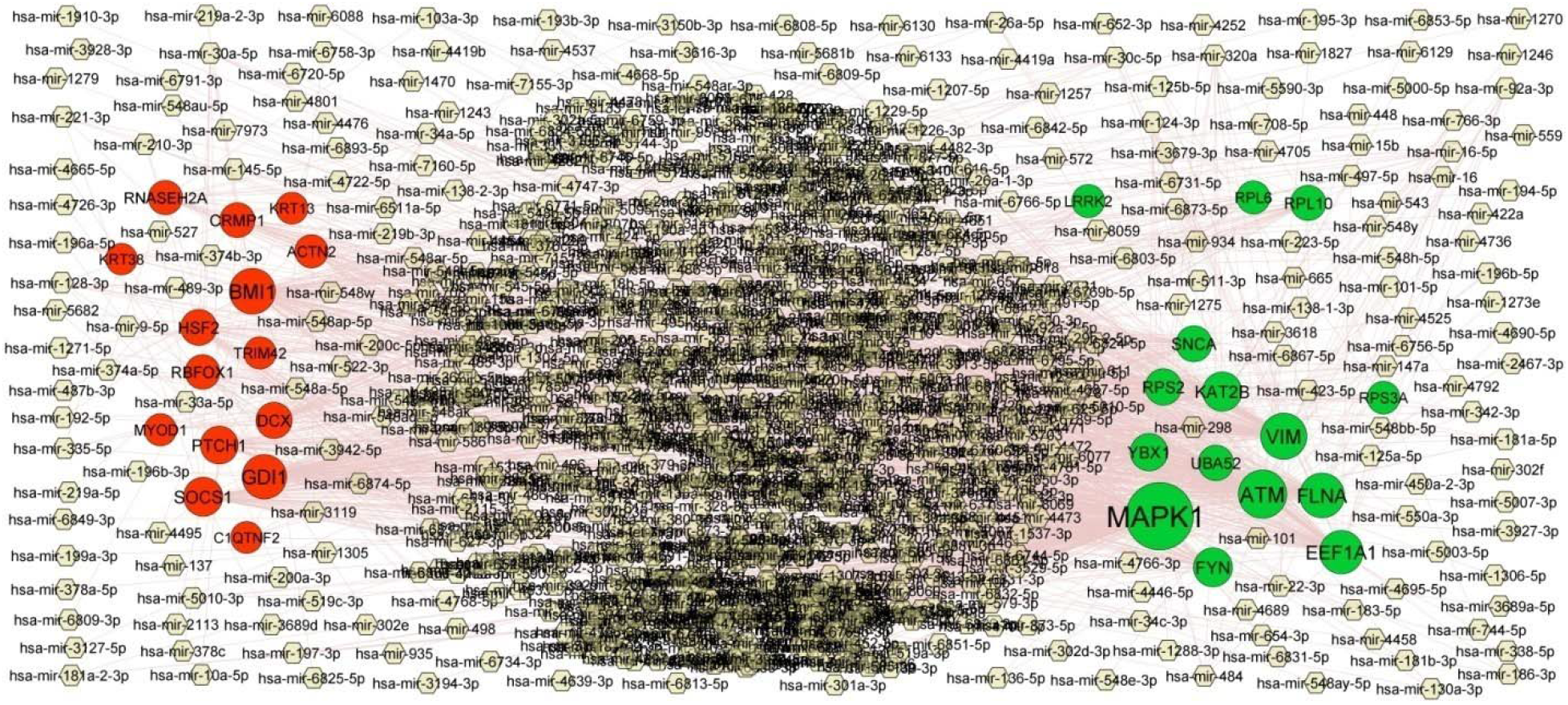
Target gene - miRNA regulatory network between target genes. The white color diamond nodes represent the key miRNAs; up regulated genes are marked in green; down regulated genes are marked in red.

**Table 5.** miRNA - target gene and TF - target gene interaction.

| Regulation | Target Genes | Degree | MicroRNA | Regulation | Target Genes | Degree | TF |
| --- | --- | --- | --- | --- | --- | --- | --- |
| Up | MAPK1 | 286 | hsa-mir-320a | Up | UBA52 | 12 | SREBF1 |
| Up | ATM | 134 | hsa-mir-1910-3p | Up | SNCA | 11 | ELK1 |
| Up | VIM | 117 | hsa-mir-766-3p | Up | FYN | 10 | NFYA |
| Up | FLNA | 106 | hsa-mir-124-3p | Up | ATM | 10 | ELF5 |
| Up | EEF1A1 | 96 | hsa-mir-4736 | Up | FLNA | 10 | TP53 |
| Up | KAT2B | 62 | hsa-mir-1246 | Up | VIM | 9 | POU2F2 |
| Up | FYN | 61 | hsa-mir-147a | Up | YBX1 | 9 | CEBPB |
| Up | YBX1 | 48 | hsa-mir-16-5p | Up | MAPK1 | 9 | USF2 |
| Up | RPS2 | 40 | hsa-mir-34a-5p | Up | RPL6 | 8 | NR3C1 |
| Up | SNCA | 37 | hsa-mir-221-3p | Up | LRRK2 | 8 | FOXC1 |
| Up | UBA52 | 32 | hsa-mir-484 | Up | RPL10 | 7 | TFAP2A |
| Up | RPL10 | 31 | hsa-mir-22-3p | Up | KAT2B | 6 | STAT1 |
| Up | RPL6 | 14 | hsa-mir-193b-3p | Up | RPS3A | 6 | HOXA5 |
| Up | LRRK2 | 14 | hsa-mir-3127-5p | Up | EEF1A1 | 4 | GATA2 |
| Up | RPS3A | 11 | hsa-mir-342-3p | Up | RPS2 | 4 | IRF2 |
| Down | BMI1 | 115 | hsa-mir-559 | Down | RBFOX1 | 19 | EGR1 |
| Down | GDI1 | 104 | hsa-mir-186-3p | Down | PTCH1 | 18 | E2F6 |
| Down | SOCs1 | 62 | hsa-mir-4495 | Down | DCX | 15 | FOXL1 |
| Down | PTCH1 | 52 | hsa-mir-3689a-5p | Down | KRT13 | 12 | STAT3 |
| Down | DCX | 39 | hsa-mir-6842-5p | Down | HSF2 | 12 | NFIC |
| Down | HSF2 | 38 | hsa-mir-26a-5p | Down | TRIM42 | 11 | GATA2 |
| Down | CRMP1 | 29 | hsa-mir-30a-5p | Down | ACTN2 | 11 | USF1 |
| Down | RBFOX1 | 27 | hsa-mir-219a-5p | Down | CRMP1 | 10 | ZNF354C |
| Down | RNASEH2A | 24 | hsa-mir-548h-5p | Down | MYOD1 | 9 | GATA3 |
| Down | ACTN2 | 16 | hsa-mir-3119 | Down | RNASEH2A | 8 | PAX2 |
| Down | MYOD1 | 8 | hsa-mir-101-3p | Down | KRT38 | 7 | HINFP |
| Down | TRIM42 | 8 | hsa-mir-6867-5p | Down | GDI1 | 6 | SOX5 |
| Down | C1QTNF2 | 5 | hsa-mir-520c-3p | Down | BMI1 | 6 | SP1 |
| Down | KRT13 | 5 | hsa-mir-1343-3p | Down | C1QTNF2 | 4 | PRDM1 |
| Down | KRT38 | 2 | hsa-mir-335-5p | Down | SOCS1 | 2 | EN1 |

### TF-hub gene regulatory network construction

NetworkAnalyst was applied to screen the targeted TFs of the hub genes. As illustrated in Fig. 6, the interaction network consists of nodes 379 (294 hub genes and 85 TFs) and 2385 edges. The top targeted hub genes for TFs were UBA52 that was modulated by 12 TFs (ex; SREBF1), SNCA that was modulated by 11 TFs (ex; ELK1), FYN that was modulated by 10 TFs (ex; NFYA), ATM that was modulated by 10 TFs (ex; ELF5), FLNA that was modulated by 10 TFs (ex; TP53), RBFOX1 that was modulated by 19 TFs (ex; EGR1), PTCH1 that was modulated by 18 TFs (ex; E2F6), DCX that was modulated by 15 TFs (ex; FOXL1), KRT13 that was modulated by 12 TFs (ex; STAT3) and HSF2 that was modulated by 12 TFs (ex; NFIC), and are listed in Table 5.

**Fig. 6.**
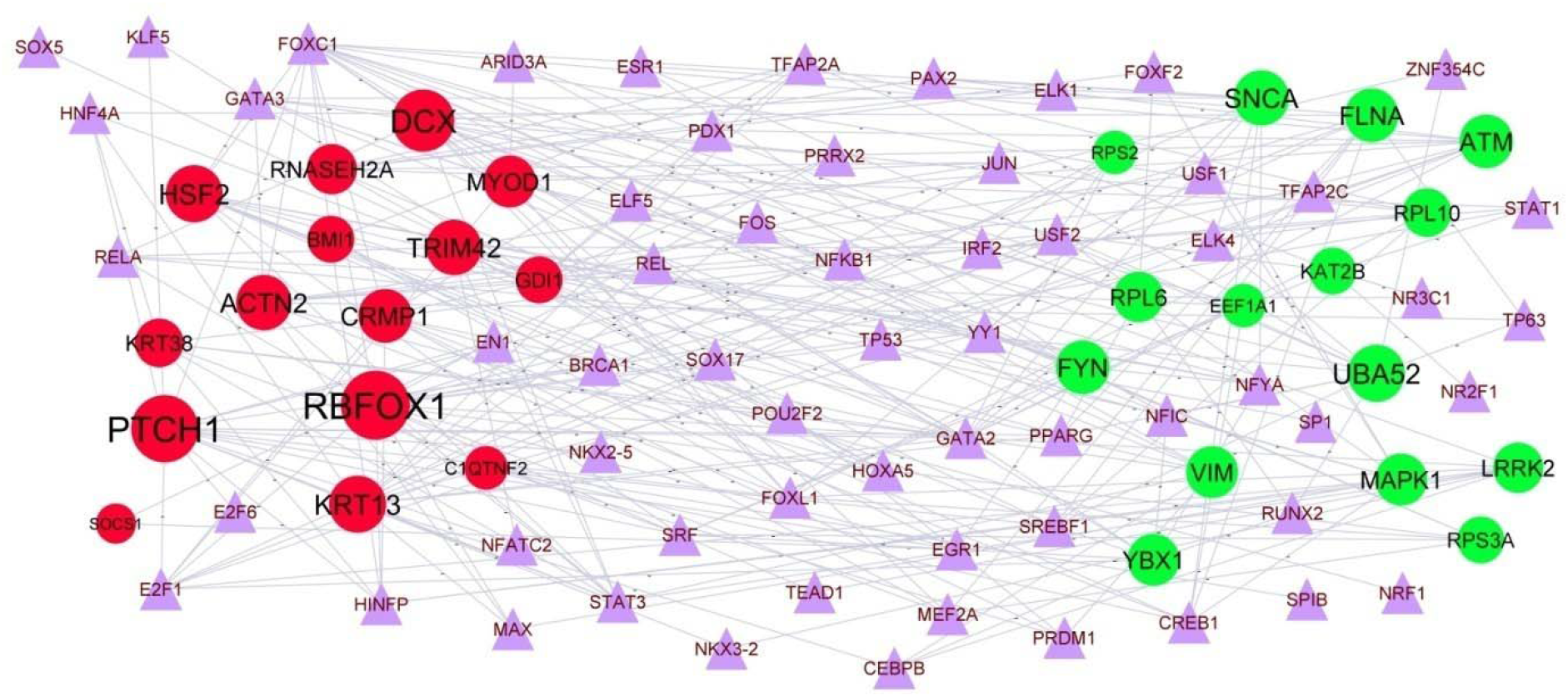
Target gene - TF regulatory network between target genes. The purple color triangle nodes represent the key TFs; up regulated genes are marked in green; down regulated genes are marked in red.

### Receiver operating characteristic curve (ROC) analysis

To identify novel biomarkers for COVID-19, ROC curves of data derived from COVID-19 and non COVID-19 from dataset GSE163151 was analyzed using the pROC package in R statistical software (Fig.7). ROC curves were generated, and the AUC was used to compare the different hub genes. RPL10 (AUC = 0.938), FYN (AUC = 0.960), FLNA (AUC = 0.944), EEF1A1 (AUC = 0.910), UBA52 (AUC = 0.941), BMI1 (AUC = 0.951), ACTN2 (AUC = 0.926), CRMP1 (AUC = 0.954), TRIM42 (AUC = 0.895) and PTCH1 (AUC = 0.923) have the largest AUC area. These findings suggest that RPL10, FYN, FLNA, EEF1A1, UBA52, BMI1, ACTN2, CRMP1, TRIM42 and PTCH1 are the most relevant biomarker for COVID-19.

**Fig. 7.**
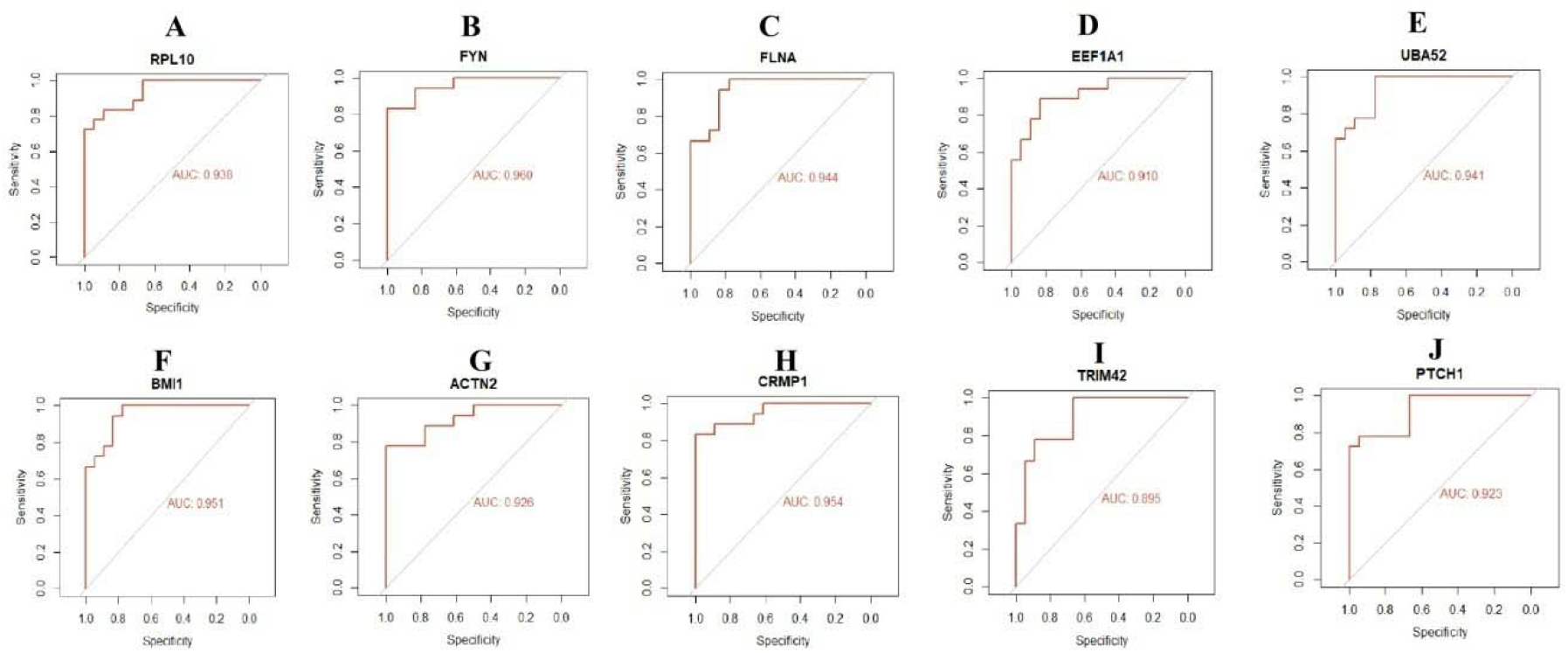
ROC curve analyses of hub genes. A) RPL10 B) FYN C) FLNA D) EEF1A1 E) UBA52 F) BMI1 G) ACTN2 H) CRMP1 I) TRIM42 J) PTCH1

## Discussion

COVID-19 infection and its secondary complications are the one of the leading causes of death worldwide and thus place heavy burdens on the healthcare systems and global economy [55]. However, the molecular mechanisms of COVID-19 infection and its secondary complications remain poorly understood. Most cases of COVID-19 infection and its secondary complications without an early discovery are not candidates for curative therapies, which might be one of the reasons for the poor patient prognosis. Thus, molecular markers for diagnosis and treatment with high performance are urgently demanded. NGS technology enables us to search the genetic modifications in COVID-19 infection and its secondary complications, and has been proved to be a useful approach to identify novel molecular markers in other diseases.

In the current investigation, NGS dataset was analyzed to obtain DEGs between COVID-19 and non COVID-19. A total of 954 DEGs were identified, including 477 up regulated genes and 477 down regulated genes. A recent study found that the altered expression of BNIP3L [56], NOX4 [57] and SOCS1 [58] are crucial for the progression of diseases related to lungs, but these genes might be novel targets for COVID-19. BNIP3L [59], NOX4 [60] and SOCS1 [61] plays an important role in the renal diseases, but these genes might be involved in the progression of COVID-19. BNIP3L [62], HBA2 [63], SNCA (synuclein alpha) [64], NAT2 [65] and DAO (D-amino acid oxidase) [66] have been reported to have a crucial role in CNS disorders, but these genes might be involved in the progression of COVID-19. Ouyang et al. [67], Vendrov et al. [68], Alikhah et al. [69], Khelil et al. [70] and Xia et al. [71] found that BNIP3L, NOX4, SOCS1, NAT2, MYOCD (myocardin) and expression were altered in cardiovascular diseases, but these genes might be involved in the progression of COVID-19. HBA2 [72], NOX4 [73], NAT2 [74] and MYOCD (myocardin) [75] are important in the development of hypertension, but these genes might be involved in the progression of COVID-19. NOX4 [76], SOCS1 [69] and NAT2 [77] expressions were shown to play an important role in diabetes mellitus, but these genes might be involved in the progression of COVID-19. NOX4 [78] and SOCS1 [79] were reported to be associated with liver injury, but these genes might be novel targets for COVID-19. NOX4 [80] and SOCS1 [81] were shown to participate in facilitating the secondary infections. A previous study revealed that SOCS1 [82] are involved in progression COVID-19 infection.

GO and REACTOME pathway enrichment analyses were performed to explore interactions among the DEGs. Pathways include immune system [83], adaptive immune system [84], neutrophil degranulation [85], cytokine signaling in immune system [86] and GPCR ligand binding [87] were responsible for progression of COVID-19 infection. A previous study demonstrated that the expression levels of AHSP (alpha hemoglobin stabilizing protein) [88], IL7R [89], FBXO7 [90], KLRB1 [91], PIP4K2A [92], NFE2 [93], CCR2 [94], CLEC12A [95], NLRP12 [96], PECAM1 [97], TRIM10 [98], ICAM3 [99], EEF1A1 [100], CCR4 [101], PTPRC (protein tyrosine phosphatase receptor type C) [102], CX3CR1 [103], TSPAN32 [104], EOMES (eomesodermin) [105], ATM (ATM serine/threonine kinase) [106], CD28 [107], LRRK2 [108], CCL5 [109], CD33 [110], FCRL3 [111], CCR3 [112], FGL2 [113], GZMA (granzyme A) [114], PICALM (phosphatidylinositol binding clathrin assembly protein) [115], ALOX5 [116], MME (membrane metalloendopeptidase) [117], VIM (vimentin) [118], CD93 [119], GCA (grancalcin) [120], CD226 [121], CD1D [122], TNFSF4 [123], LEF1 [124], TLR4 [125], CCR7 [126], DPP4 [127], NLRC4 [128], ITGB3 [129], RASGRP1 [130], TLR2 [131], DOCK2 [132], CSF1R [133], PRKCB (protein kinase C beta) [134], CAMK4 [135], CXCL5 [136], CD36 [137], P2RY12 [138], LILRB2 [139], CD5 [140], SLC25A37 [141], ADIPOR1 [142], PECAM1 [143], RGS10 [144], RGS18 [145], ANK1 [146], RNF182 [147], NPRL3 [148], NINJ2 [149], KCNA3 [150], ABCG2 [151], MS4A6A [152], WDR45 [153], RAB39B [154], SORL1 [155], LRRN3 [156], DPM2 [157], SLC4A10 [158], PINK1 [159], RPL6 [160], STAT4 [161], DUSP6 [162], MYOCD (myocardin) [163], OPRD1 [164], CLCN1 [165], CXCL10 [166], KLF15 [167], RBFOX1 [168], DDC (dopa decarboxylase) [169], APOA4 [170], BMI1 [171], GRIA1 [172], SLC24A2 [173], NPAS4 [174], GAP43 [175], GABRB2 [176], TSLP (thymic stromal lymphopoietin) [177], PADI6 [178], CCL2 [179], STC1 [180], SLC6A4 [181] IL17A [182], CNTN6 [183], GRM8 [184], CRMP1 [185], HAPLN2 [186], PCK1 [187], GABRG2 [188], EPHA5 [189], ST8SIA2 [190], FAIM2 [191], SLC6A3 [192], PDYN (prodynorphin) [193], HAMP (hepcidin antimicrobial peptide) [194], CXCL9 [195], CSMD2 [196], SLC6A15 [197], SLC10A2 [198], ATP1B4 [199], SIGLECL1 [200] and SLC18A1 [201] were altered in patients with CNS disorders, but these genes might be involved in the progression of COVID-19. IL7R [202], S1PR1 [203], NFE2 [204], CCR2 [205], NLRP12 [206], PECAM1 [207], PIM1 [208], AP1S2 [209], ITK (IL2 inducible T cell kinase) [210], CCR4 [211], CX3CR1 [212], FOXO3 [213], CD28 [214], GPX1 [215], PRDX6 [216], LRRK2 [217], CCL5 [218], ETS1 [219], CCR3 [220], SELPLG (selectin P ligand) [221], VIM (vimentin) [222], CD1D [223], SIGLEC9 [224], TLR4 [225], FLNA (filamin A) [226], CCR7 [227], DPP4 [228], NLRC4 [229], TLR2 [230], DOCK2 [231], CD40LG [232], CD36 [233], MAPK1 [234], TGFBR2 [235], LTA4H [236], PECAM1 [237], VCAN (versican) [238], NPRL3 [239], CDC27 [240], PINK1 [241], TGFBI (transforming growth factor beta induced) [242], CR1 [243], AKT3 [244], STAT4 [245], CXCL10 [246], CRP (C-reactive protein) [247], IL17F [248], CD34 [249], BCL3 [250], TSLP (thymic stromal lymphopoietin) [251], CCL2 [252], F10 [253], STC1 [254], POSTN (periostin) [255], IL17A [256], SMOC2 [257], SFTPA2 [258], FAIM2 [259], LHX9 [260], HRG (histidine rich glycoprotein) [261] and CA9 [262] have been reported to be related to lung diseases, but these genes might be involved in the progression of COVID-19. SLC4A1 [263], CCR2 [264], FGL2 [265], CD84 [266], TLR4 [267], ITGB3 [268], TLR2 [267], CD36 [269], MFSD2B [270], PLXDC2 [271], KLF15 [272], C6 [273] and HRG (histidine rich glycoprotein) [274] plays important roles in blood clotting processes. Altered expression of SLC4A1 [275], S1PR1 [276], NFE2 [277], CCR2 [278], NLRP12 [279], PECAM1 [280], ICAM3 [281], CX3CR1 [282], CD247 [283], FOXO3 [284], CD28 [285], GPX1 [286], TLR1 [287], PRDX6 [288], CCL5 [289], FCRL3 [290], ETS1 [291], FGL2 [292], CD93 [293], CD1D [294], TNFSF4 [295], TAL1 [296], TLR4 [297], CCR7 [298], DPP4 [299], NLRC4 [300], RASGRP1 [301], FYN (FYN proto-oncogene, Src family tyrosine kinase) [302], TLR2 [303], CAMK4 [304], FCGR3B [305], CXCL5 [306], CD27 [307], CD36 [308], MGAM (maltase-glucoamylase) [309], CA2 [310], TGFBR2 [311], PADI4 [312], CD5 [313], ADIPOR1 [314], VCAN (versican) [315], SPTB (spectrin beta, erythrocytic) [316], ABCG2 [317], PINK1 [318], OGT (O-linked N-acetylglucosamine (GlcNAc) transferase) [319], AKT3 [320], STAT4 [321], FSTL3 [322], CXCL10 [323], KLF15 [324], FABP1 [325], ACTA2 [326], CRP (C-reactive protein) [327], BMI1 [328], WNT7A [329], IL17F [330], UBD (ubiquitin D) [331], CD34 [332], BCL3 [333], SERPINA5 [334], ADGRF5 [335], CCL2 [336], SCT (secretin) [337], CHAC1 [338], STC1 [339], POSTN (periostin) [340], SAA1 [341], ABCG8 [342], IL17A [343], PTCH1 [344], SMOC2 [345], AHSG (alpha 2-HS glycoprotein) [346], CXCL9 [347], GPR176 [348] and CLCA1 [349] contributes to the progression of renal diseases, but these genes might be involved in the progression of COVID-19. S1PR1 [350], CCR2 [351], CLEC12A [352], KLRG1 [353], NLRP12 [354], PECAM1 [355], ICAM3 [356], PIM1 [357], EEF1A1 [358], MPEG1 [359], CCR4 [360], CX3CR1 [361], EOMES (eomesodermin) [362], LYZ (lysozyme) [363], CD244 [364], GNLY (granulysin) [365], ZAP70 [366], FOXO3 [367], CD28 [368], GPX1 [369], TLR1 [370], PRDX6 [371], TLR6 [370], C5AR2 [372], LRRK2 [373], CCL5 [374], ETS1 [375], SH2D1A [376], CCR3 [377], GZMA (granzyme A) [378], CD48 [379], LTB (lymphotoxin beta) [380], THEMIS (thymocyte selection associated) [381], PICALM (phosphatidylinositol binding clathrin assembly protein) [382], PTAFR (platelet activating factor receptor) [383], BSG (basigin (Ok blood group)) [384], ALOX5 [385], VIM (vimentin) [386], CD1D [387], TNFSF4 [388], CORO1A [389], TAL1 [390], ICAM2 [391], TLR4 [392], FLNA (filamin A) [393], CCR7 [394], DPP4 [395], NLRC4 [396], TLR2 [397], CSF1R [398], LGALS1 [399], CXCL5 [400], CD27 [401], CD36 [402], LYST (lysosomal trafficking regulator) [403], RNF125 [404], GPR183 [405], FCGR2A [406], CD5 [407], RGS10 [408], VCAN (versican) [409], ANXA6 [410], FKBP8 [411], PPM1A [412], MAF1 [413], GPR183 [414], OGT (O-linked N-acetylglucosamine (GlcNAc) transferase) [415], MKRN1 [416], STAT4 [417], DUSP6 [418], COL5A1 [419], CXCL10 [420], SPRR2A [421], KLF15 [422], DDC (dopa decarboxylase) [423], CRP (C-reactive protein) [424], CD34 [425], BCL3 [426], GAP43 [427], TSLP (thymic stromal lymphopoietin) [428], CCL2 [429], IL17A [430], AHSG (alpha 2-HS glycoprotein) [431], DLL4 [432], HRG (histidine rich glycoprotein) [433], FGF4 [434], CXCL9 [435], ANKS4B [436], C8B [437], SLC39A2 [438] and LAMP3 [439] were shown to participate in facilitating the secondary infections. S1PR1 [440], PPBP (pro-platelet basic protein) [441], CCR2 [442], NLRP12 [443], PECAM1 [444], TRIM10 [445], DEFA3 [446], FCN1 [447], PIM1 [448], KAT2B [449], S1PR4 [447], CCR4 [450], PTPRC (protein tyrosine phosphatase receptor type C) [451], CX3CR1 [452], FOXO3 [453], KLRD1 [454], CD28 [455], GPX1 [456], TLR6 [457], CCL5 [458], ETS1 [459], CCR3 [460], FGL2 [461], ALOX5 [462], SELPLG (selectin P ligand) [463], VIM (vimentin) [464], CD93 [465], CD1D [466], TNFSF4 [467], LEF1 [468], ICAM2 [469], TLR4 [470], FLNA (filamin A) [471], CCR7 [472], DPP4 [473], NLRC4 [474], PLCL2 [475], ITGB3 [476], TLR2 [477], CSF1R [478], CXCL5 [479], CD27 [480], CD36 [481], MAPK1 [482], P2RY12 [483], TGFBR2 [484], FCGR2A [485], CD53 [486], LTA4H [487], LILRB2 [488], ADIPOR1 [489], GPR146 [490], RGS10 [491], VCAN (versican) [492], NINJ2 [493], PDE3B [494], LRRC25 [495], AIDA (axininteractor, dorsalization associated) [496], ABCC4 [497], RPL7 [498], MAF1 [499], HACD1 [500], PINK1 [501], CR1 [502], OGT (O-linked N-acetylglucosamine (GlcNAc) transferase) [503], AKT3 [504], TUBB1 [505], F13A1 [506], DUSP6 [507], MYOCD (myocardin) [508], PEAR1 [509], FSTL3 [510], NTRK3 [511], ECM1 [512], CXCL10 [513], KLF15 [514], RBFOX1 [515], TFAP2B [516], NFKBIL1 [517], ACTN2 [518], APOA4 [519], ACTA2 [520], CRP (C-reactive protein) [521], BMI1 [522], IL17F [523], BCL3 [524], HSF2 [525], TSLP (thymic stromal lymphopoietin) [526], KCNJ11 [527], CCL2 [528], APLNR (apelin receptor) [529], STC1 [530], DAND5 [531], POSTN (periostin) [532], S100A7 [533], SPRR2B [534], SLC6A4 [535], MYOZ1 [536], ABCG8 [537], IL17A [538], PTCH1 [539], NDRG4 [540], FAIM2 [541], TNFRSF11B [542], TDGF1 [543], CXCL9 [544], SLCO1B1 [545], EMP1 [546] and SLC39A2 [547] have been reported to be expressed in cardiovascular diseases, but these genes might be involved in the progression of COVID-19. Hu et al. [548], Zhang et al. [549], Abid et al. [550], Zhang et al. [551], Satoh et al. [552], Ahadzadeh et al. [553], Rudemiller et al. [283], Perros et al. [554], Morris et al. [555], Vinh et al. [556], Ardanaz et al. [557], Nie et al. [558], Song et al. [559], Fan et al. [560], Almodovar et al. [561], Grylls et al. [562], Liu et al. [563], Cai et al. [564], Li et al. [565], Liu et al. [566], Kurahara et al. [567], Holmes et al. [568], Santulli et al. [569], Beitelshees et al. [570], Pravenec et al. [571], Chen et al. [572], Chang et al. [573], Pei et al. [574], Park et al. [575], Claude et al. [576], Saraji et al. [577], Acuña et al. [578], Barnes et al. [579], Sahoo et al. [580], Olivi et al. [581], Antonelli et al. [582], Jiang et al. [583], Kominami et al. [584], Marks et al. [585], Hage, [586], Li et al. [587], Diana et al. [588], Liu et al. [589], Singh et al. [590], Shimizu et al. [591], Huang et al. [525], Zhancheng et al. [592], Alsheikh et al. [593], Zaw et al. [594], Nowzari et al. [529], Zhang et al. [595], Davis et al. [596], Le et al. [597] and Kim et al. [598] found that S1PR1, KRT1, CCR2, NLRP12, PECAM1, PIM1, CX3CR1, CD247, GNLY (granulysin), FOXO3, CD28, GPX1, CCL5, ETS1, FGL2, ALOX5, TLR4, FLNA (filamin A), CCR7, ITGB3, FYN (FYN proto-oncogene, Src family tyrosine kinase), TLR2, CAMK4, CXCL5, CD36, TGFBR2, VCAN (versican), ABCG2, IL10RA, ABCC4, PINK1, SLC12A1, OGT (O-linked N-acetylglucosamine (GlcNAc) transferase), MYOCD (myocardin), PEAR1, CXCL10, KLF15, NFKBIL1, ACTA2, CRP (C-reactive protein), BMI1, GRIA1, WNT7A, IL17F, CD34, HSF2, KCNJ11, CCL2, SCT (secretin), APLNR (apelin receptor), SLC6A4, IL17A, SLC2A2 and SLC6A3 were related to the progression of hypertension, but these genes might be involved in the progression of COVID-19. Mahallawi and Suliman [599], Vanderheiden et al [600], Moustaqil et al [601], Li et al [602], McGee et al [603], Spoerl et al [604], Seale et al [605], Jeong et al [606], Pérez-García et al [607], Murch, [608], Virant-Klun and Strle [609], Bonyek-Silva et al [610], Neri et al [611], Delaveris et al [612], Brandão et al [613], Solerte et al [614], Duncan-Lowey et al [615], Zheng et al [616], Jamaly et al [617], Nassir et al [618], Liang et al [619], Vietzen et al [620], Kisserli et al [621], Ercan et al [622], Oliviero Ercan et al [623], Mpekoulis et al [624], Acar et al [625], Maione et al [626], Ehsani, [627] and Callahan et al [628] showed that TLR8, CCR2, NLRP12, PECAM1, ITK (IL2 inducible T cell kinase), CCR4, GPX1, TLR1, CCL5, CD33, BSG (basigin (Ok blood group)), ALOX5, SELPLG (selectin P ligand), SIGLEC9, TLR4, DPP4, NLRC4, TLR2, CAMK4, FCGR3B, CXCL5, KLRC2, CR1, F13A1, CXCL10, DDC (dopa decarboxylase), CRP (C-reactive protein), IL17A, HAMP (hepcidin antimicrobial peptide) and CXCL9 were associated with progression of COVID 19. Ahmad et al [629], Sireesh et al [630], Di Prospero et al [631], Anjosa et al [632], Aparicio et al [633], Rau et al [634], Kim et al [635], Thude et al [636], Njerve et al. [452], Eldor et al [637], Ferjeni et al [638], Zurawek et al [639], Yang et al. [106], Li et al. [285], Liu et al [640], Pacifici et al [641], Huang et al [642], Zhang et al [643], Pawłowicz et al [644], Ramos-Lopez et al [645], Bonyek-Silva et al [610], Abu El-Ella et al [646], Mi et al [647], Swafford et al [648], Doody et al [649], Barchetta et al [650], Xu et al [651], Li et al [652], Demirci et al [653], Hasani Ranjbar et al [654], Puchałowicz and Rać [655], Fejes et al [656], Kawabata et al [657], Haseda et al [658], Fernández-Real et al [659], Peters et al [660], Sun et al [661], Mao et al [662], Sano et al [663], Szabó et al [664], Shalaby et al [665], Yu et al [666], Xiang et al [667], Abdelmajed et al [668], Antonelli et al [669], Maeda et al [670], Wong et al. [519], Lee et al [671], Kochetova et al [672], Wang et al [673], Tripaldi et al [674], Izumi et al [675], Pehlić et al [676], Bursova et al [677], Haghvirdizadeh et al [678], Igoillo-Esteve et al [679], Xiu et al [680], Lorenzo et al [681], Abed et al [682], Qiu et al [683], Tabassum et al [684], Rees et al [685], Mancuso et al [686], Laukkanen et al [687], Kang et al [688], Wu et al [689], Licito et al [690], Salek Maghsoudi et al [691] and Porta et al [692] were reported that TLR8, NFE2, CCR2, PECAM1, FCN1, CD3D, LST1, CCR4, PTPRC (protein tyrosine phosphatase receptor type C), CX3CR1, CD247, ZAP70, FOXO3, ATM (ATM serine/threonine kinase), CD28, GPX1, PRDX6, LRRK2, CCL5, FCRL3, ETS1, CD48, ALOX5, CD226, CD1D, IKZF1, TLR4, DPP4, NLRC4, RASGRP1, TLR2, CXCL5, CD36, P2RY12, ITGB7, CD300E, TFRC (transferrin receptor), ADIPOR1, ANK1, CD52, PDE3B, ABCG2, KLRC3, SORL1, PINK1, STAT4, CXCL10, TFAP2B, APOA4, CRP (C-reactive protein), GRIA1, WNT7A, CD34, STATH (statherin), IL12RB2, GAP43, KCNJ11, CCL2, SLC6A4, PAX4, ABCG8, IL17A, DOK5, PCK1, AHSG (alpha 2-HS glycoprotein), SLC2A2, FAIM2, HAMP (hepcidin antimicrobial peptide), SLCO1B1, SERPINA12 and SLC19A3 an important target gene of diabetes mellitus, but these genes might be involved in the progression of COVID-19. She et al. [693], Malik et al. [694], Ma et al. [695], Sutti et al. [696], Piccolo et al. [697], Yan et al. [698], Xiao et al. [699], Selzner et al. [700], Ito et al. [701], Zhang and Niu [702], Wang et al. [703], DeSantis et al. [704], Roh et al. [705], Wang et al. [232], Tacke et al. [706], Fernández-Real et al. [707], Fu et al. [708], Dibra et al. [709], Goodwin et al. [710], Malik et al. [711], Bukong et al. [712], Sachar and Ma [713], Shi et al. [714], Zhang et al. [715], Chalin et al. [716], Kawakami et al. [717], Zhang et al. [718], Li et al. [719], Mak and Uetrecht [720], Wu et al. [721], Huang et al. [722], Tan et al. [723], Sun et al. [724], Zhang et al. [725], Bartneck et al. [726], Shams et al. [727], Mustafa and Clarke [728] and Song et al. [729] suggested that activity of CCR2, PECAM1, PIM1, CX3CR1, FOXO3, GPX1, CCL5, SIGLEC9, TLR4, DPP4, NLRC4, TLR2, CD40LG, CXCL5, CD36, TGFBR2, IL27RA, FECH (ferrochelatase), PECAM1, VCAN (versican), ABCG2, PINK1, OGT (O-linked N-acetylglucosamine (GlcNAc) transferase), CXCL10, FABP1, CRP (C-reactive protein), TSLP (thymic stromal lymphopoietin), CCL2, SCT (secretin), POSTN (periostin), IL17A, DLL4, EMCN (endomucin), HRG (histidine rich glycoprotein), FGF4, OTC (ornithine carbamoyltransferase) and CXCL9 were involved in liver injury, but these genes might be involved in the progression of COVID-19.

The PPI network of the DEGs was constructed, and modules were extracted and analyzed. The hub genes were identified by using topological parameters includes node degree, betweenness centrality, stress centrality and closeness centrality. Thus, RPL10, UBA52, TRIM42, THEMIS, TRAT1, LCK, GRIA2, CALB2, CAMKV, DCLK1 and SNAP91 can be a novel molecular markers of COVID-19 and provide new insight to improve therapeutic strategies for COVID-19 complications.

As a prediction tool, the miRNA-hub gene regulatory network (miRNet) and TF-hub gene regulatory network (NetworkAnalyst) databases were utilized to confirm the potential genes, miRNA and TF. hsa-mir-1910-3p [730] and STAT3 [731] are associated with COVID-19. Cao et al. [732], Krause et al. [733], Haghnazari et al. [734], Ao et al. [735] and Zhang et al. [736] showed that the hsa-mir-766-3p, SREBF1, TP53, EGR1 and STAT3 biomarkers might be related to the pathophysiology of diabetes mellitus, but these genes might be involved in the progression of COVID-19. hsa-mir-124-3p [737], SREBF1 [738], ELK1 [739], NFYA (Nuclear transcription factor Y subunit alpha) [740], TP53 [741], EGR1 [742] and STAT3 [743] have been shown as a promising biomarkers in hypertension, but these genes might be involved in the progression of COVID-19. SREBF1 [744], TP53 [745] and EGR1 [746] have been revealed to be highly expressed in CNS disorders, but these genes might be involved in the progression of COVID-19. SREBF1 [747], ELF5 [748], TP53 [749], EGR1 [750], E2F6 [751] and STAT3 [752] have been demonstrated to function in cardiovascular diseases, but these genes might be involved in the progression of COVID-19. ELK1 [753], EGR1 [754] and STAT3 [755] plays a key role in the development of the lung diseases, but these genes might be involved in the progression of COVID-19. Previous studies have reported that EGR1 [756], FOXL1 [757] and STAT3 [758] exhibit significant associations with liver injury, but these genes might be involved in the progression of COVID-19. EGR1 [759] and STAT3 [760] plays important roles in blood clotting processes. Studies have shown that EGR1 [761] and STAT3 [762] altered expressed in secondary infections. Altered expression of EGR1 [763] and STAT3 [764] were observed to be associated with the progression of renal diseases, but these genes might be involved in the progression of COVID-19. Thus, GDI1, DCX, KRT13, hsa-mir-320a, hsa-mir-4736, hsa-mir-559, hsa-mir-186-3p, hsa-mir-4495, hsa-mir-3689a-5p and hsa-mir-6842-5p can be a novel molecular markers of COVID-19 and provide new insight to improve therapeutic strategies for COVID-19 complications.

In conclusion, a comprehensive bioinformatics analysis of DEGs and pathways involved in the occurrence and development of COVID-19 and its secondary complications was performed, and we explored and obtained key regulatory genes and pathways contributing to the progression of COVID-19 and its secondary complications to improve prognosis. Moreover, these results may promote the understanding of molecular mechanisms and clinically related molecular targets for prognosis in COVID-19 and its secondary complications, and provide new insight into the occurrence and development of COVID-19 and its secondary complications.

## Acknowledgement

I thank Charles Chiu, University of California, Laboratory Medicine,San Francisco, USA, very much, the author who deposited their NGS dataset GSE163151, into the public GEO database.

## Ethical approval

This article does not contain any studies with human participants or animals performed by any of the authors.

## Informed consent

No informed consent because this study does not contain human or animals participants.

## Availability of data and materials

The datasets supporting the conclusions of this article are available in the GEO (Gene Expression Omnibus) (https://www.ncbi.nlm.nih.gov/geo/) repository. [(GSE163151) https://www.ncbi.nlm.nih.gov/geo/query/acc.cgi?acc=GSE163151)]

## Consent for publication

Not applicable.

## Author Contributions

Muttanagouda Giriyappagoudar **-** Acquisition of resources and investigation

Basavaraj Vastrad - Writing original draft, and review and editing

Rajeshwari Horakeri - Formal analysis and validation

Chanabasayya Vastrad - Software and investigation

## References

1. Alsharif W, Qurashi A. Effectiveness of COVID-19 diagnosis and management tools: A review. Radiography (Lond). 2021;27(2):682–687. doi:10.1016/j.radi.2020.09.010

2. Sanders JM, Monogue ML, Jodlowski TZ, Cutrell JB. Pharmacologic Treatments for Coronavirus Disease 2019 (COVID-19): A Review. JAMA. 2020;323(18):1824–1836. doi:10.1001/jama.2020.6019

3. Sadegh Beigee F, Pourabdollah Toutkaboni M, Khalili N, Nadji SA, Dorudinia A, Rezaei M, Askari E, Farzanegan B, Marjani M, Rafiezadeh A. Diffuse alveolar damage and thrombotic microangiopathy are the main histopathological findings in lung tissue biopsy samples of COVID-19 patients. Pathol Res Pract. 2020;216(10):153228. doi:10.1016/j.prp.2020.153228

4. Babadaei MMN, Hasan A, Bloukh SH, Edis Z, Sharifi M, Kachooei E, Falahati M. The expression level of angiotensin-converting enzyme 2 determines the severity of COVID-19: lung and heart tissue as targets. J Biomol Struct Dyn. 2021;39(10):3780–3786. doi:10.1080/07391102.2020.1767211

5. Baig AM, Khaleeq A, Ali U, Syeda H. Evidence of the COVID-19 Virus Targeting the CNS: Tissue Distribution, Host-Virus Interaction, and Proposed Neurotropic Mechanisms. ACS Chem Neurosci. 2020;11(7):995–998. doi:10.1021/acschemneuro.0c00122

6. Zhao B, Ni C, Gao R, Wang Y, Yang L, Wei J, Lv T, Liang J, Zhang Q, Xu W, et al. Recapitulation of SARS-CoV-2 infection and cholangiocyte damage with human liver ductal organoids. Protein Cell. 2020;11(10):771–775. doi:10.1007/s13238-020-00718-6

7. Ahmadian E, Hosseiniyan Khatibi SM, Razi Soofiyani S, Abediazar S, Shoja MM, Ardalan M, Zununi Vahed S. Covid-19 and kidney injury: Pathophysiology and molecular mechanisms. Rev Med Virol. 2021;31(3):e2176. doi:10.1002/rmv.2176

8. Carpagnano GE, Di Lecce V, Quaranta VN, Zito A, Buonamico E, Capozza E, Palumbo A, Di Gioia G, Valerio VN, Resta O. et al. Vitamin D deficiency as a predictor of poor prognosis in patients with acute respiratory failure due to COVID-19. J Endocrinol Invest. 2021;44(4):765–771. doi:10.1007/s40618-020-01370-x

9. Gattinoni L, Chiumello D, Caironi P, Busana M, Romitti F, Brazzi L, Camporota L. COVID-19 pneumonia: different respiratory treatments for different phenotypes?. Intensive Care Med. 2020;46(6):1099–1102. doi:10.1007/s00134-020-06033-2

10. Fan E, Beitler JR, Brochard L, Calfee CS, Ferguson ND, Slutsky AS, Brodie D. COVID-19-associated acute respiratory distress syndrome: is a different approach to management warranted?. Lancet Respir Med. 2020;8(8):816–821. doi:10.1016/S2213-2600(20)30304-0

11. Effenberger M, Grander C, Grabherr F, Griesmacher A, Ploner T, Hartig F, Bellmann-Weiler R, Joannidis M, Zoller H, Weiss G, et al. Systemic inflammation as fuel for acute liver injury in COVID-19. Dig Liver Dis. 2021;53(2):158–165. doi:10.1016/j.dld.2020.08.004

12. Vakhshoori M, Heidarpour M, Shafie D, Taheri M, Rezaei N, Sarrafzadegan N. Acute Cardiac Injury in COVID-19: A Systematic Review and Meta-analysis. Arch Iran Med. 2020;23(11):801–812. doi:10.34172/aim.2020.107

13. Ripa M, Galli L, Poli A, Oltolini C, Spagnuolo V, Mastrangelo A, Muccini C, Monti G, De Luca G, Landoni G, et al. Secondary infections in patients hospitalized with COVID-19: incidence and predictive factors. Clin Microbiol Infect. 2021;27(3):451–457. doi:10.1016/j.cmi.2020.10.021

14. Virhammar J, Nääs A, Fällmar D, Cunningham JL, Klang A, Ashton NJ, Jackmann S, Westman G, Frithiof R, Blennow K, et al. Biomarkers for central nervous system injury in cerebrospinal fluid are elevated in COVID-19 and associated with neurological symptoms and disease severity. Eur J Neurol. 2021;28(10):3324–3331. doi:10.1111/ene.14703

15. Legrand M, Bell S, Forni L, Joannidis M, Koyner JL, Liu K, Cantaluppi V. Pathophysiology of COVID-19-associated acute kidney injury. Nat Rev Nephrol. 2021;17(11):751–764. doi:10.1038/s41581-021-00452-0

16. Hantoushzadeh S, Norooznezhad AH. Possible Cause of Inflammatory Storm and Septic Shock in Patients Diagnosed with (COVID-19). Arch Med Res. 2020;51(4):347–348. doi:10.1016/j.arcmed.2020.03.015

17. Sardu C, Gargiulo G, Esposito G, Paolisso G, Marfella R. Impact of diabetes mellitus on clinical outcomes in patients affected by Covid-19. Cardiovasc Diabetol. 2020;19(1):76. doi:10.1186/s12933-020-01047-y

18. Zuin M, Rigatelli G, Zuliani G, Rigatelli A, Mazza A, Roncon L. Arterial hypertension and risk of death in patients with COVID-19 infection: Systematic review and meta-analysis. J Infect. 2020;81(1):e84–e86. doi:10.1016/j.jinf.2020.03.059

19. Zhou X, Cheng Z, Luo L, Zhu Y, Lin W, Ming Z, Chen W, Hu Y. Incidence and impact of disseminated intravascular coagulation in COVID-19 a systematic review and meta-analysis. Thromb Res. 2021;201:23–29. doi:10.1016/j.thromres.2021.02.010

20. Biswas S, Thakur V, Kaur P, Khan A, Kulshrestha S, Kumar P. Blood clots in COVID-19 patients: Simplifying the curious mystery. Med Hypotheses. 2021;146:110371. doi:10.1016/j.mehy.2020.110371

21. Riollano-Cruz M, Akkoyun E, Briceno-Brito E, Kowalsky S, Reed J, Posada R, Sordillo EM, Tosi M, Trachtman R, Paniz-Mondolfi A. Multisystem inflammatory syndrome in children related to COVID-19: A New York City experience. J Med Virol. 2021;93(1):424–433. doi:10.1002/jmv.26224

22. Wostyn P. COVID-19 and chronic fatigue syndrome: Is the worst yet to come?. Med Hypotheses. 2021;146:110469. doi:10.1016/j.mehy.2020.110469

23. Geng Y, Ma Q, Du YS, Peng N, Yang T, Zhang SY, Wu FF, Lin HL, Su L. Rhabdomyolysis is Associated with In-Hospital Mortality in Patients with COVID-19. Shock. 2021;56(3):360–367. doi:10.1097/SHK.0000000000001725

24. Keni R, Alexander A, Nayak PG, Mudgal J, Nandakumar K. COVID-19: Emergence, Spread, Possible Treatments, and Global Burden. Front Public Health. 2020;8:216. doi:10.3389/fpubh.2020.00216

25. Jamwal S, Gautam A, Elsworth J, Kumar M, Chawla R, Kumar P. An updated insight into the molecular pathogenesis, secondary complications and potential therapeutics of COVID-19 pandemic. Life Sci. 2020;257:118105. doi:10.1016/j.lfs.2020.118105

26. Wu H, Larsen CP, Hernandez-Arroyo CF, Mohamed MMB, Caza T, Sharshir M, Chughtai A, Xie L, Gimenez JM, Sandow TA, et al. AKI and Collapsing Glomerulopathy Associated with COVID-19 and APOL1 High-Risk Genotype. J Am Soc Nephrol. 2020;31(8):1688–1695. doi:10.1681/ASN.2020050558

27. Zhang H, Rostami MR, Leopold PL, Mezey JG, O’Beirne SL, Strulovici-Barel Y, Crystal RG. Expression of the SARS-CoV-2 ACE2 Receptor in the Human Airway Epithelium. Am J Respir Crit Care Med. 2020;202(2):219–229. doi:10.1164/rccm.202003-0541OC

28. Zhang N, Zhao YD, Wang XM. CXCL10 an important chemokine associated with cytokine storm in COVID-19 infected patients. Eur Rev Med Pharmacol Sci. 2020;24(13):7497–7505. doi:10.26355/eurrev_202007_21922

29. Smieszek SP, Polymeropoulos VM, Xiao C, Polymeropoulos CM, Polymeropoulos MH. Loss-of-function mutations in IFNAR2 in COVID-19 severe infection susceptibility. J Glob Antimicrob Resist. 2021;26:239–240. doi:10.1016/j.jgar.2021.06.005

30. Medetalibeyoglu A, Bahat G, Senkal N, Kose M, Avci K, Sayin GY, Isoglu-Alkac U, Tukek T, Pehlivan S. Mannose binding lectin gene 2 (rs1800450) missense variant may contribute to development and severity of COVID-19 infection. Infect Genet Evol. 2021;89:104717. doi:10.1016/j.meegid.2021.104717

31. Luo J, Lu S, Yu M, Zhu L, Zhu C, Li C, Fang J, Zhu X, Wang X. The potential involvement of JAK-STAT signaling pathway in the COVID-19 infection assisted by ACE2. Gene. 2021;768:145325. doi:10.1016/j.gene.2020.145325

32. Motaghinejad M, Gholami M. Possible Neurological and Mental Outcomes of COVID-19 Infection: A Hypothetical Role of ACE-2\Mas\BDNF Signaling Pathway. Int J Prev Med. 2020;11:84. doi:10.4103/ijpvm.IJPVM_114_20

33. Liu Y, Lv J, Liu J, Li M, Xie J, Lv Q, Deng W, Zhou N, Zhou Y, Song J, et al. Mucus production stimulated by IFN-AhR signaling triggers hypoxia of COVID-19. Cell Res. 2020;30(12):1078–1087. doi:10.1038/s41422-020-00435-z

34. Sohn KM, Lee SG, Kim HJ, Cheon S, Jeong H, Lee J, Kim IS, Silwal P, Kim YJ, Paik S, et al. COVID-19 Patients Upregulate Toll-like Receptor 4-mediated Inflammatory Signaling That Mimics Bacterial Sepsis. J Korean Med Sci. 2020;35(38):e343. doi:10.3346/jkms.2020.35.e343

35. Chen X, Kang Y, Luo J, Pang K, Xu X, Wu J, Li X, Jin S. Next-Generation Sequencing Reveals the Progression of COVID-19. Front Cell Infect Microbiol. 2021;11:632490. doi:10.3389/fcimb.2021.632490

36. Prashanth G, Vastrad B, Tengli A, Vastrad C, Kotturshetti I. Identification of hub genes related to the progression of type 1 diabetes by computational analysis. BMC Endocr Disord. 2021;21(1):61. doi:10.1186/s12902-021-00709-6

37. Vastrad B, Vastrad C, Tengli A. Bioinformatics analyses of significant genes, related pathways, and candidate diagnostic biomarkers and molecular targets in SARS-CoV-2/COVID-19. Gene Rep. 2020;21:100956. doi:10.1016/j.genrep.2020.100956

38. Ng DL, Granados AC, Santos YA, Servellita V, Goldgof GM, Meydan C, Sotomayor-Gonzalez A, Levine AG, Balcerek J, Han LM, et al. A diagnostic host response biosignature for COVID-19 from RNA profiling of nasal swabs and blood. Sci Adv. 2021;7(6):eabe5984. doi:10.1126/sciadv.abe5984

39. Clough E, Barrett T. The Gene Expression Omnibus Database. Methods Mol Biol. 2016;1418:93–110. doi:10.1007/978-1-4939-3578-9_5

40. Love MI, Huber W, Anders S. Moderated estimation of fold change and dispersion for RNA-seq data with DESeq2. Genome Biol. 2014;15(12):550. doi:10.1186/s13059-014-0550-8

41. Gauthier M, Agniel D, Thiébaut R, Hejblum BP. dearseq: a variance component score test for RNA-seq differential analysis that effectively controls the false discovery rate. NAR Genom Bioinform. 2020;2(4):lqaa093. doi:10.1093/nargab/lqaa093

42. Reimand J, Kull M, Peterson H, Hansen J, Vilo J. g:Profiler--a web-based toolset for functional profiling of gene lists from large-scale experiments. Nucleic Acids Res. 2007;35(Web Server issue):W193–W200. doi:10.1093/nar/gkm226

43. Thomas PD. The Gene Ontology and the Meaning of Biological Function. Methods Mol Biol. 2017;1446:15L24. doi:10.1007/978-1-4939-3743-1_2

44. Fabregat A, Jupe S, Matthews L, Sidiropoulos K, Gillespie M, Garapati P, Haw R, Jassal B, Korninger F, May B et al The Reactome Pathway Knowledgebase. Nucleic Acids Res. 2018;46(D1):D649–D655. doi:10.1093/nar/gkx1132

45. Pastrello C, Kotlyar M, Jurisica I. Informed Use of Protein-Protein Interaction Data: A Focus on the Integrated Interactions Database (IID). Methods Mol Biol. 2020;2074:125–134. doi:10.1007/978-1-4939-9873-9_10

46. Shannon P, Markiel A, Ozier O, Baliga NS, Wang JT, Ramage D, Amin N, Schwikowski B, Ideker T Cytoscape: a software environment for integrated models of biomolecular interaction networks. Genome Res 2003;13(11):2498–2504. doi:10.1101/gr.1239303

47. Przulj N, Wigle DA, Jurisica I. Functional topology in a network of protein interactions. Bioinformatics. 2004;20(3):340–348. doi:10.1093/bioinformatics/btg415

48. Nguyen TP, Liu WC, Jordán F. Inferring pleiotropy by network analysis: linked diseases in the human PPI network. BMC Syst Biol. 2011;5:179. doi:10.1186/1752-0509-5-179

49. Shi Z, Zhang B. Fast network centrality analysis using GPUs. BMC Bioinformatics. 2011;12:149. doi:10.1186/1471-2105-12-149

50. Fadhal E, Gamieldien J, Mwambene EC. Protein interaction networks as metric spaces: a novel perspective on distribution of hubs. BMC Syst Biol. 2014;8:6. doi:10.1186/1752-0509-8-6

51. Zaki N, Efimov D, Berengueres J. Protein complex detection using interaction reliability assessment and weighted clustering coefficient. BMC Bioinformatics. 2013;14:163. doi:10.1186/1471-2105-14

52. Fan Y, Xia J (2018) miRNet-Functional Analysis and Visual Exploration of miRNA-Target Interactions in a Network Context. Methods Mol Biol 1819:215–233. doi:10.1007/978-1-4939-8618-7_10

53. Zhou G, Soufan O, Ewald J, Hancock REW, Basu N, Xia J (2019) NetworkAnalyst 3.0: a visual analytics platform for comprehensive gene expression profiling and meta-analysis. Nucleic Acids Res 47:W234–W241. doi:10.1093/nar/gkz240

54. Robin X, Turck N, Hainard A, Tiberti N, Lisacek F, Sanchez JC, Müller M. pROC: an open-source package for R and S+ to analyze and compare ROC curves. BMC Bioinformatics 2011;12:77. doi:10.1186/1471-2105-12-77

55. Sahraian MA, Azimi A, Navardi S, Ala S, Naser Moghadasi A. Evaluation of the rate of COVID-19 infection, hospitalization and death among Iranian patients with multiple sclerosis. Mult Scler Relat Disord. 2020;46:102472. doi:10.1016/j.msard.2020.102472

56. Zhang M, Shi R, Zhang Y, et al. Nix/BNIP3L-dependent mitophagy accounts for airway epithelial cell injury induced by cigarette smoke. J Cell Physiol. 2019;234(8):14210–14220. doi:10.1002/jcp.28117

57. Jiang J, Huang K, Xu S, Garcia JGN, Wang C, Cai H. Targeting NOX4 alleviates sepsis-induced acute lung injury via attenuation of redox-sensitive activation of CaMKII/ERK1/2/MLCK and endothelial cell barrier dysfunction. Redox Biol. 2020;36:101638. doi:10.1016/j.redox.2020.101638

58. Karki P, Cha B, Zhang CO, Li Y, Ke Y, Promnares K, Kaibuchi K, Yoshimura A, Birukov KG, Birukova AA. Microtubule-dependent mechanism of anti-inflammatory effect of SOCS1 in endothelial dysfunction and lung injury. FASEB J. 2021;35(4):e21388. doi:10.1096/fj.202001477RR

59. Qin LY, Wang MX, Zhang H. MiR-133a alleviates renal injury caused by sepsis by targeting BNIP3L. Eur Rev Med Pharmacol Sci. 2020;24(5):2632–2639. doi:10.26355/eurrev_202003_20532

60. Rajaram RD, Dissard R, Faivre A, Ino F, Delitsikou V, Jaquet V, Cagarelli T, Lindenmeyer M, Jansen-Duerr P, Cohen C, et al. Tubular NOX4 expression decreases in chronic kidney disease but does not modify fibrosis evolution. Redox Biol. 2019;26:101234. doi:10.1016/j.redox.2019.101234

61. Luan J, Fu J, Wang D, Jiao C, Cui X, Chen C, Liu D, Zhang Y, Wang Y, Yuen PST, et al. miR-150-Based RNA Interference Attenuates Tubulointerstitial Fibrosis through the SOCS1/JAK/STAT Pathway In Vivo and In Vitro. Mol Ther Nucleic Acids. 2020;22:871–884. doi:10.1016/j.omtn.2020.10.008

62. Yuan Y, Zheng Y, Zhang X, Chen Y, Wu X, Wu J, Shen Z, Jiang L, Wang L, et al. BNIP3L/NIX-mediated mitophagy protects against ischemic brain injury independent of PARK2. Autophagy. 2017;13(10):1754–1766. doi:10.1080/15548627.2017.1357792

63. İnce B, Guloksuz S, Altınbaş K, Oral ET, Alpkan LR, Altinoz MA. Minor hemoglobins HbA2 and HbF associate with disease severity in bipolar disorder with a likely protective role of HbA2 against post-partum episodes. J Affect Disord. 2013;151(1):405–408. doi:10.1016/j.jad.2013.06.042

64. Chen V, Moncalvo M, Tringali D, Tagliafierro L, Shriskanda A, Ilich E, Dong W, Kantor B, Chiba-Falek O. The mechanistic role of alpha-synuclein in the nucleus: impaired nuclear function caused by familial Parkinson’s disease SNCA mutations. Hum Mol Genet. 2020;29(18):3107–3121. doi:10.1093/hmg/ddaa183

65. Gołab-Janowska M, Honczarenko K, Gawrońska-Szklarz B, Potemkowski A. The role of NAT2 gene polymorphism in aetiology of the most frequent neurodegenerative diseases with dementia. Neurol Neurochir Pol. 2007;41(5):388–394.

66. Lin H, Hu H, Duan W, Liu Y, Tan G, Li Z, Liu Y, Deng B, Song X, Wang W, et al. Intramuscular Delivery of scAAV9-hIGF1 Prolongs Survival in the hSOD1G93A ALS Mouse Model via Upregulation of D-Amino Acid Oxidase. Mol Neurobiol. 2018;55(1):682–695. doi:10.1007/s12035-016-0335-z

67. Ouyang S, Chen W, Zeng G, Lei C, Tian G, Zhu M, Liu Y, Yang M.MicroRNA-183-3p up-regulated by vagus nerve stimulation mitigates chronic systolic heart failure via the reduction of BNIP3L-mediated autophagy. Gene. 2020;726:144136. doi:10.1016/j.gene.2019.144136

68. Vendrov AE, Vendrov KC, Smith A, Yuan J, Sumida A, Robidoux J, Runge MS, Madamanchi NR. NOX4 NADPH Oxidase-Dependent Mitochondrial Oxidative Stress in Aging-Associated Cardiovascular Disease. Antioxid Redox Signal. 2015;23(18):1389–1409. doi:10.1089/ars.2014.6221

69. Alikhah A, Pahlevan Kakhki M, Ahmadi A, Dehghanzad R, Boroumand MA, Behmanesh M. The role of lnc-DC long non-coding RNA and SOCS1 in the regulation of STAT3 in coronary artery disease and type 2 diabetes mellitus. J Diabetes Complications. 2018;32(3):258–265. doi:10.1016/j.jdiacomp.2017.12.001

70. Khelil M, Zenati A, Makrelouf M, Otmane A, Tayebi B. Polymorphisms in NAT2 gene and atherosclerosis in an Algerian population. Arch Med Res. 2010;41(3):215–220. doi:10.1016/j.arcmed.2010.03.008

71. Xia XD, Yu XH, Chen LY, Xie SL, Feng YG, Yang RZ, Zhao ZW, Li H, Wang G, Tang CK. Myocardin suppression increases lipid retention and atherosclerosis via downregulation of ABCA1 in vascular smooth muscle cells. Biochim Biophys Acta Mol Cell Biol Lipids. 2021;1866(4):158824. doi:10.1016/j.bbalip.2020.158824

72. Hill QA, Farrar L, Lordan J, Gallienne A, Henderson S. A combination of two novel alpha globin variants Hb Bridlington (HBA1) and Hb Taybe (HBA2) resulting in severe hemolysis, pulmonary hypertension, and death. Hematology. 2015;20(1):50–52. doi:10.1179/1607845414Y.0000000164

73. Barman SA, Fulton D. Adventitial Fibroblast Nox4 Expression and ROS Signaling in Pulmonary Arterial Hypertension. Adv Exp Med Biol. 2017;967:1–11. doi:10.1007/978-3-319-63245-2_1

74. Li XP, Liu Y, Zhang CQ. Correlation between NAT2 gene polymorphism and cirrhotic portal hypertension in the Chinese population. Genet Test Mol Biomarkers. 2015;19(3):138–143. doi:10.1089/gtmb.2014.0283

75. Sahoo S, Meijles DN, Al Ghouleh I, Tandon M, Cifuentes-Pagano E, Sembrat J, Rojas M, Goncharova E, Pagano PJ. MEF2C-MYOCD and Leiomodin1 Suppression by miRNA-214 Promotes Smooth Muscle Cell Phenotype Switching in Pulmonary Arterial Hypertension. PLoS One. 2016;11(5):e0153780. doi:10.1371/journal.pone.0153780

76. Papadimitriou A, Peixoto EB, Silva KC, Lopes de Faria JM, Lopes de Faria JB. Increase in AMPK brought about by cocoa is renoprotective in experimental diabetes mellitus by reducing NOX4/TGFβ-1 signaling. J Nutr Biochem. 2014;25(7):773–784. doi:10.1016/j.jnutbio.2014.03.010

77. Irshaid YM, Abujbara MA, Ajlouni KM, El-Khateeb M, Jarrar YB. N-acetyltransferase-2 genotypes among Jordanian patients with diabetes mellitus. Int J Clin Pharmacol Ther. 2013;51(7):593–599. doi:10.5414/CP201883

78. Cheng Q, Li C, Yang CF, Zhong YJ, Wu D, Shi L, Chen L, Li YW, Li L. Methyl ferulic acid attenuates liver fibrosis and hepatic stellate cell activation through the TGF-β1/Smad and NOX4/ROS pathways. Chem Biol Interact. 2019;299:131–139. doi:10.1016/j.cbi.2018.12.006

79. Zhu H, Zhao H, Xu S, Zhang Y, Ding Y, Li J, Huang C, Ma T.Sennoside A alleviates inflammatory responses by inhibiting the hypermethylation of SOCS1 in CCl4-induced liver fibrosis. Pharmacol Res. 2021;174:105926. doi:10.1016/j.phrs.2021.105926

80. Yu WY, Li L, Wu F, Zhang HH, Fang J, Zhong YS, Yu CH. Moslea Herba flavonoids alleviated influenza A virus-induced pulmonary endothelial barrier disruption via suppressing NOX4/NF-κB/MLCK pathway. J Ethnopharmacol. 2020;253:112641. doi:10.1016/j.jep.2020.112641

81. Mishra R, Krishnamoorthy P, Kumar H. MicroRNA-30e-5p Regulates SOCS1 and SOCS3 During Bacterial Infection. Front Cell Infect Microbiol. 2021;10:604016. doi:10.3389/fcimb.2020.604016

82. Girona-Alarcon M, Argüello G, Esteve-Sole A, Bobillo-Perez S, Paolo-Burgos X, Bonet-Carne E, Mensa-Vilaró A, Codina A, Hernández-Garcia M, Jou C, et al. Low levels of CIITA and high levels of SOCS1 predict COVID-19 disease severity in children and adults. iScience. 2021;103595. doi:10.1016/j.isci.2021.103595

83. Gorji A, Khaleghi Ghadiri M. Potential roles of micronutrient deficiency and immune system dysfunction in the coronavirus disease 2019 (COVID-19) pandemic. Nutrition. 2021;82:111047. doi:10.1016/j.nut.2020.111047

84. Mudd PA, Remy KE. Prolonged adaptive immune activation in COVID-19: implications for maintenance of long-term immunity?. J Clin Invest. 2021;131(1):e143928. doi:10.1172/JCI143928

85. Bankar R, Suvarna K, Ghantasala S, Banerjee A, Biswas D, Choudhury M, Palanivel V, Salkar A, Verma A, Singh A, et al. Proteomic investigation reveals dominant alterations of neutrophil degranulation and mRNA translation pathways in patients with COVID-19. iScience. 2021;24(3):102135. doi:10.1016/j.isci.2021.102135

86. Nidadavolu LS, Walston JD. Underlying Vulnerabilities to the Cytokine Storm and Adverse COVID-19 Outcomes in the Aging Immune System. J Gerontol A Biol Sci Med Sci. 2021;76(3):e13–e18. doi:10.1093/gerona/glaa209

87. Zhang Q, Friedman PA. Receptor-Loaded Virion Endangers GPCR Signaling: Mechanistic Exploration of SARS-CoV-2 Infections and Pharmacological Implications. Int J Mol Sci. 2021;22(20):10963. doi:10.3390/ijms222010963

88. Appleford NE, Wilson K, Houston F, Bruce LJ, Morrison A, Bishop M, Chalmers K, Miele G, Massey E, Prowse C, et al. alpha-Hemoglobin stabilizing protein is not a suitable marker for a screening test for variant Creutzfeldt-Jakob disease. Transfusion. 2008;48(8):1616–1626. doi:10.1111/j.1537-2995.2008.01759.x

89. Imani SZH, Hojati Z, Khalilian S, Dehghanian F, Kheirollahi M, Khorrami M, Shaygannejad V, Mirmosayyeb O. Expression and clinical significance of IL7R, NFATc2, and RNF213 in familial and sporadic multiple sclerosis. Sci Rep. 2021;11(1):19260. doi:10.1038/s41598-021-98691-5

90. Lorenzo-Betancor O, Lin YH, Samii A, Jayadev S, Kim HM, Longfellow K, Distad BJ, Yearout D, Mata IF, Zabetian CP. Novel compound heterozygous FBXO7 mutations in a family with early onset Parkinson’s disease. Parkinsonism Relat Disord. 2020;80:142–147. doi:10.1016/j.parkreldis.2020.09.035

91. Søndergaard HB, Sellebjerg F, Hillert J, Olsson T, Kockum I, Lindén M, Mero IL, Myhr KM, Celius EG, Harbo HF, et al. Alterations in KLRB1 gene expression and a Scandinavian multiple sclerosis association study of the KLRB1 SNP rs4763655. Eur J Hum Genet. 2011;19(10):1100–1103. doi:10.1038/ejhg.2011.88

92. Levchenko A, Vyalova NM, Nurgaliev T, Pozhidaev IV, Simutkin GG, Bokhan NA, Ivanova SA. NRG1, PIP4K2A, and HTR2C as Potential Candidate Biomarker Genes for Several Clinical Subphenotypes of Depression and Bipolar Disorder. Front Genet. 2020;11:936. doi:10.3389/fgene.2020.00936

93. Oksanen M, Hyötyläinen I, Trontti K, Rolova T, Wojciechowski S, Koskuvi M, Viitanen M, Levonen AL, Hovatta I, Roybon L, et al. NF-E2-related factor 2 activation boosts antioxidant defenses and ameliorates inflammatory and amyloid properties in human Presenilin-1 mutated Alzheimer’s disease astrocytes. Glia. 2020;68(3):589–599. doi:10.1002/glia.23741

94. Bose S, Cho J. Role of chemokine CCL2 and its receptor CCR2 in neurodegenerative diseases. Arch Pharm Res. 2013;36(9):1039–1050. doi:10.1007/s12272-013-0161-z

95. Sagar D, Singh NP, Ginwala R, Huang X, Philip R, Nagarkatti M, Nagarkatti P, Neumann K, Ruland J, Andrews AM, et al. Antibody blockade of CLEC12A delays EAE onset and attenuates disease severity by impairing myeloid cell CNS infiltration and restoring positive immunity. Sci Rep. 2017;7(1):2707. doi:10.1038/s41598-017-03027-x

96. Gharagozloo M, Mahvelati TM, Imbeault E, Gris P, Zerif E, Bobbala D, Ilangumaran S, Amrani A, Gris D. The nod-like receptor, Nlrp12, plays an anti-inflammatory role in experimental autoimmune encephalomyelitis. J Neuroinflammation. 2015;12:198. doi:10.1186/s12974-015-0414-5

97. Williams KC, Zhao RW, Ueno K, Hickey WF. PECAM-1 (CD31) expression in the central nervous system and its role in experimental allergic encephalomyelitis in the rat. J Neurosci Res. 1996;45(6):747–757. doi:10.1002/(SICI)1097-4547(19960915)45:6<747::AID-JNR11>3.0.CO;2-T

98. Huang Q, Zhu X, Xu M. Silencing of TRIM10 alleviates apoptosis in cellular model of Parkinson’s disease. Biochem Biophys Res Commun. 2019;518(3):451–458. doi:10.1016/j.bbrc.2019.08.041

99. Chatzimanolis N, Kraus J, Bauer R, Engelhardt B, Bregenzer T, Kuehne BS, Tofighi J, Laske C, Stolz E, Blaes F, et al. CD45RA+ ICAM-3+ lymphocytes in interferon-beta1b-treated and -untreated patients with relapsing-remitting multiple sclerosis. Acta Neurol Scand. 2004;110(6):377–385. doi:10.1111/j.1600-0404.2004.00346.x

100. Zhang Y, Chen J, Li F, Li D, Xiong Q, Lin Y, Zhang D, Wang XF, Yang P, Rui YC. A pentapeptide monocyte locomotion inhibitory factor protects brain ischemia injury by targeting the eEF1A1/endothelial nitric oxide synthase pathway. Stroke. 2012;43(10):2764–2773. doi:10.1161/STROKEAHA.112.657908

101. Moriguchi K, Miyamoto K, Tanaka N, Yoshie O, Kusunoki S. The importance of CCR4 and CCR6 in experimental autoimmune encephalomyelitis. J Neuroimmunol. 2013;257(1-2):53–58. doi:10.1016/j.jneuroim.2013.02.002

102. Szvetko AL, Jones A, Mackenzie J, Tajouri L, Csurhes PA, Greer JM, Pender MP, Griffiths LR. An investigation of the C77G and C772T variations within the human protein tyrosine phosphatase receptor type C gene for association with multiple sclerosis in an Australian population. Brain Res. 2009;1255:148–152. doi:10.1016/j.brainres.2008.12.017

103. Subbarayan MS, Joly-Amado A, Bickford PC, Nash KR. CX3CL1/CX3CR1 signaling targets for the treatment of neurodegenerative diseases. Pharmacol Ther. 2021;107989. doi:10.1016/j.pharmthera.2021.107989

104. Basile MS, Mazzon E, Mangano K, Pennisi M, Petralia MC, Lombardo SD, Nicoletti F, Fagone P, Cavalli E. Impaired Expression of Tetraspanin 32 (TSPAN32) in Memory T Cells of Patients with Multiple Sclerosis. Brain Sci. 2020;10(1):52. doi:10.3390/brainsci10010052

105. Chen S, Zhang J, Yu WB, Zhuang JC, Xiao W, Wu ZY, Xiao BG. Eomesodermin in CD4+T cells is essential for Ginkgolide K ameliorating disease progression in experimental autoimmune encephalomyelitis. Int J Biol Sci. 2021;17(1):50–61. doi:10.7150/ijbs.50041

106. Yang DQ, Halaby MJ, Li Y, Hibma JC, Burn P. Cytoplasmic ATM protein kinase: an emerging therapeutic target for diabetes, cancer and neuronal degeneration. Drug Discov Today. 2011;16(7-8):332–338. doi:10.1016/j.drudis.2011.02.001

107. Markovic-Plese S, Cortese I, Wandinger KP, McFarland HF, Martin R. CD4+CD28-costimulation-independent T cells in multiple sclerosis. J Clin Invest. 2001;108(8):1185–1194. doi:10.1172/JCI12516

108. Wallings RL, Herrick MK, Tansey MG. LRRK2 at the Interface Between Peripheral and Central Immune Function in Parkinson’s. Front Neurosci. 2020;14:443. doi:10.3389/fnins.2020.00443

109. Pittaluga A. CCL5-Glutamate Cross-Talk in Astrocyte-Neuron Communication in Multiple Sclerosis. Front Immunol. 2017;8:1079. doi:10.3389/fimmu.2017.01079

110. Heidari F, Ansstas G, Ajamian F. CD33 mRNA Has Elevated Expression Levels in the Leukocytes of Peripheral Blood in Patients with Late-Onset Alzheimer’s Disease. Gerontology. 2021;1–10. doi:10.1159/000518820

111. Yuan M, Wei L, Zhou R, Bai Q, Wei Y, Zhang W, Huang Y. Four FCRL3 Gene Polymorphisms (FCRL3_3, _5, _6, _8) Confer Susceptibility to Multiple Sclerosis: Results from a Case-Control Study. Mol Neurobiol. 2016;53(3):2029–2035. doi:10.1007/s12035-015-9149-7

112. Simpson J, Rezaie P, Newcombe J, Cuzner ML, Male D, Woodroofe MN. Expression of the beta-chemokine receptors CCR2, CCR3 and CCR5 in multiple sclerosis central nervous system tissue. J Neuroimmunol. 2000;108(1-2):192–200. doi:10.1016/s0165-5728(00)00274-5

113. Yuan B, Fu F, Huang S, Lin C, Yang G, Ma K, Shi H, Yang Z. C5a/C5aR Pathway Plays a Vital Role in Brain Inflammatory Injury via Initiating Fgl-2 in Intracerebral Hemorrhage. Mol Neurobiol. 2017;54(8):6187–6197. doi:10.1007/s12035-016-0141-7

114. Iłżecka J. Granzymes A and B levels in serum of patients with amyotrophic lateral sclerosis. Clin Biochem. 2011;44(8-9):650–653. doi:10.1016/j.clinbiochem.2011.02.006

115. Seripa D, Panza F, Paroni G, D’Onofrio G, Bisceglia P, Gravina C, Urbano M, Lozupone M, Solfrizzi V, Bizzarro A, et al. Role of CLU, PICALM, and TNK1 Genotypes in Aging With and Without Alzheimer’s Disease. Mol Neurobiol. 2018;55(5):4333–4344. doi:10.1007/s12035-017-0547-x

116. Šerý O, Hlinecká L, Povová J, Bonczek O, Zeman T, Janout V, Ambroz P, Khan NA, Balcar VJ. Arachidonate 5-lipoxygenase (ALOX5) gene polymorphism is associated with Alzheimer’s disease and body mass index. J Neurol Sci. 2016;362:27–32. doi:10.1016/j.jns.2016.01.022

117. Zhang S, Xiao T, Yu Y, Qiao Y, Xu Z, Geng J, Liang Y, Mei Y, Dong Q, Wang B, et al. The extracellular matrix enriched with membrane metalloendopeptidase and insulin-degrading enzyme suppresses the deposition of amyloid-beta peptide in Alzheimer’s disease cell models. J Tissue Eng Regen Med. 2019;13(10):1759–1769. doi:10.1002/term.2906

118. Pekny M, Johansson CB, Eliasson C, Stakeberg J, Wallén A, Perlmann T, Lendahl U, Betsholtz C, Berthold CH, Frisén J. Abnormal reaction to central nervous system injury in mice lacking glial fibrillary acidic protein and vimentin. J Cell Biol. 1999;145(3):503–514. doi:10.1083/jcb.145.3.50

119. Griffiths MR, Botto M, Morgan BP, Neal JW, Gasque P. CD93 regulates central nervous system inflammation in two mouse models of autoimmune encephalomyelitis. Immunology. 2018;155(3):346–355. doi:10.1111/imm.12974

120. Martínez A, Santiago JL, Cénit MC, de Las Heras V, de la Calle H, Fernández-Arquero M, Arroyo R, de la Concha EG, Urcelay E. IFIH1-GCA-KCNH7 locus: influence on multiple sclerosis risk. Eur J Hum Genet. 2008;16(7):861–864. doi:10.1038/ejhg.2008.16

121. Ghavimi R, Alsahebfosoul F, Salehi R, Kazemi M, Etemadifar M, Zavaran Hosseini A. High-resolution melting curve analysis of polymorphisms within CD58, CD226, HLA-G genes and association with multiple sclerosis susceptibility in a subset of Iranian population: a case-control study. Acta Neurol Belg. 2020;120(3):645–652. doi:10.1007/s13760-018-0992-y

122. Mars LT, Gautron AS, Novak J, Beaudoin L, Diana J, Liblau RS, Lehuen A. Invariant NKT cells regulate experimental autoimmune encephalomyelitis and infiltrate the central nervous system in a CD1d-independent manner. J Immunol. 2008;181(4):2321–2329. doi:10.4049/jimmunol.181.4.2321

123. Lian Z, Liu J, Shi Z, Chen H, Zhang Q, Feng H, Du Q, Miao X, Zhou H. Association of TNFSF4 Polymorphisms with Neuromyelitis Optica Spectrum Disorders in a Chinese Population. J Mol Neurosci. 2017;63(3-4):396–402. doi:10.1007/s12031-017-0990-1

124. Santos R, Linker SB, Stern S, Mendes APD, Shokhirev MN, Erikson G, Randolph-Moore L, Racha V, Kim Y, Kelsoe JR, et al. Deficient LEF1 expression is associated with lithium resistance and hyperexcitability in neurons derived from bipolar disorder patients. Mol Psychiatry. 2021;26(6):2440–2456. doi:10.1038/s41380-020-00981-3

125. Leitner GR, Wenzel TJ, Marshall N, Gates EJ, Klegeris A. Targeting toll-like receptor 4 to modulate neuroinflammation in central nervous system disorders. Expert Opin Ther Targets. 2019;23(10):865–882. doi:10.1080/14728222.2019.1676416

126. Bielecki B, Jatczak-Pawlik I, Wolinski P, Bednarek A, Glabinski A. Central Nervous System and Peripheral Expression of CCL19, CCL21 and Their Receptor CCR7 in Experimental Model of Multiple Sclerosis. Arch Immunol Ther Exp (Warsz). 2015;63(5):367–376. doi:10.1007/s00005-015-0339-9

127. Tejera-Alhambra M, Casrouge A, de Andrés C, Ramos-Medina R, Alonso B, Vega J, Albert ML, Sánchez-Ramón S. Low DPP4 expression and activity in multiple sclerosis. Clin Immunol. 2014;150(2):170–183. doi:10.1016/j.clim.2013.11.011

128. Soares JL, Oliveira EM, Pontillo A. Variants in NLRP3 and NLRC4 inflammasome associate with susceptibility and severity of multiple sclerosis. Mult Scler Relat Disord. 2019;29:26–34. doi:10.1016/j.msard.2019.01.023

129. Singh AS, Chandra R, Guhathakurta S, Sinha S, Chatterjee A, Ahmed S, Ghosh S, Rajamma U. Genetic association and gene-gene interaction analyses suggest likely involvement of ITGB3 and TPH2 with autism spectrum disorder (ASD) in the Indian population. Prog Neuropsychopharmacol Biol Psychiatry. 2013;45:131–143. doi:10.1016/j.pnpbp.2013.04.015

130. Eshraghi M, Ramírez-Jarquín UN, Shahani N, Nuzzo T, De Rosa A, Swarnkar S, Galli N, Rivera O, Tsaprailis G, Scharager-Tapia C, et al. RasGRP1 is a causal factor in the development of l-DOPA-induced dyskinesia in Parkinson’s disease. Sci Adv. 2020;6(18):eaaz7001. doi:10.1126/sciadv.aaz7001

131. Hayward JH, Lee SJ. A Decade of Research on TLR2 Discovering Its Pivotal Role in Glial Activation and Neuroinflammation in Neurodegenerative Diseases. Exp Neurobiol. 2014;23(2):138–147. doi:10.5607/en.2014.23.2.138

132. Cimino PJ, Yang Y, Li X, Hemingway JF, Cherne MK, Khademi SB, Fukui Y, Montine KS, Montine TJ, Keene CD. et al. Ablation of the microglial protein DOCK2 reduces amyloid burden in a mouse model of Alzheimer’s disease. Exp Mol Pathol. 2013;94(2):366–371. doi:10.1016/j.yexmp.2013.01.002

133. Hagan N, Kane JL, Grover D, Woodworth L, Madore C, Saleh J, Sancho J, Liu J, Li Y, Proto J, et al. CSF1R signaling is a regulator of pathogenesis in progressive MS. Cell Death Dis. 2020;11(10):904. doi:10.1038/s41419-020-03084-7

134. Zhou Z, Chen F, Zhong S, Zhou Y, Zhang R, Kang K, Zhang X, Xu Y, Zhao M, Zhao C. Molecular identification of protein kinase C beta in Alzheimer’s disease. Aging (Albany NY). 2020;12(21):21798–21808. doi:10.18632/aging.103994

135. Zhang ZH, Wang YR, Li F, Liu XL, Zhang H, Zhu ZZ, Huang H, Xu XH. Circ-camk4 involved in cerebral ischemia/reperfusion induced neuronal injury. Sci Rep. 2020;10(1):7012. doi:10.1038/s41598-020-63686-1

136. Yu M, Ma X, Jiang D, Wang L, Zhan Q, Zhao J. CXC chemokine ligand 5 (CXCL5) disrupted the permeability of human brain microvascular endothelial cells via regulating p38 signal. Microbiol Immunol. 2021;65(1):40–47. doi:10.1111/1348-0421.12854

137. Dobri AM, Dudău M, Enciu AM, Hinescu ME. CD36 in Alzheimer’s Disease: An Overview of Molecular Mechanisms and Therapeutic Targeting. Neuroscience. 2021;453:301–311. doi:10.1016/j.neuroscience.2020.11.003

138. Walker DG, Tang TM, Mendsaikhan A, Tooyama I, Serrano GE, Sue LI, Beach TG, Lue LF. Patterns of Expression of Purinergic Receptor P2RY12, a Putative Marker for Non-Activated Microglia, in Aged and Alzheimer’s Disease Brains. Int J Mol Sci. 2020;21(2):678. doi:10.3390/ijms21020678

139. Yue J, Li W, Liang C, Chen B, Chen X, Wang L, Zang Z, Yu S, Liu S, Li S, et al. Activation of LILRB2 signal pathway in temporal lobe epilepsy patients and in a pilocarpine induced epilepsy model. Exp Neurol. 2016;285(Pt A):51–60. doi:10.1016/j.expneurol.2016.09.006

140. Seidi OA, Semra YK, Sharief MK. Expression of CD5 on B lymphocytes correlates with disease activity in patients with multiple sclerosis. J Neuroimmunol. 2002;133(1-2):205–210. doi:10.1016/s0165-5728(02)00360-0

141. Huo YX, Huang L, Zhang DF, Yao YG, Fang YR, Zhang C, Luo XJ. Identification of SLC25A37 as a major depressive disorder risk gene. J Psychiatr Res. 2016;83:168–175. doi:10.1016/j.jpsychires.2016.09.011

142. Yoon G, Shah SA, Ali T, Kim MO. The Adiponectin Homolog Osmotin Enhances Neurite Outgrowth and Synaptic Complexity via AdipoR1/NgR1 Signaling in Alzheimer’s Disease. Mol Neurobiol. 2018;55(8):6673–6686. doi:10.1007/s12035-017-0847-1

143. Kalinowska A, Losy J. PECAM-1, a key player in neuroinflammation. Eur J Neurol. 2006;13(12):1284–1290. doi:10.1111/j.1468-1331.2006.01640.x

144. Lee JK, Bou Dagher J. Regulator of G-protein Signaling (RGS)1 and RGS10 Proteins as Potential Drug Targets for Neuroinflammatory and Neurodegenerative Diseases. AAPS J. 2016;18(3):545–549. doi:10.1208/s12248-016-9883-4

145. Kim JO, Lee KO, Kim HW, Park HS, Kim J, Sung JH, Oh D, Kim OJ, Kim NK. ssociation between KCNQ2, TCF4 and RGS18 polymorphisms and silent brain infarction based on wholeLexome sequencing. Mol Med Rep. 2020;21(4):1973–1983. doi:10.3892/mmr.2020.10975

146. Smith AR, Smith RG, Burrage J, Troakes C, Al-Sarraj S, Kalaria RN, Sloan C, Robinson AC, Mill J, Lunnon K. A cross-brain regions study of ANK1 DNA methylation in different neurodegenerative diseases. Neurobiol Aging. 2019;74:70–76. doi:10.1016/j.neurobiolaging.2018.09.024

147. Liu QY, Lei JX, Sikorska M, Liu R. A novel brain-enriched E3 ubiquitin ligase RNF182 is up regulated in the brains of Alzheimer’s patients and targets ATP6V0C for degradation. Mol Neurodegener. 2008;3:4. doi:10.1186/1750-1326-3-4

148. Fang Y, Jiang Q, Li S, Zhu H, Xu R, Song N, Ding X, Liu J, Chen M, Song M, et al. Opposing functions of β-arrestin 1 and 2 in Parkinson’s disease via microglia inflammation and Nprl3. Cell Death Differ. 2021;28(6):1822–1836. doi:10.1038/s41418-020-00704-9

149. Peroni S, Sorosina M, Malhotra S, Clarelli F, Osiceanu AM, Ferrè L, Roostaei T, Rio J, Midaglia L, Villar LM, et al. A pharmacogenetic study implicates NINJ2 in the response to Interferon-β in multiple sclerosis. Mult Scler. 2020;26(9):1074–1082. doi:10.1177/1352458519851428

150. Lioudyno V, Abdurasulova I, Negoreeva I, Stoliarov I, Kudriavtsev I, Serebryakova M, Klimenko V, Lioudyno M. A common genetic variant rs2821557 in KCNA3 is linked to the severity of multiple sclerosis. J Neurosci Res. 2021;99(1):200–208. doi:10.1002/jnr.24596

151. Fehér Á, Juhász A, László A, Pákáski M, Kálmán J, Janka Z. Association between the ABCG2 C421A polymorphism and Alzheimer’s disease. Neurosci Lett. 2013;550:51–54. doi:10.1016/j.neulet.2013.06.044

152. Hou XH, Bi YL, Tan MS, Xu W, Li JQ, Shen XN, Dou KX, Tan CC, Tan L. Genome-wide association study identifies Alzheimer’s risk variant in MS4A6A influencing cerebrospinal fluid sTREM2 levels. Neurobiol Aging. 2019;84:241.e13–241.e20. doi:10.1016/j.neurobiolaging.2019.05.008

153. Cong Y, So V, Tijssen MAJ, Verbeek DS, Reggiori F, Mauthe M. WDR45, one gene associated with multiple neurodevelopmental disorders. Autophagy. 2021;17(12):3908–3923. doi:10.1080/15548627.2021.1899669

154. Gao Y, Wilson GR, Salce N, Romano A, Mellick GD, Stephenson SEM, Lockhart PJ.Genetic Analysis of RAB39B in an Early-Onset Parkinson’s Disease Cohort. Front Neurol. 2020;11:523. doi:10.3389/fneur.2020.00523

155. Rovelet-Lecrux A, Feuillette S, Miguel L, Schramm C, Pernet S, Quenez O, Ségalas-Milazzo I, Guilhaudis L, Rousseau S, Riou G, et al. Impaired SorLA maturation and trafficking as a new mechanism for SORL1 missense variants in Alzheimer disease. Acta Neuropathol Commun. 2021;9(1):196. doi:10.1186/s40478-021-01294-4

156. Hutcheson HB, Olson LM, Bradford Y, Folstein SE, Santangelo SL, Sutcliffe JS, Haines JL. Examination of NRCAM, LRRN3, KIAA0716, and LAMB1 as autism candidate genes. BMC Med Genet. 2004;5:12. doi:10.1186/1471-2350-5-12

157. Barone R, Aiello C, Race V, Morava E, Foulquier F, Riemersma M, Passarelli C, Concolino D, Carella M, Santorelli F, et al. DPM2-CDG: a muscular dystrophy-dystroglycanopathy syndrome with severe epilepsy. Ann Neurol. 2012;72(4):550–558. doi:10.1002/ana.23632

158. Gurnett CA, Veile R, Zempel J, Blackburn L, Lovett M, Bowcock A. Disruption of sodium bicarbonate transporter SLC4A10 in a patient with complex partial epilepsy and mental retardation. Arch Neurol. 2008;65(4):550–553. doi:10.1001/archneur.65.4.550

159. Ge P, Dawson VL, Dawson TM. PINK1 and Parkin mitochondrial quality control: a source of regional vulnerability in Parkinson’s disease. Mol Neurodegener. 2020;15(1):20. doi:10.1186/s13024-020-00367-7

160. Anirudhan A, Angulo-Bejarano PI, Paramasivam P, Manokaran K, Kamath SM, Murugesan R, Sharma A, Ahmed SSSJ. RPL6: A Key Molecule Regulating Zinc- and Magnesium-Bound Metalloproteins of Parkinson’s Disease. Front Neurosci. 2021;15:631892. doi:10.3389/fnins.2021.631892

161. Shi Z, Zhang Q, Chen H, Lian Z, Liu J, Feng H, Miao X, Du Q, Zhou H. STAT4 Polymorphisms are Associated with Neuromyelitis Optica Spectrum Disorders. Neuromolecular Med. 2017;19(4):493–500. doi:10.1007/s12017-017-8463-9

162. Kim SH, Shin SY, Lee KY, Joo EJ, Song JY, Ahn YM, Lee YH, Kim YS. The genetic association of DUSP6 with bipolar disorder and its effect on ERK activity. Prog Neuropsychopharmacol Biol Psychiatry. 2012;37(1):41–49. doi:10.1016/j.pnpbp.2011.11.014

163. Chow N, Bell RD, Deane R, Streb JW, Chen J, Brooks A, Van Nostrand W, Miano JM, Zlokovic BV. Serum response factor and myocardin mediate arterial hypercontractility and cerebral blood flow dysregulation in Alzheimer’s phenotype. Proc Natl Acad Sci U S A. 2007;104(3):823–828. doi:10.1073/pnas.0608251104

164. Ji H, Wang Y, Liu G, Chang L, Chen Z, Zhou D, Xu X, Cui W, Hong Q, Jiang L, et al. Elevated OPRD1 promoter methylation in Alzheimer’s disease patients. PLoS One. 2017;12(3):e0172335. doi:10.1371/journal.pone.0172335

165. Hori A, Ai T, Isshiki M, Motoi Y, Yano K, Tabe Y, Hattori N, Miida T. Novel Variants in the CLCN1, RYR2, and DCTN1 Found in Elderly Japanese Dementia Patients: A Case Series. Geriatrics (Basel). 2021;6(1):14. doi:10.3390/geriatrics6010014

166. Skinner D, Marro BS, Lane TE. Chemokine CXCL10 and Coronavirus-Induced Neurologic Disease. Viral Immunol. 2019;32(1):25–37. doi:10.1089/vim.2018.0073

167. Massopust R, Juros D, Shapiro D, Lopes M, Haldar SM, Taetzsch T, Valdez G. KLF15 overexpression in myocytes fails to ameliorate ALS-related pathology or extend the lifespan of SOD1G93A mice. Neurobiol Dis. 2022;162:105583. doi:10.1016/j.nbd.2021.105583

168. Raghavan NS, Dumitrescu L, Mormino E, Mahoney ER, Lee AJ, Gao Y, Bilgel M, Goldstein D, Harrison T, Engelman CD, et al. Association Between Common Variants in RBFOX1, an RNA-Binding Protein, and Brain Amyloidosis in Early and Preclinical Alzheimer Disease. JAMA Neurol. 2020;77(10):1288–1298. doi:10.1001/jamaneurol.2020.1760

169. Devos D, Lejeune S, Cormier-Dequaire F, Tahiri K, Charbonnier-Beaupel F, Rouaix N, Duhamel A, Sablonnière B, Bonnet AM, Bonnet C, et al. Dopa-decarboxylase gene polymorphisms affect the motor response to L-dopa in Parkinson’s disease. Parkinsonism Relat Disord. 2014;20(2):170–175. doi:10.1016/j.parkreldis.2013.10.017

170. Lin Q, Cao Y, Gao J. Decreased expression of the APOA1-APOC3-APOA4 gene cluster is associated with risk of Alzheimer’s disease. Drug Des Devel Ther. 2015;9:5421–5431. doi:10.2147/DDDT.S89279

171. Hogan R, Flamier A, Nardini E, Bernier G. The Role of BMI1 in Late-Onset Sporadic Alzheimer’s Disease. Genes (Basel). 2020;11(7):825. doi:10.3390/genes11070825

172. Kang WS, Park JK, Kim SK, Park HJ, Lee SM, Song JY, Chung JH, Kim JW. Genetic variants of GRIA1 are associated with susceptibility to schizophrenia in Korean population. Mol Biol Rep. 2012;39(12):10697–10703. doi:10.1007/s11033-012-1960-x

173. Zhou XG, He H, Wang PJ. A critical role for miRL135aL5pLmediated regulation of SLC24A2 in neuropathic pain. Mol Med Rep. 2020;22(3):2115–2122. doi:10.3892/mmr.2020.11262

174. Shepard R, Heslin K, Hagerdorn P, Coutellier L. Downregulation of Npas4 in parvalbumin interneurons and cognitive deficits after neonatal NMDA receptor blockade: relevance for schizophrenia. Transl Psychiatry. 2019;9(1):99. doi:10.1038/s41398-019-0436-3

175. Sjögren M, Minthon L, Davidsson P, Granérus A-K, Clarberg A, Vanderstichele H, Vanmechelen E, Wallin A, Blennow K. CSF levels of tau, beta-amyloid(1-42) and GAP-43 in frontotemporal dementia, other types of dementia and normal aging. J Neural Transm (Vienna). 2000;107(5):563–579. doi:10.1007/s007020070079

176. Barki M, Xue H. GABRB2, a key player in neuropsychiatric disorders and beyond. Gene. 2022;809:146021. doi:10.1016/j.gene.2021.146021

177. Eckhardt J, Döbbeler M, König C, Kuczera K, Kuhnt C, Ostalecki C, Zinser E, Mak TW, Steinkasserer A, Lechmann M. Thymic stromal lymphopoietin deficiency attenuates experimental autoimmune encephalomyelitis. Clin Exp Immunol. 2015;181(1):51–64. doi:10.1111/cei.12621

178. Buono RJ, Bradfield JP, Wei Z, Sperling MR, Dlugos DJ, Privitera MD, French JA, Lo W, Cossette P, Schachter SC, et al. Genetic Variation in PADI6-PADI4 on 1p36.13 Is Associated with Common Forms of Human Generalized Epilepsy. Genes (Basel). 2021;12(9):1441. doi:10.3390/genes12091441

179. Bose S, Cho J. Role of chemokine CCL2 and its receptor CCR2 in neurodegenerative diseases. Arch Pharm Res. 2013;36(9):1039–1050. doi:10.1007/s12272-013-0161-z

180. Wang P, Li XL, Cao ZH. STC1 ameliorates cognitive impairment and neuroinflammation of Alzheimer’s disease mice via inhibition of ERK1/2 pathway. Immunobiology. 2021;226(3):152092. doi:10.1016/j.imbio.2021.152092

181. Xu FL, Wang BJ, Yao J. Association between the SLC6A4 gene and schizophrenia: an updated meta-analysis. Neuropsychiatr Dis Treat. 2018;15:143–155. doi:10.2147/NDT.S190563

182. Kolbinger F, Huppertz C, Mir A, Padova FD. IL-17A and Multiple Sclerosis: Signaling Pathways, Producing Cells and Target Cells in the Central Nervous System. Curr Drug Targets. 2016;17(16):1882–1893. doi:10.2174/1389450117666160307144027

183. Mercati O, Huguet G, Danckaert A, André-Leroux G, Maruani A, Bellinzoni M, Rolland T, Gouder L, Mathieu A, Buratti J, et al. CNTN6 mutations are risk factors for abnormal auditory sensory perception in autism spectrum disorders. Mol Psychiatry. 2017;22(4):625–633. doi:10.1038/mp.2016.61

184. Sangu N, Shimojima K, Takahashi Y, Ohashi T, Tohyama J, Yamamoto T. A 7q31.33q32.1 microdeletion including LRRC4 and GRM8 is associated with severe intellectual disability and characteristics of autism. Hum Genome Var. 2017;4:17001. doi:10.1038/hgv.2017.1

185. McLean CK, Narayan S, Lin SY, Rai N, Chung Y, Hipolito MS, Cascella NG, Nurnberger JI Jr, Ishizuka K, Sawa AS, et al. Lithium-associated transcriptional regulation of CRMP1 in patient-derived olfactory neurons and symptom changes in bipolar disorder. Transl Psychiatry. 2018;8(1):81. doi:10.1038/s41398-018-0126-6

186. Wang Q, Wang C, Ji B, Zhou J, Yang C, Chen J. Hapln2 in Neurological Diseases and Its Potential as Therapeutic Target. Front Aging Neurosci. 2019;11:60. doi:10.3389/fnagi.2019.00060

187. Xia Z, Chibnik LB, Glanz BI, Liguori M, Shulman JM, Tran D, Khoury SJ, Chitnis T, Holyoak T, Weiner HL, et al. A putative Alzheimer’s disease risk allele in PCK1 influences brain atrophy in multiple sclerosis. PLoS One. 2010;5(11):e14169. doi:10.1371/journal.pone.0014169

188. Kang JQ, Shen W, Zhou C, Xu D, Macdonald RL. The human epilepsy mutation GABRG2(Q390X) causes chronic subunit accumulation and neurodegeneration. Nat Neurosci. 2015;18(7):988–996. doi:10.1038/nn.4024

189. Pascolini G, Majore S, Valiante M, Bottillo I, Laino L, Agolini E, Novelli A, Grammatico B, Calvani M, Grammatico P. Autism spectrum disorder in a patient with a genomic rearrangement that only involves the EPHA5 gene. Psychiatr Genet. 2019;29(3):86–90. doi:10.1097/YPG.0000000000000217

190. Sato C, Hane M. Mental disorders and an acidic glycan-from the perspective of polysialic acid (PSA/polySia) and the synthesizing enzyme, ST8SIA2. Glycoconj J. 2018;35(4):353–373. doi:10.1007/s10719-018-9832-9

191. Reich A, Spering C, Gertz K, Harms C, Gerhardt E, Kronenberg G, Nave KA, Schwab M, Tauber SC, Drinkut A, et al. Fas/CD95 regulatory protein Faim2 is neuroprotective after transient brain ischemia. J Neurosci. 2011;31(1):225–233. doi:10.1523/JNEUROSCI.2188-10.2011

192. Dos Santos EUD, Sampaio TF, Tenório Dos Santos AD, Bezerra Leite FC, da Silva RC, Crovella S, Asano AGC, Asano NMJ, de Souza PRE. The influence of SLC6A3 and DRD2 polymorphisms on levodopa-therapy in patients with sporadic Parkinson’s disease. J Pharm Pharmacol. 2019;71(2):206–212. doi:10.1111/jphp.13031

193. Bakalkin G, Watanabe H, Jezierska J, Depoorter C, Verschuuren-Bemelmans C, Bazov I, Artemenko KA, Yakovleva T, Dooijes D, Van de Warrenburg BP, et al. Prodynorphin mutations cause the neurodegenerative disorder spinocerebellar ataxia type 23. Am J Hum Genet. 2010;87(5):593–603. doi:10.1016/j.ajhg.2010.10.001

194. Skonieczna-Żydecka K, Jamioł-Milc D, Borecki K, Stachowska E, Zabielska P, Kamińska M, Karakiewicz B. The Prevalence of Insomnia and the Link between Iron Metabolism Genes Polymorphisms, TF rs1049296 C>T, TF rs3811647 G>A, TFR rs7385804 A>C, HAMP rs10421768 A>G and Sleep Disorders in Polish Individuals with ASD. Int J Environ Res Public Health. 2020;17(2):400. doi:10.3390/ijerph17020400

195. Koper OM, Kamińska J, Sawicki K, Kemona H. CXCL9, CXCL10, CXCL11, and their receptor (CXCR3) in neuroinflammation and neurodegeneration. Adv Clin Exp Med. 2018;27(6):849–856. doi:10.17219/acem/68846

196. Håvik B, Le Hellard S, Rietschel M, Lybæk H, Djurovic S, Mattheisen M, Mühleisen TW, Degenhardt F, Priebe L, et al. The complement control-related genes CSMD1 and CSMD2 associate to schizophrenia. Biol Psychiatry. 2011;70(1):35–42. doi:10.1016/j.biopsych.2011.01.030

197. Choi S, Han KM, Kang J, Won E, Chang HS, Tae WS, Son KR, Kim SJ, Lee MS, Ham BJ. Effects of a Polymorphism of the Neuronal Amino Acid Transporter SLC6A15 Gene on Structural Integrity of White Matter Tracts in Major Depressive Disorder. PLoS One. 2016;11(10):e0164301. doi:10.1371/journal.pone.0164301

198. Mez J, Chung J, Jun G, Kriegel J, Bourlas AP, Sherva R, Logue MW, Barnes LL, Bennett DA, Buxbaum JD, et al. Two novel loci, COBL and SLC10A2, for Alzheimer’s disease in African Americans. Alzheimers Dement. 2017;13(2):119–129. doi:10.1016/j.jalz.2016.09.002

199. Gao K, Song Z, Liang H, Zheng W, Deng X, Yuan Y, Zhao Y, Deng H. Genetic analysis of the ATP1B4 gene in Chinese Han patients with Parkinson’s disease. Mol Biol Rep. 2014;41(4):2307–2311. doi:10.1007/s11033-014-3084-y

200. Rendina A, Drongitis D, Donizetti A, Fucci L, Milan G, Tripodi F, Giustezza F, Postiglione A, Pappatà S, Ferrari R, et al. CD33 and SIGLECL1 Immunoglobulin Superfamily Involved in Dementia. J Neuropathol Exp Neurol. 2020;79(8):891–901. doi:10.1093/jnen/nlaa055

201. Lohoff FW. Genetic variants in the vesicular monoamine transporter 1 (VMAT1/SLC18A1) and neuropsychiatric disorders. Methods Mol Biol. 2010;637:165–180. doi:10.1007/978-1-60761-700-6_9

202. Wan B, Xu WJ, Xu WN, Zhan P, Wu GN, Jin JJ, Xi GM, Yin JT, Zhang H, Chen YK, et al. Plasma long noncoding RNA IL-7R as a prognostic biomarker for clinical outcomes in patients with acute respiratory distress syndrome. Clin Respir J. 2018;12(4):1607–1614. doi:10.1111/crj.12717

203. Chen L, Li L, Song Y, Lv T. Blocking SphK1/S1P/S1PR1 Signaling Pathway Alleviates Lung Injury Caused by Sepsis in Acute Ethanol Intoxication Mice. Inflammation. 2021;44(6):2170–2179. doi:10.1007/s10753-021-01490-3

204. Mathew B, Jacobson JR, Siegler JH, Moitra J, Blasco M, Xie L, Unzueta C, Zhou T, Evenoski C, Al-Sakka M, et al. Role of migratory inhibition factor in age-related susceptibility to radiation lung injury via NF-E2-related factor-2 and antioxidant regulation. Am J Respir Cell Mol Biol. 2013;49(2):269–278. doi:10.1165/rcmb.2012-0291OC

205. Osterholzer JJ, Olszewski MA, Murdock BJ, Chen GH, Erb-Downward JR, Subbotina N, Browning K, Lin Y, Morey RE, Dayrit JK, et al. Implicating exudate macrophages and Ly-6C(high) monocytes in CCR2-dependent lung fibrosis following gene-targeted alveolar injury. J Immunol. 2013;190(7):3447–3457. doi:10.4049/jimmunol.1200604

206. Jin Y, Wu W, Zhang W, Zhao Y, Wu Y, Ge G, Ba Y, Guo Q, Gao T, Chi X, et al. Involvement of EGF receptor signaling and NLRP12 inflammasome in fine particulate matter-induced lung inflammation in mice. Environ Toxicol. 2017;32(4):1121–1134. doi:10.1002/tox.22308

207. Villar J, Zhang H, Slutsky AS. Lung Repair and Regeneration in ARDS: Role of PECAM1 and Wnt Signaling. Chest. 2019;155(3):587–594. doi:10.1016/j.chest.2018.10.022a

208. Cao Y, Chen X, Liu Y, Zhang X, Zou Y, Li J. PIM1 inhibition attenuated endotoxin-induced acute lung injury through modulating ELK3/ICAM1 axis on pulmonary microvascular endothelial cells. Inflamm Res. 2021;70(1):89–98. doi:10.1007/s00011-020-01420-3

209. Zhu L, Chen Y, Chen M, Wang W. Mechanism of miR-204-5p in exosomes derived from bronchoalveolar lavage fluid on the progression of pulmonary fibrosis via AP1S2. Ann Transl Med. 2021;9(13):1068. doi:10.21037/atm-20-8033

210. Nadeem A, Al-Harbi NO, Ahmad SF, Al-Harbi MM, Alhamed AS, Alfardan AS, Assiri MA, Ibrahim KE, Albassam H. Blockade of interleukin-2-inducible T-cell kinase signaling attenuates acute lung injury in mice through adjustment of pulmonary Th17/Treg immune responses and reduction of oxidative stress. Int Immunopharmacol. 2020;83:106369. doi:10.1016/j.intimp.2020.106369

211. Adegunsoye A, Hrusch CL, Bonham CA, Jaffery MR, Blaine KM, Sullivan M, Churpek MM, Strek ME, Noth I, Sperling AI. Skewed Lung CCR4 to CCR6 CD4+ T Cell Ratio in Idiopathic Pulmonary Fibrosis Is Associated with Pulmonary Function. Front Immunol. 2016;7:516. doi:10.3389/fimmu.2016.00516

212. Chen Y, Zhang H, Li F, Wang X. Inhibition of CX3C receptor 1-mediated autophagy in macrophages alleviates pulmonary fibrosis in hyperoxic lung injury. Life Sci. 2020;259:118286. doi:10.1016/j.lfs.2020.118286

213. Mahlooji MA, Heshmati A, Kheiripour N, Ghasemi H, Asl SS, Solgi G, Ranjbar A, Hosseini A. Evaluation of Protective Effects of Curcumin and Nanocurcumin on Aluminium PhosphideLInduced Subacute Lung Injury in Rats: Modulation of Oxidative Stress through SIRT1/FOXO3 Signalling Pathway. Drug Res (Stuttg). 2021;10.1055/a-1647-2418. doi:10.1055/a-1647-2418

214. Liu Y, Tong C, Xu Y, Cong P, Liu Y, Shi L, Shi X, Zhao Y, Bi G, Jin H, et al. CD28 Deficiency Ameliorates Blast Exposure-Induced Lung Inflammation, Oxidative Stress, Apoptosis, and T Cell Accumulation in the Lungs via the PI3K/Akt/FoxO1 Signaling Pathway. Oxid Med Cell Longev. 2019;2019:4848560. doi:10.1155/2019/4848560

215. Bouch S, O’Reilly M, de Haan JB, Harding R, Sozo F. Does lack of glutathione peroxidase 1 gene expression exacerbate lung injury induced by neonatal hyperoxia in mice?. Am J Physiol Lung Cell Mol Physiol. 2017;313(1):L115–L125. doi:10.1152/ajplung.00039.2016

216. Rushefski M, Aplenc R, Meyer N, Li M, Feng R, Lanken PN, Gallop R, Bellamy S, Localio AR, Feinstein SI, et al. Novel variants in the PRDX6 Gene and the risk of Acute Lung Injury following major trauma. BMC Med Genet. 2011;12:77. doi:10.1186/1471-2350-12-77

217. Tian Y, Lv J, Su Z, Wu T, Li X, Hu X, Zhang J, Wu L. LRRK2 plays essential roles in maintaining lung homeostasis and preventing the development of pulmonary fibrosis. Proc Natl Acad Sci U S A. 2021;118(35):e2106685118. doi:10.1073/pnas.2106685118

218. Zhou Z, Zhu Y, Gao G, Zhang Y. Long noncoding RNA SNHG16 targets miR-146a-5p/CCL5 to regulate LPS-induced WI-38 cell apoptosis and inflammation in acute pneumonia. Life Sci. 2019;228:189–197. doi:10.1016/j.lfs.2019.05.008

219. Yang M, Gao XR, Meng YN, Shen F, Chen YP. ETS1 Ameliorates Hyperoxia-Induced Alveolar Epithelial Cell Injury by Regulating the TGM2-Mediated Wnt/β-Catenin Pathway. Lung. 2021;199(6):681–690. doi:10.1007/s00408-021-00489-9

220. Li B, Dong C, Wang G, Zheng H, Wang X, Bai C. Pulmonary epithelial CCR3 promotes LPS-induced lung inflammation by mediating release of IL-8. J Cell Physiol. 2011;226(9):2398–2405. doi:10.1002/jcp.22577

221. Bime C, Pouladi N, Sammani S, Batai K, Casanova N, Zhou T, Kempf CL, Sun X, Camp SM, Wang T, et al. Genome-Wide Association Study in African Americans with Acute Respiratory Distress Syndrome Identifies the Selectin P Ligand Gene as a Risk Factor. Am J Respir Crit Care Med. 2018;197(11):1421–1432. doi:10.1164/rccm.201705-0961OC

222. Li FJ, Surolia R, Li H, Wang Z, Liu G, Kulkarni T, Massicano AVF, Mobley JA, Mondal S, de Andrade JA, et al. Citrullinated vimentin mediates development and progression of lung fibrosis. Sci Transl Med. 2021;13(585):eaba2927. doi:10.1126/scitranslmed.aba2927

223. Kim JH, Chung DH. CD1d-restricted IFN-γ-secreting NKT cells promote immune complex-induced acute lung injury by regulating macrophage-inflammatory protein-1α production and activation of macrophages and dendritic cells. J Immunol. 2011;186(3):1432–1441. doi:10.4049/jimmunol.1003140

224. Retamal J, Sörensen J, Lubberink M, Suarez-Sipmann F, Borges JB, Feinstein R, Jalkanen S, Antoni G, Hedenstierna G, Roivainen A, et al. Feasibility of (68)Ga-labeled Siglec-9 peptide for the imaging of acute lung inflammation: a pilot study in a porcine model of acute respiratory distress syndrome. Am J Nucl Med Mol Imaging. 2016;6(1):18–31.

225. Ye R, Liu Z. ACE2 exhibits protective effects against LPS-induced acute lung injury in mice by inhibiting the LPS-TLR4 pathway. Exp Mol Pathol. 2020;113:104350. doi:10.1016/j.yexmp.2019.104350

226. Eltahir S, Ahmad KS, Al-Balawi MM, Bukhamsien H, Al-Mobaireek K, Alotaibi W, Al-Shamrani A. Lung disease associated with filamin A gene mutation: a case report. J Med Case Rep. 2016;10:97. doi:10.1186/s13256-016-0871-1

227. Trujillo G, Hartigan AJ, Hogaboam CM. T regulatory cells and attenuated bleomycin-induced fibrosis in lungs of CCR7-/-mice. Fibrogenesis Tissue Repair. 2010;3:18. doi:10.1186/1755-1536-3-18

228. Zientara A, Stephan M, von Hörsten S, Schmiedl A. Differential severity of LPS-induced lung injury in CD26/DPP4 positive and deficient F344 rats. Histol Histopathol. 2019;34(10):1151–1171. doi:10.14670/HH-18-117

229. He Y, Xu K, Wang Y, Chao X, Xu B, Wu J, Shen J, Ren W, Hu Y. AMPK as a potential pharmacological target for alleviating LPS-induced acute lung injury partly via NLRC4 inflammasome pathway inhibition. Exp Gerontol. 2019;125:110661. doi:10.1016/j.exger.2019.110661

230. Lee W, Hahn D, Sim H, Choo S, Lee S, Lee T, Bae JS. Inhibitory functions of cardamonin against particulate matter-induced lung injury through TLR2,4-mTOR-autophagy pathways. Fitoterapia. 2020;146:104724. doi:10.1016/j.fitote.2020.104724

231. Xu X, Su Y, Wu K, Pan F, Wang A. DOCK2 contributes to endotoxemia-induced acute lung injury in mice by activating proinflammatory macrophages. Biochem Pharmacol. 2021;184:114399. doi:10.1016/j.bcp.2020.114399

232. Wang TJ, Wu LF, Chen J, Zhu W, Wang H, Liu XL, Teng YQ. X-linked hyper-IgM syndrome complicated with interstitial pneumonia and liver injury: a new mutation locus in the CD40LG gene. Immunol Res. 2019;67(4-5):454–459. doi:10.1007/s12026-019-09098-4

233. Lagassé HA, Anidi IU, Craig JM, Limjunyawong N, Poupore AK, Mitzner W, Scott AL. Recruited monocytes modulate malaria-induced lung injury through CD36-mediated clearance of sequestered infected erythrocytes. J Leukoc Biol. 2016;99(5):659–671. doi:10.1189/jlb.4HI0315-130RRR

234. Zhu S, Song W, Sun Y, Zhou Y, Kong F. MiR-342 attenuates lipopolysaccharide-induced acute lung injury via inhibiting MAPK1 expression. Clin Exp Pharmacol Physiol. 2020;47(8):1448–1454. doi:10.1111/1440-1681.13315

235. Cao X, Zhang C, Zhang X, Chen Y, Zhang H. MiR-145 negatively regulates TGFBR2 signaling responsible for sepsis-induced acute lung injury. Biomed Pharmacother. 2019;111:852–858. doi:10.1016/j.biopha.2018.12.138

236. Xiao Q, Dong N, Yao X, Wu D, Lu Y, Mao F, Zhu J, Li J, Huang J, Chen A, et al. Bufexamac ameliorates LPS-induced acute lung injury in mice by targeting LTA4H. Sci Rep. 2016;6:25298. doi:10.1038/srep25298

237. Christofidou-Solomidou M, Kennel S, Scherpereel A, Wiewrodt R, Solomides CC, Pietra GG, Murciano JC, Shah SA, Ischiropoulos H, Albelda SM, et al. Vascular immunotargeting of glucose oxidase to the endothelial antigens induces distinct forms of oxidant acute lung injury: targeting to thrombomodulin, but not to PECAM-1, causes pulmonary thrombosis and neutrophil transmigration. Am J Pathol. 2002;160(3):1155–1169. doi:10.1016/S0002-9440(10)64935-8

238. Andersson-Sjöland A, Hallgren O, Rolandsson S, Weitoft M, Tykesson E, Larsson-Callerfelt AK, Rydell-Törmänen K, Bjermer L, Malmström A, Karlsson JC, et al. Versican in inflammation and tissue remodeling: the impact on lung disorders. Glycobiology. 2015;25(3):243–251. doi:10.1093/glycob/cwu120

239. Zou Z, Wang Q, Zhou M, Li W, Zheng Y, Li F, Zheng S, He Z. Protective effects of P2X7R antagonist in sepsis-induced acute lung injury in mice via regulation of circ_0001679 and circ_0001212 and downstream Pln, Cdh2, and Nprl3 expression. J Gene Med. 2020;22(12):e3261. doi:10.1002/jgm.3261

240. Qi F, Li Y, Yang X, Wu Y, Lin L, Liu X. Hsa_circ_0044226 knockdown attenuates progression of pulmonary fibrosis by inhibiting CDC27. Aging (Albany NY). 2020;12(14):14808–14818. doi:10.18632/aging.103543

241. Zhuang R, Yang X, Cai W, Xu R, Lv L, Sun Y, Guo Y, Ni J, Zhao G, Lu Z. MCTR3 reduces LPS-induced acute lung injury in mice via the ALX/PINK1 signaling pathway. Int Immunopharmacol. 2021;90:107142. doi:10.1016/j.intimp.2020.107142

242. Ahlfeld SK, Wang J, Gao Y, Snider P, Conway SJ. Initial Suppression of Transforming Growth Factor-β Signaling and Loss of TGFBI Causes Early Alveolar Structural Defects Resulting in Bronchopulmonary Dysplasia. Am J Pathol. 2016;186(4):777–793. doi:10.1016/j.ajpath.2015.11.024

243. Zimmerman JL, Dellinger RP, Straube RC, Levin JL. Phase I trial of the recombinant soluble complement receptor 1 in acute lung injury and acute respiratory distress syndrome. Crit Care Med. 2000;28(9):3149–3154. doi:10.1097/00003246-200009000-00004

244. Li P, Yao Y, Ma Y, Chen Y. MiR-150 attenuates LPS-induced acute lung injury via targeting AKT3. Int Immunopharmacol. 2019;75:105794. doi:10.1016/j.intimp.2019.105794

245. Fu C, Jiang L, Xu X, Zhu F, Zhang S, Wu X, Liu Z, Yang X, Li S. STAT4 knockout protects LPS-induced lung injury by increasing of MDSC and promoting of macrophage differentiation. Respir Physiol Neurobiol. 2016;223:16–22. doi:10.1016/j.resp.2015.11.016

246. Xie Y, Zheng H, Mou Z, Wang Y, Li X. High Expression of CXCL10/CXCR3 in Ventilator-Induced Lung Injury Caused by High Mechanical Power. Biomed Res Int. 2022;2022:6803154. doi:10.1155/2022/6803154

247. Kapur R, Kim M, Shanmugabhavananthan S, Liu J, Li Y, Semple JW. C-reactive protein enhances murine antibody-mediated transfusion-related acute lung injury. Blood. 2015;126(25):2747–2751. doi:10.1182/blood-2015-09-672592

248. You QH, Zhang D, Niu CC, Zhu ZM, Wang N, Yue Y, Sun GY.Expression of IL-17A and IL-17F in lipopolysaccharide-induced acute lung injury and the counteraction of anisodamine or methylprednisolone. Cytokine. 2014;66(1):78–86. doi:10.1016/j.cyto.2013.12.019

249. Lo BC, Gold MJ, Scheer S, Hughes MR, Cait J, Debruin E, Chu FSF, Walker DC, Soliman H, Rossi FM, et al. Loss of Vascular CD34 Results in Increased Sensitivity to Lung Injury. Am J Respir Cell Mol Biol. 2017;57(6):651–661. doi:10.1165/rcmb.2016-0386OC

250. Kreisel D, Sugimoto S, Tietjens J, Zhu J, Yamamoto S, Krupnick AS, Carmody RJ, Gelman AE. Bcl3 prevents acute inflammatory lung injury in mice by restraining emergency granulopoiesis. J Clin Invest. 2011;121(1):265–276. doi:10.1172/JCI42596

251. Lee JU, Chang HS, Lee HJ, Jung CA, Bae DJ, Song HJ, Park JS, Uh ST, Kim YH, Seo KH, et al. Upregulation of interleukin-33 and thymic stromal lymphopoietin levels in the lungs of idiopathic pulmonary fibrosis. BMC Pulm Med. 2017;17(1):39. doi:10.1186/s12890-017-0380-z

252. Yang J, Agarwal M, Ling S, Teitz-Tennenbaum S, Zemans RL, Osterholzer JJ, Sisson TH, Kim KK. Diverse Injury Pathways Induce Alveolar Epithelial Cell CCL2/12, Which Promotes Lung Fibrosis. Am J Respir Cell Mol Biol. 2020;62(5):622–632. doi:10.1165/rcmb.2019-0297OC

253. Scotton CJ, Krupiczojc MA, Königshoff M, Mercer PF, Lee YC, Kaminski N, Morser J, Post JM, Maher TM, Nicholson AG, et al. Increased local expression of coagulation factor X contributes to the fibrotic response in human and murine lung injury. J Clin Invest. 2009;119(9):2550–2563. doi:10.1172/JCI33288

254. Zhang Y, Shan P, Srivastava A, Li Z, Lee PJ. Endothelial Stanniocalcin 1 Maintains Mitochondrial Bioenergetics and Prevents Oxidant-Induced Lung Injury via Toll-Like Receptor 4. Antioxid Redox Signal. 2019;30(15):1775–1796. doi:10.1089/ars.2018.7514

255. Okamoto M, Izuhara K, Ohta S, Ono J, Hoshino T. Ability of Periostin as a New Biomarker of Idiopathic Pulmonary Fibrosis. Adv Exp Med Biol. 2019;1132:79–87. doi:10.1007/978-981-13-6657-4_9

256. Wang Y, Wang X, Li Y, Xue Z, Shao R, Li L, Zhu Y, Zhang H, Yang J. uanfei Baidu Decoction reduces acute lung injury by regulating infiltration of neutrophils and macrophages via PD-1/IL17A pathway. Pharmacol Res. 2022;176:106083. doi:10.1016/j.phrs.2022.106083

257. Luo L, Wang CC, Song XP, Wang HM, Zhou H, Sun Y, Wang XK, Hou S, Pei FY. Suppression of SMOC2 reduces bleomycin (BLM)-induced pulmonary fibrosis by inhibition of TGF-β1/SMADs pathway. Biomed Pharmacother. 2018;105:841–847. doi:10.1016/j.biopha.2018.03.058

258. Choi EH, Ehrmantraut M, Foster CB, Moss J, Chanock SJ. Association of common haplotypes of surfactant protein A1 and A2 (SFTPA1 and SFTPA2) genes with severity of lung disease in cystic fibrosis. Pediatr Pulmonol. 2006;41(3):255–262. doi:10.1002/ppul.20361

259. Shen W, Liu J, Fan M, Wang S, Zhang Y, Wen L, Wang R, Wei W, Li N, Zhang Y, et al. MiR-3202 protects smokers from chronic obstructive pulmonary disease through inhibiting FAIM2: An in vivo and in vitro study. Exp Cell Res. 2018;362(2):370–377. doi:10.1016/j.yexcr.2017.11.038

260. Okutomo K, Fujino N, Yamada M, Saito T, Ono Y, Okada Y, Ichinose M, Sugiura H. Increased LHX9 expression in alveolar epithelial type 2 cells of patients with chronic obstructive pulmonary disease. Respir Investig. 2022;60(1):119–128. doi:10.1016/j.resinv.2021.08.007

261. Ernst G, Dantas E, Sabatté J, Caro F, Salvado A, Grynblat P, Geffner J. Histidine-rich glycoprotein and idiopathic pulmonary fibrosis. Respir Med. 2015;109(12):1589–1591. doi:10.1016/j.rmed.2015.10.010

262. Leligdowicz A, Matthay MA. Carbonic Anhydrase IX: Scaring Away the Grim Reaper in Acute Lung Injury?. Am J Respir Cell Mol Biol. 2021;65(6):573–575. doi:10.1165/rcmb.2021-0310ED

263. Chang WA, Sheu CC, Liu KT, Shen JH, Yen MC, Kuo PL. Identification of mutations in SLC4A1, GP1BA and HFE in a family with venous thrombosis of unknown cause by next-generation sequencing. Exp Ther Med. 2018;16(5):4172–4180. doi:10.3892/etm.2018.6693

264. Henke PK, Pearce CG, Moaveni DM, Moore AJ, Lynch EM, Longo C, Varma M, Dewyer NA, Deatrick KB, Upchurch GR Jr, et al. Targeted deletion of CCR2 impairs deep vein thombosis resolution in a mouse model. J Immunol. 2006;177(5):3388–3397. doi:10.4049/jimmunol.177.5.3388

265. Yang G, Hooper WC. Physiological functions and clinical implications of fibrinogen-like 2: A review. World J Clin Infect Dis. 2013;3(3):37–46. doi:10.5495/wjcid.v3.i3.37

266. Hofmann S, Braun A, Pozgaj R, Morowski M, Vögtle T, Nieswandt B. Mice lacking the SLAM family member CD84 display unaltered platelet function in hemostasis and thrombosis. PLoS One. 2014;9(12):e115306. doi:10.1371/journal.pone.0115306

267. Semeraro F, Ammollo CT, Morrissey JH, Dale GL, Friese P, Esmon NL, Esmon CT. Extracellular histones promote thrombin generation through platelet-dependent mechanisms: involvement of platelet TLR2 and TLR4. Blood. 2011;118(7):1952–1961. doi:10.1182/blood-2011-03-343061

268. Liu SY, Yuan D, Sun RJ, Zhu JJ, Shan NN. Significant reductions in apoptosis-related proteins (HSPA6, HSPA8, ITGB3, YWHAH, and PRDX6) are involved in immune thrombocytopenia. J Thromb Thrombolysis. 2021;51(4):905–914. doi:10.1007/s11239-020-02310-5

269. Lee BC, Lin KH, Hu CY, Lo SC. Thromboelastography characterized CD36 null subjects as slow clot formation and indicative of hypocoagulability. Thromb Res. 2019;178:79–84. doi:10.1016/j.thromres.2019.04.006

270. Chandrakanthan M, Nguyen TQ, Hasan Z, Muralidharan S, Vu TM, Li AWL, Le UTN, Thi Thuy Ha H, Baik SH, Tan SH, et al. Deletion of Mfsd2b impairs thrombotic functions of platelets. Nat Commun. 2021;12(1):2286. doi:10.1038/s41467-021-22642-x

271. Thibord F, Hardy L, Ibrahim-Kosta M, Saut N, Pulcrano-Nicolas AS, Goumidi L, Civelek M, Eriksson P, Deleuze JF, Le Goff W, et al. A Genome Wide Association Study on plasma FV levels identified PLXDC2 as a new modifier of the coagulation process. J Thromb Haemost. 2019;17(11):1808–1814. doi:10.1111/jth.14562

272. Zhou J, Zhao X, Xie S, Zhou R. Transcriptome analysis of Klf15Lmediated inhibitory functions in a mouse deep venous thrombosis model. Int J Mol Med. 2020;45(6):1735–1752. doi:10.3892/ijmm.2020.4538

273. Zimmerman TS, Arroyove CM, Müller-Eberhard HJ. A blood coagulation abnormality in rabbits deficient in the sixth component of complement (C6) and its correction by purified C6. J Exp Med. 1971;134(6):1591–1600. doi:10.1084/jem.134.6.1591

274. MacQuarrie JL, Stafford AR, Yau JW, Leslie BA, Vu TT, Fredenburgh JC, Weitz JI. Histidine-rich glycoprotein binds factor XIIa with high affinity and inhibits contact-initiated coagulation. Blood. 2011;117(15):4134–4141. doi:10.1182/blood-2010-07-290551

275. Vichot AA, Zsengellér ZK, Shmukler BE, Adams ND, Dahl NK, Alper SL. Loss of kAE1 expression in collecting ducts of end-stage kidneys from a family with SLC4A1 G609R-associated distal renal tubular acidosis. Clin Kidney J. 2017;10(1):135–140. doi:10.1093/ckj/sfw074

276. Taylor Meadows KR, Steinberg MW, Clemons B, Stokes ME, Opiteck GJ, Peach R, Scott FL. Ozanimod (RPC1063), a selective S1PR1 and S1PR5 modulator, reduces chronic inflammation and alleviates kidney pathology in murine systemic lupus erythematosus. PLoS One. 2018;13(4):e0193236. doi:10.1371/journal.pone.0193236

277. Zhang J, Zhao X, Zhu H, Wang J, Ma J, Gu M. Apigenin Protects Against Renal Tubular Epithelial Cell Injury and Oxidative Stress by High Glucose via Regulation of NF-E2-Related Factor 2 (Nrf2) Pathway. Med Sci Monit. 2019;25:5280–5288. doi:10.12659/MSM.915038

278. Xia Y, Entman ML, Wang Y. CCR2 regulates the uptake of bone marrow-derived fibroblasts in renal fibrosis. PLoS One. 2013;8(10):e77493. doi:10.1371/journal.pone.0077493

279. Başaran Ö, Uncu N, Çakar N, Turanlı ET, Kiremitci S, Aydın F, Bayrakcı US. C3 glomerulopathy in NLRP12-related autoinflammatory disorder: case-based review. Rheumatol Int. 2018;38(8):1571–1576. doi:10.1007/s00296-018-4092-3

280. Završnik M, Kariž S, Makuc J, Šeruga M, Cilenšek I, Petrovič D. PECAM-1 Leu125Val (rs688) Polymorphism and Diabetic Nephropathy in Caucasians with Type 2 Diabetes Mellitus. Anal Cell Pathol (Amst). 2016;2016:3152967. doi:10.1155/2016/3152967

281. Chen WY, Wu SY, Lin TC, Lin SL, Wu-Hsieh BA. Human dendritic cell-specific ICAM-3-grabbing non-integrin downstream signaling alleviates renal fibrosis via Raf-1 activation in systemic candidiasis. Cell Mol Immunol. 2019;16(3):288–301. doi:10.1038/s41423-018-0161-5

282. von Vietinghoff S, Kurts C. Regulation and function of CX3CR1 and its ligand CX3CL1 in kidney disease. Cell Tissue Res. 2021;385(2):335–344. doi:10.1007/s00441-021-03473-0

283. Rudemiller N, Lund H, Jacob HJ, Geurts AM, Mattson DL; PhysGen Knockout Program. CD247 modulates blood pressure by altering T-lymphocyte infiltration in the kidney. Hypertension. 2014;63(3):559–564. doi:10.1161/HYPERTENSIONAHA.113.02191

284. Choi S, Kim DY, Ahn Y, Lee EJ, Park JH. Suppression of Foxo3-Gatm by miR-132-3p Accelerates Cyst Formation by Up-Regulating ROS in Autosomal Dominant Polycystic Kidney Disease. Biomol Ther (Seoul). 2021;29(3):311–320. doi:10.4062/biomolther.2020.197

285. Li Y, Jin L, Yan J, Zhang H, Zhang R, Hu C. CD28 Genetic Variants Increase Susceptibility to Diabetic Kidney Disease in Chinese Patients with Type 2 Diabetes: A Cross-Sectional Case Control Study. Mediators Inflamm. 2021;2021:5521050. doi:10.1155/2021/5521050

286. Jerotic D, Matic M, Suvakov S, Vucicevic K, Damjanovic T, Savic-Radojevic A, Pljesa-Ercegovac M, Coric V, Stefanovic A, et al. Association of Nrf2, SOD2 and GPX1 Polymorphisms with Biomarkers of Oxidative Distress and Survival in End-Stage Renal Disease Patients. Toxins (Basel). 2019;11(7):431. doi:10.3390/toxins11070431

287. Gao J, Wei L, Wei J, Yao G, Wang L, Wang M, Liu X, Dai C, et al. TLR1 polymorphism rs4833095 as a risk factor for IgA nephropathy in a Chinese Han population: A case-control study. Oncotarget. 2016;7(50):83031–83039. doi:10.18632/oncotarget.12965

288. Lee DH, Park JH, Han SB, Yoon DY, Jung YY, Hong JT. Peroxiredoxin 6 overexpression attenuates lipopolysaccharide-induced acute kidney injury. Oncotarget. 2017;8(31):51096–51107. doi:10.18632/oncotarget.17002

289. Krensky AM, Ahn YT. Mechanisms of disease: regulation of RANTES (CCL5) in renal disease. Nat Clin Pract Nephrol. 2007;3(3):164–170. doi:10.1038/ncpneph0418

290. Zhang H, He Y, He X, Wang L, Jin T, Yuan D. Three SNPs of FCRL3 and one SNP of MTMR3 are associated with immunoglobulin A nephropathy risk. Immunobiology. 2020;225(1):151869. doi:10.1016/j.imbio.2019.11.004

291. Di J, Jiang L, Zhou Y, Cao H, Fang L, Wen P, Li X, Dai C, Yang J. Ets-1 targeted by microrna-221 regulates angiotensin II-induced renal fibroblast activation and fibrosis. Cell Physiol Biochem. 2014;34(4):1063–1074. doi:10.1159/000366321

292. Shalev I, Liu H, Koscik C, Bartczak A, Javadi M, Wong KM, Maknojia A, He W, Liu MF, Diao J, et al. Targeted deletion of fgl2 leads to impaired regulatory T cell activity and development of autoimmune glomerulonephritis. J Immunol. 2008;180(1):249–260. doi:10.4049/jimmunol.180.1.249

293. Lee M, Park HS, Choi MY, Kim HZ, Moon SJ, Ha JY, Choi A, Park YW, Park JS, Shin EC, et al. Significance of Soluble CD93 in Type 2 Diabetes as a Biomarker for Diabetic Nephropathy: Integrated Results from Human and Rodent Studies. J Clin Med. 2020;9(5):1394. doi:10.3390/jcm9051394

294. Yang JQ, Singh AK, Wilson MT, Satoh M, Stanic AK, Park JJ, Hong S, Gadola SD, Mizutani A, Kakumanu SR, et al. Immunoregulatory role of CD1d in the hydrocarbon oil-induced model of lupus nephritis. J Immunol. 2003;171(4):2142–2153. doi:10.4049/jimmunol.171.4.2142

295. Zhou XJ, Cheng FJ, Qi YY, Zhao MH, Zhang H. A replication study from Chinese supports association between lupus-risk allele in TNFSF4 and renal disorder. Biomed Res Int. 2013;2013:597921. doi:10.1155/2013/597921

296. Hosur V, Cox ML, Burzenski LM, Riding RL, Alley L, Lyons BL, Kavirayani A, Martin KA, Cox GA, Johnson KR, et al. Retrotransposon insertion in the T-cell acute lymphocytic leukemia 1 (Tal1) gene is associated with severe renal disease and patchy alopecia in Hairpatches (Hpt) mice. PLoS One. 2013;8(1):e53426. doi:10.1371/journal.pone.0053426

297. Sun T, Dong W, Jiang G, Yang J, Liu J, Zhao L, Ma P. Cordyceps militaris Improves Chronic Kidney Disease by Affecting TLR4/NF-κB Redox Signaling Pathway. Oxid Med Cell Longev. 2019;2019:7850863. doi:10.1155/2019/7850863

298. Sakai N, Wada T, Yokoyama H, Lipp M, Ueha S, Matsushima K, Kaneko S. Secondary lymphoid tissue chemokine (SLC/CCL21)/CCR7 signaling regulates fibrocytes in renal fibrosis. Proc Natl Acad Sci U S A. 2006;103(38):14098–14103. doi:10.1073/pnas.0511200103

299. Kanasaki K. The role of renal dipeptidyl peptidase-4 in kidney disease: renal effects of dipeptidyl peptidase-4 inhibitors with a focus on linagliptin. Clin Sci (Lond). 2018;132(4):489–507. doi:10.1042/CS20180031

300. Wang Y, Gou R, Yu L, Wang L, Yang Z, Guo Y, Tang L.Activation of the NLRC4 inflammasome in renal tubular epithelial cell injury in diabetic nephropathy. Exp Ther Med. 2021;22(2):814. doi:10.3892/etm.2021.10246

301. Zhou XJ, Nath SK, Qi YY, Sun C, Hou P, Zhang YM, Lv JC, Shi SF, Liu LJ, Chen R, et al. Novel identified associations of RGS1 and RASGRP1 variants in IgA Nephropathy. Sci Rep. 2016;6:35781. doi:10.1038/srep35781

302. Yu CC, Yen TS, Lowell CA, DeFranco AL. Lupus-like kidney disease in mice deficient in the Src family tyrosine kinases Lyn and Fyn. Curr Biol. 2001;11(1):34–38. doi:10.1016/s0960-9822(00)00024-5

303. Andrade-Silva M, Cenedeze MA, Perandini LA, Felizardo RJF, Watanabe IKM, Agudelo JSH, Castoldi A, Gonçalves GM, Origassa CST, Semedo P, et al. TLR2 and TLR4 play opposite role in autophagy associated with cisplatin-induced acute kidney injury. Clin Sci (Lond). 2018;132(16):1725–1739. doi:10.1042/CS20170262

304. Maeda K, Otomo K, Yoshida N, Abu-Asab MS, Ichinose K, Nishino T, Kono M, Ferretti A, Bhargava R, Maruyama S, et al. CaMK4 compromises podocyte function in autoimmune and nonautoimmune kidney disease. J Clin Invest. 2018;128(8):3445–3459. doi:10.1172/JCI99507

305. Yang H, Zhou Q, Chen Dj, Jiang H, Mao Y, Chen J. The single nucleotide polymorphisms gene but not the copy number variation of Fcgr3B is associated with lupus nephritis in Chinese people. Lupus. 2010;19(5):662–664. doi:10.1177/0961203309350754

306. Higurashi M, Ohya Y, Joh K, Muraguchi M, Nishimura M, Terawaki H, Yagui K, Hashimoto N, Saito Y, Yamada K. Increased urinary levels of CXCL5, CXCL8 and CXCL9 in patients with Type 2 diabetic nephropathy. J Diabetes Complications. 2009;23(3):178–184. doi:10.1016/j.jdiacomp.2007.12.001

307. Nakatsuji T. High levels of serum soluble CD27 correlated with renal dysfunction. Clin Exp Med. 2003;2(4):192–196. doi:10.1007/s102380300006

308. Kato H, Watanabe H, Imafuku T, Arimura N, Fujita I, Noguchi I, Tanaka S, Nakano T, Tokumaru K, Enoki Y, et al. Advanced oxidation protein products contribute to chronic kidney disease-induced muscle atrophy by inducing oxidative stress via CD36/NADPH oxidase pathway. J Cachexia Sarcopenia Muscle. 2021;12(6):1832–1847. doi:10.1002/jcsm.12786

309. Awdishu L, Tsunoda S, Pearlman M, Kokoy-Mondragon C, Ghassemian M, Naviaux RK, Patton HM, Mehta RL, Vijay B, RamachandraRao SP.Identification of Maltase Glucoamylase as a Biomarker of Acute Kidney Injury in Patients with Cirrhosis. Crit Care Res Pract. 2019;2019:5912804. doi:10.1155/2019/5912804

310. Hains DS, Chen X, Saxena V, Barr-Beare E, Flemming W, Easterling R, Becknell B, Schwartz GJ, Schwaderer AL. Carbonic anhydrase 2 deficiency leads to increased pyelonephritis susceptibility. Am J Physiol Renal Physiol. 2014;307(7):F869–F880. doi:10.1152/ajprenal.00344.2014

311. McKnight AJ, Savage DA, Patterson CC, Sadlier D, Maxwell AP. Resequencing of genes for transforming growth factor beta1 (TGFB1) type 1 and 2 receptors (TGFBR1, TGFBR2), and association analysis of variants with diabetic nephropathy. BMC Med Genet. 2007;8:5. doi:10.1186/1471-2350-8-5

312. Costa NA, Polegato BF, Pereira AG, Paiva SAR, Gut AL, Balbi AL, Ponce D, Zornoff LAM, Azevedo PS, Minicucci MF. Evaluation of peptidylarginine deiminase 4 and PADI4 polymorphisms in sepsis-induced acute kidney injury. Rev Assoc Med Bras (1992). 2020;66(11):1515–1520. doi:10.1590/1806-9282.66.11.1515

313. Jones BM, Cheng IK, Wong RW, Kung AW. CD5-positive and CD5-negative rheumatoid factor-secreting B cells in IgA nephropathy, rheumatoid arthritis and Graves’ disease. Scand J Immunol. 1993;38(6):575–580. doi:10.1111/j.1365-3083.1993.tb03244.x

314. Sopić M, Joksić J, Spasojević-Kalimanovska V, Bogavac-Stanojević N, Simić-Ogrizović S, Kravljača M, Jelić Ivanović Z. Downregulation of AdipoR1 is Associated with increased Circulating Adiponectin Levels in Serbian Chronic Kidney Disease Patients. J Med Biochem. 2016;35(4):436–442. doi:10.1515/jomb-2016-0007

315. Rudnicki M, Perco P, Neuwirt H, Noppert SJ, Leierer J, Sunzenauer J, Eder S, Zoja C, Eller K, Rosenkranz AR, et al.Increased renal versican expression is associated with progression of chronic kidney disease. PLoS One. 2012;7(9):e44891. doi:10.1371/journal.pone.0044891

316. Meglic A, Debeljak M, Kovac J, Trampus Bakija A, Rajic V, Kojc N, Trebusak Podkrajsek K. SPTB related spherocytosis in a three-generation family presenting with kidney failure in adulthood due to co-occurrence of UMOD disease causing variant. Nefrologia (Engl Ed). 2020;40(4):421–428. doi:10.1016/j.nefro.2019.10.009

317. Bhatnagar V, Richard EL, Wu W, Nievergelt CM, Lipkowitz MS, Jeff J, Maihofer AX, Nigam SK. Analysis of ABCG2 and other urate transporters in uric acid homeostasis in chronic kidney disease: potential role of remote sensing and signaling. Clin Kidney J. 2016;9(3):444–453. doi:10.1093/ckj/sfw010

318. Wang Y, Zhu J, Liu Z, Shu S, Fu Y, Liu Y, Cai J, Tang C, Liu Y, Yin X, et al. The PINK1/PARK2/optineurin pathway of mitophagy is activated for protection in septic acute kidney injury. Redox Biol. 2021;38:101767. doi:10.1016/j.redox.2020.101767

319. Cao Y, Chen X, Sun H. Silencing of O-linked N-acetylglucosamine transferase ameliorates hypercalcemia-induced neurotoxicity in renal failure by regulating EZH2/KLF2/CXCL1 axis. Cell Death Dis. 2021;12(9):819. doi:10.1038/s41419-021-04022-x

320. Xu Y, Jiang W, Zhong L, Li H, Bai L, Chen X, Lin Y, Zheng D.circ-AKT3 aggravates renal ischaemia-reperfusion injury via regulating miR-144-5p /Wnt/β-catenin pathway and oxidative stress. J Cell Mol Med. 2020;10.1111/jcmm.16072. doi:10.1111/jcmm.16072

321. Bolin K, Sandling JK, Zickert A, Jönsen A, Sjöwall C, Svenungsson E, Bengtsson AA, Eloranta ML, Rönnblom L, Syvänen AC, et al. Association of STAT4 polymorphism with severe renal insufficiency in lupus nephritis. PLoS One. 2013;8(12):e84450. doi:10.1371/journal.pone.0084450

322. Kralisch S, Hoffmann A, Klöting N, Bachmann A, Kratzsch J, Stolzenburg JU, Dietel A, Beige J, Anders M, Bast I, et al. FSTL3 is increased in renal dysfunction. Nephrol Dial Transplant. 2017;32(10):1637–1644. doi:10.1093/ndt/gfw472

323. Huang H, Zhou H, Wang W, Dai X, Li W, Chen J, Bai Z, Pan J, Li X, Wang J, et al. Prediction of acute kidney injury, sepsis and mortality in children with urinary CXCL10. Pediatr Res. 2021;10.1038/s41390-021-01813-y. doi:10.1038/s41390-021-01813-y

324. Guo Y, Pace J, Li Z, Ma’ayan A, Wang Z, Revelo MP, Chen E, Gu X, Attalah A, Yang Y, et al. Podocyte-Specific Induction of Krüppel-Like Factor 15 Restores Differentiation Markers and Attenuates Kidney Injury in Proteinuric Kidney Disease. J Am Soc Nephrol. 2018;29(10):2529–2545. doi:10.1681/ASN.2018030324

325. Tsai IT, Wu CC, Hung WC, Lee TL, Hsuan CF, Wei CT, Lu YC, Yu TH, Chung FM, Lee YJ, et al. FABP1 and FABP2 as markers of diabetic nephropathy. Int J Med Sci. 2020;17(15):2338–2345. doi:10.7150/ijms.49078

326. Han CS, Liu K, Zhang N, Li SW, Gao HC. Rutin suppresses high glucose-induced ACTA2 and p38 protein expression in diabetic nephropathy. Exp Ther Med. 2017;14(1):181–186. doi:10.3892/etm.2017.4509

327. Lee BT, Ahmed FA, Hamm LL, Teran FJ, Chen CS, Liu Y, Shah K, Rifai N, Batuman V, Simon EE, et al. Association of C-reactive protein, tumor necrosis factor-alpha, and interleukin-6 with chronic kidney disease. BMC Nephrol. 2015;16:77. doi:10.1186/s12882-015-0068-7

328. Yang X, Bai M, Ning X, Ma F, Liu L, Liu T, Liu M, Wang H, Sun S. The associations of Bmi-1 with progression of glomerular chronic kidney diseaseL. Clin Nephrol. 2018;89(2):93–103. doi:10.5414/CN109117

329. Qin S, Taglienti M, Cai L, Zhou J, Kreidberg JA. c-Met and NF-κB-dependent overexpression of Wnt7a and −7b and Pax2 promotes cystogenesis in polycystic kidney disease. J Am Soc Nephrol. 2012;23(8):1309–1318. doi:10.1681/ASN.2011030277

330. Riedel JH, Paust HJ, Krohn S, Turner JE, Kluger MA, Steinmetz OM, Krebs CF, Stahl RA, Panzer U. IL-17F Promotes Tissue Injury in Autoimmune Kidney Diseases. J Am Soc Nephrol. 2016;27(12):3666–3677. doi:10.1681/ASN.2015101077

331. Zhang JY, Wang M, Tian L, Genovese G, Yan P, Wilson JG, Thadhani R, Mottl AK, Appel GB, Bick AG, et al. UBD modifies APOL1-induced kidney disease risk. Proc Natl Acad Sci U S A. 2018;115(13):3446–3451. doi:10.1073/pnas.1716113115

332. Shimizu Y, Yamanashi H, Noguchi Y, Koyamatsu J, Nagayoshi M, Kiyoura K, Fukui S, Tamai M, Kawashiri SY, Kondo H, et al. Association between chronic kidney disease and carotid intima-media thickness in relation to circulating CD34-positive cell count among community-dwelling elderly Japanese men. Atherosclerosis. 2019;283:85–91. doi:10.1016/j.atherosclerosis.2019.02.004

333. Poveda J, Sanz AB, Carrasco S, Ruiz-Ortega M, Cannata-Ortiz P, Sanchez-Niño MD, Ortiz A. Bcl3: a regulator of NF-κB inducible by TWEAK in acute kidney injury with anti-inflammatory and antiapoptotic properties in tubular cells. Exp Mol Med. 2017;49(7):e352. doi:10.1038/emm.2017.89

334. Vilander LM, Kaunisto MA, Vaara ST, Pettilä V; FINNAKI study group. Genetic variants in SERPINA4 and SERPINA5, but not BCL2 and SIK3 are associated with acute kidney injury in critically ill patients with septic shock. Crit Care. 2017;21(1):47. doi:10.1186/s13054-017-1631-3

335. Lu S, Liu S, Wietelmann A, Kojonazarov B, Atzberger A, Tang C, Schermuly RT, Gröne HJ, Offermanns S. Developmental vascular remodeling defects and postnatal kidney failure in mice lacking Gpr116 (Adgrf5) and Eltd1 (Adgrl4). PLoS One. 2017;12(8):e0183166. doi:10.1371/journal.pone.0183166

336. Feng Y, Lv LL, Wu WJ, Li ZL, Chen J, Ni HF, Zhou LT, Tang TT, Wang FM, Wang B, et al. Urinary Exosomes and Exosomal CCL2 mRNA as Biomarkers of Active Histologic Injury in IgA Nephropathy. Am J Pathol. 2018;188(11):2542–2552. doi:10.1016/j.ajpath.2018.07.017

337. Wang X, Ye H, Ward CJ, Chu JY, Masyuk TV, Larusso NF, Harris PC, Chow BK, Torres VE. Insignificant effect of secretin in rodent models of polycystic kidney and liver disease. Am J Physiol Renal Physiol. 2012;303(7):F1089–F1098. doi:10.1152/ajprenal.00242.2012

338. Li S, Jia Y, Xue M, Hu F, Zheng Z, Zhang S, Ren S, Yang Y, Si Z, Wang L, et al. Inhibiting Rab27a in renal tubular epithelial cells attenuates the inflammation of diabetic kidney disease through the miR-26a-5p/CHAC1/NF-kB pathway. Life Sci. 2020;261:118347. doi:10.1016/j.lfs.2020.118347

339. Liu Z, Liu H, Xiao L, Liu G, Sun L, He L. STC-1 ameliorates renal injury in diabetic nephropathy by inhibiting the expression of BNIP3 through the AMPK/SIRT3 pathway. Lab Invest. 2019;99(5):684–697. doi:10.1038/s41374-018-0176-7

340. An JN, Yang SH, Kim YC, Hwang JH, Park JY, Kim DK, Kim JH, Kim DW, Hur DG, Oh YK, et al. Periostin induces kidney fibrosis after acute kidney injury via the p38 MAPK pathway. Am J Physiol Renal Physiol. 2019;316(3):F426–F437. doi:10.1152/ajprenal.00203.2018

341. Kelly KJ, Zhang J, Han L, Wang M, Zhang S, Dominguez JH. Intravenous renal cell transplantation with SAA1-positive cells prevents the progression of chronic renal failure in rats with ischemic-diabetic nephropathy. Am J Physiol Renal Physiol. 2013;305(12):F1804–F1812. doi:10.1152/ajprenal.00097.2013

342. Nicolas A, Fatima S, Lamri A, Bellili-Muñoz N, Halimi JM, Saulnier PJ, Hadjadj S, Velho G, Marre M, Roussel R, et al. ABCG8 polymorphisms and renal disease in type 2 diabetic patients. Metabolism. 2015;64(6):713–719. doi:10.1016/j.metabol.2015.03.005

343. Chan AJ, Alikhan MA, Odobasic D, Gan PY, Khouri MB, Steinmetz OM, Mansell AS, Kitching AR, Holdsworth SR, Summers SA. Innate IL-17A-producing leukocytes promote acute kidney injury via inflammasome and Toll-like receptor activation. Am J Pathol. 2014;184(5):1411–1418. doi:10.1016/j.ajpath.2014.01.023

344. Tang S, Wang Y, Xie G, Li W, Chen Y, Liang J, Liu P, Song F, Zhou J. Regulation of Ptch1 by miR-342-5p and FoxO3 Induced Autophagy Involved in Renal Fibrosis. Front Bioeng Biotechnol. 2020;8:583318. doi:10.3389/fbioe.2020.583318

345. Xin C, Lei J, Wang Q, Yin Y, Yang X, Moran Guerrero JA, Sabbisetti V, Sun X, Vaidya VS, Bonventre JV. Therapeutic silencing of SMOC2 prevents kidney function loss in mouse model of chronic kidney disease. iScience. 2021;24(10):103193. doi:10.1016/j.isci.2021.103193

346. Axelsson J, Wang X, Ketteler M, Qureshi AR, Heimbürger O, Bárány P, Lindholm B, Nordfors L, Stenvinkel P. Is fetuin-A/alpha2-Heremans-Schmid glycoprotein associated with the metabolic syndrome in patients with chronic kidney disease?. Am J Nephrol. 2008;28(4):669–676. doi:10.1159/000121358

347. Menke J, Zeller GC, Kikawada E, Means TK, Huang XR, Lan HY, Lu B, Farber J, Luster AD, Kelley VR. CXCL9, but not CXCL10, promotes CXCR3-dependent immune-mediated kidney disease. J Am Soc Nephrol. 2008;19(6):1177–1189. doi:10.1681/ASN.2007111179

348. Ruiz-Hernández A, Cabrera-Becerra S, Vera-Juárez G, Hong E, Fengyang H, Arauz J, Villafaña S. Diabetic nephropathy produces alterations in the tissue expression profile of the orphan receptors GPR149, GPR153, GPR176, TAAR3, TAAR5 and TAAR9 in Wistar rats. Nucleosides Nucleotides Nucleic Acids. 2020;39(8):1150–1161. doi:10.1080/15257770.2020.1780437

349. Lee HJ, Donati A, Feliers D, Sun Y, Ding Y, Madesh M, Salmon AB, Ikeno Y, Ross C, O’Connor CL, Ju W, et al. Chloride channel accessory 1 integrates chloride channel activity and mTORC1 in aging-related kidney injury. Aging Cell. 2021;20(7):e13407. doi:10.1111/acel.13407

350. Zhao J, Zhu M, Jiang H, Shen S, Su X, Shi Y. Combination of sphingosine-1-phosphate receptor 1 (S1PR1) agonist and antiviral drug: a potential therapy against pathogenic influenza virus. Sci Rep. 2019;9(1):5272. doi:10.1038/s41598-019-41760-7

351. Kurtz SL, De Pascalis R, Meierovics AI, Elkins KL. Deficiency in CCR2 increases susceptibility of mice to infection with an intracellular pathogen, Francisella tularensis LVS, but does not impair development of protective immunity. PLoS One. 2021;16(3):e0249142. doi:10.1371/journal.pone.0249142

352. Raulf MK, Johannssen T, Matthiesen S, Neumann K, Hachenberg S, Mayer-Lambertz S, Steinbeis F, Hegermann J, Seeberger PH, Baumgärtner W, et al. The C-type Lectin Receptor CLEC12A Recognizes Plasmodial Hemozoin and Contributes to Cerebral Malaria Development. Cell Rep. 2019;28(1):30–38.e5. doi:10.1016/j.celrep.2019.06.015

353. Lindenstrøm T, Knudsen NP, Agger EM, Andersen P. Control of chronic mycobacterium tuberculosis infection by CD4 KLRG1-IL-2-secreting central memory cells. J Immunol. 2013;190(12):6311–6319. doi:10.4049/jimmunol.1300248

354. Allen IC, McElvania-TeKippe E, Wilson JE, Lich JD, Arthur JC, Sullivan JT, Braunstein M, Ting JP. Characterization of NLRP12 during the in vivo host immune response to Klebsiella pneumoniae and Mycobacterium tuberculosis. PLoS One. 2013;8(4):e60842. doi:10.1371/journal.pone.0060842

355. Berger SS, Turner L, Wang CW, Petersen JE, Kraft M, Lusingu JP, Mmbando B, Marquard AM, Bengtsson DB, Hviid L, et al. Plasmodium falciparum expressing domain cassette 5 type PfEMP1 (DC5-PfEMP1) bind PECAM1. PLoS One. 2013;8(7):e69117. doi:10.1371/journal.pone.0069117

356. Navarro-Sanchez E, Altmeyer R, Amara A, Schwartz O, Fieschi F, Virelizier JL, Arenzana-Seisdedos F, Desprès P. Dendritic-cell-specific ICAM3-grabbing non-integrin is essential for the productive infection of human dendritic cells by mosquito-cell-derived dengue viruses. EMBO Rep. 2003;4(7):723–728. doi:10.1038/sj.embor.embor866

357. Zhou F, Wan Q, Lu J, Chen Y, Lu G, He ML. Pim1 Impacts Enterovirus A71 Replication and Represents a Potential Target in Antiviral Therapy. iScience. 2019;19:715–727. doi:10.1016/j.isci.2019.08.008

358. Shi N, Zhang S, Guo Y, Yu X, Zhao W, Zhang M, Guan Z, Duan M. CircRNA_0050463 promotes influenza A virus replication by sponging miR-33b-5p to regulate EEF1A1. Vet Microbiol. 2021;254:108995. doi:10.1016/j.vetmic.2021.108995

359. McCormack RM, Szymanski EP, Hsu AP, Perez E, Olivier KN, Fisher E, Goodhew EB, Podack ER, Holland SM. MPEG1/perforin-2 mutations in human pulmonary nontuberculous mycobacterial infections. JCI Insight. 2017;2(8):e89635. doi:10.1172/jci.insight.89635

360. Bozza S, Iannitti RG, Pariano M, Renga G, Costantini C, Romani L, Bayry J. Small Molecule CCR4 Antagonists Protect Mice from Aspergillus Infection and Allergy. Biomolecules. 2021;11(3):351. doi:10.3390/biom11030351

361. Zhivaki D, Lemoine S, Lim A, Morva A, Vidalain PO, Schandene L, Casartelli N, Rameix-Welti MA, Hervé PL, Dériaud E, et al. Respiratory Syncytial Virus Infects Regulatory B Cells in Human Neonates via Chemokine Receptor CX3CR1 and Promotes Lung Disease Severity. Immunity. 2017;46(2):301–314. doi:10.1016/j.immuni.2017.01.010

362. Cui N, Su LX, Wang H, Xiao M, Yang F, Zheng M, Li X, Xu YC, Liu DW. mTOR Modulates Lymphocyte Differentiation through T-bet and Eomesodermin in Response to Invasive Pulmonary Aspergillosis in Rats. Chin Med J (Engl). 2016;129(14):1704–1710. doi:10.4103/0366-6999.185858

363. Bel S, Pendse M, Wang Y, Li Y, Ruhn KA, Hassell B, Leal T, Winter SE, Xavier RJ, Hooper LV. Paneth cells secrete lysozyme via secretory autophagy during bacterial infection of the intestine. Science. 2017;357(6355):1047–1052. doi:10.1126/science.aal4677

364. Wang Y, Zhong H, Xie X, Chen CY, Huang D, Shen L, Zhang H, Chen ZW, Zeng G. Long noncoding RNA derived from CD244 signaling epigenetically controls CD8+ T-cell immune responses in tuberculosis infection. Proc Natl Acad Sci U S A. 2015;112(29):E3883–E3892. doi:10.1073/pnas.1501662112

365. Hernández-Castañeda MA, Lavergne M, Casanova P, Nydegger B, Merten C, Subramanian BY, Matthey P, Lannes N, Mantel PY, Walch M. Profound Membrane Reorganization Defines Susceptibility of Plasmodium falciparum Infected Red Blood Cells to Lysis by Granulysin and Perforin. Front Immunol. 2021;12:643746. doi:10.3389/fimmu.2021.643746

366. Wu Y, Wu M, Ming S, Zhan X, Hu S, Li X, Yin H, Cao C, Liu J, Li J, et al. TREM-2 promotes Th1 responses by interacting with the CD3ζ-ZAP70 complex following Mycobacterium tuberculosis infection. J Clin Invest. 2021;131(17):e137407. doi:10.1172/JCI137407

367. Park M, Baker W, Cambow D, Gogerty D, Leda AR, Herlihy B, Pavlenko D, Van Den Nieuwenhuizen S, Toborek M. Methamphetamine Enhances HIV-Induced Aberrant Proliferation of Neural Progenitor Cells via the FOXO3-Mediated Mechanism. Mol Neurobiol. 2021;58(11):5421–5436. doi:10.1007/s12035-021-02407-9

368. Ndlovu H, Darby M, Froelich M, Horsnell W, Lühder F, Hünig T, Brombacher F. Inducible deletion of CD28 prior to secondary nippostrongylus brasiliensis infection impairs worm expulsion and recall of protective memory CD4L T cell responses. PLoS Pathog. 2014;10(2):e1003906. doi:10.1371/journal.ppat.1003906

369. Chen X, Ren F, Hesketh J, Shi X, Li J, Gan F, Huang K. Selenium blocks porcine circovirus type 2 replication promotion induced by oxidative stress by improving GPx1 expression. Free Radic Biol Med. 2012;53(3):395–405. doi:10.1016/j.freeradbiomed.2012.04.035

370. D’Onofrio V, Monnier AA, Kremer C, Stappers MHT, Netea MG, Gyssens IC. Lesion size is associated with genetic polymorphisms in TLR1, TLR6, and TIRAP genes in patients with major abscesses and diabetic foot infections. Eur J Clin Microbiol Infect Dis. 2020;39(2):353–360. doi:10.1007/s10096-019-03732-7

371. Wang LL, Chen XF, Hu P, Lu SY, Fu BQ, Li YS, Zhai FF, Ju DD, Zhang SJ, Shui YM, et al. Host Prdx6 contributing to the intracellular survival of Brucella suis S2 strain. BMC Vet Res. 2019;15(1):304. doi:10.1186/s12917-019-2049-8

372. Muenstermann M, Strobel L, Klos A, Wetsel RA, Woodruff TM, Köhl J, Johswich KO. Distinct roles of the anaphylatoxin receptors C3aR, C5aR1 and C5aR2 in experimental meningococcal infections. Virulence. 2019;10(1):677–694. doi:10.1080/21505594.2019.1640035

373. Weindel CG, Bell SL, Vail KJ, West KO, Patrick KL, Watson RO. LRRK2 maintains mitochondrial homeostasis and regulates innate immune responses to Mycobacterium tuberculosis. Elife. 2020;9:e51071. doi:10.7554/eLife.51071

374. Vesosky B, Rottinghaus EK, Stromberg P, Turner J, Beamer G. CCL5 participates in early protection against Mycobacterium tuberculosis. J Leukoc Biol. 2010;87(6):1153–1165. doi:10.1189/jlb.1109742

375. Teng Y, Cang B, Mao F, Chen W, Cheng P, Peng L, Luo P, Lu D, You N, Zou Q, et al. Expression of ETS1 in gastric epithelial cells positively regulate inflammatory response in Helicobacter pylori-associated gastritis. Cell Death Dis. 2020;11(7):498. doi:10.1038/s41419-020-2705-8

376. Sumegi J, Huang D, Lanyi A, Davis JD, Seemayer TA, Maeda A, Klein G, Seri M, Wakiguchi H, Purtilo DT, et al. Correlation of mutations of the SH2D1A gene and epstein-barr virus infection with clinical phenotype and outcome in X-linked lymphoproliferative disease. Blood. 2000;96(9):3118–3125.

377. Turner JD, Pionnier N, Furlong-Silva J, Sjoberg H, Cross S, Halliday A, Guimaraes AF, Cook DAN, Steven A, Van Rooijen N, et al. Interleukin-4 activated macrophages mediate immunity to filarial helminth infection by sustaining CCR3-dependent eosinophilia. PLoS Pathog. 2018;14(3):e1006949. doi:10.1371/journal.ppat.1006949

378. Schanoski AS, Le TT, Kaiserman D, Rowe C, Prow NA, Barboza DD, Santos CA, Zanotto PMA, Magalhães KG, Aurelio L, et al. Granzyme A in Chikungunya and Other Arboviral Infections. Front Immunol. 2020;10:3083. doi:10.3389/fimmu.2019.03083

379. Muñoz S, Hernández-Pando R, Abraham SN, Enciso JA. Mast cell activation by Mycobacterium tuberculosis: mediator release and role of CD48. J Immunol. 2003;170(11):5590–5596. doi:10.4049/jimmunol.170.11.5590

380. Wang Y, Koroleva EP, Kruglov AA, Kuprash DV, Nedospasov SA, Fu YX, Tumanov AV. Lymphotoxin beta receptor signaling in intestinal epithelial cells orchestrates innate immune responses against mucosal bacterial infection. Immunity. 2010;32(3):403–413. doi:10.1016/j.immuni.2010.02.011

381. Torre S, Faucher SP, Fodil N, Bongfen SE, Berghout J, Schwartzentruber JA, Majewski J, Lathrop M, Cooper AM, Vidal SM, et al. THEMIS is required for pathogenesis of cerebral malaria and protection against pulmonary tuberculosis. Infect Immun. 2015;83(2):759–768. doi:10.1128/IAI.02586-14

382. Wu J, Gu J, Shen L, Fang D, Zou X, Cao Y, Wang S, Mao L. Exosomal MicroRNA-155 Inhibits Enterovirus A71 Infection by Targeting PICALM. Int J Biol Sci. 2019;15(13):2925–2935. doi:10.7150/ijbs.36388

383. Bhalla M, Hui Yeoh J, Lamneck C, Herring SE, Tchalla EYI, Heinzinger LR, Leong JM, Bou Ghanem EN. A1 adenosine receptor signaling reduces Streptococcus pneumoniae adherence to pulmonary epithelial cells by targeting expression of platelet-activating factor receptor. Cell Microbiol. 2020;22(2):e13141. doi:10.1111/cmi.13141

384. Muramatsu T. Basigin: a multifunctional membrane protein with an emerging role in infections by malaria parasites. Expert Opin Ther Targets. 2012;16(10):999–1011. doi:10.1517/14728222.2012.711818

385. Shen C, Wu XR, Wang BB, Sun L, Jiao WW, Wang J, Feng WX, Xiao J, Miao Q, Liu F, et al. ALOX5 is associated with tuberculosis in a subset of the pediatric population of North China. Genet Test Mol Biomarkers. 2013;17(4):284–288. doi:10.1089/gtmb.2012.0426

386. Schäfer G, Graham LM, Lang DM, Blumenthal MJ, Bergant Marušič M, Katz AA. Vimentin Modulates Infectious Internalization of Human Papillomavirus 16 Pseudovirions. J Virol. 2017;91(16):e00307–17. doi:10.1128/JVI.00307-17

387. Rao P, Pham HT, Kulkarni A, Yang Y, Liu X, Knipe DM, Cresswell P, Yuan W. Herpes simplex virus 1 glycoprotein B and US3 collaborate to inhibit CD1d antigen presentation and NKT cell function. J Virol. 2011;85(16):8093–8104. doi:10.1128/JVI.02689-10

388. Fu Z, Cai W, Shao J, Xue H, Ge Z, Fan H, Dong C, Wang C, Zhang J, Shen C, et al. Genetic Variants in TNFSF4 and TNFSF8 Are Associated With the Risk of HCV Infection Among Chinese High-Risk Population. Front Genet. 2021;12:630310. doi:10.3389/fgene.2021.630310

389. Tanigawa K, Suzuki K, Kimura H, Takeshita F, Wu H, Akama T, Kawashima A, Ishii N. Tryptophan aspartate-containing coat protein (CORO1A) suppresses Toll-like receptor signalling in Mycobacterium leprae infection. Clin Exp Immunol. 2009;156(3):495–501. doi:10.1111/j.1365-2249.2009.03930.x

390. Banerjee A, Tripathi A, Duggal S, Banerjee A, Vrati S. Dengue virus infection impedes megakaryopoiesis in MEG-01 cells where the virus envelope protein interacts with the transcription factor TAL-1. Sci Rep. 2020;10(1):19587. doi:10.1038/s41598-020-76350-5

391. Hobden JA. Intercellular adhesion molecule-2 (ICAM-2) and Pseudomonas aeruginosa ocular infection. DNA Cell Biol. 2003;22(10):649–655. doi:10.1089/104454903770238120

392. Nguyen CT, Kim EH, Luong TT, Pyo S, Rhee DK. TLR4 mediates pneumolysin-induced ATF3 expression through the JNK/p38 pathway in Streptococcus pneumoniae-infected RAW 264.7 cells. Mol Cells. 2015;38(1):58–64. doi:10.14348/molcells.2015.2231

393. Bhalla M, Van Ngo H, Gyanwali GC, Ireton K. The Host Scaffolding Protein Filamin A and the Exocyst Complex Control Exocytosis during InlB-Mediated Entry of Listeria monocytogenes. Infect Immun. 2018;87(1):e00689–18. doi:10.1128/IAI.00689-18

394. Heer AK, Harris NL, Kopf M, Marsland BJ. CD4+ and CD8+ T cells exhibit differential requirements for CCR7-mediated antigen transport during influenza infection. J Immunol. 2008;181(10):6984–6994. doi:10.4049/jimmunol.181.10.6984

395. Peck KM, Cockrell AS, Yount BL, Scobey T, Baric RS, Heise MT. Glycosylation of mouse DPP4 plays a role in inhibiting Middle East respiratory syndrome coronavirus infection. J Virol. 2015;89(8):4696–4699. doi:10.1128/JVI.03445-14

396. Verma V, Gupta S, Kumar P, Yadav S, Dhanda RS, Gaind R, Arora R, Frimodt-Møller N, Yadav M. Involvement of NLRP3 and NLRC4 Inflammasome in Uropathogenic E. coli Mediated Urinary Tract Infections. Front Microbiol. 2019;10:2020. doi:10.3389/fmicb.2019.02020

397. Aguilar-Briseño JA, Upasani V, Ellen BMT, Moser J, Pauzuolis M, Ruiz-Silva M, Heng S, Laurent D, Choeung R, Dussart P, et al. TLR2 on blood monocytes senses dengue virus infection and its expression correlates with disease pathogenesis. Nat Commun. 2020;11(1):3177. doi:10.1038/s41467-020-16849-7

398. Funk KE, Klein RS. CSF1R antagonism limits local restimulation of antiviral CD8+ T cells during viral encephalitis. J Neuroinflammation. 2019;16(1):22. doi:10.1186/s12974-019-1397-4

399. Chen Y, Zhou J, Cheng Z, Yang S, Chu H, Fan Y, Li C, Wong BH, Zheng S, Zhu Y, et al. Functional variants regulating LGALS1 (Galectin 1) expression affect human susceptibility to influenza A(H7N9). Sci Rep. 2015;5:8517. doi:10.1038/srep08517

400. Mancuso RI, Miyaji EN, Silva CCF, Portaro FV, Soares-Schanoski A, Ribeiro OG, Oliveira MLS. Impaired expression of CXCL5 and matrix metalloproteinases in the lungs of mice with high susceptibility to Streptococcus pneumoniae infection. Immun Inflamm Dis. 2018;6(1):128–142. doi:10.1002/iid3.205

401. Welten SP, Redeker A, Franken KL, Benedict CA, Yagita H, Wensveen FM, Borst J, Melief CJ, van Lier RA, van Gisbergen KP, et al. CD27-CD70 costimulation controls T cell immunity during acute and persistent cytomegalovirus infection. J Virol. 2013;87(12):6851–6865. doi:10.1128/JVI.03305-12

402. Okuda K, Tong M, Dempsey B, Moore KJ, Gazzinelli RT, Silverman N. Leishmania amazonensis Engages CD36 to Drive Parasitophorous Vacuole Maturation. PLoS Pathog. 2016;12(6):e1005669. doi:10.1371/journal.ppat.1005669

403. Rahman M, Haberman A, Tracy C, Ray S, Krämer H. Drosophila mauve mutants reveal a role of LYST homologs late in the maturation of phagosomes and autophagosomes. Traffic. 2012;13(12):1680–1692. doi:10.1111/tra.12005

404. Zhu B, Ye J, Nie Y, Ashraf U, Zohaib A, Duan X, Fu ZF, Song Y, Chen H, Cao S. MicroRNA-15b Modulates Japanese Encephalitis Virus-Mediated Inflammation via Targeting RNF125. J Immunol. 2015;195(5):2251–2262. doi:10.4049/jimmunol.1500370

405. Bartlett S, Gemiarto AT, Ngo MD, Sajiir H, Hailu S, Sinha R, Foo CX, Kleynhans L, Tshivhula H, Webber T, et al. GPR183 Regulates Interferons, Autophagy, and Bacterial Growth During Mycobacterium tuberculosis Infection and Is Associated With TB Disease Severity. Front Immunol. 2020;11:601534. doi:10.3389/fimmu.2020.601534

406. Alagarasu K, Bachal RV, Damle I, Shah PS, Cecilia D. Association of FCGR2A p.R131H and CCL2 c.-2518 A>G gene variants with thrombocytopenia in patients with dengue virus infection. Hum Immunol. 2015;76(11):819–822. doi:10.1016/j.humimm.2015.09.042

407. Mourglia-Ettlin G, Miles S, Velasco-De-Andrés M, Armiger-Borràs N, Cucher M, Dematteis S, Lozano F. The ectodomains of the lymphocyte scavenger receptors CD5 and CD6 interact with tegumental antigens from Echinococcus granulosus sensu lato and protect mice against secondary cystic echinococcosis. PLoS Negl Trop Dis. 2018;12(11):e0006891. doi:10.1371/journal.pntd.0006891

408. Almutairi F, Sarr D, Tucker SL, Fantone K, Lee JK, Rada B. RGS10 Reduces Lethal Influenza Infection and Associated Lung Inflammation in Mice. Front Immunol. 2021;12:772288. doi:10.3389/fimmu.2021.772288

409. Kellar GG, Barrow KA, Rich LM, Debley JS, Wight TN, Ziegler SF, Reeves SR. Loss of versican and production of hyaluronan in lung epithelial cells are associated with airway inflammation during RSV infection. J Biol Chem. 2021;296:100076. doi:10.1074/jbc.RA120.016196

410. Komissarov A, Sergeeva M, Zhuravlev E, Medvedev S, Malakhova A, Andreeva E, Shurygina AP, Gorshkov A, Timofeeva M, Balakhonova E, et al. CRISPR-Cas9 mediated knockout of AnxA6 gene enhances influenza A virus replication in low-permissive HEK293FT cell line. Gene. 2022;809:146024. doi:10.1016/j.gene.2021.146024

411. Li H, Zhang C, Cui H, Guo K, Wang F, Zhao T, Liang W, Lv Q, Zhang Y. FKBP8 interact with classical swine fever virus NS5A protein and promote virus RNA replication. Virus Genes. 2016;52(1):99–106. doi:10.1007/s11262-015-1286-6

412. Schaaf K, Smith SR, Duverger A, Wagner F, Wolschendorf F, Westfall AO, Kutsch O, Sun J.Mycobacterium tuberculosis exploits the PPM1A signaling pathway to block host macrophage apoptosis. Sci Rep. 2017;7:42101. doi:10.1038/srep42101

413. Liu B, Gao TT, Fu XY, Xu ZH, Ren H, Zhao P, Qi ZT, Qin ZL. PTEN Lipid Phosphatase Activity Enhances Dengue Virus Production through Akt/FoxO1/Maf1 Signaling. Virol Sin. 2021;36(3):412–423. doi:10.1007/s12250-020-00291-6

414. Bartlett S, Gemiarto AT, Ngo MD, Sajiir H, Hailu S, Sinha R, Foo CX, Kleynhans L, Tshivhula H, Webber T, et al. GPR183 Regulates Interferons, Autophagy, and Bacterial Growth During Mycobacterium tuberculosis Infection and Is Associated With TB Disease Severity. Front Immunol. 2020;11:601534. doi:10.3389/fimmu.2020.601534

415. Angelova M, Ortiz-Meoz RF, Walker S, Knipe DM. Inhibition of O-Linked N-Acetylglucosamine Transferase Reduces Replication of Herpes Simplex Virus and Human Cytomegalovirus. J Virol. 2015;89(16):8474–8483. doi:10.1128/JVI.01002-15

416. Subrahmanian M, Marimuthu J, Sairam T, Sankaran R. In vitro ubiquitination of Mycobacterium tuberculosis by E3 ubiquitin ligase, MKRN1. Biotechnol Lett. 2020;42(8):1527–1534. doi:10.1007/s10529-020-02873-6

417. Eva MM, Yuki KE, Dauphinee SM, Schwartzentruber JA, Pyzik M, Paquet M, Lathrop M, Majewski J, Vidal SM, Malo D. Altered IFN-γ-mediated immunity and transcriptional expression patterns in N-Ethyl-N-nitrosourea-induced STAT4 mutants confer susceptibility to acute typhoid-like disease. J Immunol. 2014;192(1):259–270. doi:10.4049/jimmunol.1301370

418. Wang H, Liu D, Sun Y, Meng C, Tan L, Song C, Qiu X, Liu W, Ding C, Ying L. Upregulation of DUSP6 impairs infectious bronchitis virus replication by negatively regulating ERK pathway and promoting apoptosis. Vet Res. 2021;52(1):7. doi:10.1186/s13567-020-00866-x

419. Zhetkenev S, Khassan A, Khamzina A, Issanov A, Crape B, Akilzhanova A, Yerezhepov D, Kozhamkulov U, Chan CK. Association of rs12722 COL5A1 with pulmonary tuberculosis: a preliminary case-control study in a Kazakhstani population. Mol Biol Rep. 2021;48(1):691–699. doi:10.1007/s11033-020-06121-y

420. Semmler G, Griebler H, Aberle SW, Stiasny K, Richter L, Holzmann H, Weseslindtner L. Elevated CXCL10 Serum Levels in Measles Virus Primary Infection and Reinfection Correlate With the Serological Stage and Hospitalization Status. J Infect Dis. 2020;222(12):2030–2034. doi:10.1093/infdis/jiaa326

421. Hu Z, Zhang C, Sifuentes-Dominguez L, Zarek CM, Propheter DC, Kuang Z, Wang Y, Pendse M, Ruhn KA, Hassell B, et al. Small proline-rich protein 2A is a gut bactericidal protein deployed during helminth infection. Science. 2021;374(6568):eabe6723. doi:10.1126/science.abe6723

422. El-Mayet FS, Harrison KS, Jones C. Regulation of Krüppel-Like Factor 15 Expression by Herpes Simplex Virus Type 1 or Bovine Herpesvirus 1 Productive Infection. Viruses. 2021;13(6):1148. doi:10.3390/v13061148

423. Frakolaki E, Kalliampakou KI, Kaimou P, Moraiti M, Kolaitis N, Boleti H, Koskinas J, Vassilacopoulou D, Vassilaki N. Emerging Role of l-Dopa Decarboxylase in Flaviviridae Virus Infections. Cells. 2019;8(8):837. doi:10.3390/cells8080837

424. Hansson LO, Axelsson G, Linné T, Aurelius E, Lindquist L. Serum C-reactive protein in the differential diagnosis of acute meningitis. Scand J Infect Dis. 1993;25(5):625–630. doi:10.3109/00365549309008552

425. Grassl GA, Faustmann M, Gill N, Zbytnuik L, Merkens H, So L, Rossi FM, McNagny KM, Finlay BB. CD34 mediates intestinal inflammation in Salmonella-infected mice. Cell Microbiol. 2010;12(11):1562–1575. doi:10.1111/j.1462-5822.2010.01488.x

426. Tassi I, Claudio E, Wang H, Tang W, Ha HL, Saret S, Sher A, Jankovic D, Siebenlist U. Adaptive immune-mediated host resistance to Toxoplasma gondii is governed by the NF-κB regulator Bcl-3 in dendritic cells. Eur J Immunol. 2015;45(7):1972–1979. doi:10.1002/eji.201445045

427. Pyo KH, Kang EY, Jung BK, Moon JH, Chai JY, Shin EH. Depressed neuronal growth associated protein (GAP)-43 expression in the small intestines of mice experimentally infected with Neodiplostomum seoulense. Korean J Parasitol. 2012;50(1):89–93. doi:10.3347/kjp.2012.50.1.89

428. Stier MT, Bloodworth MH, Toki S, Newcomb DC, Goleniewska K, Boyd KL, Quitalig M, Hotard AL, Moore ML, Hartert TV, et al. Respiratory syncytial virus infection activates IL-13-producing group 2 innate lymphoid cells through thymic stromal lymphopoietin. J Allergy Clin Immunol. 2016;138(3):814–824.e11. doi:10.1016/j.jaci.2016.01.050

429. Alagarasu K, Bachal RV, Damle I, Shah PS, Cecilia D. Association of FCGR2A p.R131H and CCL2 c.-2518 A>G gene variants with thrombocytopenia in patients with dengue virus infection. Hum Immunol. 2015;76(11):819–822. doi:10.1016/j.humimm.2015.09.042

430. Hassane M, Jouan Y, Creusat F, Soulard D, Boisseau C, Gonzalez L, Patin EC, Heuzé-Vourc’h N, Sirard JC, Faveeuw C, et al. Interleukin-7 protects against bacterial respiratory infection by promoting IL-17A-producing innate T-cell response. Mucosal Immunol. 2020;13(1):128–139. doi:10.1038/s41385-019-0212-y

431. Zhu X, Wang Y, Zhang H, Liu X, Chen T, Yang R, Shi Y, Cao W, Li P, Ma Q, et al. Genetic variation of the human α-2-Heremans-Schmid glycoprotein (AHSG) gene associated with the risk of SARS-CoV infection. PLoS One. 2011;6(8):e23730. doi:10.1371/journal.pone.0023730

432. Ting HA, Schaller MA, de Almeida Nagata DE, Rasky AJ, Maillard IP, Lukacs NW. Notch Ligand Delta-like 4 Promotes Regulatory T Cell Identity in Pulmonary Viral Infection. J Immunol. 2017;198(4):1492–1502. doi:10.4049/jimmunol.1601654

433. Rydengård V, Shannon O, Lundqvist K, Kacprzyk L, Chalupka A, Olsson AK, Mörgelin M, Jahnen-Dechent W, Malmsten M, Schmidtchen A. Histidine-rich glycoprotein protects from systemic Candida infection. PLoS Pathog. 2008;4(8):e1000116. doi:10.1371/journal.ppat.1000116

434. Bultmann H, Busse JS, Brandt CR. Modified FGF4 signal peptide inhibits entry of herpes simplex virus type 1. J Virol. 2001;75(6):2634–2645. doi:10.1128/JVI.75.6.2634-2645.2001

435. Hasan Z, Jamil B, Khan J, Ali R, Khan MA, Nasir N, Yusuf MS, Jamil S, Irfan M, Hussain R. Relationship between circulating levels of IFN-gamma, IL-10, CXCL9 and CCL2 in pulmonary and extrapulmonary tuberculosis is dependent on disease severity. Scand J Immunol. 2009;69(3):259–267. doi:10.1111/j.1365-3083.2008.02217.x

436. Lin Q, Zhou S, Huang Y, Huo Z, Chen C, Luo X, He J, Liu C, Zhang P. ANKS4B Restricts Replication of Zika Virus by Downregulating the Autophagy. Front Microbiol. 2020;11:1745. doi:10.3389/fmicb.2020.01745

437. Rao CP, Minta JO, Laski B, Alper CA, Gelfand EW. Inherited C8 beta subunit deficiency in a patient with recurrent meningococcal infections: in vivo functional kinetic analysis of C8. Clin Exp Immunol. 1985;60(1):183–190.

438. Tao YT, Huang Q, Jiang YL, Wang XL, Sun P, Tian Y, Wu HL, Zhang M, Meng SB, Wang YS, et al. Up-regulation of Slc39A2(Zip2) mRNA in peripheral blood mononuclear cells from patients with pulmonary tuberculosis. Mol Biol Rep. 2013;40(8):4979–4984. doi:10.1007/s11033-013-2598-z

439. Lee EJ, Park KS, Jeon IS, Choi JW, Lee SJ, Choy HE, Song KD, Lee HK, Choi JK. LAMP-3 (Lysosome-Associated Membrane Protein 3) Promotes the Intracellular Proliferation of Salmonella typhimurium. Mol Cells. 2016;39(7):566–572. doi:10.14348/molcells.2016.0112

440. Deng S, Zhou X, Ge Z, Song Y, Wang H, Liu X, Zhang D. Exosomes from adipose-derived mesenchymal stem cells ameliorate cardiac damage after myocardial infarction by activating S1P/SK1/S1PR1 signaling and promoting macrophage M2 polarization. Int J Biochem Cell Biol. 2019;114:105564. doi:10.1016/j.biocel.2019.105564

441. Maneerat Y, Prasongsukarn K, Benjathummarak S, Dechkhajorn W. PPBP and DEFA1/DEFA3 genes in hyperlipidaemia as feasible synergistic inflammatory biomarkers for coronary heart disease. Lipids Health Dis. 2017;16(1):80. doi:10.1186/s12944-017-0471-0

442. França CN, Izar MCO, Hortêncio MNS, do Amaral JB, Ferreira CES, Tuleta ID, Fonseca FAH. Monocyte subtypes and the CCR2 chemokine receptor in cardiovascular disease. Clin Sci (Lond). 2017;131(12):1215–1224. doi:10.1042/CS20170009

443. Akosile W, Voisey J, Lawford B, Colquhounc D, Young RM, Mehta D. The inflammasome NLRP12 is associated with both depression and coronary artery disease in Vietnam veterans. Psychiatry Res. 2018;270:775–779. doi:10.1016/j.psychres.2018.10.051

444. Harry BL, Sanders JM, Feaver RE, Lansey M, Deem TL, Zarbock A, Bruce AC, Pryor AW, Gelfand BD, Blackman BR, et al. Endothelial cell PECAM-1 promotes atherosclerotic lesions in areas of disturbed flow in ApoE-deficient mice. Arterioscler Thromb Vasc Biol. 2008;28(11):2003–2008. doi:10.1161/ATVBAHA.108.164707

445. Yang H, Wang XX, Zhou CY, Xiao X, Tian C, Li HH, Yin CL, Wang HX. Tripartite motif 10 regulates cardiac hypertrophy by targeting the PTEN/AKT pathway. J Cell Mol Med. 2020;24(11):6233–6241. doi:10.1111/jcmm.15257

446. Maneerat Y, Prasongsukarn K, Benjathummarak S, Dechkhajorn W. PPBP and DEFA1/DEFA3 genes in hyperlipidaemia as feasible synergistic inflammatory biomarkers for coronary heart disease. Lipids Health Dis. 2017;16(1):80. doi:10.1186/s12944-017-0471-0

447. Mo XG, Liu W, Yang Y, Imani S, Lu S, Dan G, Nie X, Yan J, Zhan R, Li X, et al. NCF2, MYO1F, S1PR4, and FCN1 as potential noninvasive diagnostic biomarkers in patients with obstructive coronary artery: A weighted gene co-expression network analysis. J Cell Biochem. 2019;120(10):18219–18235. doi:10.1002/jcb.29128

448. Liu N, Wang BJ, Broughton KM, Alvarez R, Siddiqi S, Loaiza R, Nguyen N, Quijada P, Gude N, Sussman MA. PIM1-minicircle as a therapeutic treatment for myocardial infarction. PLoS One. 2017;12(3):e0173963. doi:10.1371/journal.pone.0173963

449. Hou YS, Wang JZ, Shi S, Han Y, Zhang Y, Zhi JX, Xu C, Li FF, Wang GY, Liu SL. Identification of epigenetic factor KAT2B gene variants for possible roles in congenital heart diseases. Biosci Rep. 2020;40(4):BSR20191779. doi:10.1042/BSR20191779

450. Elmén L, Volpato CB, Kervadec A, Pineda S, Kalvakuri S, Alayari NN, Foco L, Pramstaller PP, Ocorr K, Rossini A, et al. Silencing of CCR4-NOT complex subunits affects heart structure and function. Dis Model Mech. 2020;13(7):dmm044727. doi:10.1242/dmm.044727

451. Thude H, Gerlach K, Richartz B, Krack A, Brenke B, Pethig K, Figulla HR, Barz D. No association between transmembrane protein-tyrosine phosphatase receptor type C (CD45) exon A point mutation (77C>G) and idiopathic dilated cardiomyopathy. Hum Immunol. 2005;66(9):1008–1012. doi:10.1016/j.humimm.2005.07.004

452. Njerve IU, Pettersen AÅ, Opstad TB, Arnesen H, Seljeflot I. Fractalkine and its receptor (CX3CR1) in patients with stable coronary artery disease and diabetes mellitus. Metab Syndr Relat Disord. 2012;10(6):400–406. doi:10.1089/met.2012.0052

453. Su Y, Zhu C, Wang B, Zheng H, McAlister V, Lacefield JC, Quan D, Mele T, Greasley A, Liu K, et al. Circular RNA Foxo3 in cardiac ischemia-reperfusion injury in heart transplantation: A new regulator and target. Am J Transplant. 2021;21(9):2992–3004. doi:10.1111/ajt.16475

454. Zhang Q, Zheng Y, Ning M, Li T. KLRD1, FOSL2 and LILRB3 as potential biomarkers for plaques progression in acute myocardial infarction and stable coronary artery disease. BMC Cardiovasc Disord. 2021;21(1):344. doi:10.1186/s12872-021-01997-5

455. Wang H, Kwak D, Fassett J, Hou L, Xu X, Burbach BJ, Thenappan T, Xu Y, Ge JB, Shimizu Y, et al. CD28/B7 Deficiency Attenuates Systolic Overload-Induced Congestive Heart Failure, Myocardial and Pulmonary Inflammation, and Activated T Cell Accumulation in the Heart and Lungs. Hypertension. 2016;68(3):688–696. doi:10.1161/HYPERTENSIONAHA.116.07579

456. Yeh HL, Kuo LT, Sung FC, Yeh CC. Association between Polymorphisms of Antioxidant Gene (MnSOD, CAT, and GPx1) and Risk of Coronary Artery Disease. Biomed Res Int. 2018;2018:5086869. doi:10.1155/2018/5086869

457. Wang C, Li Q, Yang H, Gao C, Du Q, Zhang C, Zhu L, Li Q. MMP9, CXCR1, TLR6, and MPO participant in the progression of coronary artery disease. J Cell Physiol. 2020;235(11):8283–8292. doi:10.1002/jcp.29485

458. Mohtavinejad N, Nakhaee A, Harati H, Gholipour N, Mahmoodzade Y. Association of CCL5 rs2107538, and CCL2 rs3760396 Gene Polymorphisms with the Risk of Cardiovascular Disease. Iran J Public Health. 2021;50(7):1436–1444. doi:10.18502/ijph.v50i7.6634

459. Lee TL, Lai TC, Lin SR, Lin SW, Chen YC, Pu CM, Lee IT, Tsai JS, Lee CW, Chen YL.Conditioned medium from adipose-derived stem cells attenuates ischemia/reperfusion-induced cardiac injury through the microRNA-221/222/PUMA/ETS-1 pathway. Theranostics. 2021;11(7):3131–3149. doi:10.7150/thno.52677

460. Haley KJ, Lilly CM, Yang JH, Feng Y, Kennedy SP, Turi TG, Thompson JF, Sukhova GH, Libby P, Lee RT. Overexpression of eotaxin and the CCR3 receptor in human atherosclerosis: using genomic technology to identify a potential novel pathway of vascular inflammation. Circulation. 2000;102(18):2185–2189. doi:10.1161/01.cir.102.18.2185

461. Jia P, Wang J, Wang L, Chen X, Chen Y, Li WZ, Long R, Chen J, Shu YW, Liu K, et al. TNF-α upregulates Fgl2 expression in rat myocardial ischemia/reperfusion injury. Microcirculation. 2013;20(6):524–533. doi:10.1111/micc.12050

462. Camacho-Mejorado R, Gómez R, Torres-Sánchez LE, Alhelí Hernández-Tobías E, Noris G, Santana C, Magaña JJ, Orozco L, de la Peña-Díaz A, de la Luz Arenas-Sordo M, et al.ALOX5, LPA, MMP9 and TPO gene polymorphisms increase atherothrombosis susceptibility in middle-aged Mexicans. R Soc Open Sci. 2020;7(1):190775. doi:10.1098/rsos.190775

463. Volcik KA, Catellier D, Folsom AR, Matijevic N, Wasserman B, Boerwinkle E. SELP and SELPLG genetic variation is associated with cell surface measures of SELP and SELPLG: the Atherosclerosis Risk in Communities Carotid MRI Study. Clin Chem. 2009;55(6):1076–1082. doi:10.1373/clinchem.2008.119487

464. Håversen L, Sundelin JP, Mardinoglu A, Rutberg M, Ståhlman M, Wilhelmsson U, Hultén LM, Pekny M, Fogelstrand P, Bentzon JF, et al. Vimentin deficiency in macrophages induces increased oxidative stress and vascular inflammation but attenuates atherosclerosis in mice. Sci Rep. 2018;8(1):16973. doi:10.1038/s41598-018-34659-2

465. Mälarstig A, Silveira A, Wågsäter D, Öhrvik J, Bäcklund A, Samnegård A, Khademi M, Hellenius ML, Leander K, Olsson T, et al. Plasma CD93 concentration is a potential novel biomarker for coronary artery disease. J Intern Med. 2011;270(3):229–236. doi:10.1111/j.1365-2796.2011.02364.x

466. van Puijvelde GHM, Foks AC, van Bochove RE, Bot I, Habets KLL, de Jager SC, Ter Borg MND, van Osch P, Boon L, Vos M, et al. CD1d deficiency inhibits the development of abdominal aortic aneurysms in LDL receptor deficient mice. PLoS One. 2018;13(1):e0190962. doi:10.1371/journal.pone.0190962

467. Liu S, Wang X, Yu S, Yan M, Peng Y, Zhang G, Xu Z. A Meta-Analysis on the Association Between TNFSF4 Polymorphisms (rs3861950 T > C and rs1234313 A > G) and Susceptibility to Coronary Artery Disease. Front Physiol. 2020;11:539288. doi:10.3389/fphys.2020.539288

468. Cho HM, Lee KH, Shen YM, Shin TJ, Ryu PD, Choi MC, Kang KS, Cho JY. Transplantation of hMSCs Genome Edited with LEF1 Improves Cardio-Protective Effects in Myocardial Infarction. Mol Ther Nucleic Acids. 2020;19:1186–1197. doi:10.1016/j.omtn.2020.01.007

469. Głogowska-Ligus J, Dąbek J, Zych-Twardowska E, Tkacz M. Expression analysis of intercellular adhesion molecule-2 (ICAM-2) in the context of classical cardiovascular risk factors in acute coronary syndrome patients. Arch Med Sci. 2013;9(6):1035–1039. doi:10.5114/aoms.2012.28808

470. de Vicente LG, Pinto AP, da Rocha AL, Pauli JR, de Moura LP, Cintra DE, Ropelle ER, da Silva ASR. Role of TLR4 in physical exercise and cardiovascular diseases. Cytokine. 2020;136:155273. doi:10.1016/j.cyto.2020.155273

471. Bandaru S, Ala C, Salimi R, Akula MK, Ekstrand M, Devarakonda S, Karlsson J, Van den Eynden J, Bergström G, Larsson E, et al. Targeting Filamin A Reduces Macrophage Activity and Atherosclerosis. Circulation. 2019;140(1):67–79. doi:10.1161/CIRCULATIONAHA.119.039697

472. Hueso M, Mallén A, Casas Á, Guiteras J, Sbraga F, Blasco-Lucas A, Lloberas N, Torras J, Cruzado JM, Navarro E. Integrated miRNA/mRNA Counter-Expression Analysis Highlights Oxidative Stress-Related Genes CCR7 and FOXO1 as Blood Markers of Coronary Arterial Disease. Int J Mol Sci. 2020;21(6):1943. doi:10.3390/ijms21061943

473. Zhong J, Maiseyeu A, Davis SN, Rajagopalan S. DPP4 in cardiometabolic disease: recent insights from the laboratory and clinical trials of DPP4 inhibition. Circ Res. 2015;116(8):1491–1504. doi:10.1161/CIRCRESAHA.116.305665

474. Johansson Å, Eriksson N, Becker RC, Storey RF, Himmelmann A, Hagström E, Varenhorst C, Axelsson T, Barratt BJ, James SK, et al. NLRC4 Inflammasome Is an Important Regulator of Interleukin-18 Levels in Patients With Acute Coronary Syndromes: Genome-Wide Association Study in the PLATelet inhibition and patient Outcomes Trial (PLATO). Circ Cardiovasc Genet. 2015;8(3):498–506. doi:10.1161/CIRCGENETICS.114.000724

475. Ramezanpour N, Nasiri M, Akbarpour OR. Association of rs4618210A>G variant in PLCL2 gene with myocardial infarction: A case-control study in Iran. J Cardiovasc Thorac Res. 2020;12(4):303–306. doi:10.34172/jcvtr.2020.49

476. Shabana NA, Ashiq S, Ijaz A, Khalid F, Saadat IU, Khan K, Sarwar S, Shahid SU. Genetic risk score (GRS) constructed from polymorphisms in the PON1, IL-6, ITGB3, and ALDH2 genes is associated with the risk of coronary artery disease in Pakistani subjects. Lipids Health Dis. 2018;17(1):224. doi:10.1186/s12944-018-0874-6

477. Guven M, Ismailoglu Z, Batar B, Unal S, Onaran I, Karadag B, Ongen Z. The effect of genetic polymorphisms of TLR2 and TLR4 in Turkish patients with coronary artery disease. Gene. 2015;558(1):99–102. doi:10.1016/j.gene.2014.12.047

478. Wei Y, Zhu M, Corbalán-Campos J, Heyll K, Weber C, Schober A. Regulation of Csf1r and Bcl6 in macrophages mediates the stage-specific effects of microRNA-155 on atherosclerosis. Arterioscler Thromb Vasc Biol. 2015;35(4):796–803. doi:10.1161/ATVBAHA.114.304723

479. Ravi S, Schuck RN, Hilliard E, Lee CR, Dai X, Lenhart K, Willis MS, Jensen BC, Stouffer GA, Patterson C, et al. Clinical Evidence Supports a Protective Role for CXCL5 in Coronary Artery Disease. Am J Pathol. 2017;187(12):2895–2911. doi:10.1016/j.ajpath.2017.08.006

480. Li W, Zhang F, Ju C, Lv S, Huang K. The role of CD27-CD70 signaling in myocardial infarction and cardiac remodeling. Int J Cardiol. 2019;278:210–216. doi:10.1016/j.ijcard.2018.11.132

481. Shu H, Peng Y, Hang W, Nie J, Zhou N, Wang DW. The role of CD36 in cardiovascular disease. Cardiovasc Res. 2020;cvaa319. doi:10.1093/cvr/cvaa319

482. Zhang Q, Wu X, Yang J. miR-194-5p protects against myocardial ischemia/reperfusion injury via MAPK1/PTEN/AKT pathway. Ann Transl Med. 2021;9(8):654. doi:10.21037/atm-21-807

483. Karazhanova L, Zhukusheva S, Akilzhanova A. Association Between the P2RY12 Receptor Gene Polymorphism and Aspirin Resistance in Patients with Coronary Artery Disease. Cent Asian J Glob Health. 2014;3(Suppl):160. doi:10.5195/cajgh.2014.160

484. Du Y, Hu Y, Wen N, Fu S, Zhang G, Li L, Liu T, Lv X, Zhang W. Abnormal expression of TGFBR2, EGF, LRP10, and IQGAP1 is involved in the pathogenesis of coronary artery disease. Rev Cardiovasc Med. 2021;22(3):947–958. doi:10.31083/j.rcm2203103

485. Kroupis C, Theodorou M, Chaidaroglou A, Dalamaga M, Oliveira SC, Cokkinos DV, Degiannis D, Manginas A. The association between a common FCGR2A polymorphism and C-reactive protein and coronary artery disease revisited. Genet Test Mol Biomarkers. 2010;14(6):839–846. doi:10.1089/gtmb.2010.0108

486. Liu C, Zhang H, Chen Y, Wang S, Chen Z, Liu Z, Wang J. Identifying RBM47, HCK, CD53, TYROBP, and HAVCR2 as Hub Genes in Advanced Atherosclerotic Plaques by Network-Based Analysis and Validation. Front Genet. 2021;11:602908. doi:10.3389/fgene.2020.602908

487. van der Laan SW, Foroughi Asl H, van den Borne P, van Setten J, van der Perk ME, van de Weg SM, Schoneveld AH, de Kleijn DP, Michoel T, et al. Variants in ALOX5, ALOX5AP and LTA4H are not associated with atherosclerotic plaque phenotypes: the Athero-Express Genomics Study. Atherosclerosis. 2015;239(2):528–538. doi:10.1016/j.atherosclerosis.2015.01.018

488. Xie M, Hu C, Li D, Li S. MicroRNA-377 Alleviates Myocardial Injury Induced by Hypoxia/Reoxygenation via Downregulating LILRB2 Expression. Dose Response. 2020;18(2):1559325820936124. doi:10.1177/1559325820936124

489. Shen X, Li H, Li W, Wu X, Sun Z, Ding X. Telmisartan ameliorates adipoR1 and adipoR2 expression via PPAR-γ activation in the coronary artery and VSMCs. Biomed Pharmacother. 2017;95:129–136. doi:10.1016/j.biopha.2017.08.041

490. Yu H, Rimbert A, Palmer AE, Toyohara T, Xia Y, Xia F, Ferreira LMR, Chen Z, Chen T, Loaiza N, et al. GPR146 Deficiency Protects against Hypercholesterolemia and Atherosclerosis. Cell. 2019;179(6):1276–1288.e14. doi:10.1016/j.cell.2019.10.034

491. Patel VB, McLean BA, Chen X, Oudit GY. Regulators of G-Protein Signaling 10 and Heart Failure: The Importance of Negative Regulators of Heart Disease. Hypertension. 2016;67(1):38–40. doi:10.1161/HYPERTENSIONAHA.115.06109

492. Chabior A, Gąsecka A, Marchel M, Gozdowska R, Makowska A, Maciak K, Góra M, Filipiak KJ, Burzyńska B, Opolski G. Expression of versican mRNA transcript to predict cardiac remodelling after myocardial infarction. Kardiol Pol. 2021;79(7-8):833–840. doi:10.33963/KP.a2021.0042

493. Wang P, Wang Y, Peng H, Wang J, Zheng Q, Wang P, Wang J, Zhang H, Huang Y, Xiong L, et al. Functional rare variant in a C/EBPbeta binding site in NINJ2 gene increases the risk of coronary artery disease. Aging (Albany NY). 2021;13(23):25393–25407. doi:10.18632/aging.203755

494. He Q, Shao D, Hao S, Yuan Y, Liu H, Liu F, Mu Q. CircSCAP Aggravates Oxidized Low-density Lipoprotein-induced Macrophage Injury by Upregulating PDE3B by miR-221-5p in Atherosclerosis. J Cardiovasc Pharmacol. 2021;78(5):e749–e760. doi:10.1097/FJC.0000000000001118

495. Zhang X, Zhang MC, Wang CT. Loss of LRRC25 accelerates pathological cardiac hypertrophy through promoting fibrosis and inflammation regulated by TGF-β1. Biochem Biophys Res Commun. 2018;506(1):137–144. doi:10.1016/j.bbrc.2018.09.065

496. Lalonde S, Codina-Fauteux VA, de Bellefon SM, Leblanc F, Beaudoin M, Simon MM, Dali R, Kwan T, Lo KS, Pastinen T, et al. Integrative analysis of vascular endothelial cell genomic features identifies AIDA as a coronary artery disease candidate gene. Genome Biol. 2019;20(1):133. doi:10.1186/s13059-019-1749-5

497. Belleville-Rolland T, Sassi Y, Decouture B, Dreano E, Hulot JS, Gaussem P, Bachelot-Loza C. MRP4 (ABCC4) as a potential pharmacologic target for cardiovascular disease. Pharmacol Res. 2016;107:381–389. doi:10.1016/j.phrs.2016.04.002

498. Linke AT, Marchant B, Marsh P, Frampton G, Murphy J, Rose ML. Screening of a HUVEC cDNA library with transplant-associated coronary artery disease sera identifies RPL7 as a candidate autoantigen associated with this disease. Clin Exp Immunol. 2001;126(1):173–179. doi:10.1046/j.1365-2249.2001.01654.x

499. Sun Y, Chen C, Xue R, Wang Y, Dong B, Li J, Chen C, Jiang J, Fan W, Liang Z, et al. Maf1 ameliorates cardiac hypertrophy by inhibiting RNA polymerase III through ERK1/2. Theranostics. 2019;9(24):7268–7281. doi:10.7150/thno.33006

500. Konishi H, Okuda A, Ohno Y, Kihara A. Characterization of HACD1 K64Q mutant found in arrhythmogenic right ventricular dysplasia patients. J Biochem. 2010;148(5):617–622. doi:10.1093/jb/mvq092

501. Zhang CL, Long TY, Bi SS, Sheikh SA, Li F. CircPAN3 ameliorates myocardial ischaemia/reperfusion injury by targeting miR-421/Pink1 axis-mediated autophagy suppression. Lab Invest. 2021;101(1):89–103. doi:10.1038/s41374-020-00483-4

502. de Vries MA, Trompet S, Mooijaart SP, Smit RA, Böhringer S, Castro Cabezas M, Jukema JW. Complement receptor 1 gene polymorphisms are associated with cardiovascular risk. Atherosclerosis. 2017;257:16–21. doi:10.1016/j.atherosclerosis.2016.12.017

503. Watson LJ, Facundo HT, Ngoh GA, Ameen M, Brainard RE, Lemma KM, Long BW, Prabhu SD, Xuan YT, Jones SP. O-linked β-N-acetylglucosamine transferase is indispensable in the failing heart. Proc Natl Acad Sci U S A. 2010;107(41):17797–17802. doi:10.1073/pnas.1001907107

504. Ni J, Huang Z, Wang D. LncRNA TP73-AS1 promotes oxidized low-density lipoprotein-induced apoptosis of endothelial cells in atherosclerosis by targeting the miR-654-3p/AKT3 axis. Cell Mol Biol Lett. 2021;26(1):27. doi:10.1186/s11658-021-00264-x

505. Navarro-Núñez L, Roldán V, Lozano ML, Rivera J, Marin F, Vicente V, Martínez C.TUBB1 Q43P polymorphism does not protect against acute coronary syndrome and premature myocardial infarction. Thromb Haemost. 2008;100(6):1211–1213.

506. Vishwajeet V, Jamwal M, Sharma P, Das R, Ahluwalia J, Dogra RK, Rohit MK. Coagulation F13A1 V34L, fibrinogen and homocysteine versus conventional risk factors in the pathogenesis of MI in young persons. Acta Cardiol. 2018;73(4):328–334. doi:10.1080/00015385.2017.1384172

507. Maillet M, Purcell NH, Sargent MA, York AJ, Bueno OF, Molkentin JD. DUSP6 (MKP3) null mice show enhanced ERK1/2 phosphorylation at baseline and increased myocyte proliferation in the heart affecting disease susceptibility. J Biol Chem. 2008;283(45):31246–31255. doi:10.1074/jbc.M806085200

508. Xia XD, Zhou Z, Yu XH, Zheng XL, Tang CK. Myocardin: A novel player in atherosclerosis. Atherosclerosis. 2017;257:266–278. doi:10.1016/j.atherosclerosis.2016.12.002

509. Nie XY, Li JL, Qin SB, Fu Y, Liang GK, Shi LW, Shao H, Liu J, Lu Y. Genetic mutations in PEAR1 associated with cardiovascular outcomes in Chinese patients with acute coronary syndrome. Thromb Res. 2018;163:77–82. doi:10.1016/j.thromres.2018.01.026

510. Runhua M, Qiang J, Yunqing S, Wenjun D, Chunsheng W. FSTL3 Induces Lipid Accumulation and Inflammatory Response in Macrophages and Associates With Atherosclerosis. J Cardiovasc Pharmacol. 2019;74(6):566–573. doi:10.1097/FJC.0000000000000742

511. Werner P, Paluru P, Simpson AM, Latney B, Iyer R, Brodeur GM, Goldmuntz E. Mutations in NTRK3 suggest a novel signaling pathway in human congenital heart disease. Hum Mutat. 2014;35(12):1459–1468. doi:10.1002/humu.22688

512. Hardy SA, Mabotuwana NS, Murtha LA, Coulter B, Sanchez-Bezanilla S, Al-Omary MS, Senanayake T, Loering S, Starkey M, Lee RJ, et al. Novel role of extracellular matrix protein 1 (ECM1) in cardiac aging and myocardial infarction. PLoS One. 2019;14(2):e0212230. doi:10.1371/journal.pone.0212230

513. Tavakolian Ferdousie V, Mohammadi M, Hassanshahi G, Khorramdelazad H, Khanamani Falahati-Pour S, Mirzaei M, Allah Tavakoli M, Kamiab Z, Ahmadi Z, Vazirinejad R, et al. Serum CXCL10 and CXCL12 chemokine levels are associated with the severity of coronary artery disease and coronary artery occlusion. Int J Cardiol. 2017;233:23–28. doi:10.1016/j.ijcard.2017.02.011

514. Gao L, Guo Y, Liu X, Shang D, Du Y. KLF15 protects against isoproterenol-induced cardiac hypertrophy via regulation of cell death and inhibition of Akt/mTOR signaling. Biochem Biophys Res Commun. 2017;487(1):22–27. doi:10.1016/j.bbrc.2017.03.087

515. Wang J, Li G, Yu D, Wong YP, Yong TF, Liang MC, Liao P, Foo R, Hoppe UC, Soong TW. Characterization of CaV1.2 exon 33 heterozygous knockout mice and negative correlation between Rbfox1 and CaV1.2 exon 33 expressions in human heart failure. Channels (Austin). 2018;12(1):51–57. doi:10.1080/19336950.2017.1381805

516. Xiong F, Li Q, Zhang C, Chen Y, Li P, Wei X, Li Q, Zhou W, Li L, Shang X, et al. Analyses of GATA4, NKX2.5, and TFAP2B genes in subjects from southern China with sporadic congenital heart disease. Cardiovasc Pathol. 2013;22(2):141–145. doi:10.1016/j.carpath.2012.07.001

517. Koch W, Hoppmann P, Michou E, Jung V, Pfeufer A, Mueller JC, Gieger C, Wichmann HE, Meitinger T, Schömig A, et al. Association of variants in the BAT1-NFKBIL1-LTA genomic region with protection against myocardial infarction in Europeans. Hum Mol Genet. 2007;16(15):1821–1827. doi:10.1093/hmg/ddm130

518. Arvanitis M, Tampakakis E, Zhang Y, Wang W, Auton A; 23andMe Research Team, Dutta D, Glavaris S, Keramati A, Chatterjee N, et al. Genome-wide association and multi-omic analyses reveal ACTN2 as a gene linked to heart failure. Nat Commun. 2020;11(1):1122. doi:10.1038/s41467-020-14843-7

519. Wong WM, Stephens JW, Acharya J, Hurel SJ, Humphries SE, Talmud PJ. The APOA4 T347S variant is associated with reduced plasma TAOS in subjects with diabetes mellitus and cardiovascular disease. J Lipid Res. 2004;45(8):1565–1571. doi:10.1194/jlr.M400130-JLR200

520. Guo DC, Papke CL, Tran-Fadulu V, Regalado ES, Avidan N, Johnson RJ, Kim DH, Pannu H, Willing MC, Sparks E, et al. Mutations in smooth muscle alpha-actin (ACTA2) cause coronary artery disease, stroke, and Moyamoya disease, along with thoracic aortic disease. Am J Hum Genet. 2009;84(5):617–627. doi:10.1016/j.ajhg.2009.04.007

521. Avan A, Tavakoly Sany SB, Ghayour-Mobarhan M, Rahimi HR, Tajfard M, Ferns G. Serum C-reactive protein in the prediction of cardiovascular diseases: Overview of the latest clinical studies and public health practice. J Cell Physiol. 2018;233(11):8508–8525. doi:10.1002/jcp.26791

522. Yang W, Wu Z, Yang K, Han Y, Chen Y, Zhao W, Huang F, Jin Y, Jin W. BMI1 promotes cardiac fibrosis in ischemia-induced heart failure via the PTEN-PI3K/Akt-mTOR signaling pathway. Am J Physiol Heart Circ Physiol. 2019;316(1):H61–H69. doi:10.1152/ajpheart.00487.2018

523. Geng GY, Liu HL, Zhao YJ, Wu L, Mao L, Ba N. Correlation between polymorphisms in the IL-17A and IL-17F genes and development of coronary artery disease. Genet Mol Res. 2015;14(3):11488–11494. doi:10.4238/2015.September.25.15

524. Prasongsukarn K, Dechkhajorn W, Benjathummarak S, Maneerat Y. TRPM2, PDLIM5, BCL3, CD14, GBA Genes as Feasible Markers for Premature Coronary Heart Disease Risk. Front Genet. 2021;12:598296. doi:10.3389/fgene.2021.598296

525. Huang CY, Kuo CH, Pai PY, Ho TJ, Lin YM, Chen RJ, Tsai FJ, Vijaya Padma V, Kuo WW, Huang CY. Inhibition of HSF2 SUMOylation via MEL18 upregulates IGF-IIR and leads to hypertension-induced cardiac hypertrophy. Int J Cardiol. 2018;257:283–290. doi:10.1016/j.ijcard.2017.10.102

526. Peng Y, Meng K, Jiang L, Zhong Y, Yang Y, Lan Y, Zeng Q, Cheng L. Thymic stromal lymphopoietin-induced HOTAIR activation promotes endothelial cell proliferation and migration in atherosclerosis. Biosci Rep. 2017;37(4):BSR20170351. doi:10.1042/BSR20170351

527. Kane GC, Behfar A, Dyer RB, O’Cochlain DF, Liu XK, Hodgson DM, Reyes S, Miki T, Seino S, Terzic A. KCNJ11 gene knockout of the Kir6.2 KATP channel causes maladaptive remodeling and heart failure in hypertension. Hum Mol Genet. 2006;15(15):2285–2297. doi:10.1093/hmg/ddl154

528. Hedayati-Moghadam M, Hosseinian S, Paseban M, Shabgah AG, Gholizadeh J, Jamialahmadi T, Sathyapalan T, Sahebkar A. The Role of Chemokines in Cardiovascular Diseases and the Therapeutic Effect of Curcumin on CXCL8 and CCL2 as Pathological Chemokines in Atherosclerosis. Adv Exp Med Biol. 2021;1328:155–170. doi:10.1007/978-3-030-73234-9_11

529. Nowzari Z, Masoumi M, Nazari-Robati M, Akbari H, Shahrokhi N, Asadikaram G. Association of polymorphisms of leptin, leptin receptor and apelin receptor genes with susceptibility to coronary artery disease and hypertension. Life Sci. 2018;207:166–171. doi:10.1016/j.lfs.2018.06.007

530. Yang L, Zhu X, Ni Y, Wu D, Tian Y, Chen Z, Li M, Zhang H, Liang D. MicroRNA-34c Inhibits Osteogenic Differentiation and Valvular Interstitial Cell Calcification via STC1-Mediated JNK Pathway in Calcific Aortic Valve Disease. Front Physiol. 2020;11:829. doi:10.3389/fphys.2020.00829

531. Cristo F, Inácio JM, de Almeida S, Mendes P, Martins DS, Maio J, Anjos R, Belo JA. Functional study of DAND5 variant in patients with Congenital Heart Disease and laterality defects. BMC Med Genet. 2017;18(1):77. doi:10.1186/s12881-017-0444-1

532. Landry NM, Cohen S, Dixon IMC. Periostin in cardiovascular disease and development: a tale of two distinct roles. Basic Res Cardiol. 2017;113(1):1. doi:10.1007/s00395-017-0659-5

533. Awad SM, Attallah DA, Salama RH, Mahran AM, Abu El-Hamed E. Serum levels of psoriasin (S100A7) and koebnerisin (S100A15) as potential markers of atherosclerosis in patients with psoriasis. Clin Exp Dermatol. 2018;43(3):262–267. doi:10.1111/ced.13370

534. Burke RM, Lighthouse JK, Quijada P, Dirkx RA Jr, Rosenberg A, Moravec CS, Alexis JD, Small EM. Small proline-rich protein 2B drives stress-dependent p53 degradation and fibroblast proliferation in heart failure. Proc Natl Acad Sci U S A. 2018;115(15):E3436–E3445. doi:10.1073/pnas.1717423115

535. Moyer AM, Walker DL, Avula R, Lapid MI, Kung S, Bryant SC, Edwards KK, Black JL, Karpyak VM, Shinozaki G, et al. Relationship of genetic variation in the serotonin transporter gene (SLC6A4) and congenital and acquired cardiovascular diseases. Genet Test Mol Biomarkers. 2015;19(3):115–123. doi:10.1089/gtmb.2014.0250

536. Martin RI, Babaei MS, Choy MK, Owens WA, Chico TJ, Keenan D, Yonan N, Koref MS, Keavney BD. Genetic variants associated with risk of atrial fibrillation regulate expression of PITX2, CAV1, MYOZ1, C9orf3 and FANCC. J Mol Cell Cardiol. 2015;85:207–214. doi:10.1016/j.yjmcc.2015.06.005

537. Silbernagel G, Chapman MJ, Genser B, Kleber ME, Fauler G, Scharnagl H, Grammer TB, Boehm BO, Mäkelä KM, Kähönen M, et al. High intestinal cholesterol absorption is associated with cardiovascular disease and risk alleles in ABCG8 and ABO: evidence from the LURIC and YFS cohorts and from a meta-analysis. J Am Coll Cardiol. 2013;62(4):291–299. doi:10.1016/j.jacc.2013.01.100

538. Mormile R. Multiple sclerosis and susceptibility to cardiovascular diseases: Implications of ethnicity-related interleukin-17A gene polymorphism?. Med Hypotheses. 2015;85(3):365–366. doi:10.1016/j.mehy.2015.06.006

539. Li SH, Zhang YY, Sun YL, Zhao HJ, Wang Y. Inhibition of microRNA-802-5p inhibits myocardial apoptosis after myocardial infarction via Sonic Hedgehog signaling pathway by targeting PTCH1. Eur Rev Med Pharmacol Sci. 2021;25(1):326–334. doi:10.26355/eurrev_202101_24398

540. Cheng H, Yan W. MiR-433 Regulates Myocardial Ischemia Reperfusion Injury by Targeting NDRG4 Via the PI3K/Akt Pathway. Shock. 2020;54(6):802–809. doi:10.1097/SHK.0000000000001532

541. Huang H, Zeng Z, Zhang L, Liu R, Li X, Qiang O, Chen Y. Implication of genetic variants near TMEM18, BCDIN3D/FAIM2, and MC4R with coronary artery disease and obesity in Chinese: a angiography-based study. Mol Biol Rep. 2012;39(2):1739–1744. doi:10.1007/s11033-011-0914-z

542. Wang M, Gu M, Li Z, Lian X, Shen H, Dai Z, Zhang Z, Liu X. TNFRSF11B polymorphisms predict poor outcome after large artery atherosclerosis stroke. Gene. 2020;743:144617. doi:10.1016/j.gene.2020.144617

543. Wang B, Yan J, Peng Z, Wang J, Liu S, Xie X, Ma X. Teratocarcinoma-derived growth factor 1 (TDGF1) sequence variants in patients with congenital heart defect. Int J Cardiol. 2011;146(2):225–227. doi:10.1016/j.ijcard.2009.08.046

544. Lin CF, Su CJ, Liu JH, Chen ST, Huang HL, Pan SL. Potential Effects of CXCL9 and CCL20 on Cardiac Fibrosis in Patients with Myocardial Infarction and Isoproterenol-Treated Rats. J Clin Med. 2019;8(5):659. doi:10.3390/jcm8050659

545. Brunette CA, Vassy JL. The role of SLCO1B1 genotyping in lowering cardiovascular risk. Pharmacogenomics. 2021;22(11):649–656. doi:10.2217/pgs-2021-0075

546. Zhao B, Li G, Peng J, Ren L, Lei L, Ye H, Wang Z, Zhao S. CircMACF1 Attenuates Acute Myocardial Infarction Through miR-500b-5p-EMP1 Axis. J Cardiovasc Transl Res. 2021;14(1):161–172. doi:10.1007/s12265-020-09976-5

547. Du L, Zhang H, Zhao H, Cheng X, Qin J, Teng T, Yang Q, Xu Z. The critical role of the zinc transporter Zip2 (SLC39A2) in ischemia/reperfusion injury in mouse hearts. J Mol Cell Cardiol. 2019;132:136–145. doi:10.1016/j.yjmcc.2019.05.011

548. Hu G, Zhu Q, Wang W, Xie D, Chen C, Li PL, Ritter JK, Li N. Collecting duct-specific knockout of sphingosine-1-phosphate receptor 1 aggravates DOCA-salt hypertension in mice. J Hypertens. 2021;39(8):1559–1566. doi:10.1097/HJH.0000000000002809

549. Zhang L, Zeng XX, Li YM, Chen SK, Tang LY, Wang N, Yang X, Lin MJ. Keratin 1 attenuates hypoxic pulmonary artery hypertension by suppressing pulmonary artery media smooth muscle expansion. Acta Physiol (Oxf). 2021;231(2):e13558. doi:10.1111/apha.13558

550. Abid S, Marcos E, Parpaleix A, Amsellem V, Breau M, Houssaini A, Vienney N, Lefevre M, Derumeaux G, Evans S, et al. CCR2/CCR5-mediated macrophage-smooth muscle cell crosstalk in pulmonary hypertension. Eur Respir J. 2019;54(4):1802308. doi:10.1183/13993003.02308-2018

551. Zhang L, Sun Y, Zhang X, Shan X, Li J, Yao Y, Shu Y, Lin K, Huang X, Yang Z, et al. Three Novel Genetic Variants in the FAM110D, CACNA1A, and NLRP12 Genes Are Associated With Susceptibility to Hypertension Among Dai People. Am J Hypertens. 2021;34(8):874–879. doi:10.1093/ajh/hpab040

552. Satoh K, Kikuchi N, Shimokawa H. PIM1 (Provirus Integration Site For Moloney Murine Leukemia Virus) as a Novel Biomarker and Therapeutic Target in Pulmonary Arterial Hypertension: Another Evidence for Cancer Theory. Arterioscler Thromb Vasc Biol. 2020;40(3):500–502. doi:10.1161/ATVBAHA.120.313975

553. Ahadzadeh E, Rosendahl A, Czesla D, Steffens P, Prüßner L, Meyer-Schwesinger C, Wanner N, Paust HJ, Huber TB, Stahl RAK, et al. The chemokine receptor CX3CR1 reduces renal injury in mice with angiotensin II-induced hypertension. Am J Physiol Renal Physiol. 2018;315(6):F1526–F1535. doi:10.1152/ajprenal.00149.2018

554. Perros F, Cohen-Kaminsky S, Gambaryan N, Girerd B, Raymond N, Klingelschmitt I, Huertas A, Mercier O, Fadel E, Simonneau G, et al. Cytotoxic cells and granulysin in pulmonary arterial hypertension and pulmonary veno-occlusive disease. Am J Respir Crit Care Med. 2013;187(2):189–196. doi:10.1164/rccm.201208-1364OC

555. Morris BJ, Chen R, Donlon TA, Evans DS, Tranah GJ, Parimi N, Ehret GB, Newton-Cheh C, Seto T, Willcox DC, et al. Association Analysis of FOXO3 Longevity Variants With Blood Pressure and Essential Hypertension. Am J Hypertens. 2016;29(11):1292–1300. doi:10.1093/ajh/hpv171

556. Vinh A, Chen W, Blinder Y, Weiss D, Taylor WR, Goronzy JJ, Weyand CM, Harrison DG, Guzik TJ.Inhibition and genetic ablation of the B7/CD28 T-cell costimulation axis prevents experimental hypertension. Circulation. 2010;122(24):2529–2537. doi:10.1161/CIRCULATIONAHA.109.930446

557. Ardanaz N, Yang XP, Cifuentes ME, Haurani MJ, Jackson KW, Liao TD, Carretero OA, Pagano PJ. Lack of glutathione peroxidase 1 accelerates cardiac-specific hypertrophy and dysfunction in angiotensin II hypertension. Hypertension. 2010;55(1):116–123. doi:10.1161/HYPERTENSIONAHA.109.135715

558. Nie X, Tan J, Dai Y, Liu Y, Zou J, Sun J, Ye S, Shen C, Fan L, Chen J, et al. CCL5 deficiency rescues pulmonary vascular dysfunction, and reverses pulmonary hypertension via caveolin-1-dependent BMPR2 activation. J Mol Cell Cardiol. 2018;116:41–56. doi:10.1016/j.yjmcc.2018.01.016

559. Song R, Lei S, Yang S, Wu SJ. LncRNA PAXIP1-AS1 fosters the pathogenesis of pulmonary arterial hypertension via ETS1/WIPF1/RhoA axis. J Cell Mol Med. 2021;25(15):7321–7334. doi:10.1111/jcmm.16761

560. Fan C, Wang J, Mao C, Li W, Liu K, Wang Z. The FGL2 prothrombinase contributes to the pathological process of experimental pulmonary hypertension. J Appl Physiol (1985). 2019;127(6):1677–1687. doi:10.1152/japplphysiol.00396.2019

561. Almodovar S, Wade BE, Porter KM, Smith JM, Lopez-Astacio RA, Bijli K, Kang BY, Cribbs SK, Guidot DM, Molehin D, et al. HIV X4 Variants Increase Arachidonate 5-Lipoxygenase in the Pulmonary Microenvironment and are associated with Pulmonary Arterial Hypertension. Sci Rep. 2020;10(1):11696. doi:10.1038/s41598-020-68060-9

562. Grylls A, Seidler K, Neil J. Link between microbiota and hypertension: Focus on LPS/TLR4 pathway in endothelial dysfunction and vascular inflammation, and therapeutic implication of probiotics. Biomed Pharmacother. 2021;137:111334. doi:10.1016/j.biopha.2021.111334

563. Liu C, Tang W, Zhao H, Yang S, Ren Z, Li J, Chen Y, Zhao X, Xu D, Zhao Y, et al. The variants at FLNA and FLNB contribute to the susceptibility of hypertension and stroke with differentially expressed mRNA. Pharmacogenomics J. 2021;21(4):458–466. doi:10.1038/s41397-021-00222-y

564. Cai M, Li X, Dong H, Wang Y, Huang X. CCR7 and its related molecules may be potential biomarkers of pulmonary arterial hypertension. FEBS Open Bio. 2021;11(6):1565–1578. doi:10.1002/2211-5463.13130

565. Li Y, Yang L, Dong L, Yang ZW, Zhang J, Zhang SL, Niu MJ, Xia JW, Gong Y, Zhu N, et al. Crosstalk between the Akt/mTORC1 and NF-κB signaling pathways promotes hypoxia-induced pulmonary hypertension by increasing DPP4 expression in PASMCs. Acta Pharmacol Sin. 2019;40(10):1322–1333. doi:10.1038/s41401-019-0272-2

566. Liu A, Liu Y, Li B, Yang M, Liu Y, Su J. Role of miR-223-3p in pulmonary arterial hypertension via targeting ITGB3 in the ECM pathway. Cell Prolif. 2019;52(2):e12550. doi:10.1111/cpr.12550

567. Kurahara LH, Hiraishi K, Yamamura A, Zhang Y, Abe K, Yahiro E, Aoki M, Koga K, Yokomise H, Go T, et al. Eicosapentaenoic acid ameliorates pulmonary hypertension via inhibition of tyrosine kinase Fyn. J Mol Cell Cardiol. 2020;148:50–62. doi:10.1016/j.yjmcc.2020.08.013

568. Holmes L Jr, Lim A, Comeaux CR, Dabney KW, Okundaye O. DNA Methylation of Candidate Genes (ACE II, IFN-γ, AGTR 1, CKG, ADD1, SCNN1B and TLR2) in Essential Hypertension: A Systematic Review and Quantitative Evidence Synthesis. Int J Environ Res Public Health. 2019;16(23):4829. doi:10.3390/ijerph16234829

569. Santulli G, Cipolletta E, Sorriento D, Del Giudice C, Anastasio A, Monaco S, Maione AS, Condorelli G, Puca A, Trimarco B, et al. CaMK4 Gene Deletion Induces Hypertension. J Am Heart Assoc. 2012;1(4):e001081. doi:10.1161/JAHA.112.001081

570. Beitelshees AL, Aquilante CL, Allayee H, Langaee TY, Welder GJ, Schofield RS, Zineh I. CXCL5 polymorphisms are associated with variable blood pressure in cardiovascular disease-free adults. Hum Genomics. 2012;6(1):9. doi:10.1186/1479-7364-6-9

571. Pravenec M, Churchill PC, Churchill MC, Viklicky O, Kazdova L, Aitman TJ, Petretto E, Hubner N, Wallace CA, Zimdahl H, et al. Identification of renal Cd36 as a determinant of blood pressure and risk for hypertension. Nat Genet. 2008;40(8):952–954. doi:10.1038/ng.164

572. Chen J, Zhao X, Wang H, Chen Y, Wang W, Zhou W, Wang X, Tang J, Zhao Y, Lu X, et al. Common variants in TGFBR2 and miR-518 genes are associated with hypertension in the Chinese population. Am J Hypertens. 2014;27(10):1268–1276. doi:10.1093/ajh/hpu047

573. Chang YT, Chan CK, Eriksson I, Johnson PY, Cao X, Westöö C, Norvik C, Andersson-Sjöland A, Westergren-Thorsson G, Johansson S, et al. Versican accumulates in vascular lesions in pulmonary arterial hypertension. Pulm Circ. 2016;6(3):347–359. doi:10.1086/686994

574. Pei Q, Yang L, Tan HY, Liu SK, Liu Y, Huang L, Li RH, Wan Q, Huang J, Guo CX, et al. Effects of genetic variants in UGT1A1, SLCO1B3, ABCB1, ABCC2, ABCG2, ORM1 on PK/PD of telmisartan in Chinese patients with mild to moderate essential hypertensionL. Int J Clin Pharmacol Ther. 2017;55(8):659–665. doi:10.5414/CP202744

575. Park HK, Kim DH, Yun DH, Ban JY. Association between IL10, IL10RA, and IL10RB SNPs and ischemic stroke with hypertension in Korean population. Mol Biol Rep. 2013;40(2):1785–1790. doi:10.1007/s11033-012-2232-5

576. Claude C, Mougenot N, Bechaux J, Hadri L, Brockschnieder D, Clergue M, Atassi F, Lompré AM, Hulot JS. Inhalable delivery of AAV-based MRP4/ABCC4 silencing RNA prevents monocrotaline-induced pulmonary hypertension. Mol Ther Methods Clin Dev. 2015;2:14065. doi:10.1038/mtm.2014.65

577. Saraji A, Sydykov A, Schäfer K, Garcia-Castro CF, Henneke I, Alebrahimdehkordi N, Kosanovic D, Hadzic S, Guenther A, Hecker M, et al. PINK1-mediated Mitophagy Contributes to Pulmonary Vascular Remodeling in Pulmonary Hypertension. Am J Respir Cell Mol Biol. 2021;65(2):226–228. doi:10.1165/rcmb.2021-0082LE

578. Acuña R, Martínez-de-la-Maza L, Ponce-Coria J, Vázquez N, Ortal-Vite P, Pacheco-Alvarez D, Bobadilla NA, Gamba G. Rare mutations in SLC12A1 and SLC12A3 protect against hypertension by reducing the activity of renal salt cotransporters. J Hypertens. 2011;29(3):475–483. doi:10.1097/HJH.0b013e328341d0fd

579. Barnes JW, Tian L, Heresi GA, Farver CF, Asosingh K, Comhair SA, Aulak KS, Dweik RA. O-linked β-N-acetylglucosamine transferase directs cell proliferation in idiopathic pulmonary arterial hypertension. Circulation. 2015;131(14):1260–1268. doi:10.1161/CIRCULATIONAHA.114.01387

580. Sahoo S, Meijles DN, Al Ghouleh I, Tandon M, Cifuentes-Pagano E, Sembrat J, Rojas M, Goncharova E, Pagano PJ. MEF2C-MYOCD and Leiomodin1 Suppression by miRNA-214 Promotes Smooth Muscle Cell Phenotype Switching in Pulmonary Arterial Hypertension. PLoS One. 2016;11(5):e0153780. doi:10.1371/journal.pone.0153780

581. Olivi L, Vandenbriele C, Gu YM, Salvi E, Carpini SD, Liu YP, Jacobs L, Jin Y, Thijs L, Citterio L, et al. PEAR1 is not a human hypertension-susceptibility gene. Blood Press. 2015;24(1):61–64. doi:10.3109/08037051.2014.986928

582. Antonelli A, Fallahi P, Ferrari SM, Ghiadoni L, Virdis A, Mancusi C, Centanni M, Taddei S, Ferrannini E. High serum levels of CXC (CXCL10) and CC (CCL2) chemokines in untreated essential hypertension. Int J Immunopathol Pharmacol. 2012;25(2):387–395. doi:10.1177/039463201202500208

583. Jiang J, Xia Y, Liang Y, Yang M, Zeng W, Zeng X. miR-190a-5p participates in the regulation of hypoxia-induced pulmonary hypertension by targeting KLF15 and can serve as a biomarker of diagnosis and prognosis in chronic obstructive pulmonary disease complicated with pulmonary hypertension. Int J Chron Obstruct Pulmon Dis. 2018;13:3777–3790. doi:10.2147/COPD.S182504

584. Kominami S, Tanabe N, Ota M, Naruse TK, Katsuyama Y, Nakanishi N, Tomoike H, Sakuma M, Shirato K, Takahashi M, et al. HLA-DPB1 and NFKBIL1 may confer the susceptibility to chronic thromboembolic pulmonary hypertension in the absence of deep vein thrombosis. J Hum Genet. 2009;54(2):108–114. doi:10.1038/jhg.2008.15

585. Marks SD, Gullett AM, Brennan E, Tullus K, Jaureguiberry G, Klootwijk E, Stanescu HC, Kleta R, Woolf AS. Renal FMD may not confer a familial hypertensive risk nor is it caused by ACTA2 mutations. Pediatr Nephrol. 2011;26(10):1857–1861. doi:10.1007/s00467-011-1891-0

586. Hage FG. C-reactive protein and hypertension. J Hum Hypertens. 2014;28(7):410-415. doi:10.1038/jhh.2013.111

587. Li K, Li Y, Yu Y, Ding J, Huang H, Chu C, Hu L, Yu Y, Cao Y, Xu P, et al. Bmi-1 alleviates adventitial fibroblast senescence by eliminating ROS in pulmonary hypertension. BMC Pulm Med. 2021;21(1):80. doi:10.1186/s12890-021-01439-0

588. Diana MC, Santoro ML, Xavier G, Santos CM, Spindola LN, Moretti PN, Ota VK, Bressan RA, Abilio VC, Belangero SI. Low expression of Gria1 and Grin1 glutamate receptors in the nucleus accumbens of Spontaneously Hypertensive Rats (SHR). Psychiatry Res. 2015;229(3):690–694. doi:10.1016/j.psychres.2015.08.021

589. Liu G, Wan N, Liu Q, Chen Y, Cui H, Wang Y, Ren J, Shen X, Lu W, Yu Y, et al. Resolvin E1 Attenuates Pulmonary Hypertension by Suppressing Wnt7a/β-Catenin Signaling. Hypertension. 2021;78(6):1914–1926. doi:10.1161/HYPERTENSIONAHA.121.17809

590. Singh MV, Cicha MZ, Kumar S, Meyerholz DK, Irani K, Chapleau MW, Abboud FM. Abnormal CD161+ immune cells and retinoic acid receptor-related orphan receptor γt-mediate enhanced IL-17F expression in the setting of genetic hypertension. J Allergy Clin Immunol. 2017;140(3):809–821.e3. doi:10.1016/j.jaci.2016.11.039

591. Shimizu Y, Sato S, Koyamatsu J, Yamanashi H, Nagayoshi M, Kadota K, Kawashiri SY, Maeda T. Association between high-density lipoprotein-cholesterol and hypertension in relation to circulating CD34-positive cell levels. J Physiol Anthropol. 2017;36(1):26. doi:10.1186/s40101-017-0143-9

592. Zhancheng W, Wenhui J, Yun J, Lingli W, Huijun H, Yan S, Jin L. The dominant models of KCNJ11 E23K and KCNMB1 E65K are associated with essential hypertension (EH) in Asian: Evidence from a meta-analysis. Medicine (Baltimore). 2019;98(23):e15828. doi:10.1097/MD.0000000000015828

593. Alsheikh AJ, Dasinger JH, Abais-Battad JM, Fehrenbach DJ, Yang C, Cowley AW Jr, Mattson DL. CCL2 mediates early renal leukocyte infiltration during salt-sensitive hypertension. Am J Physiol Renal Physiol. 2020;318(4):F982–F993. doi:10.1152/ajprenal.00521.2019

594. Zaw AM, Sekar R, Mak SOK, Law HKW, Chow BKC. Loss of secretin results in systemic and pulmonary hypertension with cardiopulmonary pathologies in mice. Sci Rep. 2019;9(1):14211. doi:10.1038/s41598-019-50634-x

595. Zhang F, Yang M, Xiao T, Hua Y, Chen Y, Xu S, Ni C. SLC6A4 gene L/S polymorphism and susceptibility to pulmonary arterial hypertension: a meta-analysis. J Int Med Res. 2020;48(9):300060520935309. doi:10.1177/0300060520935309

596. Davis GK, Fehrenbach DJ, Madhur MS. Interleukin 17A: Key Player in the Pathogenesis of Hypertension and a Potential Therapeutic Target. Curr Hypertens Rep. 2021;23(3):13. doi:10.1007/s11906-021-01128-7

597. Le MT, Lobmeyer MT, Campbell M, Cheng J, Wang Z, Turner ST, Chapman AB, Boerwinkle E, Gums JG, Gong Y, et al. Impact of genetic polymorphisms of SLC2A2, SLC2A5, and KHK on metabolic phenotypes in hypertensive individuals. PLoS One. 2013;8(1):e52062. doi:10.1371/journal.pone.0052062

598. Kim WT, Lee SR, Roh YG, Kim SI, Choi YH, Mun MH, Jeong MS, Koh SS, Leem SH. Characterization of VNTRs Within the Entire Region of SLC6A3 and Its Association with Hypertension. DNA Cell Biol. 2017;36(3):227–236. doi:10.1089/dna.2016.3448

599. Mahallawi WH, Suliman BA. TLR8 is highly conserved among the Saudi population and its mutations have no effect on the severity of COVID-19 symptoms. Am J Clin Exp Immunol. 2021;10(3):71–76.

600. Vanderheiden A, Thomas J, Soung AL, Davis-Gardner ME, Floyd K, Jin F, Cowan DA, Pellegrini K, Shi PY, Grakoui A, et al. CCR2 Signaling Restricts SARS-CoV-2 Infection. mBio. 2021;12(6):e0274921. doi:10.1128/mBio.02749-21

601. Moustaqil M, Ollivier E, Chiu HP, Van Tol S, Rudolffi-Soto P, Stevens C, Bhumkar A, Hunter DJB, Freiberg AN, Jacques D, et al. SARS-CoV-2 proteases PLpro and 3CLpro cleave IRF3 and critical modulators of inflammatory pathways (NLRP12 and TAB1): implications for disease presentation across species. Emerg Microbes Infect. 2021;10(1):178–195. doi:10.1080/22221751.2020.1870414

602. Li L, Huang M, Shen J, Wang Y, Wang R, Yuan C, Jiang L, Huang M. Serum Levels of Soluble Platelet Endothelial Cell Adhesion Molecule 1 in COVID-19 Patients Are Associated With Disease Severity. J Infect Dis. 2021;223(1):178–179. doi:10.1093/infdis/jiaa642

603. McGee MC, August A, Huang W. BTK/ITK dual inhibitors: Modulating immunopathology and lymphopenia for COVID-19 therapy. J Leukoc Biol. 2021;109(1):49–53. doi:10.1002/JLB.5COVR0620-306R

604. Spoerl S, Kremer AN, Aigner M, Eisenhauer N, Koch P, Meretuk L, Löffler P, Tenbusch M, Maier C, Überla K, et al. Upregulation of CCR4 in activated CD8+ T cells indicates enhanced lung homing in patients with severe acute SARS-CoV-2 infection. Eur J Immunol. 2021;51(6):1436–1448. doi:10.1002/eji.202049135

605. Seale LA, Torres DJ, Berry MJ, Pitts MW. A role for selenium-dependent GPX1 in SARS-CoV-2 virulence. Am J Clin Nutr. 2020;112(2):447–448. doi:10.1093/ajcn/nqaa177

606. Jeong SK, Heo YK, Jeong JH, Ham SJ, Yum JS, Ahn BC, Song CS, Chun EY. COVID-19 Subunit Vaccine with a Combination of TLR1/2 and TLR3 Agonists Induces Robust and Protective Immunity. Vaccines (Basel). 2021;9(9):957. doi:10.3390/vaccines9090957

607. Pérez-García F, Martin-Vicente M, Rojas-García RL, Castilla-García L, Muñoz-Gomez MJ, Hervás Fernández I, González Ventosa V, Vidal-Alcántara EJ, Cuadros-González J, Bermejo-Martin JF, et al. High SARS-CoV-2 viral load and low CCL5 expression levels in the upper respiratory tract are associated with COVID-19 severity. J Infect Dis. 2021;jiab604. doi:10.1093/infdis/jiab604

608. Murch SH. Common determinants of severe Covid-19 infection are explicable by SARS-CoV-2 secreted glycoprotein interaction with the CD33-related Siglecs, Siglec-3 and Siglec-5/14. Med Hypotheses. 2020;144:110168. doi:10.1016/j.mehy.2020.110168

609. Virant-Klun I, Strle F. Human Oocytes Express Both ACE2 and BSG Genes and Corresponding Proteins: Is SARS-CoV-2 Infection Possible?. Stem Cell Rev Rep. 2021;17(1):278–284. doi:10.1007/s12015-020-10101-x

610. Bonyek-Silva I, Machado AFA, Cerqueira-Silva T, Nunes S, Silva Cruz MR, Silva J, Santos RL, Barral A, Oliveira PRS, Khouri R, et al. LTB4-Driven Inflammation and Increased Expression of ALOX5/ACE2 During Severe COVID-19 in Individuals With Diabetes. Diabetes. 2021;70(9):2120–2130. doi:10.2337/db20-1260

611. Neri T, Nieri D, Celi A. P-selectin blockade in COVID-19-related ARDS. Am J Physiol Lung Cell Mol Physiol. 2020;318(6):L1237–L1238. doi:10.1152/ajplung.00202.2020

612. Delaveris CS, Wilk AJ, Riley NM, Stark JC, Yang SS, Rogers AJ, Ranganath T, Nadeau KC. Synthetic Siglec-9 Agonists Inhibit Neutrophil Activation Associated with COVID-19. Preprint. ChemRxiv. 2020;10.26434/chemrxiv.13378148.v1. doi:10.26434/chemrxiv.13378148

613. Brandão SCS, Ramos JOX, Dompieri LT, Godoi ETAM, Figueiredo JL, Sarinho ESC, Chelvanambi S, Aikawa M. Is Toll-like receptor 4 involved in the severity of COVID-19 pathology in patients with cardiometabolic comorbidities?. Cytokine Growth Factor Rev. 2021;58:102–110. doi:10.1016/j.cytogfr.2020.09.002

614. Solerte SB, Di Sabatino A, Galli M, Fiorina P. Dipeptidyl peptidase-4 (DPP4) inhibition in COVID-19. Acta Diabetol. 2020;57(7):779–783. doi:10.1007/s00592-020-01539-z

615. Duncan-Lowey JK, Chowdhary V, Dunne DW. Coronavirus Disease 2019 in a Patient With a Systemic Autoinflammatory Syndrome due to an NLRC4 Inflammasomopathy. Open Forum Infect Dis. 2021;8(8):ofab362. doi:10.1093/ofid/ofab362

616. Zheng M, Karki R, Williams EP, Yang D, Fitzpatrick E, Vogel P, Jonsson CB, Kanneganti TD. TLR2 senses the SARS-CoV-2 envelope protein to produce inflammatory cytokines. Nat Immunol. 2021;22(7):829–838. doi:10.1038/s41590-021-00937-x

617. Jamaly S, Tsokos MG, Bhargava R, Brook OR, Hecht JL, Abdi R, Moulton VR, Satyam A, Tsokos GC. Complement activation and increased expression of Syk, mucin-1 and CaMK4 in kidneys of patients with COVID-19. Clin Immunol. 2021;229:108795. doi:10.1016/j.clim.2021.108795

618. Nassir N, Tambi R, Bankapur A, Al Heialy S, Karuvantevida N, Khansaheb HH, Zehra B, Begum G, Hameid RA, Ahmed A, et al. Single-cell transcriptome identifies FCGR3B upregulated subtype of alveolar macrophages in patients with critical COVID-19. iScience. 2021;24(9):103030. doi:10.1016/j.isci.2021.103030

619. Liang Y, Li H, Li J, Yang ZN, Li JL, Zheng HW, Chen YL, Shi HJ, Guo L, Liu LD. Role of neutrophil chemoattractant CXCL5 in SARS-CoV-2 infection-induced lung inflammatory innate immune response in an in vivo hACE2 transfection mouse model. Zool Res. 2020;41(6):621–631. doi:10.24272/j.issn.2095-8137.2020.118

620. Vietzen H, Zoufaly A, Traugott M, Aberle J, Aberle SW, Puchhammer-Stöckl E. Deletion of the NKG2C receptor encoding KLRC2 gene and HLA-E variants are risk factors for severe COVID-19. Genet Med. 2021;23(5):963–967. doi:10.1038/s41436-020-01077-7

621. Kisserli A, Schneider N, Audonnet S, Tabary T, Goury A, Cousson J, Mahmoudi R, Bani-Sadr F, Kanagaratnam L, Jolly D, et al. Acquired decrease of the C3b/C4b receptor (CR1, CD35) and increased C4d deposits on erythrocytes from ICU COVID-19 patients. Immunobiology. 2021;226(3):152093. doi:10.1016/j.imbio.2021.152093

622. Ercan H, Schrottmaier WC, Pirabe A, Schmuckenschlager A, Pereyra D, Santol J, Pawelka E, Traugott MT, Schörgenhofer C, Seitz T, et al. Platelet Phenotype Analysis of COVID-19 Patients Reveals Progressive Changes in the Activation of Integrin αIIbβ3, F13A1, the SARS-CoV-2 Target EIF4A1 and Annexin A5. Front Cardiovasc Med. 2021;8:779073. doi:10.3389/fcvm.2021.779073

623. Oliviero A, de Castro F, Coperchini F, Chiovato L, Rotondi M. COVID-19 Pulmonary and Olfactory Dysfunctions: Is the Chemokine CXCL10 the Common Denominator?. Neuroscientist. 2021;27(3):214–221. doi:10.1177/1073858420939033

624. Mpekoulis G, Frakolaki E, Taka S, Ioannidis A, Vassiliou AG, Kalliampakou KI, Patas K, Karakasiliotis I, Aidinis V, Chatzipanagiotou S, et al. Alteration of L-Dopa decarboxylase expression in SARS-CoV-2 infection and its association with the interferon-inducible ACE2 isoform. PLoS One. 2021;16(6):e0253458. doi:10.1371/journal.pone.0253458

625. Acar E, Demir A, Yıldırım B, Kaya MG, Gökçek K. The role of hemogram parameters and C-reactive protein in predicting mortality in COVID-19 infection. Int J Clin Pract. 2021;75(7):e14256. doi:10.1111/ijcp.14256

626. Maione F, Casillo GM, Raucci F, Salvatore C, Ambrosini G, Costa L, Scarpa R, Caso F, Bucci M. Interleukin-17A (IL-17A): A silent amplifier of COVID-19. Biomed Pharmacother. 2021;142:111980. doi:10.1016/j.biopha.2021.111980

627. Ehsani S. COVID-19 and iron dysregulation: distant sequence similarity between hepcidin and the novel coronavirus spike glycoprotein. Biol Direct. 2020;15(1):19. doi:10.1186/s13062-020-00275-2

628. Callahan V, Hawks S, Crawford MA, Lehman CW, Morrison HA, Ivester HM, Akhrymuk I, Boghdeh N, Flor R, Finkielstein CV, et al. The Pro-Inflammatory Chemokines CXCL9, CXCL10 and CXCL11 Are Upregulated Following SARS-CoV-2 Infection in an AKT-Dependent Manner. Viruses. 2021;13(6):1062. doi:10.3390/v13061062

629. Ahmad R, Kochumon S, Thomas R, Atizado V, Sindhu S. Increased adipose tissue expression of TLR8 in obese individuals with or without type-2 diabetes: significance in metabolic inflammation. J Inflamm (Lond). 2016;13:38. doi:10.1186/s12950-016-0147-y

630. Sireesh D, Dhamodharan U, Ezhilarasi K, Vijay V, Ramkumar KM. Association of NF-E2 Related Factor 2 (Nrf2) and inflammatory cytokines in recent onset Type 2 Diabetes Mellitus. Sci Rep. 2018;8(1):5126. doi:10.1038/s41598-018-22913-6

631. Di Prospero NA, Artis E, Andrade-Gordon P, Johnson DL, Vaccaro N, Xi L, Rothenberg P. CCR2 antagonism in patients with type 2 diabetes mellitus: a randomized, placebo-controlled study. Diabetes Obes Metab. 2014;16(11):1055–1064. doi:10.1111/dom.12309

632. Anjosa ZP, Santos MM, Rodrigues NJ, Lacerda GA, Araujo J, Silva JA, Tavares NA, Guimarães RL, Crovella S, Brandão LA. Polymorphism in ficolin-1 (FCN1) gene is associated with an earlier onset of type 1 diabetes mellitus in children and adolescents from northeast Brazil. J Genet. 2016;95(4):1031–1034. doi:10.1007/s12041-016-0719-x

633. Aparicio JM, Wakisaka A, Takada A, Matsuura N, Yoshiki T. Non-HLA genetic factors and insulin dependent diabetes mellitus in the Japanese: TCRA, TCRB and TCRG, INS, THY1, CD3D and ETS1. Dis Markers. 1990;8(5):283–294.

634. Rau H, Usadel KH, Ommert S, Badenhoop K. PVU II polymorphism of LST-1 (leucocyte specific transcript-1) in type I diabetes mellitus, Graves’ disease and healthy controls. Eur J Immunogenet. 1995;22(3):277–282. doi:10.1111/j.1744-313x.1995.tb00242.x

635. Kim SH, Cleary MM, Fox HS, Chantry D, Sarvetnick N. CCR4-bearing T cells participate in autoimmune diabetes. J Clin Invest. 2002;110(11):1675–1686. doi:10.1172/JCI15547

636. Thude H, Rosenhahn S, Hunger-Dathe W, Müller UA, Barz D. A transmembrane protein-tyrosine phosphatase receptor type C (CD45) exon A point mutation (77 C to G) is not associated with the development of type 1 diabetes mellitus in a German population. Eur J Immunogenet. 2004;31(6):245–247. doi:10.1111/j.1365-2370.2004.00479.x

637. Eldor R, Klieger Y, Sade-Feldman M, Vaknin I, Varfolomeev I, Fuchs C, Baniyash M. CD247, a novel T cell-derived diagnostic and prognostic biomarker for detecting disease progression and severity in patients with type 2 diabetes. Diabetes Care. 2015;38(1):113–118. doi:10.2337/dc14-1544

638. Ferjeni Z, Bouzid D, Fourati H, Stayoussef M, Abida O, Kammoun T, Hachicha M, Penha-Gonçalves C, Masmoudi H. Association of TCR/CD3, PTPN22, CD28 and ZAP70 gene polymorphisms with type 1 diabetes risk in Tunisian population: family based association study. Immunol Lett. 2015;163(1):1–7. doi:10.1016/j.imlet.2014.11.005

639. Zurawek M, Fichna M, Fichna P, Czainska M, Rozwadowska N. Upregulation of FOXO3 in New-Onset Type 1 Diabetes Mellitus. J Immunol Res. 2020;2020:9484015. doi:10.1155/2020/9484015

640. Liu D, Liu L, Hu Z, Song Z, Wang Y, Chen Z. Evaluation of the oxidative stress-related genes ALOX5, ALOX5AP, GPX1, GPX3 and MPO for contribution to the risk of type 2 diabetes mellitus in the Han Chinese population. Diab Vasc Dis Res. 2018;15(4):336–339. doi:10.1177/1479164118755044

641. Pacifici F, Arriga R, Sorice GP, Capuani B, Scioli MG, Pastore D, Donadel G, Bellia A, Caratelli S, Coppola A, et al. Peroxiredoxin 6, a novel player in the pathogenesis of diabetes. Diabetes. 2014;63(10):3210–3220. doi:10.2337/db14-0144

642. Huang MH, Liu YF, Nfor ON, Hsu SY, Lin WY, Chang YS, Liaw YP. Interactive Association Between Intronic Polymorphism (rs10506151) of the LRRK2 Gene and Type 2 Diabetes on Neurodegenerative Diseases. Pharmgenomics Pers Med. 2021;14:839–847. doi:10.2147/PGPM.S316158

643. Zhang J, Luo J, Liu F, Wu D, Zhong Q, Zeng L, Wu Z, Zhao T, Wu L, Hao H. Diabetes mellitus potentiates diffuse large BLcell lymphoma via high levels of CCL5. Mol Med Rep. 2014;10(3):1231–1236. doi:10.3892/mmr.2014.2341

644. Pawłowicz M, Filipów R, Krzykowski G, Stanisławska-Sachadyn A, Morzuch L, Kulczycka J, Balcerska A, Limon J. Coincidence of PTPN22 c.1858CC and FCRL3 −169CC genotypes as a biomarker of preserved residual β-cell function in children with type 1 diabetes. Pediatr Diabetes. 2017;18(8):696–705. doi:10.1111/pedi.12429

645. Ramos-Lopez E, Ghebru S, Van Autreve J, Aminkeng F, Herwig J, Seifried E, Seidl C, Van der Auwera B, Badenhoop K. Neither an intronic CA repeat within the CD48 gene nor the HERV-K18 polymorphisms are associated with type 1 diabetes. Tissue Antigens. 2006;68(2):147–152. doi:10.1111/j.1399-0039.2006.00637.x

646. Abu El-Ella SS, Khattab ESAEH, El-Mekkawy MS, El-Shamy AA. CD226 gene polymorphism (rs763361 C>T) is associated with susceptibility to type 1 diabetes mellitus among Egyptian children. Arch Pediatr. 2018;25(6):378–382. doi:10.1016/j.arcped.2018.06.009

647. Mi QS, Ly D, Zucker P, McGarry M, Delovitch TL. Interleukin-4 but not interleukin-10 protects against spontaneous and recurrent type 1 diabetes by activated CD1d-restricted invariant natural killer T-cells. Diabetes. 2004;53(5):1303–1310. doi:10.2337/diabetes.53.5.1303

648. Swafford AD, Howson JM, Davison LJ, Wallace C, Smyth DJ, Schuilenburg H, Maisuria-Armer M, Mistry T, Lenardo MJ, Todd JA. An allele of IKZF1 (Ikaros) conferring susceptibility to childhood acute lymphoblastic leukemia protects against type 1 diabetes. Diabetes. 2011;60(3):1041–1044. doi:10.2337/db10-0446

649. Doody NE, Dowejko MM, Akam EC, Cox NJ, Bhatti JS, Singh P, Mastana SS. The Role of TLR4, TNF-α and IL-1β in Type 2 Diabetes Mellitus Development within a North Indian Population. Ann Hum Genet. 2017;81(4):141–146. doi:10.1111/ahg.12197

650. Barchetta I, Ciccarelli G, Barone E, Cimini FA, Ceccarelli V, Bertoccini L, Sentinelli F, Tramutola A, Del Ben M, Angelico F, et al. Greater circulating DPP4 activity is associated with impaired flow-mediated dilatation in adults with type 2 diabetes mellitus. Nutr Metab Cardiovasc Dis. 2019;29(10):1087–1094. doi:10.1016/j.numecd.2019.07.010

651. Xu L, Sun X, Xia Y, Luo S, Lin J, Xiao Y, Liu Y, Wang Y, Huang G, Li X, et al. Polymorphisms of the NLRC4 Gene are Associated with the Onset Age, Positive Rate of GADA and 2-h Postprandial C-Peptide in Patients with Type 1 Diabetes. Diabetes Metab Syndr Obes. 2020;13:811–818. doi:10.2147/DMSO.S244882

652. Li JY, Tao F, Wu XX, Tan YZ, He L, Lu H. Common RASGRP1 Gene Variants That Confer Risk of Type 2 Diabetes. Genet Test Mol Biomarkers. 2015;19(8):439–443. doi:10.1089/gtmb.2015.0005

653. Demirci M, Bahar Tokman H, Taner Z, Keskin FE, Çağatay P, Ozturk Bakar Y, Özyazar M, Kiraz N, Kocazeybek BS. Bacteroidetes and Firmicutes levels in gut microbiota and effects of hosts TLR2/TLR4 gene expression levels in adult type 1 diabetes patients in Istanbul, Turkey. J Diabetes Complications. 2020;34(2):107449. doi:10.1016/j.jdiacomp.2019.107449

654. Hasani Ranjbar S, Amiri P, Zineh I, Langaee TY, Namakchian M, Heshmet R, Sajadi M, Mirzaee M, Rezazadeh E, Balaei P, et al. CXCL5 gene polymorphism association with diabetes mellitus. Mol Diagn Ther. 2008;12(6):391–394. doi:10.1007/BF03256304

655. Puchałowicz K, Rać ME. The Multifunctionality of CD36 in Diabetes Mellitus and Its Complications-Update in Pathogenesis, Treatment and Monitoring. Cells. 2020;9(8):1877. doi:10.3390/cells9081877

656. Fejes Z, Póliska S, Czimmerer Z, Káplár M, Penyige A, Gál Szabó G, Beke Debreceni I, Kunapuli SP, Kappelmayer J, Nagy B Jr.Hyperglycaemia suppresses microRNA expression in platelets to increase P2RY12 and SELP levels in type 2 diabetes mellitus. Thromb Haemost. 2017;117(3):529–542. doi:10.1160/TH16-04-0322

657. Kawabata Y, Nishida N, Awata T, Kawasaki E, Imagawa A, Shimada A, Osawa H, Tanaka S, Takahashi K, Nagata M, et al. Genome-Wide Association Study Confirming a Strong Effect of HLA and Identifying Variants in CSAD/lnc-ITGB7-1 on Chromosome 12q13.13 Associated With Susceptibility to Fulminant Type 1 Diabetes. Diabetes. 2019;68(3):665–675. doi:10.2337/db18-0314

658. Haseda F, Imagawa A, Nishikawa H, Mitsui S, Tsutsumi C, Fujisawa R, Sano H, Murase-Mishiba Y, Terasaki J, Sakaguchi S, et al. Antibody to CMRF35-Like Molecule 2, CD300e A Novel Biomarker Detected in Patients with Fulminant Type 1 Diabetes. PLoS One. 2016;11(8):e0160576. doi:10.1371/journal.pone.0160576

659. Fernández-Real JM, Mercader JM, Ortega FJ, Moreno-Navarrete JM, López-Romero P, Ricart W. Transferrin receptor-1 gene polymorphisms are associated with type 2 diabetes. Eur J Clin Invest. 2010;40(7):600–607. doi:10.1111/j.1365-2362.2010.02306.x

660. Peters KE, Beilby J, Cadby G, Warrington NM, Bruce DG, Davis WA, Davis TM, Wiltshire S, Knuiman M, McQuillan BM, et al. A comprehensive investigation of variants in genes encoding adiponectin (ADIPOQ) and its receptors (ADIPOR1/R2), and their association with serum adiponectin, type 2 diabetes, insulin resistance and the metabolic syndrome. BMC Med Genet. 2013;14:15. doi:10.1186/1471-2350-14-15

661. Sun L, Zhang X, Wang T, Chen M, Qiao H. Association of ANK1 variants with new-onset type 2 diabetes in a Han Chinese population from northeast China. Exp Ther Med. 2017;14(4):3184–3190. doi:10.3892/etm.2017.4866

662. Mao R, Yang F, Zhang Y, Liu H, Guo P, Liu Y, Zhang T. High expression of CD52 in adipocytes: a potential therapeutic target for obesity with type 2 diabetes. Aging (Albany NY). 2021;13(8):11043–11060. doi:10.18632/aging.202714

663. Sano R, Miki T, Suzuki Y, Shimada F, Taira M, Kanatsuka A, Makino H, Hashimoto N, Saito Y. Analysis of the insulin-sensitive phosphodiesterase 3B gene in type 2 diabetes. Diabetes Res Clin Pract. 2001;54(2):79–88. doi:10.1016/s0168-8227(01)00287-x

664. Szabó E, Kulin A, Mózner O, Korányi L, Literáti-Nagy B, Vitai M, Cserepes J, Sarkadi B, Várady G. Potential role of the ABCG2-Q141K polymorphism in type 2 diabetes. PLoS One. 2021;16(12):e0260957. doi:10.1371/journal.pone.0260957

665. Shalaby D, Saied M, Khater D, Abou Zeid A. The Expression of Activating Receptor Gene of Natural Killer Cells (KLRC3) in Patients with LType 1 Diabetes Mellitus (T1DM). Oman Med J. 2017;32(4):316–321. doi:10.5001/omj.2017.60

666. Yu Y, Mingjiao W, Yang X, Sui M, Zhang T, Liang J, Gu X, Wang X. Association between DNA methylation of SORL1 5’-flanking region and mild cognitive impairment in type 2 diabetes mellitus. Ann Endocrinol (Paris). 2016;77(6):625–632. doi:10.1016/j.ando.2016.02.008

667. Xiang RL, Huang Y, Zhang Y, Cong X, Zhang ZJ, Wu LL, Yu GY. Type 2 diabetes-induced hyposalivation of the submandibular gland through PINK1/Parkin-mediated mitophagy. J Cell Physiol. 2020;235(1):232–244. doi:10.1002/jcp.28962

668. Abdelmajed SS, El-Dessouky MA, SalahElDin DS, Hassan NA, Zaki ME, Ismail S. Assessing the association of rs7574865 STAT4 gene variant and type 1 diabetes mellitus among Egyptian patients. J Genet Eng Biotechnol. 2021;19(1):112. doi:10.1186/s43141-021-00214-2

669. Antonelli A, Ferrari SM, Corrado A, Ferrannini E, Fallahi P. CXCR3, CXCL10 and type 1 diabetes. Cytokine Growth Factor Rev. 2014;25(1):57–65. doi:10.1016/j.cytogfr.2014.01.006

670. Maeda S, Tsukada S, Kanazawa A, Sekine A, Tsunoda T, Koya D, Maegawa H, Kashiwagi A, Babazono T, Matsuda M, et al. Genetic variations in the gene encoding TFAP2B are associated with type 2 diabetes mellitus. J Hum Genet. 2005;50(6):283–292. doi:10.1007/s10038-005-0253-9

671. Lee CC, Adler AI, Sandhu MS, Sharp SJ, Forouhi NG, Erqou S, Luben R, Bingham S, Khaw KT, Wareham NJ. Association of C-reactive protein with type 2 diabetes: prospective analysis and meta-analysis. Diabetologia. 2009;52(6):1040–1047. doi:10.1007/s00125-009-1338-3

672. Kochetova OV, Avzaletdinova DS, Korytina GF, Morugova TV, Mustafina OE. The association between eating behavior and polymorphisms in GRIN2B, GRIK3, GRIA1 and GRIN1 genes in people with type 2 diabetes mellitus. Mol Biol Rep. 2020;47(3):2035–2046. doi:10.1007/s11033-020-05304-x

673. Wang W, Yan X, Lin Y, Ge H, Tan Q. Wnt7a promotes wound healing by regulation of angiogenesis and inflammation: Issues on diabetes and obesity. J Dermatol Sci. 2018;S0923-1811(18)30103–8. doi:10.1016/j.jdermsci.2018.02.007

674. Tripaldi R, Lanuti P, Simeone PG, Liani R, Bologna G, Ciotti S, Simeone P, Di Castelnuovo A, Marchisio M, Cipollone F, et al. Endogenous PCSK9 may influence circulating CD45neg/CD34bright and CD45neg/CD34bright/CD146neg cells in patients with type 2 diabetes mellitus. Sci Rep. 2021;11(1):9659. doi:10.1038/s41598-021-88941-x

675. Izumi M, Zhang BX, Dean DD, Lin AL, Saunders MJ, Hazuda HP, Yeh CK. Secretion of salivary statherin is compromised in uncontrolled diabetic patients. BBA Clin. 2015;3:135–140. doi:10.1016/j.bbacli.2015.01.002

676. Pehlić M, Vrkić D, Skrabić V, Jerončić A, Stipančić G, Urojić AŠ, Marjanac I, Jakšić J, Kačić Z, Boraska V, et al. IL12RB2 gene is associated with the age of type 1 diabetes onset in Croatian family Trios. PLoS One. 2012;7(11):e49133. doi:10.1371/journal.pone.0049133

677. Bursova S, Dubovy P, Vlckova-Moravcova E, Nemec M, Klusakova I, Belobradkova J, Bednarik J. Expression of growth-associated protein 43 in the skin nerve fibers of patients with type 2 diabetes mellitus. J Neurol Sci. 2012;315(1-2):60–63. doi:10.1016/j.jns.2011.11.038

678. Haghvirdizadeh P, Mohamed Z, Abdullah NA, Haghvirdizadeh P, Haerian MS, Haerian BS. KCNJ11: Genetic Polymorphisms and Risk of Diabetes Mellitus. J Diabetes Res. 2015;2015:908152. doi:10.1155/2015/908152

679. Igoillo-Esteve M, Marselli L, Cunha DA, Ladrière L, Ortis F, Grieco FA, Dotta F, Weir GC, Marchetti P, Eizirik DL, et al. Palmitate induces a pro-inflammatory response in human pancreatic islets that mimics CCL2 expression by beta cells in type 2 diabetes. Diabetologia. 2010;53(7):1395–1405. doi:10.1007/s00125-010-1707-y

680. Xiu L, Lin M, Liu W, Kong D, Liu Z, Zhang Y, Ouyang P, Liang Y, Zhong S, Chen C, et al. Association of DRD3, COMT, and SLC6A4 Gene Polymorphisms with Type 2 Diabetes in Southern Chinese: A Hospital-Based Case-Control Study. Diabetes Technol Ther. 2015;17(8):580–586. doi:10.1089/dia.2014.0344

681. Lorenzo PI, Cobo-Vuilleumier N, Gauthier BR. Therapeutic potential of pancreatic PAX4-regulated pathways in treating diabetes mellitus. Curr Opin Pharmacol. 2018;43:1–10. doi:10.1016/j.coph.2018.07.004

682. Abed E, Jarrar Y, Alhawari H, Abdullah S, Zihlif M. How the cytochrome 7a1 (CYP7A1) and ATP-binding cassette G8 (ABCG8) genetic variants affect atorvastatin response among type 2 diabetic patients attending the University of Jordan Hospital. Int J Clin Pharmacol Ther. 2021;59(2):99–108. doi:10.5414/CP203779

683. Qiu AW, Cao X, Zhang WW, Liu QH. IL-17A is involved in diabetic inflammatory pathogenesis by its receptor IL-17RA. Exp Biol Med (Maywood). 2021;246(1):57–65. doi:10.1177/1535370220956943

684. Tabassum R, Mahajan A, Chauhan G, Dwivedi OP, Ghosh S, Tandon N, Bharadwaj D. Evaluation of DOK5 as a susceptibility gene for type 2 diabetes and obesity in North Indian population. BMC Med Genet. 2010;11:35.doi:10.1186/1471-2350-11-35

685. Rees SD, Britten AC, Bellary S, O’Hare JP, Kumar S, Barnett AH, Kelly MA. The promoter polymorphism −232C/G of the PCK1 gene is associated with type 2 diabetes in a UK-resident South Asian population. BMC Med Genet. 2009;10:83. doi:10.1186/1471-2350-10-83

686. Mancuso E, Spiga R, Rubino M, Averta C, Rotundo S, Segura-Garcìa C, Mannino GC, Sesti G, Andreozzi F. Effects of Alpha-2-HS-glycoprotein on cognitive and emotional assessment in prediabetic and diabetic subjects. J Affect Disord. 2021;282:700–706. doi:10.1016/j.jad.2020.12.135

687. Laukkanen O, Lindström J, Eriksson J, Valle TT, Hämäläinen H, Ilanne-Parikka P, Keinänen-Kiukaanniemi S, Tuomilehto J, Uusitupa M, Laakso M. Polymorphisms in the SLC2A2 (GLUT2) gene are associated with the conversion from impaired glucose tolerance to type 2 diabetes: the Finnish Diabetes Prevention Study. Diabetes. 2005;54(7):2256–2260. doi:10.2337/diabetes.54.7.2256

688. Kang J, Guan RC, Zhao Y, Chen Y. Obesity-related loci in TMEM18, CDKAL1 and FAIM2 are associated with obesity and type 2 diabetes in Chinese Han patients. BMC Med Genet. 2020;21(1):65. doi:10.1186/s12881-020-00999-y

689. Wu HX, Liu JY, Yan DW, Li L, Zhuang XH, Li HY, Zhou ZG, Zhou HD. Atypical juvenile hereditary hemochromatosis onset with positive pancreatic islet autoantibodies diabetes caused by novel mutations in HAMP and overall clinical management. Mol Genet Genomic Med. 2020;8(12):e1522. doi:10.1002/mgg3.1522

690. Licito A, Marotta G, Battaglia M, Benincasa G, Mentone L, Grillo MR, De Lucia V, Leonardi G, Bignucolo A, Comello F, et al. Assessment of pharmacogenomic SLCO1B1 assay for prediction of neuromuscular pain in type 2 diabetes mellitus and cardiovascular patients: preliminary results. Eur Rev Med Pharmacol Sci. 2020;24(1):469–477. doi:10.26355/eurrev_202001_19948

691. Salek Maghsoudi A, Hassani S, Rezaei Akmal M, Ganjali MR, Mirnia K, Norouzi P, Abdollahi M. An Electrochemical Aptasensor Platform Based on Flower-Like Gold Microstructure-Modified Screen-Printed Carbon Electrode for Detection of Serpin A12 as a Type 2 Diabetes Biomarker. Int J Nanomedicine. 2020;15:2219–2230. doi:10.2147/IJN.S244315

692. Porta M, Toppila I, Sandholm N, Hosseini SM, Forsblom C, Hietala K, Borio L, Harjutsalo V, Klein BE, Klein R, et al. Variation in SLC19A3 and Protection From Microvascular Damage in Type 1 Diabetes. Diabetes. 2016;65(4):1022–1030. doi:10.2337/db15-1247

693. She S, Wu X, Zheng D, Pei X, Ma J, Sun Y, Zhou J, Nong L, Guo C, Lv P, et al. PSMP/MSMP promotes hepatic fibrosis through CCR2 and represents a novel therapeutic target. J Hepatol. 2020;72(3):506–518. doi:10.1016/j.jhep.2019.09.033

694. Malik G, Wilting J, Hess CF, Ramadori G, Malik IA. PECAM-1 modulates liver damage induced by synergistic effects of TNF-α and irradiation. J Cell Mol Med. 2019;23(5):3336–3344. doi:10.1111/jcmm.14224

695. Ma N, Li S, Lin C, Cheng X, Meng Z. Mesenchymal stem cell conditioned medium attenuates oxidative stress injury in hepatocytes partly by regulating the miR-486-5p/PIM1 axis and the TGF-β/Smad pathway. Bioengineered. 2021;12(1):6434–6447. doi:10.1080/21655979.2021.1972196

696. Sutti S, Bruzzì S, Heymann F, Liepelt A, Krenkel O, Toscani A, Ramavath NN, Cotella D, Albano E, Tacke F. CX3CR1 Mediates the Development of Monocyte-Derived Dendritic Cells during Hepatic Inflammation. Cells. 2019;8(9):1099.. doi:10.3390/cells8091099

697. Piccolo P, Ferriero R, Barbato A, Attanasio S, Monti M, Perna C, Borel F, Annunziata P, Carissimo A, De Cegli R, et al. Up-regulation of miR-34b/c by JNK and FOXO3 protects from liver fibrosis. Proc Natl Acad Sci U S A. 2021;118(10):e2025242118. doi:10.1073/pnas.2025242118

698. Yan Y, Jiang W, Tan Y, Zou S, Zhang H, Mao F, Gong A, Qian H, Xu W. hucMSC Exosome-Derived GPX1 Is Required for the Recovery of Hepatic Oxidant Injury. Mol Ther. 2017;25(2):465–479. doi:10.1016/j.ymthe.2016.11.019

699. Xiao P, Li M, Zhou M, Zhao X, Wang C, Qiu J, Fang Q, Jiang H, Dong H, Zhou R. TTP protects against acute liver failure by regulating CCL2 and CCL5 through m6A RNA methylation. JCI Insight. 2021;6(23):e149276. doi:10.1172/jci.insight.149276

700. Selzner N, Liu H, Boehnert MU, Adeyi OA, Shalev I, Bartczak AM, Xue-Zhong M, Manuel J, Rotstein OD, McGilvray ID, et al. FGL2/fibroleukin mediates hepatic reperfusion injury by induction of sinusoidal endothelial cell and hepatocyte apoptosis in mice. J Hepatol. 2012;56(1):153–159. doi:10.1016/j.jhep.2011.05.033

701. Ito T, Ishigami M, Matsushita Y, Hirata M, Matsubara K, Ishikawa T, Hibi H, Ueda M, Hirooka Y, Goto H, et al. Secreted Ectodomain of SIGLEC-9 and MCP-1 Synergistically Improve Acute Liver Failure in Rats by Altering Macrophage Polarity. Sci Rep. 2017;7:44043. doi:10.1038/srep44043

702. Zhang CC, Niu F. LncRNA NEAT1 promotes inflammatory response in sepsis-induced liver injury via the Let-7a/TLR4 axis. Int Immunopharmacol. 2019;75:105731. doi:10.1016/j.intimp.2019.105731

703. Wang XM, Holz LE, Chowdhury S, Cordoba SP, Evans KA, Gall MG, Vieira de Ribeiro AJ, Zheng YZ, Levy MT, Yu DM, et al. The pro-fibrotic role of dipeptidyl peptidase 4 in carbon tetrachloride-induced experimental liver injury. Immunol Cell Biol. 2017;95(5):443–453. doi:10.1038/icb.2016.116

704. DeSantis DA, Ko CW, Liu Y, Liu X, Hise AG, Nunez G, Croniger CM. Alcohol-induced liver injury is modulated by Nlrp3 and Nlrc4 inflammasomes in mice. Mediators Inflamm. 2013;2013:751374. doi:10.1155/2013/751374

705. Roh YS, Zhang B, Loomba R, Seki E. TLR2 and TLR9 contribute to alcohol-mediated liver injury through induction of CXCL1 and neutrophil infiltration. Am J Physiol Gastrointest Liver Physiol. 2015;309(1):G30–G41. doi:10.1152/ajpgi.00031.2015

706. Tacke F, Zimmermann HW, Trautwein C, Schnabl B. CXCL5 plasma levels decrease in patients with chronic liver disease. J Gastroenterol Hepatol. 2011;26(3):523–529. doi:10.1111/j.1440-1746.2010.06436.x

707. Fernández-Real JM, Handberg A, Ortega F, Højlund K, Vendrell J, Ricart W. Circulating soluble CD36 is a novel marker of liver injury in subjects with altered glucose tolerance. J Nutr Biochem. 2009;20(6):477–484. doi:10.1016/j.jnutbio.2008.05.009

708. Fu X, Qie J, Fu Q, Chen J, Jin Y, Ding Z. miR-20a-5p/TGFBR2 Axis Affects Pro-inflammatory Macrophages and Aggravates Liver Fibrosis. Front Oncol. 2020;10:107. doi:10.3389/fonc.2020.00107

709. Dibra D, Xia X, Mitra A, Cutrera JJ, Lozano G, Li S. Mutant p53 in concert with an interleukin-27 receptor alpha deficiency causes spontaneous liver inflammation, fibrosis, and steatosis in mice. Hepatology. 2016;63(3):1000–1012. doi:10.1002/hep.28379

710. Goodwin RG, Kell WJ, Laidler P, Long CC, Whatley SD, McKinley M, Badminton MN, Burnett AK, Williams GT, Elder GH. Photosensitivity and acute liver injury in myeloproliferative disorder secondary to late-onset protoporphyria caused by deletion of a ferrochelatase gene in hematopoietic cells. Blood. 2006;107(1):60–62. doi:10.1182/blood-2004-12-4939

711. Malik G, Wilting J, Hess CF, Ramadori G, Malik IA. PECAM-1 modulates liver damage induced by synergistic effects of TNF-α and irradiation. J Cell Mol Med. 2019;23(5):3336–3344. doi:10.1111/jcmm.14224

712. Bukong TN, Maurice SB, Chahal B, Schaeffer DF, Winwood PJ. Versican: a novel modulator of hepatic fibrosis. Lab Invest. 2016;96(3):361–374. doi:10.1038/labinvest.2015.152

713. Sachar M, Ma X. Role of ABCG2 in liver injury associated with erythropoietic protoporphyria. Hepatology. 2016;64(1):305. doi:10.1002/hep.28249

714. Shi Q, Zhao G, Wei S, Guo C, Wu X, Zhao RC, Di G. Pterostilbene alleviates liver ischemia/reperfusion injury via PINK1-mediated mitophagy. J Pharmacol Sci. 2022;148(1):19–30. doi:10.1016/j.jphs.2021.09.005

715. Zhang B, Li MD, Yin R, Liu Y, Yang Y, Mitchell-Richards KA, Nam JH, Li R, Wang L, Iwakiri Y, et al. O-GlcNAc transferase suppresses necroptosis and liver fibrosis. JCI Insight. 2019;4(21):e127709. doi:10.1172/jci.insight.127709

716. Chalin A, Lefevre B, Devisme C, Pronier C, Carrière V, Thibault V, Amiot L, Samson M.Serum CXCL10, CXCL11, CXCL12, and CXCL14 chemokine patterns in patients with acute liver injury. Cytokine. 2018;111:500–504. doi:10.1016/j.cyto.2018.05.029

717. Kawakami R, Matsui M, Konno A, Kaneko R, Shrestha S, Shrestha S, Sunaga H, Hanaoka H, Goto S, Hosojima M, et al. Urinary FABP1 is a biomarker for impaired proximal tubular protein reabsorption and is synergistically enhanced by concurrent liver injury. J Pathol. 2021;255(4):362–373. doi:10.1002/path.5775

718. Zhang N, Hu Y, Ding C, Zeng W, Shan W, Fan H, Zhao Y, Shi X, Gao L, Xu T, et al. Salvianolic acid B protects against chronic alcoholic liver injury via SIRT1-mediated inhibition of CRP and ChREBP in rats. Toxicol Lett. 2017;267:1–10. doi:10.1016/j.toxlet.2016.12.010

719. Li S, Yi Z, Deng M, Scott MJ, Yang C, Li W, Lei Z, Santerre NM, Loughran P, Billiar TR.TSLP protects against liver I/R injury via activation of the PI3K/Akt pathway. JCI Insight. 2019;4(22):e129013. doi:10.1172/jci.insight.129013

720. Mak A, Uetrecht J. Involvement of CCL2/CCR2 macrophage recruitment in amodiaquine-induced liver injury. J Immunotoxicol. 2019;16(1):28–33. doi:10.1080/1547691X.2018.1516014

721. Wu N, Baiocchi L, Zhou T, Kennedy L, Ceci L, Meng F, Sato K, Wu C, Ekser B, Kyritsi K, et al. Functional Role of the Secretin/Secretin Receptor Signaling During Cholestatic Liver Injury. Hepatology. 2020;72(6):2219–2227. doi:10.1002/hep.31484

722. Huang Y, Liu W, Xiao H, Maitikabili A, Lin Q, Wu T, Huang Z, Liu F, Luo Q, Ouyang G. Matricellular protein periostin contributes to hepatic inflammation and fibrosis. Am J Pathol. 2015;185(3):786–797. doi:10.1016/j.ajpath.2014.11.002

723. Tan Z, Qian X, Jiang R, Liu Q, Wang Y, Chen C, Wang X, Ryffel B, Sun B. IL-17A plays a critical role in the pathogenesis of liver fibrosis through hepatic stellate cell activation. J Immunol. 2013;191(4):1835–1844. doi:10.4049/jimmunol.1203013

724. Sun S, Yuan L, An Z, Shi D, Xin J, Jiang J, Ren K, Chen J, Guo B, Zhou X, et al. DLL4 restores damaged liver by enhancing hBMSC differentiation into cholangiocytes. Stem Cell Res. 2020;47:101900. doi:10.1016/j.scr.2020.101900

725. Zhang P, Yue K, Liu X, Yan X, Yang Z, Duan J, Xia C, Xu X, Zhang M, Liang L, et al. Endothelial Notch activation promotes neutrophil transmigration via downregulating endomucin to aggravate hepatic ischemia/reperfusion injury. Sci China Life Sci. 2020;63(3):375–387. doi:10.1007/s11427-019-1596-4

726. Bartneck M, Fech V, Ehling J, Govaere O, Warzecha KT, Hittatiya K, Vucur M, Gautheron J, Luedde T, Trautwein C, et al. Histidine-rich glycoprotein promotes macrophage activation and inflammation in chronic liver disease. Hepatology. 2016;63(4):1310–1324. doi:10.1002/hep.28418

727. Shams S, Mohsin S, Nasir GA, Khan M, Khan SN. Mesenchymal Stem Cells Pretreated with HGF and FGF4 Can Reduce Liver Fibrosis in Mice. Stem Cells Int. 2015;2015:747245. doi:10.1155/2015/747245

728. Mustafa A, Clarke JT. Ornithine transcarbamoylase deficiency presenting with acute liver failure. J Inherit Metab Dis. 2006;29(4):586. doi:10.1007/s10545-006-0303-2

729. Song X, Shen Y, Lao Y, Tao Z, Zeng J, Wang J, Wu H. CXCL9 regulates acetaminophen-induced liver injury via CXCR3. Exp Ther Med. 2019;18(6):4845–4851. doi:10.3892/etm.2019.8122

730. Aslani M, Mortazavi-Jahromi SS, Mirshafiey A. Cytokine storm in the pathophysiology of COVID-19: Possible functional disturbances of miRNAs. Int Immunopharmacol. 2021;101(Pt A):108172. doi:10.1016/j.intimp.2021.108172

731. Jafarzadeh A, Nemati M, Jafarzadeh S. Contribution of STAT3 to the pathogenesis of COVID-19. Microb Pathog. 2021;154:104836. doi:10.1016/j.micpath.2021.104836

732. Cao Z, Yao F, Lang Y, Feng X. Elevated Circulating LINC-P21 Serves as a Diagnostic Biomarker of Type 2 Diabetes Mellitus and Regulates Pancreatic β-cell Function by Sponging miR-766-3p to Upregulate NR3C2. Exp Clin Endocrinol Diabetes. 2020;10.1055/a-1247-4978. doi:10.1055/a-1247-4978

733. Krause C, Sievert H, Geißler C, Grohs M, El Gammal AT, Wolter S, Ohlei O, Kilpert F, Krämer UM, Kasten M, et al. Critical evaluation of the DNA-methylation markers ABCG1 and SREBF1 for Type 2 diabetes stratification. Epigenomics. 2019;11(8):885–897. doi:10.2217/epi-2018-0159

734. Haghnazari L, Sabzi R. Relationship between TP53 and interleukin-6 gene variants and the risk of types 1 and 2 diabetes mellitus development in the Kermanshah province. J Med Life. 2021;14(1):37–44. doi:10.25122/jml-2019-0150

735. Ao H, Liu B, Li H, Lu L. Egr1 mediates retinal vascular dysfunction in diabetes mellitus via promoting p53 transcription. J Cell Mol Med. 2019;23(5):3345–3356. doi:10.1111/jcmm.14225

736. Zhang Y, Lin C, Chen R, Luo L, Huang J, Liu H, Chen W, Xu J, Yu H, Ding Y. Association analysis of SOCS3, JAK2 and STAT3 gene polymorphisms and genetic susceptibility to type 2 diabetes mellitus in Chinese population. Diabetol Metab Syndr. 2022;14(1):4. doi:10.1186/s13098-021-00774-w

737. Luo L, Wang Y, Hu P, Wu J. Long Non-Coding RNA Metastasis Associated Lung Adenocarcinoma Transcript 1 (MALAT1) Promotes Hypertension by Modulating the Hsa-miR-124-3p/Nuclear Receptor Subfamily 3, Group C, Member 2 (NR3C2) and Hsa-miR-135a-5p/NR3C2 Axis. Med Sci Monit. 2020;26:e920478. doi:10.12659/MSM.920478

738. Pravenec M, Kazdova L, Landa V, Zidek V, Mlejnek P, Simakova M, Jansa P, Forejt J, Kren V, Krenova D, et al. Identification of mutated Srebf1 as a QTL influencing risk for hepatic steatosis in the spontaneously hypertensive rat. Hypertension. 2008;51(1):148–153. doi:10.1161/HYPERTENSIONAHA.107.100743

739. Patel M, Predescu D, Tandon R, Bardita C, Pogoriler J, Bhorade S, Wang M, Comhair S, Ryan-Hemnes A, Chen J, et al. A novel p38 mitogen-activated protein kinase/Elk-1 transcription factor-dependent molecular mechanism underlying abnormal endothelial cell proliferation in plexogenic pulmonary arterial hypertension. J Biol Chem. 2013;288(36):25701–25716. doi:10.1074/jbc.M113.502674

740. Satoh T, Wang L, Espinosa-Diez C, Wang B, Hahn SA, Noda K, Rochon ER, Dent MR, Levine AR, Baust JJ, et al. Metabolic Syndrome Mediates ROS-miR-193b-NFYA-Dependent Downregulation of Soluble Guanylate Cyclase and Contributes to Exercise-Induced Pulmonary Hypertension in Heart Failure With Preserved Ejection Fraction. Circulation. 2021;144(8):615–637. doi:10.1161/CIRCULATIONAHA.121.053889

741. Feng Q, Tian T, Liu J, Zhang L, Qi J, Lin X. Deregulation of microRNAL31aL5p is involved in the development of primary hypertension by suppressing apoptosis of pulmonary artery smooth muscle cells via targeting TP53. Int J Mol Med. 2018;42(1):290–298. doi:10.3892/ijmm.2018.3597

742. Lv L, Shen J, Xu J, Wu X, Zeng C, Lin L, Mao W, Wei T. MiR-124-3p reduces angiotensin II-dependent hypertension by down-regulating EGR1. J Hum Hypertens. 2021;35(8):696–708. doi:10.1038/s41371-020-0381-x

743. Cheng G, He L, Zhang Y. LincRNA-Cox2 promotes pulmonary arterial hypertension by regulating the let-7a-mediated STAT3 signaling pathway. Mol Cell Biochem. 2020;475(1-2):239–247. doi:10.1007/s11010-020-03877-6

744. Le Hellard S, Mühleisen TW, Djurovic S, Fernø J, Ouriaghi Z, Mattheisen M, Vasilescu C, Raeder MB, Hansen T, Strohmaier J, et al. Polymorphisms in SREBF1 and SREBF2, two antipsychotic-activated transcription factors controlling cellular lipogenesis, are associated with schizophrenia in German and Scandinavian samples. Mol Psychiatry. 2010;15(5):463–472. doi:10.1038/mp.2008.110

745. Jacobs WB, Kaplan DR, Miller FD. The p53 family in nervous system development and disease. J Neurochem. 2006;97(6):1571–1584. doi:10.1111/j.1471-4159.2006.03980.x

746. Duclot F, Kabbaj M. The Role of Early Growth Response 1 (EGR1) in Brain Plasticity and Neuropsychiatric Disorders. Front Behav Neurosci. 2017;11:35. doi:10.3389/fnbeh.2017.00035

747. Pu Y, Zhao Q, Men X, Jin W, Yang M. MicroRNA-325 facilitates atherosclerosis progression by mediating the SREBF1/LXR axis via KDM1A. Life Sci. 2021;277:119464. doi:10.1016/j.lfs.2021.119464

748. Huang Y, Chen L, Feng Z, Chen W, Yan S, Yang R, Xiao J, Gao J, Zhang D, Ke X. EPC-Derived Exosomal miR-1246 and miR-1290 Regulate Phenotypic Changes of Fibroblasts to Endothelial Cells to Exert Protective Effects on Myocardial Infarction by Targeting ELF5 and SP1. Front Cell Dev Biol. 2021;9:647763. doi:10.3389/fcell.2021.647763

749. Rouhi L, Fan S, Cheedipudi SM, Braza-Boïls A, Molina MS, Yao Y, Robertson MJ, Coarfa C, Gimeno JR, Molina P, et al. The EP300/TP53 pathway, a suppressor of the Hippo and canonical WNT pathways, is activated in human hearts with arrhythmogenic cardiomyopathy in the absence of overt heart failure. Cardiovasc Res. 2021;cvab197. doi:10.1093/cvr/cvab197

750. Fan K, Huang W, Qi H, Song C, He C, Liu Y, Zhang Q, Wang L, Sun H. The Egr-1/miR-15a-5p/GPX4 axis regulates ferroptosis in acute myocardial infarction. Eur J Pharmacol. 2021;909:174403. doi:10.1016/j.ejphar.2021.174403

751. Major JL, Dewan A, Salih M, Leddy JJ, Tuana BS. E2F6 Impairs Glycolysis and Activates BDH1 Expression Prior to Dilated Cardiomyopathy. PLoS One. 2017;12(1):e0170066. doi:10.1371/journal.pone.0170066

752. Bao Q, Zhang B, Suo Y, Liu C, Yang Q, Zhang K, Yuan M, Yuan M, Zhang Y, Li G. Intermittent hypoxia mediated by TSP1 dependent on STAT3 induces cardiac fibroblast activation and cardiac fibrosis. Elife. 2020;9:e49923. doi:10.7554/eLife.49923

753. Tao Z, Jie Y, Mingru Z, Changping G, Fan Y, Haifeng W, Yuelan W. The Elk1/MMP-9 axis regulates E-cadherin and occludin in ventilator-induced lung injury. Respir Res. 2021;22(1):233. doi:10.1186/s12931-021-01829-2

754. Yan C, Chen J, Ding Y, Zhou Z, Li B, Deng C, Yuan D, Zhang Q, Wang X. The Crucial Role of PPARγ-Egr-1-Pro-Inflammatory Mediators Axis in IgG Immune Complex-Induced Acute Lung Injury. Front Immunol. 2021;12:634889. doi:10.3389/fimmu.2021.634889

755. Paris AJ, Hayer KE, Oved JH, Avgousti DC, Toulmin SA, Zepp JA, Zacharias WJ, Katzen JB, Basil MC, Kremp MM, et al. STAT3-BDNF-TrkB signalling promotes alveolar epithelial regeneration after lung injury. Nat Cell Biol. 2020;22(10):1197–1210. doi:10.1038/s41556-020-0569-x

756. Bai Q, Yan H, Sheng Y, Jin Y, Shi L, Ji L, Wang Z.Long-term acetaminophen treatment induced liver fibrosis in mice and the involvement of Egr-1. Toxicology. 2017;382:47–58. doi:10.1016/j.tox.2017.03.008

757. Shin S, Upadhyay N, Greenbaum LE, Kaestner KH. Ablation of Foxl1-Cre-labeled hepatic progenitor cells and their descendants impairs recovery of mice from liver injury. Gastroenterology. 2015;148(1):192–202.e3. doi:10.1053/j.gastro.2014.09.039

758. Chen J, Xia H, Zhang L, Zhang H, Wang D, Tao X. Protective effects of melatonin on sepsis-induced liver injury and dysregulation of gluconeogenesis in rats through activating SIRT1/STAT3 pathway. Biomed Pharmacother. 2019;117:109150. doi:10.1016/j.biopha.2019.109150

759. Camerer E, Rottingen JA, Gjernes E, et al. Coagulation factors VIIa and Xa induce cell signaling leading to up-regulation of the egr-1 gene. J Biol Chem. 1999;274(45):32225–32233. doi:10.1074/jbc.274.45.32225

760. Xu S, Pan X, Mao L, Pan H, Xu W, Hu Y, Yu X, Chen Z, Qian S, Ye Y, et al. Phospho-Tyr705 of STAT3 is a therapeutic target for sepsis through regulating inflammation and coagulation. Cell Commun Signal. 2020;18(1):104. doi:10.1186/s12964-020-00603-z

761. Hou P, Zhao M, He W, He H, Wang H. Cellular microRNA bta-miR-2361 inhibits bovine herpesvirus 1 replication by directly targeting EGR1 gene. Vet Microbiol. 2019;233:174–183. doi:10.1016/j.vetmic.2019.05.004

762. Carpio VH, Aussenac F, Puebla-Clark L, Wilson KD, Villarino AV, Dent AL, Stephens R. T Helper Plasticity Is Orchestrated by STAT3, Bcl6, and Blimp-1 Balancing Pathology and Protection in Malaria. iScience. 2020;23(7):101310. doi:10.1016/j.isci.2020.101310

763. Németh Á, Mózes MM, Calvier L, Hansmann G, Kökény G. The PPARγ agonist pioglitazone prevents TGF-β induced renal fibrosis by repressing EGR-1 and STAT3. BMC Nephrol. 2019;20(1):245. doi:10.1186/s12882-019-1431-x

764. Pan J, Shi M, Guo F, Ma L, Fu P. Pharmacologic inhibiting STAT3 delays the progression of kidney fibrosis in hyperuricemia-induced chronic kidney disease. Life Sci. 2021;285:119946. doi:10.1016/j.lfs.2021.119946

